# Convergent genome streamlining accompanies independent miniaturization in the world’s smallest fishes

**DOI:** 10.64898/2026.04.20.719654

**Authors:** Hiranya Sudasinghe, Michael Matschiner, Ralf Britz, Kevin W. Conway, Heok Hui Tan, Walter Salzburger, Catherine L. Peichel, Lukas Rüber

## Abstract

Miniaturization, the reduction of adult body size to an extreme degree, has evolved repeatedly across vertebrates. Yet its genomic underpinnings remain poorly understood. Cypriniformes, the most species-rich order of freshwater fishes, contains multiple miniaturized lineages that have evolved contrasting developmental processes. Proportioned dwarfs are tiny-bodied but otherwise morphologically similar to larger relatives, while progenetic miniatures exhibit developmental truncation thus retaining larval-like anatomical features into adulthood. Using a new time-calibrated phylogeny of 309 cypriniform species and comparative genomic analyses of 33 high-quality genome assemblies, we investigated the evolutionary history and genomic correlates of miniaturization across this order. Ancestral state reconstruction revealed multiple independent origins of both miniature types, with transitions predominantly unidirectional and non-randomly distributed across the phylogeny. The origins of the two types of miniatures differed in their timing. Progenetic miniatures arose predominantly as early as the Eocene while proportioned dwarfs arose mainly within the Miocene period. Genome size variation across Cypriniformes has been overwhelmingly driven by polyploidy. However, progenetic miniatures but not proportioned dwarfs showed consistent genome size reduction. Comparative genomic analyses revealed that all three independently-evolved progenetic miniature lineages share convergent signatures of repeat loss alongside genome-wide intron shortening, patterns absent in proportioned dwarfs. Our study provides the broadest evidence to date that progenetic miniaturization, despite independent origins, is underpinned by predictable structural genomic changes, revealing a fundamental link between developmental truncation and genome architecture in vertebrates.

## Introduction

Miniaturization, the evolution of extremely small adult body size, offers a natural experiment for understanding how developmental, morphological, and genomic systems reorganize under strong selection (Hanken & Wake, 1993). Across animals, miniaturization has repeatedly evolved alongside heterochronic shifts in developmental timing, often resulting in paedomorphosis: the retention of juvenile traits in reproductively mature adults (Gould, 1977; Hanken & Wake, 1993). Paedomorphosis occurs via two mechanisms: neoteny, where somatic development slows relative to reproductive maturation, and progenesis, where reproductive maturation accelerates relative to somatic growth (Gould, 1977). While neoteny is extensively documented in vertebrates (Bufill et al., 2011; Gould, 1977; Skulachev et al., 2017), progenesis, despite its strong association with miniaturization (Britz & Conway, 2016) remains less well studied. Progenesis, however, has been implicated in the repeated evolution of miniaturization in invertebrates, including interstitial meiofauna and, parasitic lineages (Gould, 1977; Lefebvre & Poulin, 2005; Okamura et al., 2015; Rundell & Leander, 2010; Zemann et al., 2023).

Among fishes, Cypriniformes is the most species-rich order of freshwater fishes with over 5,000 described species. This taxon provides an ideal system for studying miniaturization because it contains multiple independently derived miniaturized lineages with contrasting anatomical outcomes (Rüber et al., 2007). Miniaturized fishes are conventionally defined as species reaching sexual maturity at or below a standard length of 26 mm while often exhibiting paedomorphic features compared to larger relatives (Weitzman & Vari, 1988). Within Cypriniformes, two distinct morphological outcomes have been recognized: proportioned dwarfs, which are miniaturized but morphologically similar to larger relatives, and progenetic miniatures, which exhibit developmental truncation and retain larval-like anatomical features into adulthood (Britz et al., 2009, 2014; Britz & Conway, 2009, 2016; Rüber et al., 2007). Four genera – *Paedocypris*, *Danionella*, *Sundadanio*, and *Fangfangia* – exhibit extreme progenesis, with *Paedocypris progenetica* representing the world’s smallest fish. It reaches sexual maturity at just under 8 mm standard length (Kottelat et al., 2006). These progenetic miniatures display striking, sexually dimorphic, morphological novelties not developed in proportioned dwarfs and non-miniature cypriniform fishes (Britz et al., 2011; Britz & Conway, 2009, 2016; Conway & Britz, 2007; Kottelat et al., 2006). For example, the pectoral, and pelvic girdles of male *Paedocypris* is highly modified and is hypothesized as a clasping organ (Britz & Conway, 2009; Kottelat et al., 2006). The recurrent evolution of miniaturization and the presence or absence of novelties across progenetic miniature, and proportioned dwarf cypriniform taxa, provides a powerful comparative framework for investigating the evolutionary consequences of extreme body-size reduction.

Despite extensive morphological characterization, the genomic architecture underlying miniaturization in cypriniforms remains poorly understood. Recent work on *Paedocypris* revealed genomes (0.41–0.43 Gb) approximately one-third the size of zebrafish genomes (∼1.5 Gb), characterized by intron shortening, and reduced repetitive content (Malmstrøm et al., 2018). Whether this pattern is generally present across miniaturized lineages, however, remains unknown. Genome size reduction may reflect selective pressures related to development rate, metabolic demands, and cell-size constraints in extremely small organisms (Gregory, 2005a). Here, we make use of a new phylogenomic framework spanning all cypriniform families, including all known progenetic miniatures and most proportioned dwarfs (Sudasinghe, Britz, et al., 2026), to reconstruct a fossil-calibrated temporal framework for Cypriniformes and investigate the evolutionary history of miniaturization across this order. This is complemented by newly generated reference genome assemblies for representatives of both types of miniatures (Sudasinghe, Liu, et al., 2026). Together, these resources enable phylogenetic comparative analyses testing for correlations between body size and genome architecture across evolutionary time. Specifically, we test: (1) whether the frequency and phylogenetic distribution of independent miniaturization origins differ between the two types of miniatures; (2) whether the timing of these events reveals temporal clustering; (3) whether independent miniaturization events consistently associate with genome size reduction; and (4) whether shared genomic mechanisms, repeat loss, and intron shortening underlie convergent genome streamlining. Our findings demonstrate that convergent genome streamlining through repeat loss and intron shortening is a predictable genomic consequence of progenetic miniaturization but not of proportioned dwarfism, revealing a fundamental genomic distinction between the two types of miniatures.

## Materials and methods

### Molecular dating of the Cypriniformes phylogeny

In our previous work, we presented a phylogenomic hypothesis for Cypriniformes with complete family-level representation and broad genus-level coverage, comprising 335 species (316 cypriniform species) that included approximately 30% of all described cypriniform genera (Sudasinghe, Britz, et al., 2026). That study analyzed multiple genome-wide marker datasets using both concatenation-based and coalescent-aware phylogenetic approaches. In the present study, we extend this phylogenomic framework by inferring a molecular dated phylogeny of Cypriniformes, providing a temporal context for lineage diversification and a necessary foundation for downstream phylogenetic comparative analyses of miniaturization.

As a basis for molecular dating, we selected the topology of the ASTRAL-IV species tree inferred from the combined first and second codon positions of the Benchmarking Universal Single-Copy Orthologs (BUSCO.NT.C12) and ultraconserved element (UCE) nucleotide dataset in Sudasinghe, Britz et al. (2026). This dataset comprises 1,167 loci with a total aligned length of 541,983 bp. We chose this topology because it showed high overall congruence across alternative phylogenomic inference methods in the original study.

Molecular dating was performed under a Bayesian framework using BEAST v2.7.7 (Bouckaert et al., 2019). Given the substantial computational demands of molecular dating with large phylogenomic datasets, we reduced the alignment to a subset of 120 loci (55,332 bp) selected based on completeness across taxa from the original 1,167-locus dataset. The ASTRAL-IV species tree inferred from the full BUSCO.NT.C12 and UCE dataset was provided as a starting tree, and its topology was fixed by removing all topological operators from the XML configuration file (Drummond et al., 2012).

We implemented a Birth-Death model (Gernhard, 2008) as the tree prior, and a Gamma model of among site rate variation with four rate categories, coupled with the Optimized Relaxed Clock (ORC) model (Douglas et al., 2022) as the clock model. Fossil calibrations were selected following a review of the published Ostariophysi fossil record, and only fossils considered to be undisputed and convincing representatives of their respective clades were included (Supplementary Information 1). Ingroup fossil calibrations comprised †*Paleogobio zhongyuanensis* as a stem representative of Cyprinoidei (minimum age 47.0 million years ago (Ma)), †*Cobitis nanningensis* as a stem representative of Cobitidae (minimum age 28.1 Ma), †*Amyzon aggregatum* and †*Wilsonium brevipinne* as a stem representatives of Catostomoidei (minimum age 47.8 Ma). Outgroup calibrations included †*Astephus antiquus* constraining the split between *Ictalurus furcatus* and *Cranoglanis bouderius* (minimum age 46.0 Ma), and †*Rubiesichthys gregalis* as a stem representative of Gonorynchiformes (minimum age 132.6 Ma). All fossil calibrations were used exclusively to specify minimum ages for the respective clades, by applying uniform prior distributions ranging from the fossil age to an arbitrarily selected, unrealistically old upper end of 250.0 Ma. Additionally, the crown age of Cypriniformes was constrained according to the timeline of Hughes et al. (2018), applying a log-normal distribution prior with a mean (in real space) of 193.5 Ma and a standard deviation of 0.04 (2.5% quantile: 179 Ma; 97.5% quantile: 209 Ma).

We conducted eight independent Markov-chain Monte Carlo (MCMC) runs, each for at least 50 million generations, including a burn-in of 30%, and sampled parameters and trees every 10,000 generations. Convergence and mixing were assessed by examining effective sample size (ESS) values in Tracer v.1.7.2 (Rambaut et al., 2018), with all parameters exceeding an ESS of 200 in the combined run and all ESS values of individual replicates were above 100 or close to 100. The posterior tree distribution (after removing the burn-in proportions) from each run were combined using LogCombiner, and summarized using TreeAnnotator to generate a maximum clade credibility (MCC) tree with median node heights (Heled & Bouckaert, 2013).

To contextualize our divergence time estimates and assess their congruence with the existing literature, we compared key inferred node ages across Cypriniformes with those reported in previously published molecular dating studies of Cypriniformes (Supplementary Table 1).

### K-mer based genome size estimation

The phylogenomic dataset of Sudasinghe, Britz et al. (2026) comprises 370 genome assemblies representing Cypriniformes, including 95 publicly available genome assemblies and 275 newly assembled genomic resources. Of the newly assembled genomes, 253 were Illumina short-read assemblies Sudasinghe, Britz et al. (2026), four were high-quality reference genomes generated using long-read sequencing (Sudasinghe, Liu, et al., 2026), and the remaining 18 consisted of publicly available raw Illumina short-read sequencing datasets downloaded from the NCBI Sequence Read Archive (Sudasinghe, Britz, et al., 2026). We used these 275 datasets for k-mer– based genome profiling and genome size estimation in the present study.

K-mer profiling and genome characterization were conducted using KMC v3.0 (Kokot et al., 2017) for Illumina short-read datasets and Meryl v.1.4.1 (Rhie et al., 2020) for PacBio long-read datasets. For all datasets, k-mer spectra were generated using both k = 21 and k = 31 to evaluate the robustness of genome size estimates between the two k-mer lengths. GenomeScope 2.0 (Ranallo-Benavidez et al., 2020) was then used to infer genome size, heterozygosity, and repeat content from the k-mer frequency distributions.

During GenomeScope 2.0 analyses, diploid species were modelled using ploidy = 2, whereas known polyploid cypriniform taxa were modelled using their reported ploidy levels (p = 4 or p = 6). Ploidy levels were assigned based on published literature. For a subset of samples, GenomeScope 2.0 failed to converge on the correct haploid (1n) k-mer coverage peak when using default parameters. In these cases, genome profiles were visually inspected, and a biologically informed coverage prior (-l) was manually specified to guide model convergence (Jenike et al., 2025). Resulting estimates were re-evaluated to ensure consistency with expected k-mer distributions and biologically plausible genome characteristics.

Eleven samples exhibited poor model fit and elevated read error rates during genome profiling, and the taxonomic identities of an additional three samples were considered uncertain based on Sudasinghe, Britz et al. (2026). These 14 samples (Supplementary Table 2) were excluded from all downstream analyses. The final dataset therefore comprised 261 genomes, providing complete family-level coverage and extensive genus-level representation across Cypriniformes (Supplementary Table 3; Supplementary Figure 1a).

GenomeScope 2.0 reports genome size estimates as ranges. Therefore, for each genome, mean genome size estimates for k = 21 and k = 31 were calculated by averaging the minimum and maximum haploid genome length estimates returned by GenomeScope 2.0. Similarly, GenomeScope 2.0 reports repeat content estimates as ranges. The proportion of repeat content in the genome was estimated by dividing the minimum repeat length by the minimum haploid genome length and the maximum repeat length by the maximum haploid genome length. Then mean repeat content for each k-mer size was calculated as the average of these two values. Genome size estimates derived from k = 21 and k = 31 were strongly concordant for all species (Supplementary Figure 1b). Consequently, we used the genome size estimates derived from k = 31, which provided the most complete dataset, for all downstream comparative analyses.

Genome size variation within each cypriniform family was quantified as fold variation, calculated as the ratio of the maximum to minimum genome size estimates observed within each family. To explore potential correlates of genome size evolution, we evaluated the relationships between genome size, repeat content, and genomic GC content. GC content estimates were obtained from the corresponding genome assemblies (Sudasinghe, Britz, et al., 2026; Sudasinghe, Liu, et al., 2026).

To assess the concordance between k-mer–based genome size estimates and traditional cytometric measurements, we compiled all available genome size data for Cypriniformes from the Animal Genome Size Database (Gregory, 2025), accessed on 11 November 2024. This dataset was manually curated to match the taxonomic framework used in our study and filtered to retain only genera represented in our genomic dataset. Several records with questionable or ambiguous genome size estimates were excluded, resulting in a curated dataset of 216 cytometric genome size estimates spanning the cypriniform families Gyrinocheilidae, Catostomidae, Balitoridae, Botiidae, Cobitidae, Gastromyzontidae, Nemacheilidae, Acheilognathidae, Cyprinidae, Danionidae, Gobionidae, Leuciscidae, Tincidae, and Xenocyprididae (Supplementary Table 4). For both the k-mer–based and cytometric datasets, the mean genome size was then calculated at the genus level. Genome size estimates were log-transformed prior to comparisons, and correlations between datasets were assessed using Pearson correlation tests implemented in R. Statistical significance was evaluated at p = 0.05.

K-mer-based genome size estimates showed a strong and significant positive correlation with cytometric genome size measurements compiled from the Animal Genome Size Database (r = 0.791, p < 0.0001; Supplementary Figure 2a), confirming that our k-mer-based estimates are consistent with traditional flow cytometric and Feulgen densitometry measurements and are suitable for broad comparative analyses (Elliott & Gregory, 2015).

### Phylogenetic comparative analyses of miniaturization

Within Cypriniformes, two distinct morphological outcomes of miniaturization have been recognized: proportioned dwarfs, which are miniaturized but retain an overall anatomical structure similar to that of larger relatives, and progenetic miniatures, which have undergone profound developmental truncation and retain larval-like anatomical features into adulthood (Britz et al., 2014; Britz & Conway, 2009; Rüber et al., 2007). Progenetic miniaturization within Cypriniformes is confined to all included species of the four genera *Paedocypris*, *Danionella*, *Sundadanio*, and *Fangfangia*.

To test whether miniaturization evolved multiple times independently within Cypriniformes and to evaluate whether these events clustered temporally and/or reveal phylogenetic patterning, we performed ancestral character state reconstruction of type of miniaturization on the time-calibrated phylogeny inferred in the present study (see Molecular dating section). The MCC tree was first pruned to remove outgroup taxa and seven additional cypriniform species that exhibited poor model fit during k-mer genome profiling. The resulting pruned tree comprised 309 cypriniform species, of which 12 are progenetic miniatures, 26 are proportioned dwarfs, and 271 are non-miniature species (Supplementary Table 5). This sampling encompassed all four recognized genera of progenetic miniatures and nearly all known genera of cypriniform proportioned dwarfs (Supplementary Table 6).

Miniaturization status was coded as a discrete trait with three states: non-miniature, proportioned dwarf, and progenetic miniature. Ancestral state reconstruction was performed using stochastic character mapping implemented in the make.simmap function of the R package phytools (Revell, 2012). We specified an equal-rates (ER) model of state transitions and generated 1,000 stochastic maps. Results were then summarized and visualized by averaging posterior probabilities of ancestral states across all 1,000 simulations.

To evaluate whether genome size reduction is a lineage-specific feature of *Paedocypris* or a genomic correlate of progenetic miniaturization, or a generality of miniaturization across Cypriniformes, we mapped genome size and body size as continuous traits onto the pruned 309-species time-calibrated phylogeny. Genome sizes for species in our k-mer profiling dataset were obtained from the GenomeScope 2.0 analyses described above (k = 31); for species represented by publicly available genome assemblies not included in the k-mer profiling dataset, genome sizes were obtained from the corresponding NCBI genome pages. Maximum reported standard length was used as a proxy for body size and was compiled from published literature for each species. Both traits were log10 transformed prior to all analyses.

Ancestral states for log10 transformed genome size and log10 transformed standard length were inferred using maximum likelihood ancestral character state reconstruction via the function fastAnc in the phytools package (Revell, 2012), and results were visualized using the package’s contmap function to produce continuous-trait phylogenetic heat maps.

Before conducting formal comparative tests, we quantified phylogenetic signal for both genome size and body size to assess the degree to which trait variation reflects shared evolutionary history. Two complementary metrics were calculated using the phylosig function in phytools: Pagel’s λ (Pagel, 1999) and Blomberg’s K (Blomberg et al., 2003). Pagel’s λ ranges from 0 to 1, where λ = 0 indicates that trait variation is independent of phylogenetic relatedness and λ = 1 indicates evolution consistent with Brownian motion. Statistical significance of λ was assessed using likelihood ratio tests against the null hypothesis of λ = 0. Blomberg’s K quantifies the strength of phylogenetic signal relative to Brownian motion expectations, with K < 1 indicating weaker structuring than expected under Brownian motion and K > 1 indicating stronger structuring. Significance of K was evaluated by comparing observed values against null distributions generated by randomly permuting trait values across the phylogeny tips (1,000 permutations).

To characterize the mode of evolution underlying genome size and body size variation across Cypriniformes, we compared the fit of four evolutionary models using the fitContinuous function implemented in the R package geiger (Harmon et al., 2008). For each trait, we fitted a Brownian motion (BM) model, an Ornstein–Uhlenbeck (OU) model, an early burst (EB) model, and a white noise (WN) model (Supplementary Table 7). Model support was compared using Akaike Information Criterion (AIC) scores, with ΔAIC > 2 considered indicative of a meaningful difference in fit (Burnham, & Anderson, 2003). To test whether genome size differed significantly between miniaturized and non-miniaturized lineages while accounting for phylogenetic non-independence among species, we fitted phylogenetic generalized least squares (PGLS) regression models using the phylolm package (Ho & Ané, 2014). PGLS explicitly models the expected covariance structure among species arising from shared ancestry, thereby correcting for the inflated Type I error rates that arise when species are treated as statistically independent observations. Miniaturization status was included as a categorical predictor variable. To evaluate the relative contributions of miniaturization type, ploidy, and body size to genome size variation, we fitted a series of models incorporating these predictors individually and in combination, including models with interaction terms between body size and miniaturization status. Models were fitted separately under two evolutionary process assumptions: a Brownian motion (BM) model, in which trait variance accumulates proportionally to branch lengths via random drift (Supplementary Table 8), and an Ornstein-Uhlenbeck (OU) model, which incorporates stabilizing selection toward a trait optimum and may better capture the constrained evolution of genome size if functional or developmental limits exists between miniature and non-miniature lineages (Supplementary Table 9). Model support was compared using Akaike Information Criterion (AIC) scores, with ΔAIC > 2 considered indicative of a meaningful difference in fit (Supplementary Table 10). To assess whether the inclusion of polyploid species influenced our conclusions, all models were additionally fitted on a dataset restricted to diploid species only. As the results were consistent between the full and restricted datasets, those for the diploid-only dataset are not reported separately.

All comparative analyses were conducted under two complementary grouping schemes: (i) all miniature species combined (progenetic miniatures and proportioned dwarfs) versus non-miniatures, and (ii) progenetic miniatures versus proportioned dwarfs versus non-miniatures. This allowed us to distinguish genomic signatures that are broadly shared across miniaturization types. All analyses were conducted in R version 4.3.3 (R Core Team, 2024).

### Comparative genomics of miniaturization

Malmstrøm et al. (2018) reported two primary mechanisms underlying genome compaction in *Paedocypris*: reduced repetitive element content and intron shortening. However, that study was limited in its phylogenetic scope within Cypriniformes. It included only five non-miniature species representing three families (Danionidae, Cyprinidae, and Leuciscidae), and the two *Paedocypris* genomes analyzed were not assembled to reference-genome level. Consequently, it remained unclear whether reduced repeat content and intron shortening represent general features of miniaturization across Cypriniformes or lineage-specific phenomena.

In the present study, we tested whether these genomic signatures are consistently associated with miniaturization in a broader comparative framework. Specifically, we evaluated whether independent miniature lineages exhibit parallel patterns of genome compaction. We compiled a dataset of 33 high-quality reference genomes with available gene annotations, of which 31 are pseudo-chromosome-level assemblies, representing 11 cypriniform families spanning all four recognized suborders (Supplementary Table 11). This sampling includes four genomes generated in Sudasinghe, Liu, et al. (2026), with the remaining assemblies obtained from NCBI.

The dataset includes four miniature species: *Paedocypris* sp., *Sundadanio atomus*, *Danionella cerebrum*, and *Boraras brigittae*. The first three are progenetic miniatures, whereas *B. brigittae* represents a proportioned dwarf. The remaining 29 genomes correspond to non-miniature cypriniform species. Using this dataset, we investigated patterns of transposable element (TE) content and gene architecture across lineages.

Comparative analyses were conducted under two grouping schemes: (i) all miniature species (combining progenetic miniatures and proportioned dwarfs) versus non-miniatures, and (ii) progenetic miniatures only versus non-miniatures.

### Transposable element (TE) annotation and analysis

All genomes were annotated for TEs using the same pipeline to ensure comparability across species. For each genome, TEs were annotated de novo using Extensive de novo TE Annotator (EDTA) (Ou et al., 2019) to construct species-specific repeat libraries. EDTA was run with the --sensitive 1 option to enable RepeatModeler-based discovery of additional repeat families (Flynn et al., 2020) and with --anno 1 to perform whole-genome TE annotation following library construction. The resulting custom TE libraries were subsequently used to mask each genome assembly using RepeatMasker with the slow search option to maximize sensitivity (Smit et al., 2015).

For each species, the CpG-adjusted Kimura two-parameter (K2P) divergence from the consensus sequence of each repeat family was estimated using the RepeatMasker utility script calcDivergenceFromAlign.pl, applied to the .align files generated during masking (Kimura, 1980). These divergence estimates were used to characterize the genomic landscape of TE insertions and to approximate the relative age distribution of TE activity within each genome. In TE divergence landscapes, peaks correspond to periods of elevated TE accumulation. Low-divergence copies represent more recent insertions and highly diverged copies reflect older, inactive elements (Chalopin et al., 2015; Kapusta et al., 2017; Sotero-Caio et al., 2017). We then tested whether miniaturized lineages exhibited distinct TE age distributions compared to non-miniatures by comparing the proportion of insertions in three divergence classes, used here as proxies for relative insertion age: recent (K2P < 5%), intermediate (K2P within 5–20%), and ancient (K2P > 20%). These thresholds were chosen to broadly capture distinct phases of TE activity, ongoing or recent transposition, historically active periods, and older inactive elements respectively.

The resulting .divsum files were processed using custom R scripts to quantify the genomic proportion occupied by non-repetitive DNA and by major TE and repeat categories, including total repeats, class I (LTR, LINEs, SINEs) repeats, class II (DNA transposons, helitrons) repeats, other repeats (i.e., all repeats except class I and class II), retroelements, total interspersed repeats, LINEs, SINEs, LTR elements, DNA transposons, helitrons, small RNAs, satellites, simple repeats, low-complexity repeats, and unclassified repeats, and to generate TE divergence landscapes. Visualization was performed using the ggplot2 package (Wickham, 2016).

### Comparative analysis of gene architecture

To ensure consistency across annotations, we first extracted the longest isoform for each gene from all genome annotations using the script agat_sp_keep_longest_isoform.pl from AGAT (Dainat, 2022). Proteome completeness and annotation consistency were then assessed using OMArk (Nevers et al., 2025), which evaluates proteome quality by comparing each proteome to an inferred ancestral gene repertoire reconstructed for the focal lineage and reports both completeness and consistency based on lineage-appropriate gene family assignments. OMArk single-copy completeness and consistency metrics did not reveal any systematic bias between annotation sources or between miniature and non-miniature species (Supplementary Figure 3). We focused our gene architecture analyses on overall gene length and intron length distributions, as these metrics are most resilient to differences in annotation methodologies across genome sources. While exon lengths are highly conserved due to functional constraints on protein-coding sequences (Long & Deutsch, 1999; Nevers et al., 2023; Sakharkar et al., 2004), estimates of exon lengths, particularly at gene boundaries, are sensitive to UTR annotation practices, which could vary between NCBI and *de novo* annotations. In contrast, intron lengths exhibit substantial biological variation and are less affected by annotation pipeline differences, as intron boundaries are defined by conserved splice sites (Jeffares et al., 2006; Long & Deutsch, 1999; Sakharkar et al., 2004; Sheth et al., 2006).

From the filtered GFF annotation files, we separately extracted gene and exon coordinates for each genome and processed these datasets in R for downstream analyses. For gene-level analyses, when applicable, features were filtered to retain only protein-coding genes (gbkey = Gene; gene_biotype = protein_coding), and all RNA genes were excluded. To remove biologically implausible models (Long & Deutsch, 1999; Sakharkar et al., 2004), genes shorter than 150 bp were discarded. The longest gene in the dataset (2,462,016 bp), however, was retained as it falls in the known range for the longest genes among vertebrates (Sakharkar et al., 2004). For each species, summary statistics of gene length distributions were then calculated.

Intron coordinates were inferred from the filtered exon coordinates for each gene. Introns shorter than 30 bp or longer than 1,000,000 bp were excluded as biologically implausible (Long & Deutsch, 1999; Sakharkar et al., 2004). For each gene, total intron length was calculated, and introns were further classified by position as first (5′), last (3′), middle, or single introns. Because intron length distributions across cypriniform genomes were bimodal, with a first peak around ∼80 bp, an antimode near ∼300 bp, and a second peak around ∼1,000 bp (Supplementary Figure 4), introns were grouped into two primary size classes: compact introns (≤300 bp) and long introns (>300 bp). Compact introns were further subdivided into ultra-short introns (≤100 bp) and short introns (101–300 bp). We quantified the proportion of compact and long introns, calculated the median number of introns per gene among multi-intron genes, and summarized intron length distributions overall, by size class, by positional category, and by the combination of size class and position.

To evaluate potential biases introduced by differences in annotation sources, we compared length metrics between genomes obtained from NCBI and those generated in Sudasinghe, Liu, et al. (2026). Median gene lengths and intron lengths did not show systematic differences between annotation sources (Supplementary Figure 5–7), confirming that these metrics are comparable across our dataset.

### Phylogenetically informed comparative analyses

To test for associations between miniaturization and genomic traits while accounting for evolutionary relationships among species, we analyzed TE content and gene architecture metrics in a phylogenetic comparative framework. The time-calibrated phylogeny inferred in this study (see Molecular dating section) was pruned to retain only the 33 species represented in the comparative genomics dataset.

Three species with available genome assemblies (*Xyrauchen texanus*, *Paramisgurnus dabryanus*, *Cirrhinus molitorella*) were not sampled in the original 335-species molecular dating analysis. For *X. texanus*, we used *Catostomus commersoni* as a phylogenetic proxy, justified by phylogenetic analyses placing *Xyrauchen* as nested within *Catostomus* rather than as a distinct lineage (Bagley et al., 2018; Bangs et al., 2018). Similarly, *Paramisgurnus dabryanus* and *Misgurnus* do not recover as distinct clades in available phylogenies (Perdices et al., 2012; Yi et al., 2017), supporting the use of *Misgurnus mizolepis* as a proxy. For *Cirrhinus molitorella*, a sister group relationship with *Labeo* (Stout et al., 2016; Yang & Mayden, 2010) justified its replacement by the most phylogenetically proximate *Labeo* species in our dataset, minimizing the divergence introduced by this substitution. Additionally, we replaced several species in the time tree with congeneric species within the same genus to match available genome assemblies: *Garra periyarensis* with *G. rufa*, *Leuciscus leuciscus* with *L. chuanchicus*, *L. idus* with *L. waleckii*, and *Gyrinocheilus* sp. with *G. aymonieri*. The resulting pruned phylogeny with 33 tips retained the divergence times and branch length estimates from the full molecular dating analysis. To infer the evolutionary trajectory of total TE% and genome size, we performed maximum likelihood ancestral character state reconstruction using the function fastAnc implemented in the phytools package.

Before conducting comparative tests, we quantified phylogenetic signal for each genomic trait to assess the extent to which trait variation reflects shared evolutionary history versus independent evolution. The phylogenetic signal was quantified using Pagel’s λ and Blomberg’s K (Supplementary Table 12).

To test whether genomic traits differed significantly between miniaturized and non-miniaturized lineages while controlling for phylogenetic non-independence, we fitted PGLS regression models. For each genomic trait, we fitted separate models (BM model and OU model) with miniaturization status as a categorical predictor variable (Supplementary Table 13–14). Model support was compared using Akaike Information Criterion (AIC) scores, with ΔAIC > 2 considered indicative of a meaningful difference in fit (Supplementary Table 15). Given the limited number of miniaturized species in our dataset (n = 4), we report uncorrected p-values but interpret results conservatively, emphasizing effect sizes and consistency across biologically related metrics.

## Results

### A time-calibrated phylogeny of Cypriniformes

To provide a temporal framework for investigating the macroevolutionary history of miniaturization across Cypriniformes, we inferred a time-calibrated phylogeny using Bayesian divergence time estimation in BEAST 2. Our estimates support an early Cretaceous origin of crown Cypriniformes at approximately 112.0 Ma (95% HPD: 91.6–133.0 Ma; Fig. 1), which is in broad agreement with previously published molecular dating studies (e.g., Betancur-R et al., 2017; Chen et al., 2013; Hughes et al., 2018; Near et al., 2012; Rabosky et al., 2018). The divergence estimates among the three major suborders fall within the late Early to early Late Cretaceous, with Cyprinoidei diverging from a clade combining Cobitoidei and Catostomoidei at 101.1 Ma (95% HPD: 83.2–120.4 Ma), Cobitoidei and Catostomoidei separating from each other at 97.8 Ma (95% HPD: 81.0–117.5 Ma), and the crown ages of Cyprinoidei and Cobitoidei estimated at 88.3 Ma (95% HPD: 71.6–106.6 Ma) and 77.0 Ma (95% HPD: 62.9–92.1 Ma), respectively.

**Figure 1.**
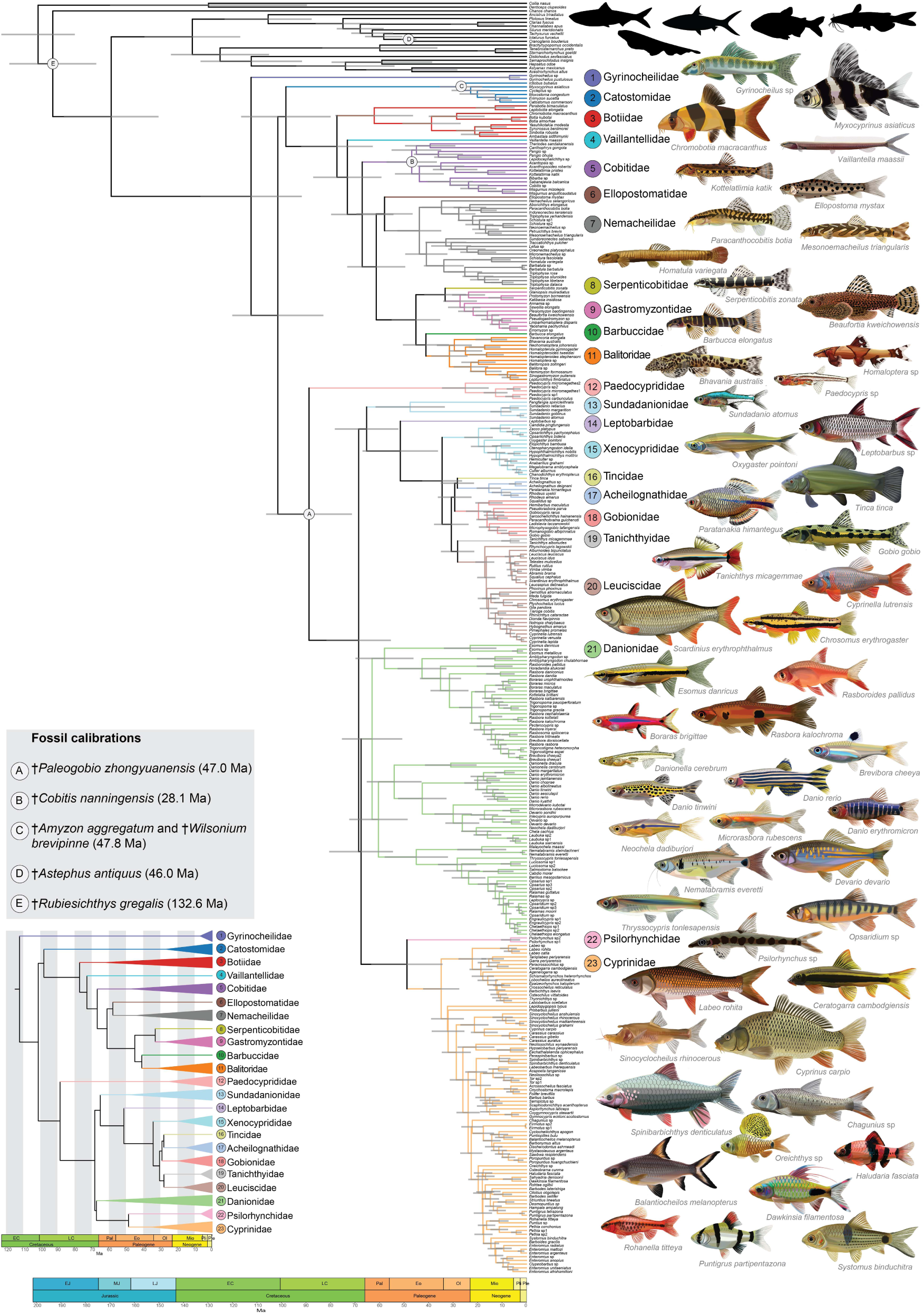
Time-calibrated phylogeny of Cypriniformes inferred from a combined dataset of combined first and second codon positions of BUSCO loci and ultraconserved elements (UCEs; 120 loci, 55,332 bp) under a relaxed clock model in BEAST 2. Branches are colored by family and node bars represent 95% highest posterior density (HPD) intervals in million years (Ma). Labeled nodes indicate fossil calibration points. Inset shows the same phylogeny collapsed to the family level.

At the family level, crown ages broadly fall into two temporal classes. Families with Paleogene origin include Botiidae (61.9 Ma), Danionidae (62.7 Ma), Cobitidae (46.5 Ma), Nemacheilidae (43.5 Ma), Sundadanionidae (47.2 Ma), Cyprinidae (34.7 Ma), Balitoridae (31.2 Ma), and Gastromyzontidae (26.9 Ma). Families with Neogene crown age include Xenocyprididae (19.3 Ma), Gobionidae (19.4 Ma), Leuciscidae (16.9 Ma), and Acheilognathidae (15.2 Ma), with Catostomidae (26.3 Ma) and Paedocyprididae (24.4 Ma) straddling the Oligocene–Miocene boundary. For families represented by only one or two species, stem ages are reported in place of crown ages and ranged from 72.6 Ma (Vaillantellidae; Late Cretaceous) to 27.8 Ma (Tanichthyidae; late Oligocene to early Miocene). Complete stem and crown age estimates with 95% HPD intervals, together with comparisons of key divergence time estimates against previously published molecular dating analyses, are provided in Supplementary Table 1.

### Genome size diversity across Cypriniformes reveals exceptionally compact genomes in loaches

To characterize genome size variation across Cypriniformes and establish a baseline for comparative analyses of miniaturization, we estimated haploid genome sizes for 261 species using k-mer profiling. Genome size showed a strong positive correlation with repeat content (r = 0.687, p < 0.0001; Supplementary Figure 2b), indicating that repeat element accumulation is a primary driver of genome size variation across the order (Gregory, 2005b; Reinar et al., 2023; Smith & Gregory, 2009), and a weak negative correlation with GC content (r = -0.360, p < 0.0001; Supplementary Figure 2c), consistent with patterns reported in other teleosts (Vinogradov, 1998).

Across the 261 species, the median haploid genome size was 752.7 Mb, with a sevenfold span ranging from 347.9 Mb in the balitorid loach *Homalopterula gymnogaster* to 2,509.8 Mb in the polyploid *Cycleptus* sp. (Catostomidae). Genome size within most families was remarkably conserved (intrafamilial fold variation: 1.0–1.5), with only Cobitidae, Cyprinidae, and Danionidae exceeding threefold variation (Supplementary Figure 1c).

A particularly striking finding were the consistently small genome sizes observed across multiple loach (Cobitoidei) families (Supplementary Figure 1b): Balitoridae (347.9–537.5 Mb), Barbuccidae (379.5 Mb), Gastromyzontidae (411.6–474.3 Mb), Vaillantellidae (427.5 Mb), Serpenticobitidae (440.3 Mb), and Ellopostomatidae (452.7 Mb), with several falling below or approaching the genome size of pufferfishes, including the genera *Fugu*, *Takifugu,* and *Tetraodon,* which have the smallest known vertebrate genomes ranging from 346.5–385.0 Mb (Aparicio et al., 2002; Liu et al., 2025).

### Progenetic miniaturization and proportioned dwarfism differ in their frequencies and times of origin

Stochastic character mapping on the time-calibrated phylogeny revealed that miniaturization is a derived condition within Cypriniformes, with the ancestral state at the root inferred as non-miniature (Fig. 2a). Transitions were predominantly unidirectional. Forward transitions from non-miniature to miniature states were substantially more frequent than reversals. Progenetic miniaturization evolved rarely, with a mean of 3.6 (SD = 0.90) independent origins per reconstruction, representing three independent origins across the four recognized genera (*Paedocypris*, *Danionella*, and *Sundadanio* + *Fangfangia*), all within Cyprinoidei. Proportioned dwarfism evolved far more frequently, with a mean of 17.4 (SD = 1.17) independent origins per reconstruction, concentrated predominantly within Danionidae, but with additional origins in Cyprinidae, Tanichthyidae, Cobitidae, Balitoridae, and Barbuccidae. The two miniaturization types differed markedly in their inferred times of origin. Progenetic miniature origins were concentrated in the Eocene (mean 44.9 Ma; Inter Quartile Range (I QR): 37.8–52.3 Ma), whereas proportioned dwarf origins occurred predominantly in the Miocene (mean 10.5 Ma; I QR: 9.2–11.6 Ma).

**Figure 2.**
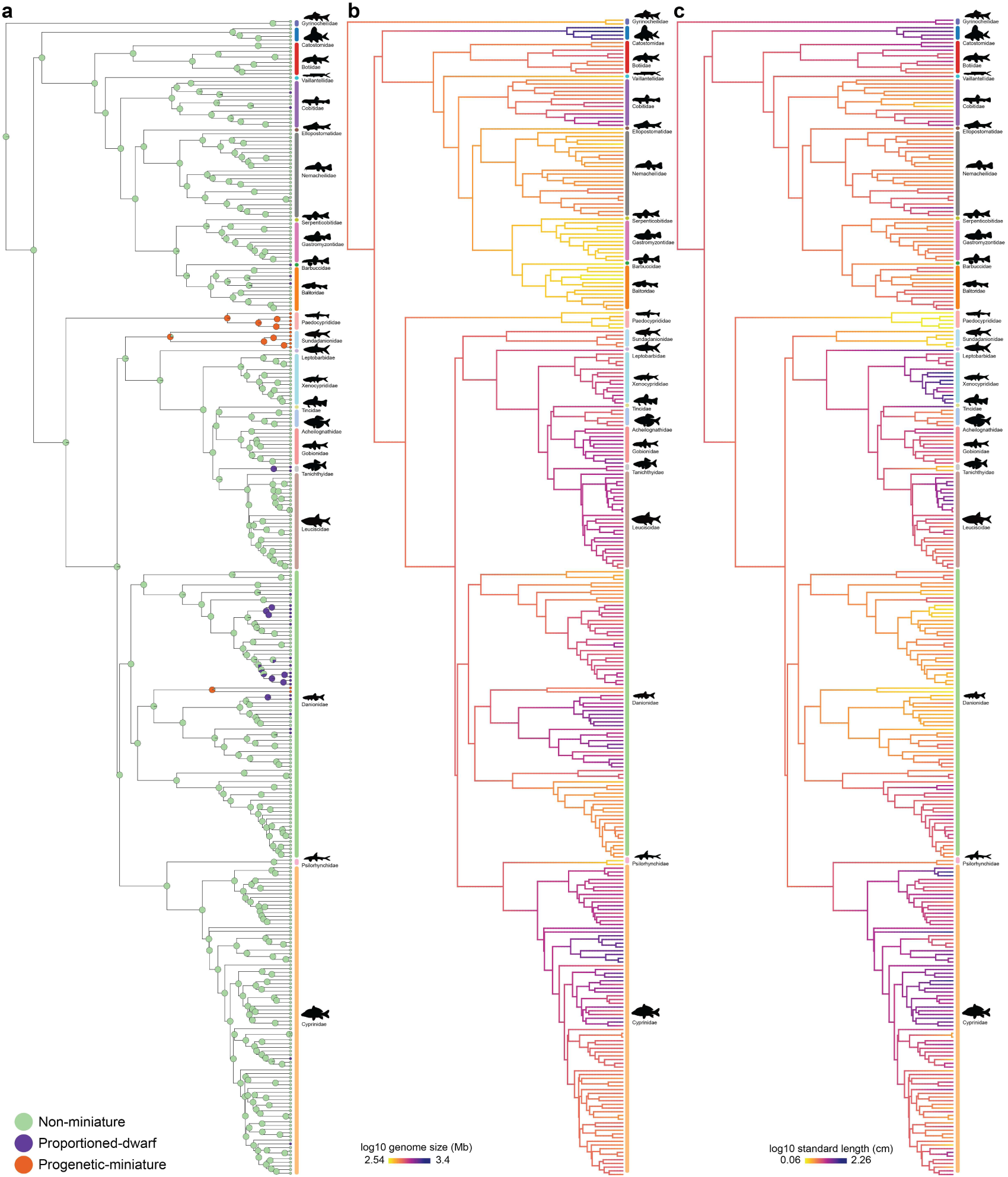
Ancestral state reconstructions of miniaturization, genome size, and body size across Cypriniformes plotted on the pruned time-calibrated phylogeny. **a)** Stochastic character mapping of miniaturization status using an equal-rates (ER) model across 1,000 reconstructions. Pie charts at nodes represent the posterior probability of each miniaturization state. Maximum likelihood ancestral state reconstruction of **b)** log-transformed haploid genome size and **c)** log-transformed standard length as a proxy for body size. In panels **b** and **c**, warmer colors indicate smaller values and cooler colors indicate larger values of the respective log-transformed trait.

### Progenetic miniaturization but not proportioned dwarfism shows a consistent trend toward genome size reduction

Ancestral state reconstruction of genome size revealed recurrent expansions and reductions across the phylogeny (Fig. 2b). Loaches showed consistently smaller genome sizes than Cyprinoidei overall, with the clade comprising Serpenticobitidae, Gastromyzontidae, Barbuccidae, and Balitoridae exhibiting particularly marked genome size reduction. The non-Cobitoidei families Gyrinocheilidae and Psilorhynchidae also exhibited genome size reduction. Ancestral state reconstruction of body size revealed a broadly different pattern, with body size and genome size decoupled in several clades, most notably Danionidae, where most species are small-bodied despite spanning a wide genome size range (Fig. 2c). Both traits showed strong phylogenetic signal (genome size: Pagel’s λ = 1.000, K = 1.110, p = 0.001; body size: Pagel’s λ = 0.946, K = 0.322, p = 0.001), though the contrasting K values indicate that genome size is more conserved within lineages while body size is more evolutionarily variable, consistent with the repeated independent evolution of miniaturization. Not surprisingly, PGLS regression of body size against miniaturization status revealed highly significant reductions in both progenetic miniatures (85% smaller than non-miniatures; p = 2.72 × 10⁻⁷: Fig.3a) and proportioned dwarfs (72% smaller; p = 3.56 × 10⁻¹¹: Fig.3a).

PGLS regression indicated that miniaturization status did not significantly predict genome size in either miniaturization category across the full 309-species dataset (Supplementary Tables 8, 9). Ploidy emerged as the overwhelmingly dominant predictor of genome size variation (β = 0.258, SE = 0.029, p < 2.2 × 10⁻^16^; R² = 0.209), while body size was not a significant predictor after accounting for ploidy and miniaturization status. Proportioned dwarfs showed no meaningful difference in genome size relative to non-miniatures in any model. However, progenetic miniatures showed a consistent negative trend that was directionally consistent across all model structures and independent of ploidy and body size as covariates (β = -0.126, SE = 0.082, t = - 1.533, p = 0.126). Progenetic miniatures had an approximate 25% genome size reduction relative to non-miniatures, which is equivalent to a reduction of roughly 190 Mb for a typical non-miniature cypriniform genome (Fig. 3b).

**Figure 3.**
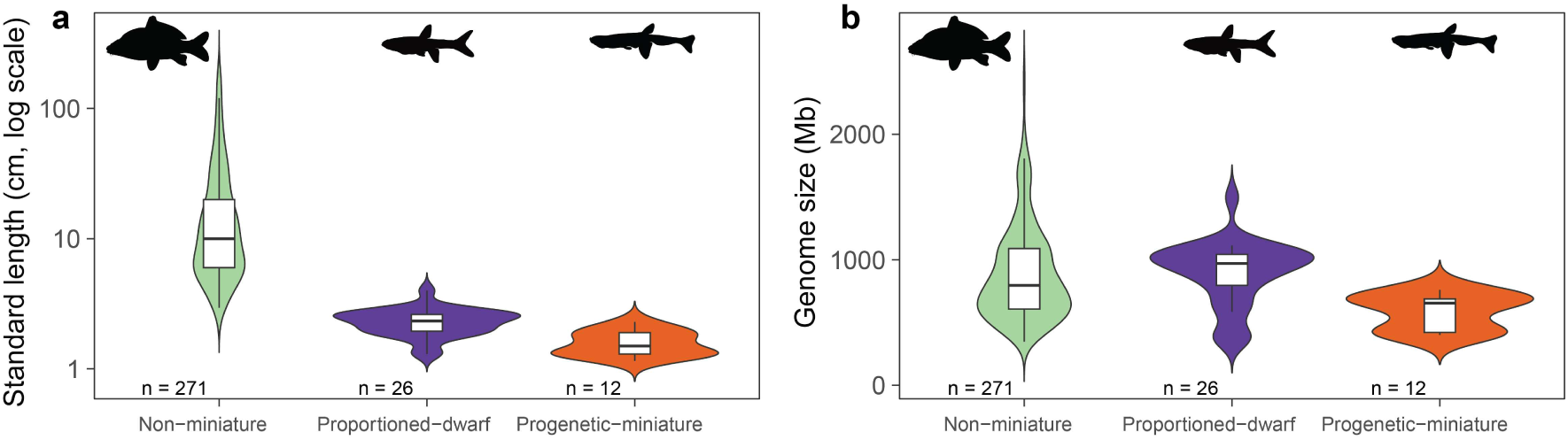
Distribution of **a)** log-transformed standard length as a proxy for body size and **b)** haploid genome size across non-miniature, proportioned dwarf, and progenetic miniature cypriniform species included in the phylogenetic comparative analyses.

### Convergent TE loss underlies genome compaction in progenetic miniatures

To test whether repeat loss represents a shared genomic mechanism of genome compaction across independently evolved miniature lineages, we annotated and compared TE content across a dataset of 33 high-quality cypriniform genome assemblies spanning miniature and non-miniature species. Progenetic miniatures occupied a consistently higher range of non-repetitive DNA content (58.5–64.7%) compared to non-miniatures (34.3–74.4%: (Supplementary Figure 8), and total repeat content showed a strong positive correlation with assembly size across all 33 species (r = 0.951, p < 2.2 × 10⁻^16^; Supplementary Figure 9a). PGLS regression indicated that while total TE content did not differ significantly between all miniatures and non-miniatures combined (β = 9.440, SE = 6.133, p = 0.134), the comparison restricted to progenetic miniatures versus non-miniatures approached significance (β = 13.877, SE = 7.984, p = 0.092), corresponding to approximately 14% more TE content in non-miniature genomes.

Maximum likelihood ancestral state reconstruction illustrated the convergent TE loss across all three independently evolved progenetic miniature lineages (*Paedocypris* sp., *Sundadanio atomus*, and *Danionella cerebrum*), each showing marked reductions in TE content relative to their closest non-miniature relatives (Fig. 4a). In contrast, the proportioned dwarf *Boraras brigittae* showed no comparable reduction, reinforcing the pattern that genome compaction is specifically associated with progenetic miniatures rather than proportioned dwarfs. However, TE loss was not exclusive to progenetic miniatures. Several non-miniature species including *Gyrinocheilus aymonieri*, nemacheilid loaches, and *Puntigrus tetrazona* also exhibited TE loss, consistent with the broader pattern of genome compaction observed across certain cypriniform lineages independently of miniaturization (Fig. 2b, 4a,b).

**Figure 4.**
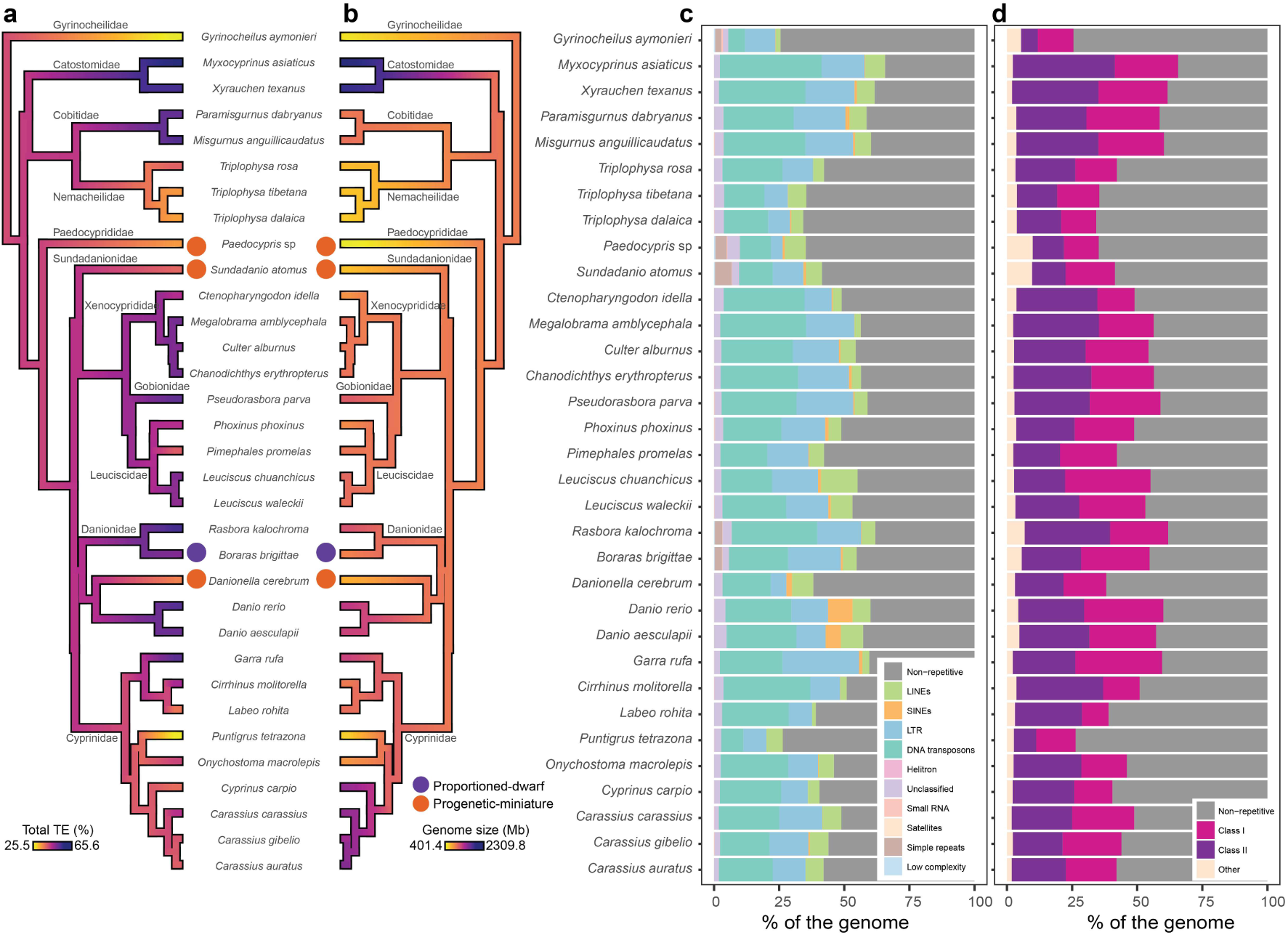
Transposable element (TE) content and genome size variation across miniature and non-miniature cypriniform species in the 33-taxon comparative genomic dataset. Maximum likelihood ancestral state reconstruction of **a)** total TE content as a percentage of the genome and **b)** haploid genome size (Mb) plotted on the 33-taxon time-calibrated phylogeny. In both panels, warmer colors indicate lower values and cooler colors indicate higher values. Distribution of **c)** major TE subclasses and **d)** major repeat classes (Class I, Class II, and other repeats) as a percentage of the genome across miniature and non-miniature cypriniform species.

At the repeat class level, Class II elements were significantly reduced in progenetic miniatures relative to non-miniatures (β = 10.507, SE = 4.876, p = 0.039), while Class I content did not differ significantly (β = 5.300, SE = 3.751, p = 0.168). Within each class, DNA transposons and LTR elements were both significantly reduced in progenetic miniatures (β = 10.554, SE = 4.875, p = 0.038; β = 7.579, SE = 2.984, p = 0.017, respectively), identifying these as the primary drivers of genome compaction in these lineages. Interestingly, other repeat content, driven primarily by simple repeats was significantly enriched in progenetic miniatures relative to non-miniatures (β = -3.207, SE = 1.035, p = 0.004; simple repeats: β = -2.819, SE = 0.701, p = 0.0004; see Fig. 4c,d), suggesting that genome compaction in progenetic miniatures reflects a complex remodelling of repeat composition rather than a uniform depletion of all repetitive elements. Results for all repeat categories are provided in Supplementary Tables 12–13 and Supplementary Figures 10–11.

To determine whether the depletion of TEs in progenetic miniature genomes reflects a reduction in recent transposition activity, a loss of older elements, or both, we examined the relative age distribution of TE insertions across the dataset using CpG-adjusted Kimura two-parameter (K2P) divergence landscapes (Supplementary Figure 12). The three progenetic miniature species, together with the non-miniature species that exhibited TE loss in the ancestral state reconstruction (*Gyrinocheilus aymonieri*, nemacheilid loaches, and *Puntigrus tetrazona*), showed broadly similar TE age distributions relative to the remaining non-miniature species, characterized by low proportions of both recent and intermediate divergence insertions, suggesting a sustained reduction in TE activity across both recent and intermediate timescales rather than a depletion confined to a specific period (Supplementary Figure 12b,c). In contrast, the proportioned dwarf *Boraras brigittae* showed a TE age distribution comparable to non-miniature species, consistent with the absence of overall TE depletion in this lineage.

PGLS comparisons across TE age classes supported this interpretation. Among intermediate divergence insertions (K2P 5–20%), LTR elements were significantly reduced in progenetic miniatures (β = 4.786, SE = 2.033, p = 0.025), while other repeat content was significantly elevated (β = -1.463, SE = 0.557, p = 0.014), with Class II elements and DNA transposons additionally approaching significance. Among recent insertions (K2P < 5%), Class II elements and DNA transposons showed higher levels in non-miniatures approaching significance. In contrast, LINEs were elevated in progenetic miniatures among recent insertions, potentially reflecting ongoing LINE activity in otherwise streamlined genomes. Among ancient insertions (K2P > 20%), LTR elements showed a trend toward reduction in progenetic miniatures (β = 1.937, SE = 1.083, p = 0.084), though no other repeat types differed significantly. Taken together, these results indicate that TE depletion in progenetic miniature genomes is not confined to a single temporal class but reflects a sustained and prolonged reduction in transposition activity spanning both recent and intermediate timescales, consistent with a genome-wide and temporally extended process of repeat loss rather than a recent or episodic purging of insertions. Full results for all TE age class comparisons are provided in Supplementary Tables 12–13 and Supplementary Figures 13–18.

### Intron shortening is a consistent genomic signature across progenetic miniature lineages in Cypriniformes

To test whether intron shortening represents a shared genomic mechanism of genome compaction alongside TE loss, we examined gene length and intron length distributions across the 33-species comparative genomic dataset. Median gene lengths were marginally but non-significantly longer in non-miniature species relative to all miniatures combined (β = 2,433.5, SE = 1,291.6, p = 0.069) and to progenetic miniatures (β = 2,302.6, SE = 1520.5, p = 0.140; Fig. 5), likely reflecting limited statistical power given the small number of progenetic miniature species.

**Figure 5.**
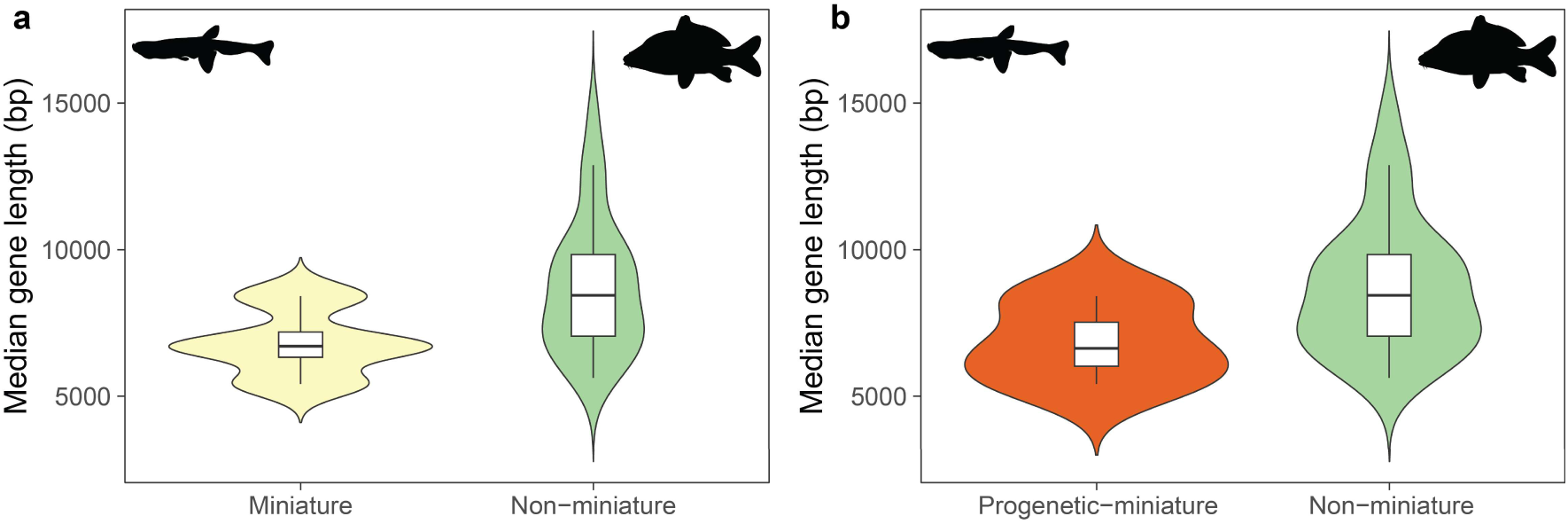
Distribution of median gene length (bp) across **a)** miniature and non-miniature and **b)** progenetic miniature and non-miniature cypriniform species included in the comparative genomic analyses.

The most striking finding emerged from the comparison of intron length distributions by size class. Plotting median long intron length (>300 bp) against median compact intron length (≤300 bp) clearly separated all three progenetic miniature species from the remainder of the dataset (Fig. 6), suggesting convergent intron shortening of compact introns across independently evolved progenetic miniature lineages. The proportioned dwarf *Boraras brigittae* clustered with non-miniature species. Furthermore, the non-miniature species exhibiting TE loss and reduced genome sizes (*Gyrinocheilus aymonieri*, nemacheilid loaches, and *Puntigrus tetrazona*) also clustered with the rest of the non-miniature species (Fig. 6). Notably, non-miniature species with compact genomes showed comparatively shorter long introns relative to other non-miniature species, suggesting that while these lineages undergo some degree of intron shortening, it is concentrated in long introns rather than compact introns (Fig. 6). This suggests that intron shortening of compact introns is specifically associated with progenetic miniaturization and does not accompany TE-mediated genome compaction in non-miniature lineages in Cypriniformes.

**Figure 6.**
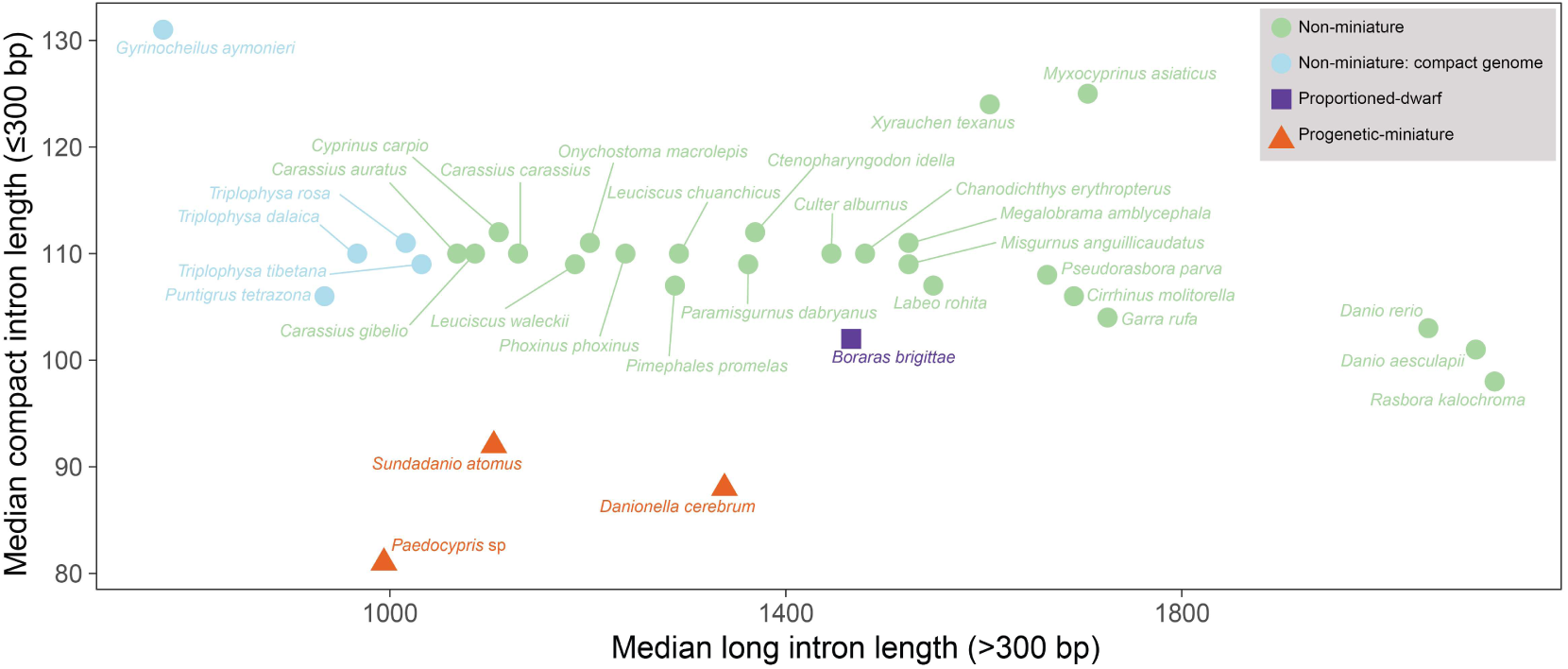
Distribution of median long intron length (>300 bp) versus median compact intron length (≤300 bp) across miniature and non-miniature cypriniform species included in the comparative genomic analyses. Each point represents a species, colored by miniaturization status.

PGLS analyses confirmed that intron length differences between progenetic miniatures and non-miniatures were most pronounced in compact intron size classes. Ultra-short introns (≤100 bp) and compact introns (≤300 bp) were significantly shorter in progenetic miniatures (β = 7.968, SE = 1.064, p = 2.405e-08; β = 20.513, SE = 3.435, p = 1.51e-06 respectively), while long introns (>300 bp) also differed significantly but less markedly (β = 429.04, SE = 201.78, p = 0.042). Intron shortening was not confined to specific positions in the genes. Compact introns showed pronounced differences across first (5′), last (3′), and middle intron positions, indicating a genome-wide rather than position-specific phenomenon, with the exception of single introns which did not differ significantly between groups (Supplementary Tables 12–13; Supplementary Figures 19–22).

In contrast to the pronounced intron length differences, the proportion of compact versus long introns and the median number of introns per gene did not differ significantly between progenetic miniatures and non-miniatures (Supplementary Figures 23–24), providing evidence for genome-wide intron shortening but not intron loss in progenetic miniature lineages.

## Discussion

Miniaturization, the reduction of adult body size to an extreme degree, is a recurrent theme across animal diversity, having evolved independently across numerous invertebrate and vertebrate lineages (Hanken & Wake, 1993). The morphological consequences of miniaturization have been extensively documented (e.g., Glynne & Adams, 2024; Gould, 1977; Hanken & Wake, 1993; Perez-Martinez & Leal, 2021; Polilov, 2015), yet its genomic underpinnings remain poorly investigated, with only a handful of studies addressing this question in invertebrates (e.g., Greenhalgh et al., 2020; Martín-Durán et al., 2020; Xu et al., 2021) and vertebrates (e.g., Decena-Segarra et al., 2020; Malmstrøm et al., 2018; Troyer et al., 2025). Miniaturization is comparatively frequent in teleost fishes (Kottelat & Vidthayanon, 1993; Rüber et al., 2007; Weitzman & Vari, 1988). Within Cypriniformes, the morphological and developmental correlates of miniaturization have been especially well characterized. Progenetic miniatures undergo profound developmental truncation, giving rise to striking morphological novelties not shown in their larger relatives, while proportioned dwarfs retain a normal developmental trajectory despite their reduced body size (e.g., Britz et al., 2009; Britz & Conway, 2009, 2016; Conway et al., 2017, 2021). Here, we build on this foundation to provide one of the most comprehensive investigations to date of the evolutionary history and genomic correlates of miniaturization in a vertebrate order.

### Evolutionary origins and timing of miniaturization

Our time-calibrated phylogeny provides a temporal framework for tracing the evolutionary history of miniaturization across the order. Miniaturization evolved independently multiple times within Cypriniformes, with transitions predominantly unidirectional from non-miniature to miniature states. This unidirectionality is most readily explained for progenetic miniaturization, which involves profound and largely irreversible developmental truncation (Britz & Conway, 2009, 2016; Hanken & Wake, 1993). The frequency of independent origins differed strikingly between the two types: progenetic miniaturization arose rarely, with only three independent origins all within Cyprinoidei, while proportioned dwarfism evolved far more frequently, concentrated predominantly within Danionidae. This non-random phylogenetic distribution suggests that the propensity for miniaturization is phylogenetically structured within Cypriniformes, indicating that certain lineages may possess developmental or ecological preconditions that facilitate body size reduction. The difference in frequency likely reflects the greater developmental constraints on progenetic miniaturization, which requires extreme heterochronic shifts resulting in the simultaneous truncation of multiple organ systems, compared to the simpler body size reduction characterizing proportioned dwarfism, which may face fewer developmental and genetic constraints (Britz & Conway, 2009, 2016; Conway et al., 2021). Furthermore, we document an apparent temporal dissociation between the two types: progenetic miniature origins were concentrated in the Eocene while proportioned dwarfs originated predominantly in the Miocene, suggesting that the two types not only differ morphologically and developmentally but may have originated in response to different macroevolutionary processes operating at different times in cypriniform history.

### Genome size evolution across Cypriniformes

Our cypriniform genome size dataset represents one of the largest comparative genome size resources assembled for any vertebrate group, capturing an exceptional range of genomic diversity within a single order. Genome size within most families was remarkably conserved, consistent with patterns reported across teleost more broadly (Canapa et al., 2015; Gregory, 2005a), with order-wide variation driven predominantly by polyploidy (Blommaert, 2020; Hardie & Hebert, 2004). Against this background, miniaturization status did not significantly predict genome size across the full cypriniform dataset. Proportioned dwarfs showed no meaningful difference in genome size relative to non-miniatures, demonstrating that body size reduction alone does not entail genome compaction. This decoupling of genome size from body size reduction is consistent with the broader C-value enigma, reinforcing that genome size evolution is governed by mechanisms largely independent of organismal size or complexity (Gregory, 2001; Liedtke et al., 2018). Progenetic miniatures, however, showed a directionally consistent ∼25% genome size reduction relative to non-miniatures, robust across all model structures and independent of ploidy and body size. Although this trend did not reach statistical significance, possibly reflecting the small number of independent progenetic miniature lineages against the broad genomic diversity of non-miniatures in our PGLS analyses, its consistency suggests a genuine biological tendency toward genome compaction in progenetic lineages. This extends the genome compaction previously reported in *Paedocypris* (Malmstrøm et al., 2018) to progenetic miniatures more broadly, while confirming that it is not a feature of proportioned dwarfism. This suggests that the mechanisms linking body size reduction and genome evolution differ fundamentally between the two types of miniaturization.

A particularly striking finding were the consistently small genome sizes across multiple loach families, with Serpenticobitidae, Gastromyzontidae, Barbuccidae, and Balitoridae showing particularly marked reductions. Given the monophyly of the clade comprising these four families, this pattern likely reflects a shared ancestral genome compaction event rather than independent reductions within the suborder. The ecological correlates of genome compaction across these lineages are intriguing. Loaches along with Psilorhynchidae (torrent minnows), which also had reduced genome sizes, are predominantly benthic fishes adapted to fast-flowing, well-oxygenated streams (Lujan & Conway, 2015), while progenetic miniatures outside of *Danionella* are confined to the chronically resource-limited, acidic peat swamp forests of Southeast Asia (Conway et al., 2011; Kottelat et al., 2006). Both groups therefore inhabit physiologically demanding environments, consistent with the long-standing hypothesis that metabolically demanding lifestyles select for smaller genomes through the relationship between cell size, nuclear DNA content, and metabolic rate (e.g., Blommaert, 2020; Gardner et al., 2020; Gregory, 2001; Hardie & Hebert, 2003; Kapusta et al., 2017). In amphibians, environmental factors such as habitat temperature and moisture have been shown to influence genome size evolution indirectly through their effects on developmental rate (Liedtke et al., 2018). Whether metabolic demands, developmental rate, or habitat extremes may similarly influence genome size evolution in Cypriniformes represents a promising avenue for future research.

### Genomic signatures of genome streamlining in progenetic miniatures

Genome compaction has been documented across diverse vertebrate and invertebrate lineages irrespective of body size, with compact genomes consistently showing predictable signatures of repeat depletion and structural simplification (e.g., Kelley et al., 2014; Lamichhaney et al., 2021; Liu et al., 2025). Our study asked whether miniaturization in Cypriniformes is associated with such genomic streamlining and whether the same mechanisms operate repeatedly across independently evolved lineages. The answer depends critically on the type of miniaturization: proportioned dwarfism shows no evidence of genome streamlining, while progenetic miniaturization is consistently associated with TE loss and intron shortening across all three independently evolved lineages.

The primary causes of TE depletion were reduction of LTR elements and DNA transposons which are highly active element classes capable of rapid genome expansion (Chalopin et al., 2015; Gao et al., 2016; Piegu et al., 2006; Rubio et al., 2025; Sotero-Caio et al., 2017; Sun & Mueller, 2014). Depletion of these elements suggests either reduced transposition activity, increased elimination of newly inserted elements, or both, consistent with mechanisms proposed for genome compaction in other vertebrates with streamlined genomes (Canapa et al., 2015; Kapusta et al., 2017; Lamichhaney et al., 2021; Liu et al., 2025). Genome compaction in progenetic miniatures involves compositional remodelling rather than uniform repeat depletion, with simple repeats significantly enriched relative to non-miniatures, indicating mechanistically specific streamlining rather than a general suppression of all repetitive elements. TE depletion spanning both recent and intermediate divergence classes further suggests a sustained process operating over an extended evolutionary period in each progenetic lineage rather than a recent episodic event. Notably, TE loss also occurred in several non-miniature species, raising the question of whether shared mechanisms underlie genome compaction across cypriniform lineages independently of miniaturization.

The most compelling evidence for a genomic mechanism associated with progenetic miniaturization came from gene architecture analyses. Intron length distributions clearly separated the three progenetic miniature species from all other species in the dataset, including the proportioned dwarf, and non-miniature species with reduced TE content with small genomes. Intron shortening was consistently associated with progenetic miniaturization and did not accompany TE-mediated genome compaction in non-miniature lineages. This intron shortening was genome-wide, affecting first, last, and middle intron positions, and was most pronounced in compact intron size classes. This is consistent with the observation that teleost introns fall into two functional classes: compact introns containing primarily sequences necessary for efficient splicing, and long introns disproportionately harbouring regulatory elements and conserved sequences maintained across vertebrates (Jakt et al., 2022). The preferential shortening of compact introns suggests that introns most tolerant of length reduction are the primary targets of genome streamlining, while long introns with greater regulatory content show comparatively less reduction. Interestingly, non-miniature species with compact genomes showed a contrasting pattern, with shortening concentrated in long introns rather than compact introns, suggesting that different mechanisms may drive intron shortening in these lineages compared to progenetic miniatures. Whether this reflects active selection for shorter introns in progenetic miniatures imposed by developmental constraints on transcription speed and splicing efficiency (Jeffares et al., 2006; Swinburne & Silver, 2008), or is driven primarily by deletion bias, remains to be determined.

Progenetic miniaturization across Cypriniformes, despite arising independently, is underpinned by convergent genomic signatures of TE loss and intron shortening. Whether proportioned dwarfs exhibit functional genomic differences from non-miniatures despite lacking structural genome compaction remains to be investigated. Gene expression differences have been reported in miniature gobies (Troyer et al., 2025) and whether similar patterns occur in cypriniform miniatures is worth exploring. The goby genus *Schindleria*, and the clupeoid genus *Sundasalanx*, offer additional striking cases of progenetic miniaturization among teleosts (Britz & Conway, 2009; Johnson & Brothers, 1993; Siebert, 1997). They provide independent examples to test whether the genomic signatures identified herein extend beyond Cypriniformes and whether there is a shared genomic basis of progenetic miniaturization in vertebrates. Taken together, our results reveal that it is the mode of developmental reduction, rather than the degree of body size reduction, that determines whether miniaturization leaves a structural genomic signature.

## Acknowledgments

We thank Nadeela Hirimuthugoda (https://nadeela.weebly.com/illustrations.html) for providing the fish illustrations. HS thanks Fabrizia Ronco, Vivien Louppe and Conor Waldock for valuable guidance on phylogenetic comparative methods, Daniel Jeffries and Zuyao Liu for helpful discussions on comparative genomic analyses, and Marius Roesti for discussions on statistical analyses.

## Supplementary Information

**Supplementary Information 1.** Fossil calibrations used in Bayesian divergence time estimation, including taxonomic assignment, minimum age, and justification for inclusion. Excluded calibrations and the reasons for their exclusion are also documented.

## Supplementary Tables

**Supplementary Table 1.** Comparison of key divergence time estimates for major clades inferred in the present study with those reported in previously published molecular dating analyses. Node ages in the present study are reported as median estimates with 95% highest posterior density (HPD) intervals in million years (Ma).

**Supplementary Table 2.** Species excluded from downstream analyses.

**Supplementary Table 3.** Genome size estimates for species included in the k-mer profiling dataset (n = 261).

**Supplementary Table 4.** Cytometric genome size estimates for cypriniform species compiled from the Animal Genome Size Database, accessed on 11 November 2024. Records were curated to retain only genera represented in the k-mer profiling dataset of the present study.

**Supplementary Table 5.** Comparative dataset of 309 cypriniform species used in the phylogenetic comparative analyses of miniaturization.

**Supplementary Table 6.** Miniature fishes within Cypriniformes, comprising progenetic miniatures and proportioned dwarfs, with indication of which species were included in the present study.

**Supplementary Table 7.** Comparison of evolutionary model fit for log-transformed genome size and body size across Cypriniformes. Models were fitted using the fitContinuous function in the R package geiger.

**Supplementary Table 8.** Results of phylogenetic generalized least squares (PGLS) regression analyses for log-transformed genome size and body size across Cypriniformes under a Brownian motion (BM) model of trait evolution.

**Supplementary Table 9.** Results of phylogenetic generalized least squares (PGLS) regression analyses for log-transformed genome size and body size across Cypriniformes under an Ornstein–Uhlenbeck (OU) model of trait evolution.

**Supplementary Table 10.** Comparison of Brownian motion (BM) and Ornstein–Uhlenbeck (OU) model fit within the phylogenetic generalized least squares (PGLS) regression framework for log-transformed genome size and body size across Cypriniformes.

**Supplementary Table 11.** Reference genome assemblies used in the analyses of transposable element content and gene architecture across miniature and non-miniature cypriniform species.

**Supplementary Table 12.** Phylogenetic signal estimates for transposable element content and gene architecture traits across Cypriniformes, reported as Pagel’s λ and Blomberg’s K with associated p values.

**Supplementary Table 13.** Results of phylogenetic generalized least squares (PGLS) regression analyses for transposable element content and gene architecture traits across Cypriniformes under a Brownian motion (BM) model of trait evolution, with miniaturization status as the categorical predictor.

**Supplementary Table 14.** Results of phylogenetic generalized least squares (PGLS) regression analyses for transposable element content and gene architecture traits across Cypriniformes under an Ornstein–Uhlenbeck (OU) model of trait evolution, with miniaturization status as the categorical predictor.

**Supplementary Table 15.** Comparison of Brownian motion (BM) and Ornstein–Uhlenbeck (OU) model fit within the phylogenetic generalized least squares (PGLS) regression framework for transposable element content and gene architecture traits across Cypriniformes.

---

**Supplementary Figure 1.**
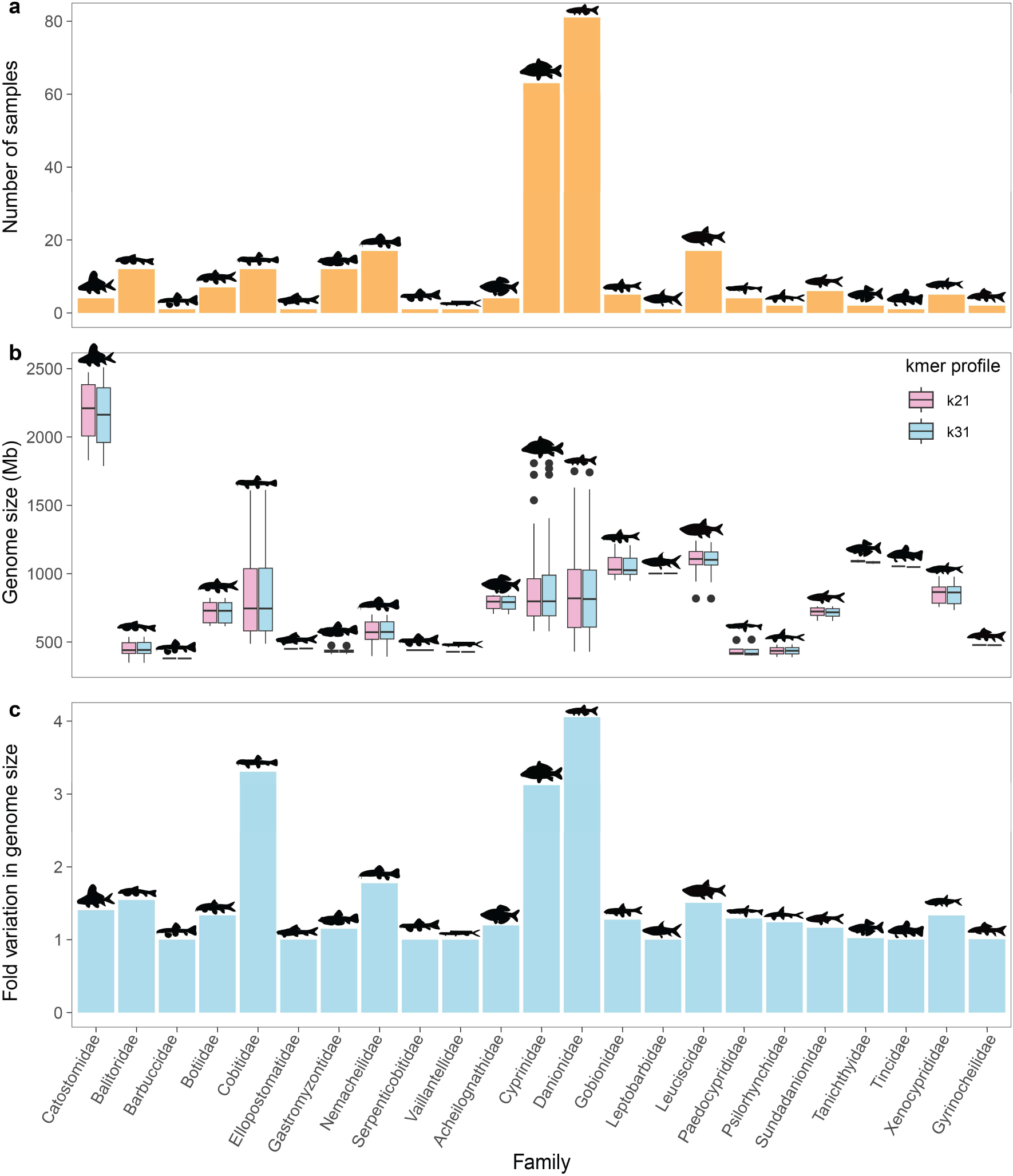
K-mer-based genome size estimation across Cypriniformes for 261 samples. **a)** Number of species sampled per family included in the k-mer profiling dataset. **b)** Comparison of haploid genome size estimates derived from k = 21 and k = 31 across cypriniform families. **c)** Intrafamilial fold variation in genome size (based on k = 31 estimates) across cypriniform families.

**Supplementary Figure 2.**
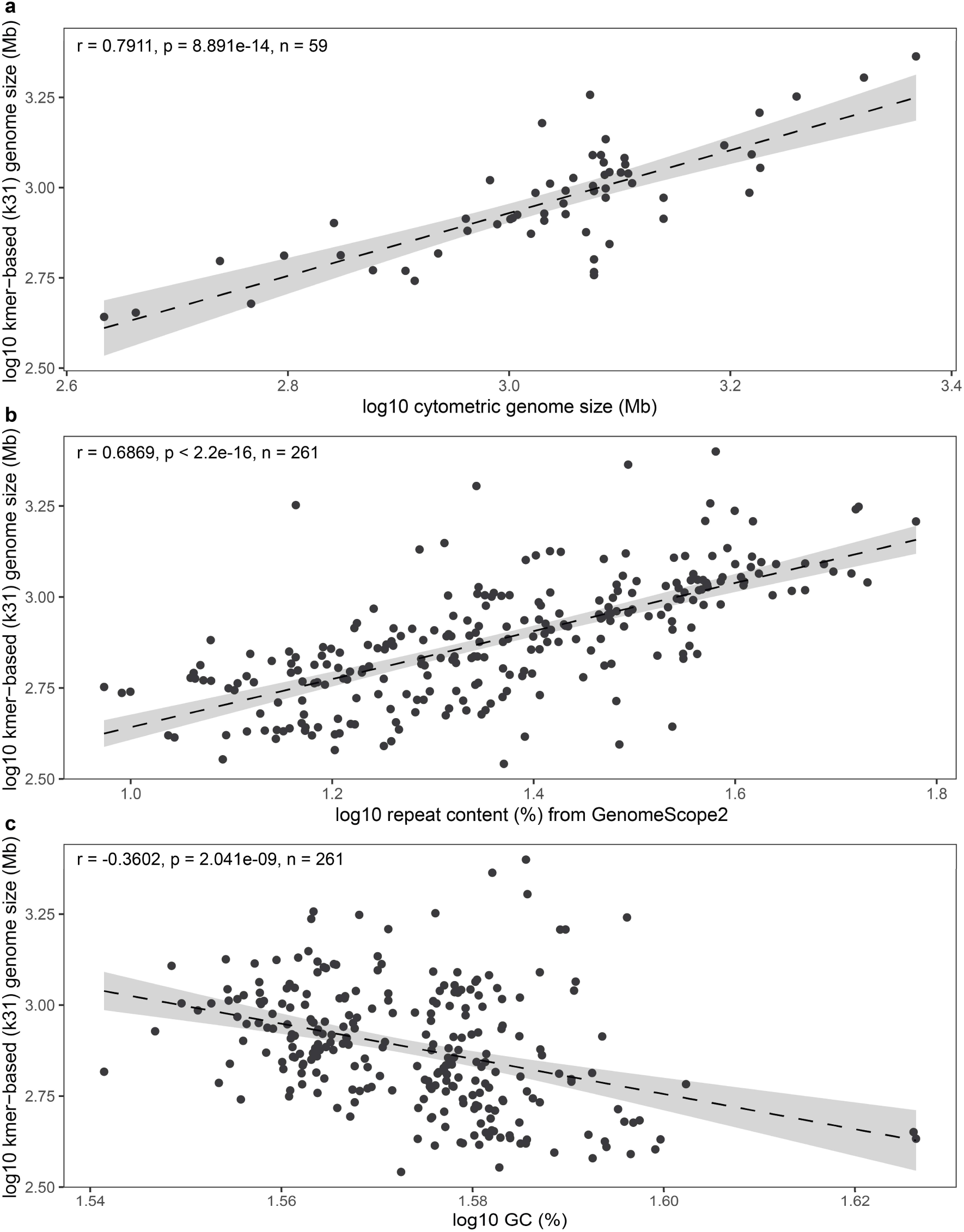
Correlates of genome size variation across Cypriniformes based on k = 31 estimates. **a)** Correlation between cytometric genome size estimates compiled from the Animal Genome Size Database and k-mer-based genome size estimates from the present study, with each data point representing a genus-level mean. **b)** Correlation between repeat content estimated from GenomeScope 2.0 k-mer profiling and haploid genome size. **c)** Correlation between genomic GC content of the genome assembly and haploid genome size. Pearson correlation coefficients and associated p values are indicated in each panel.

**Supplementary Figure 3.**
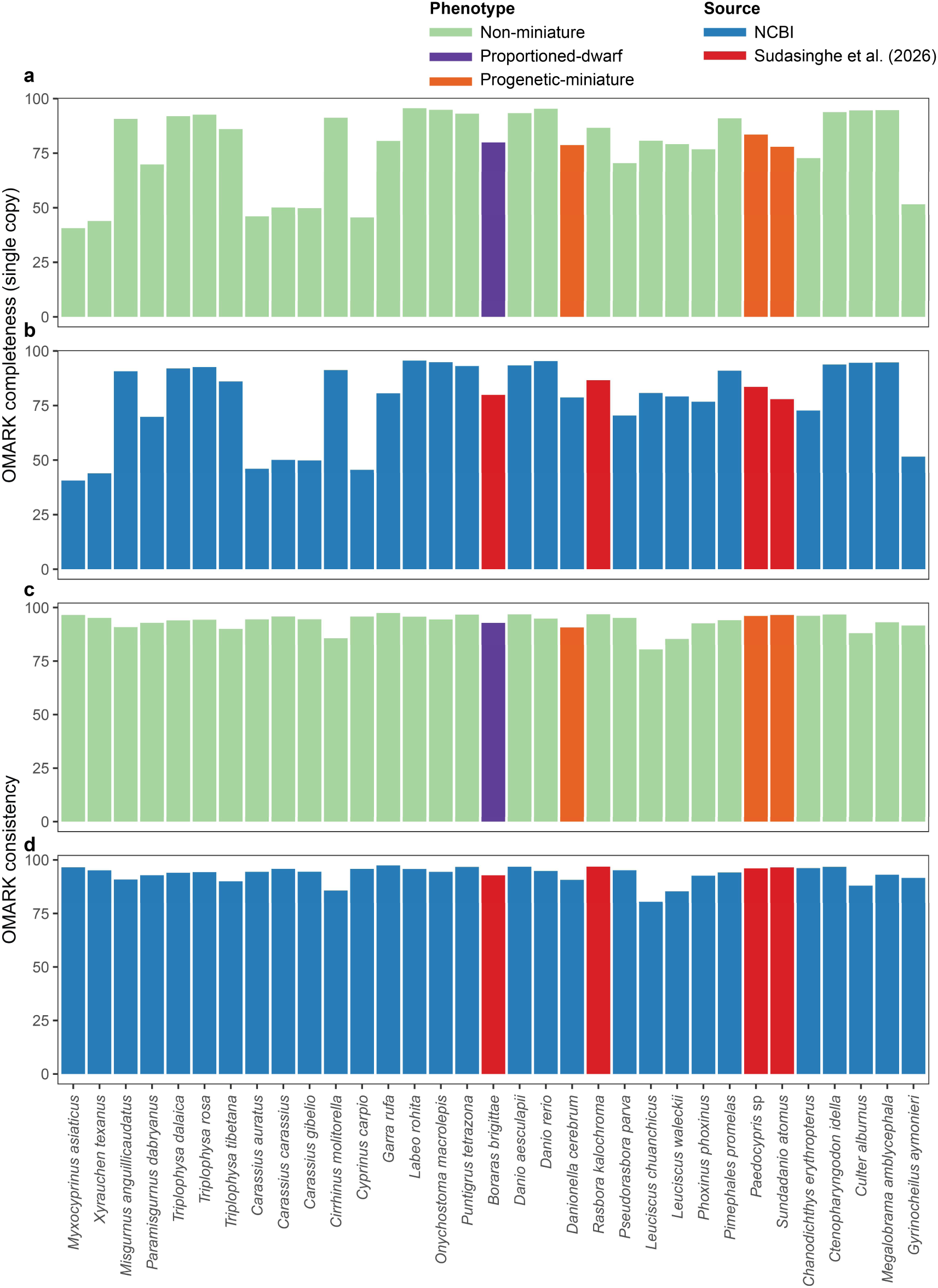
Assessment of proteome quality and annotation consistency using OMArk for the 33-species dataset used in comparative genomic analyses of gene architecture. Single-copy completeness scores grouped by **a)** miniaturization category and **b)** annotation source. OMArk consistency scores grouped by **c)** miniaturization category and **d)** annotation source.

**Supplementary Figure 4.**
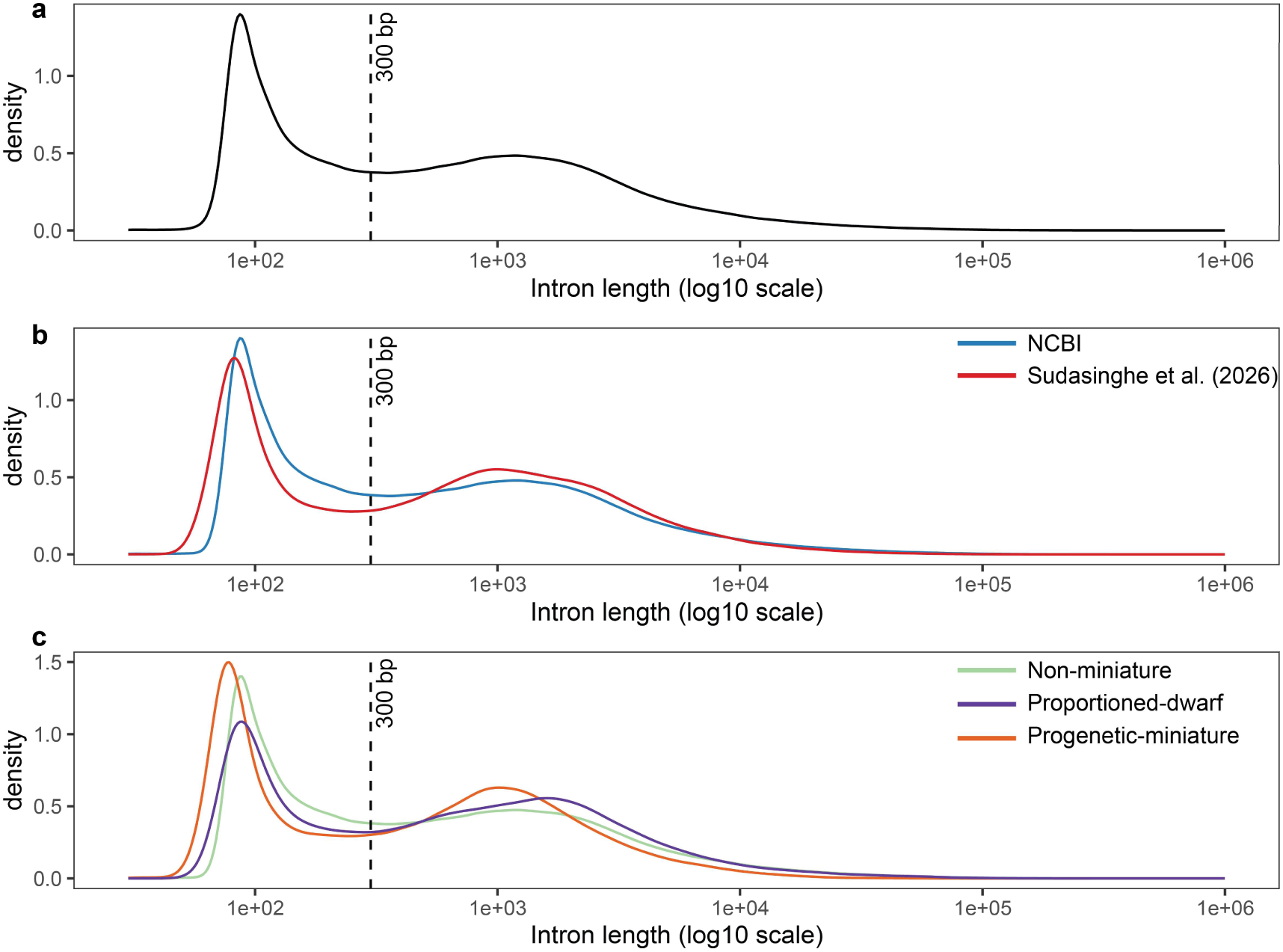
Intron length distributions across the 33-species dataset used in comparative genomic analyses of gene architecture, illustrating the bimodal distribution. **a)** Overall distribution across all species. **b)** Distributions grouped by annotation source. **c)** Distributions grouped by miniaturization category.

**Supplementary Figure 5.**
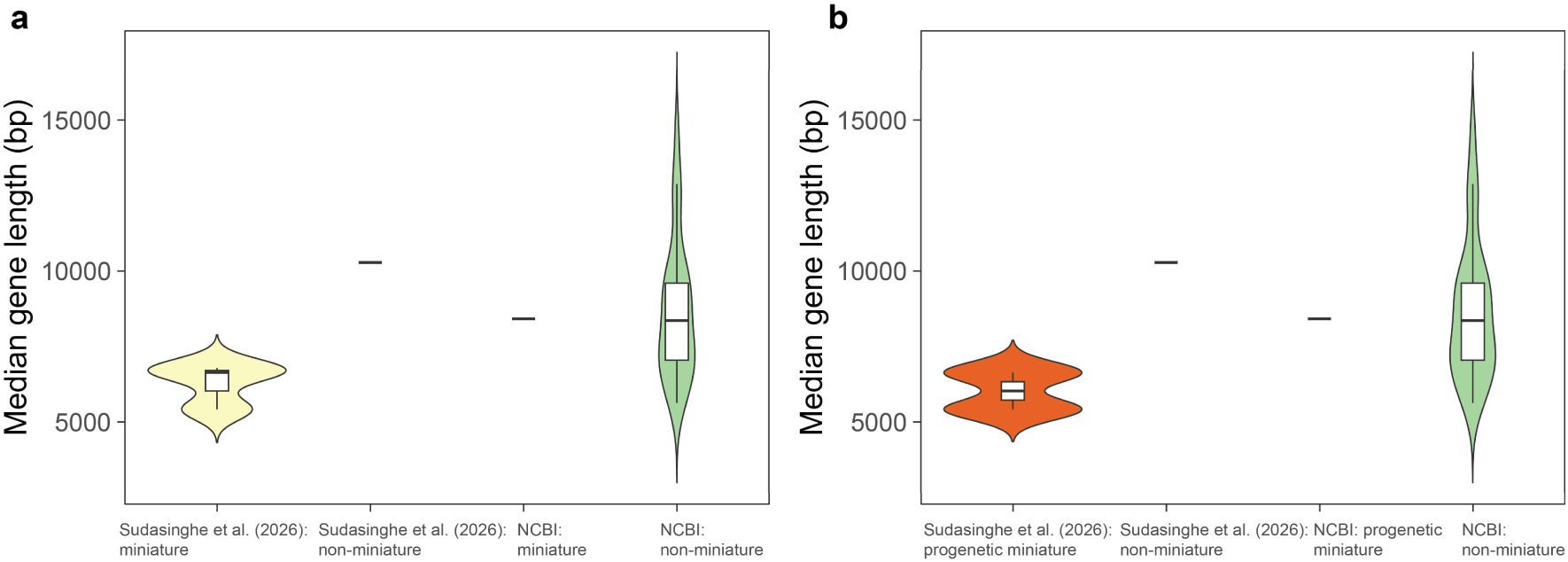
Comparison of median gene lengths between annotation sources for **a)** miniature and non-miniature and **b)** progenetic miniature and non-miniature species, confirming the absence of systematic bias in gene length estimates between annotation sources in the 33-species comparative genomic dataset.

**Supplementary Figure 6.**
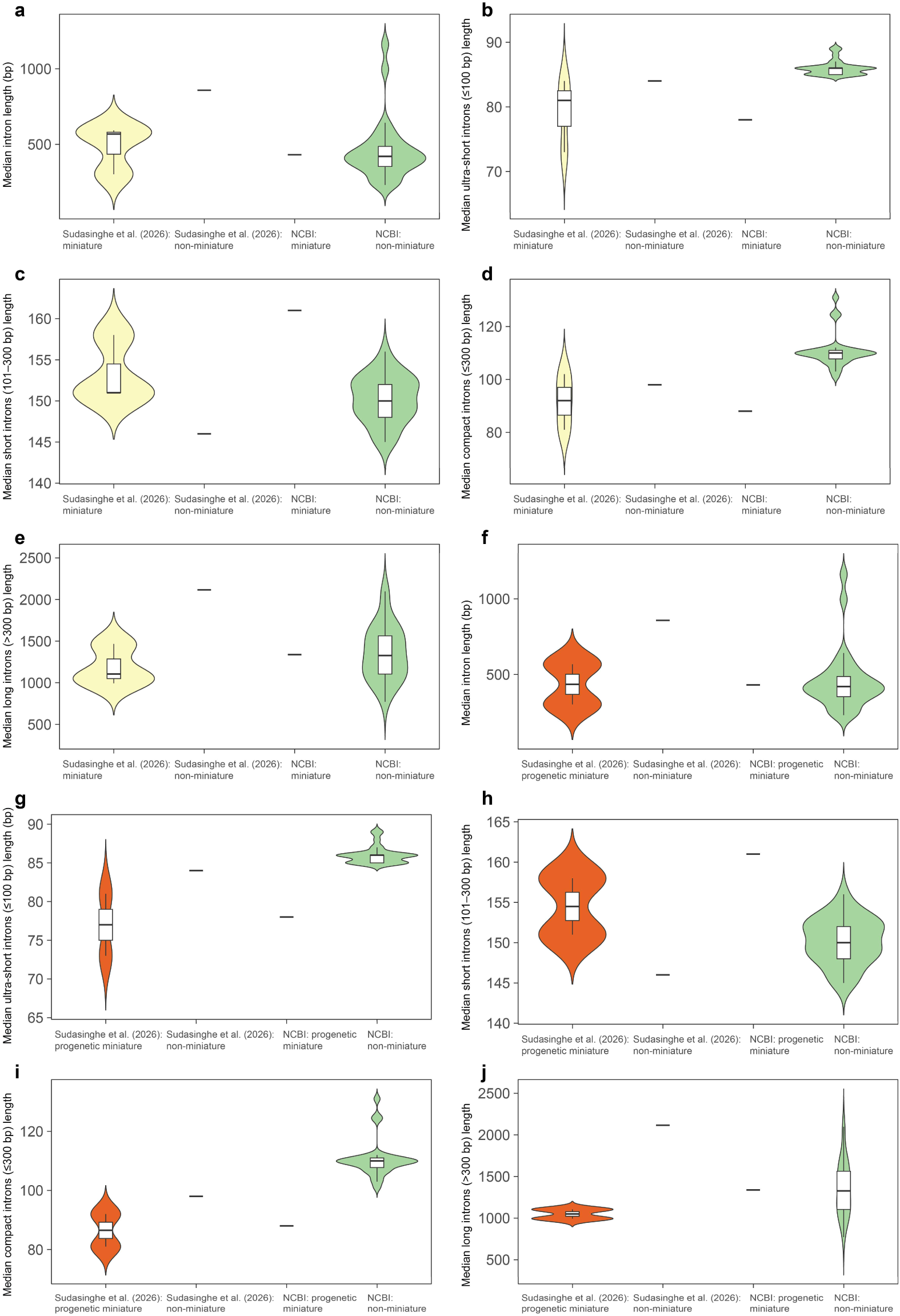
Comparison of median intron lengths by size category between annotation sources for **a–e)** miniature and non-miniature and **f–j)** progenetic miniature and non-miniature species, confirming the absence of systematic bias in intron length estimates between annotation sources in the 33-species comparative genomic dataset. Panels show **a, f)** overall median intron length; **b, g)** median ultra-short intron length (≤100 bp); **c, h)** median short intron length (101–300 bp); **d, i)** median compact intron length (≤300 bp); and **e, j)** median long intron length (>300 bp).

**Supplementary Figure 7.**
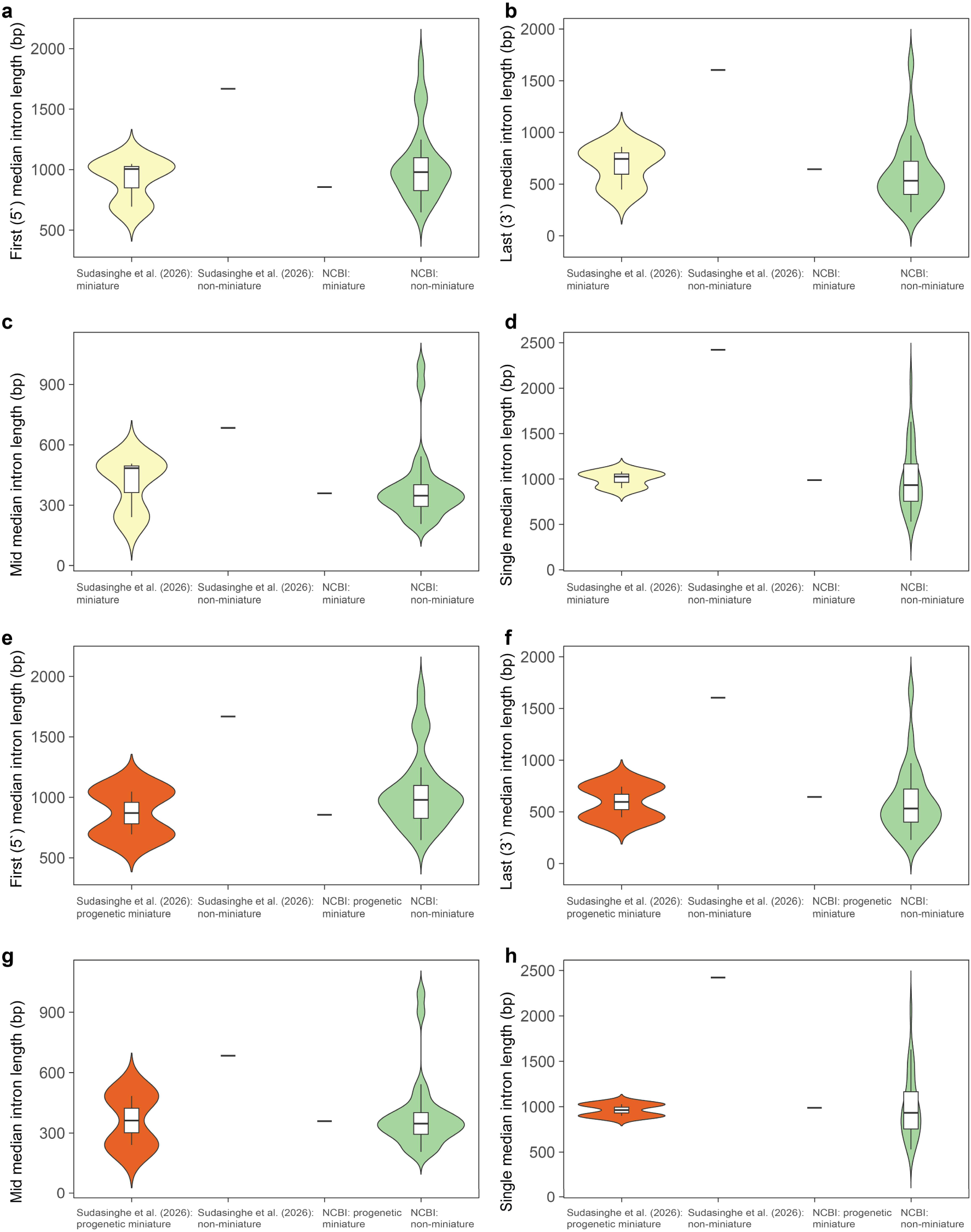
Comparison of median intron lengths by positional category between annotation sources for **a–d)** miniature and non-miniature and **e–h)** progenetic miniature and non-miniature species, confirming the absence of systematic bias in intron length estimates between annotation sources in the 33-species comparative genomic dataset. Panels show **a, e)** first (5′) intron length; **b, f)** last (3′) intron length; **c, g)** middle intron length; and **d, h)** single intron length.

**Supplementary Figure 8.**
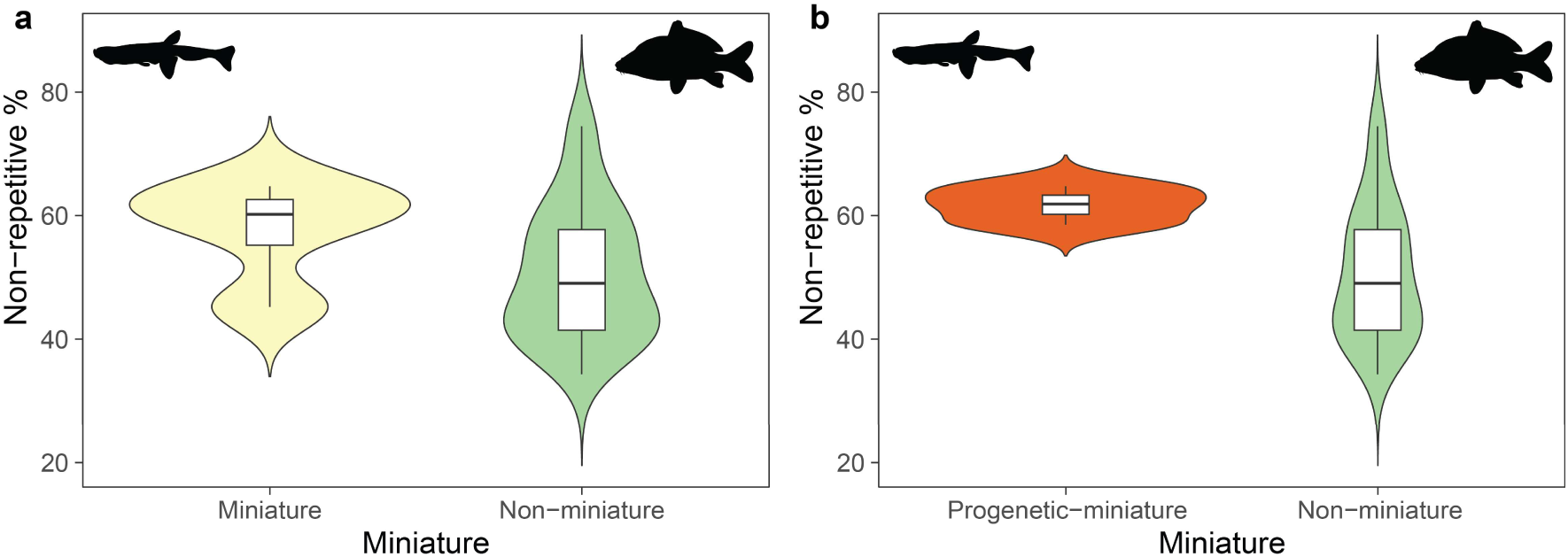
Distribution of the percentage of non-repetitive DNA across **a)** miniature and non-miniature and **b)** progenetic miniature and non-miniature cypriniform species included in the comparative genomic analyses of transposable element content.

**Supplementary Figure 9.**
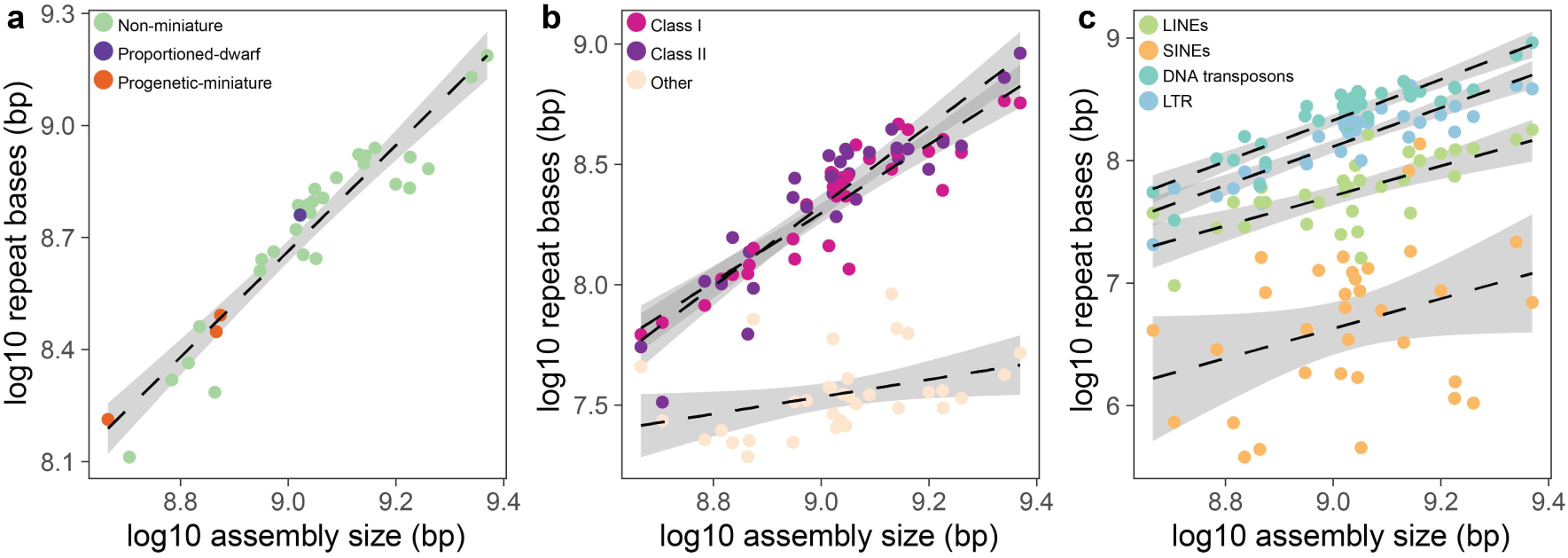
Relationship between log-transformed genome assembly size and log-transformed repeat content across the 33-species comparative genomic dataset. **a)** Data points colored by miniaturization category (non-miniature, proportioned dwarf, and progenetic miniature). **b)** Data points colored by major repeat class (Class I, Class II, and other repeats). **c)** Data points colored by repeat subclass (LINEs, SINEs, DNA transposons, and LTR elements).

**Supplementary Figure 10.**
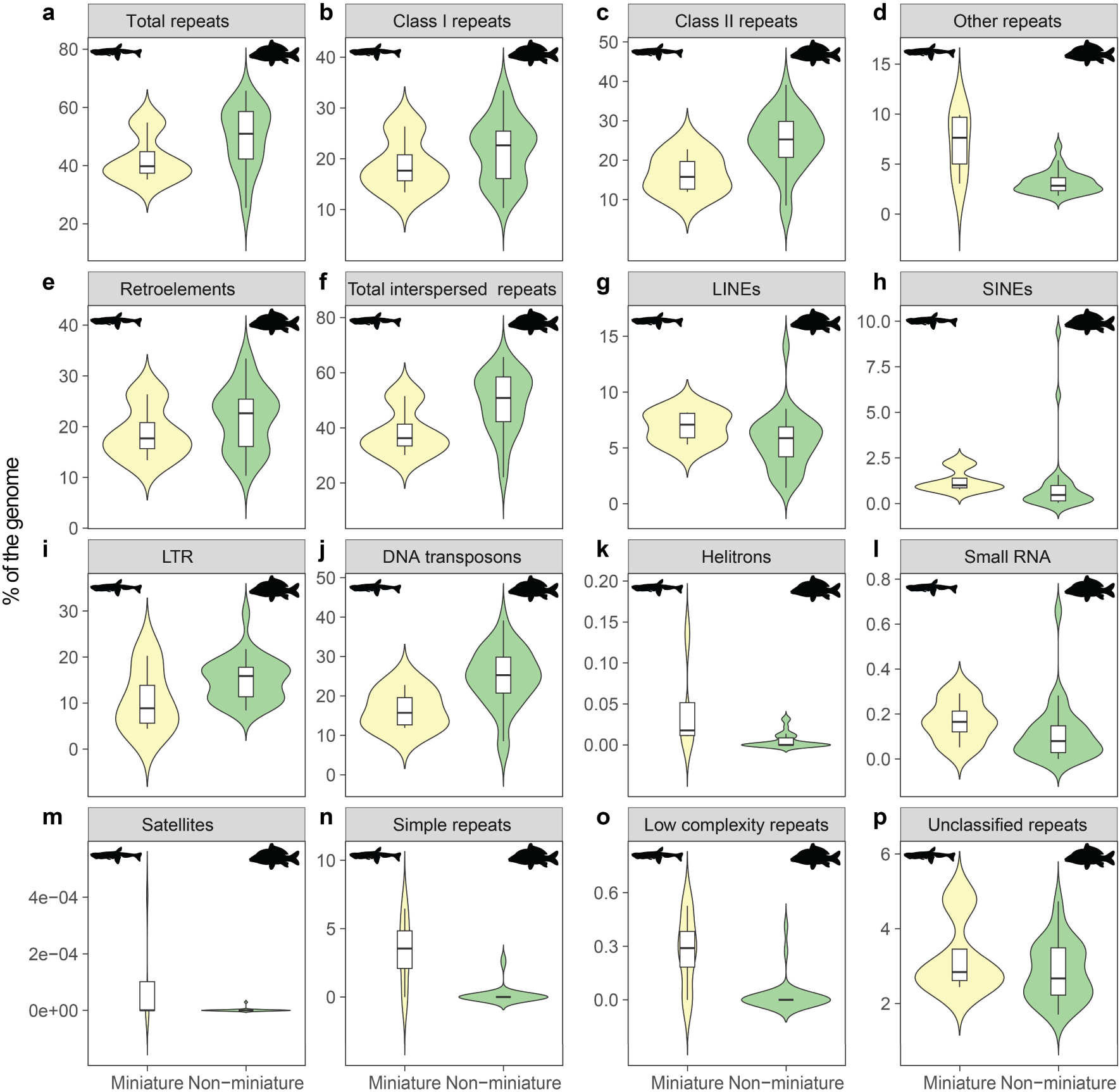
Comparison of repeat content as a percentage of the genome between miniature (n = 4) and non-miniature (n = 29) cypriniform species included in the comparative genomic analyses of transposable element content. Panels show **a)** total repeats; **b)** Class I repeats; **c)** Class II repeats; **d)** other repeats (small RNAs, satellites, simple repeats, low-complexity regions, and unclassified elements); **e)** retroelements; **f)** total interspersed repeats; **g)** LINEs; **h)** SINEs; **i)** LTR elements; **j)** DNA transposons; **k)** helitrons; **l)** small RNAs; **m)** satellites; **n)** simple repeats; **o)** low-complexity repeats; and **p)** unclassified repeats.

**Supplementary Figure 11.**
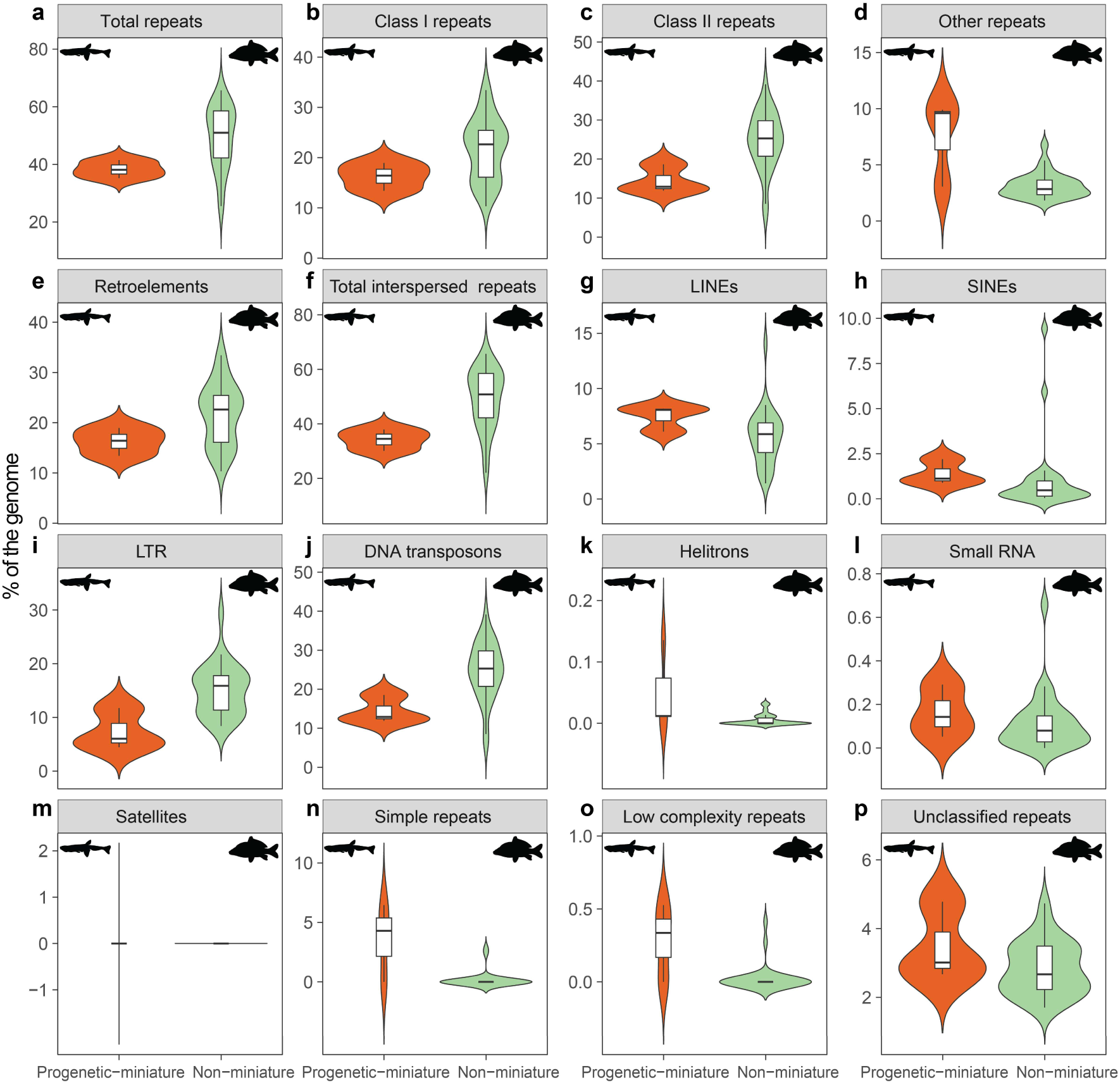
Comparison of repeat content as a percentage of the genome between progenetic-miniature (n = 3) and non-miniature (n = 29) cypriniform species included in the comparative genomic analyses of transposable element content. Panels show **a)** total repeats; **b)** Class I repeats; **c)** Class II repeats; **d)** other repeats (small RNAs, satellites, simple repeats, low-complexity regions, and unclassified elements); **e)** retroelements; **f)** total interspersed repeats; **g)** LINEs; **h)** SINEs; **i)** LTR elements; **j)** DNA transposons; **k)** helitrons; **l)** small RNAs; **m)** satellites; **n)** simple repeats; **o)** low-complexity repeats; and **p)** unclassified repeats.

**Supplementary Figure 12.**
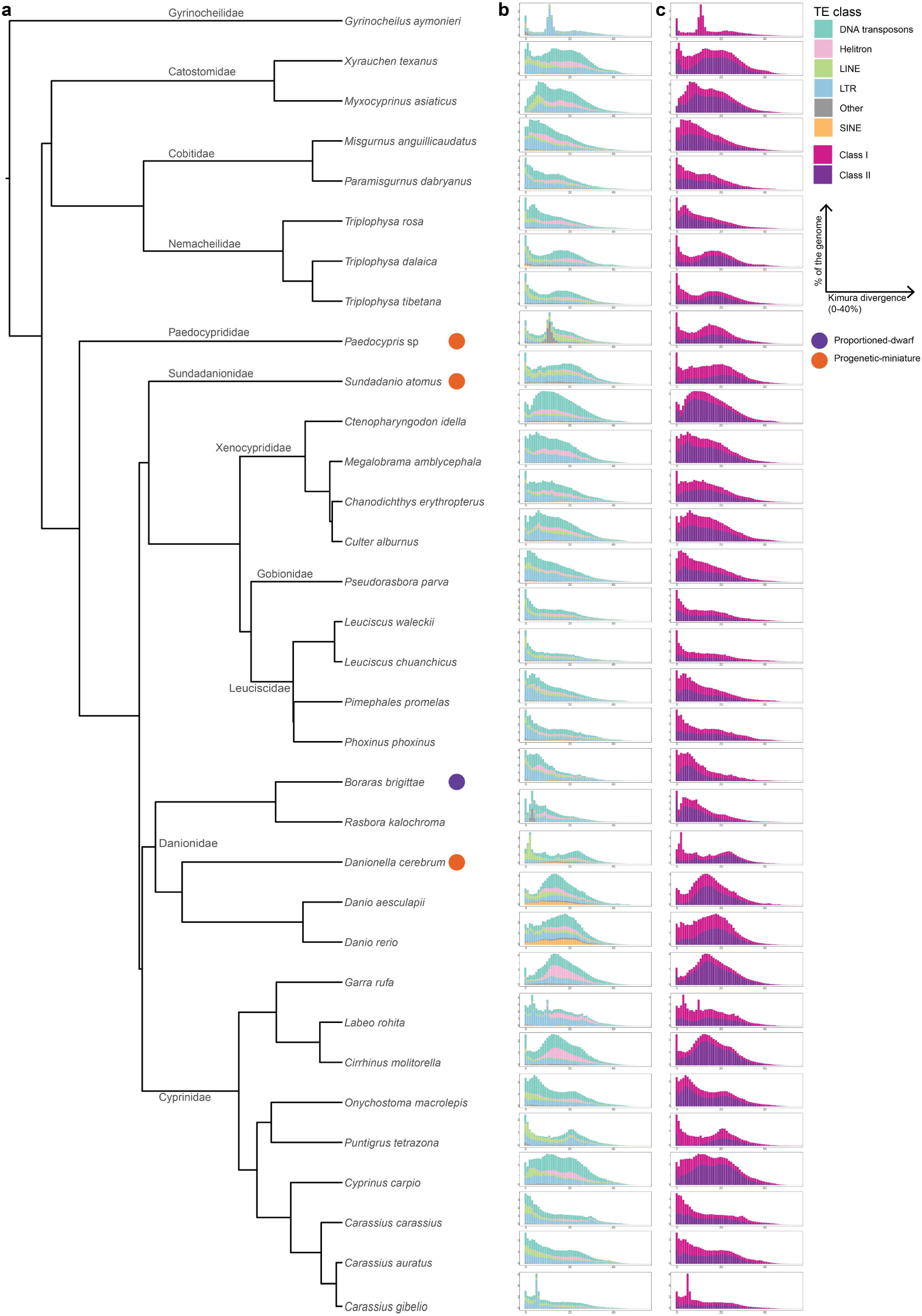
Transposable element (TE) divergence landscapes for the 33-species comparative genomic dataset. **a)** Phylogenetic relationships among the 33 taxa included in the comparative genomic analyses. **b)** Repeat landscapes showing the genomic proportion occupied by major TE subclasses and other repeat categories (small RNAs, satellites, simple repeats, low-complexity regions, and unclassified elements) for each species. **c)** Repeat landscapes showing the genomic proportion occupied by Class I (LTR elements, LINEs, SINEs) and Class II (DNA transposons, helitrons) transposable elements for each species. In panels **b** and **c**, the x-axis represents CpG-adjusted Kimura two-parameter (K2P) divergence from TE consensus sequences, where low divergence values (left) correspond to recently active insertions and high divergence values (right) reflect older, inactive transposition events.

**Supplementary Figure 13.**
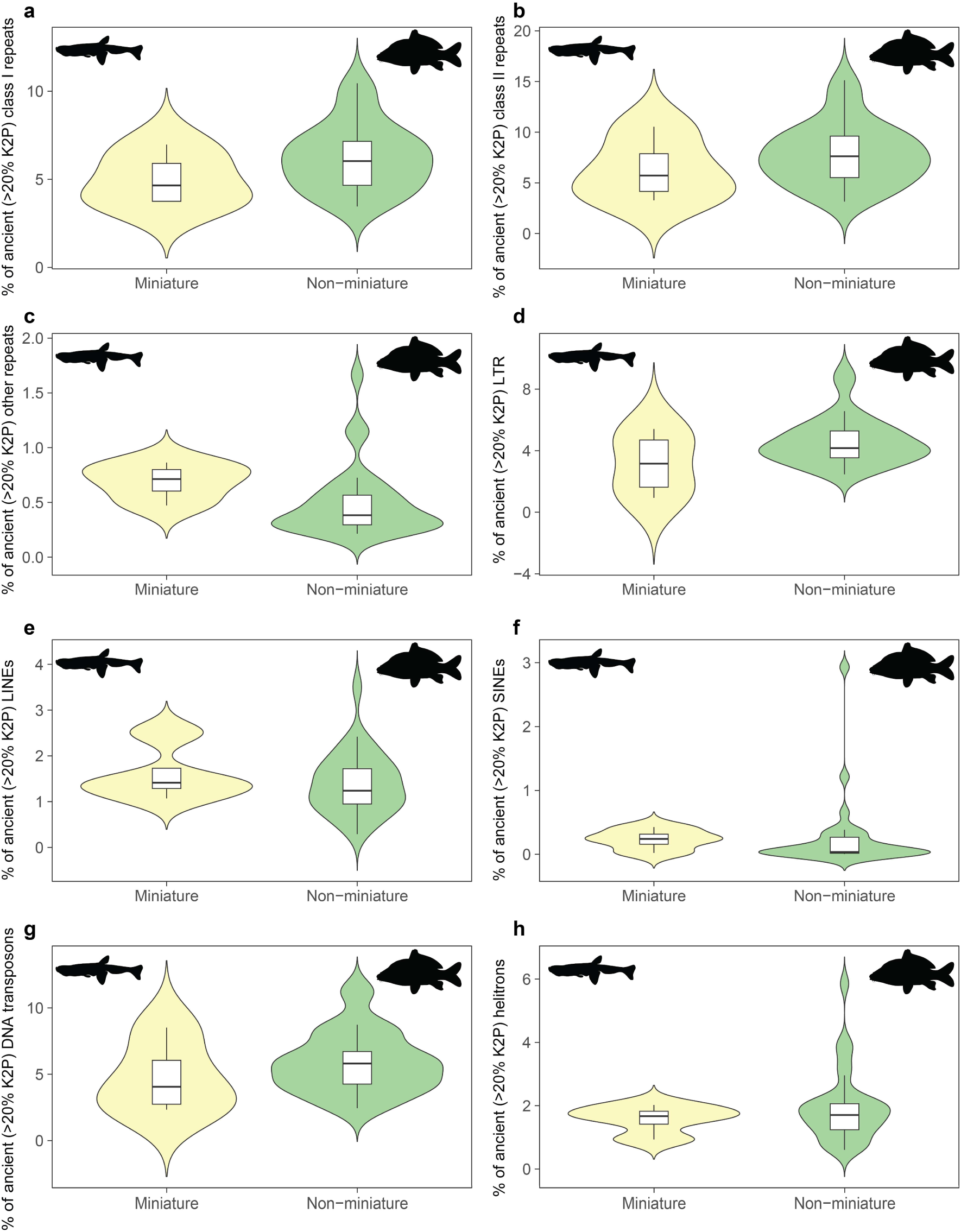
Comparison of ancient transposable element (TE) insertions (CpG-adjusted Kimura two-parameter divergence > 20% from TE consensus sequences) between miniature (n = 4) and non-miniature (n = 29) cypriniform species included in the comparative genomic analyses of TE content. Panels show **a)** Class I repeats; **b)** Class II repeats; **c)** other repeats (small RNAs, satellites, simple repeats, low-complexity regions, and unclassified elements); **d)** LTR elements; **e)** LINEs; **f)** SINEs; **g)** DNA transposons; and **h)** helitrons.

**Supplementary Figure 14.**
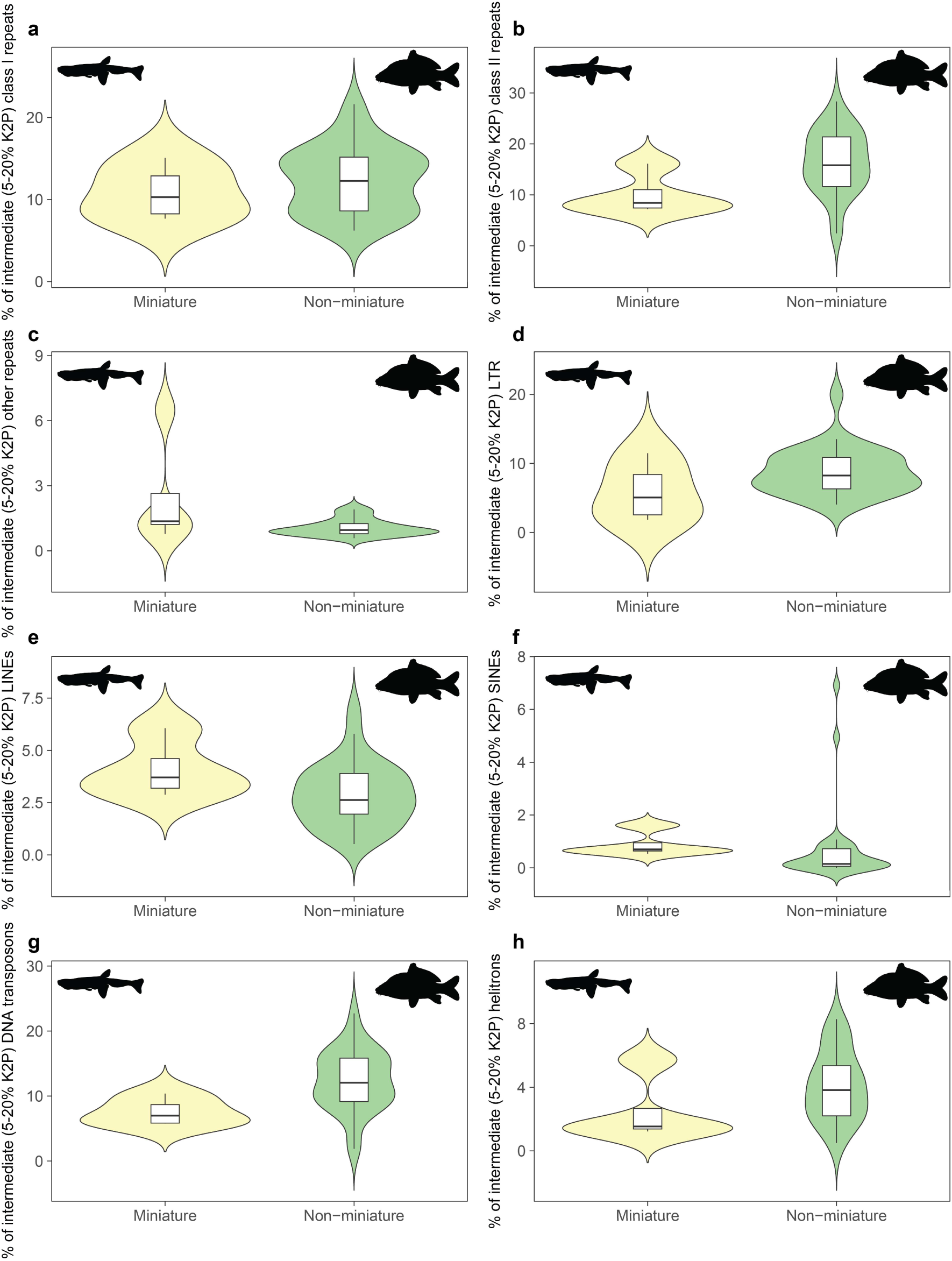
Comparison of intermediate transposable element (TE) insertions (CpG-adjusted Kimura two-parameter divergence 5-20% from TE consensus sequences) between miniature (n = 4) and non-miniature (n = 29) cypriniform species included in the comparative genomic analyses of TE content. Panels show **a)** Class I repeats; **b)** Class II repeats; **c)** other repeats (small RNAs, satellites, simple repeats, low-complexity regions, and unclassified elements); **d)** LTR elements; **e)** LINEs; **f)** SINEs; **g)** DNA transposons; and **h)** helitrons.

**Supplementary Figure 15.**
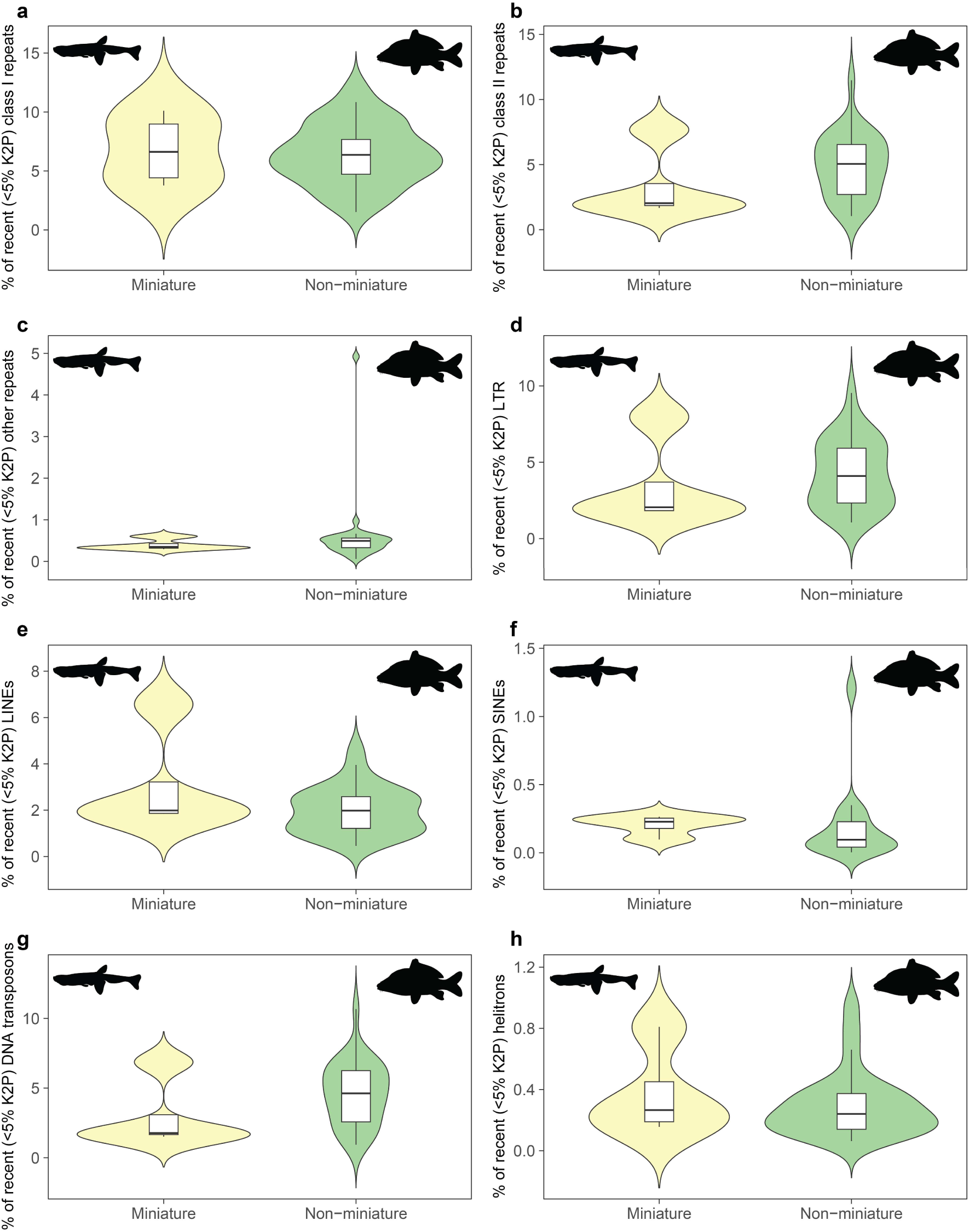
Comparison of recent transposable element (TE) insertions (CpG-adjusted Kimura two-parameter divergence < 5% from TE consensus sequences) between miniature (n = 4) and non-miniature (n = 29) cypriniform species included in the comparative genomic analyses of TE content. Panels show **a)** Class I repeats; **b)** Class II repeats; **c)** other repeats (small RNAs, satellites, simple repeats, low-complexity regions, and unclassified elements); **d)** LTR elements; **e)** LINEs; **f)** SINEs; **g)** DNA transposons; and **h)** helitrons.

**Supplementary Figure 16.**
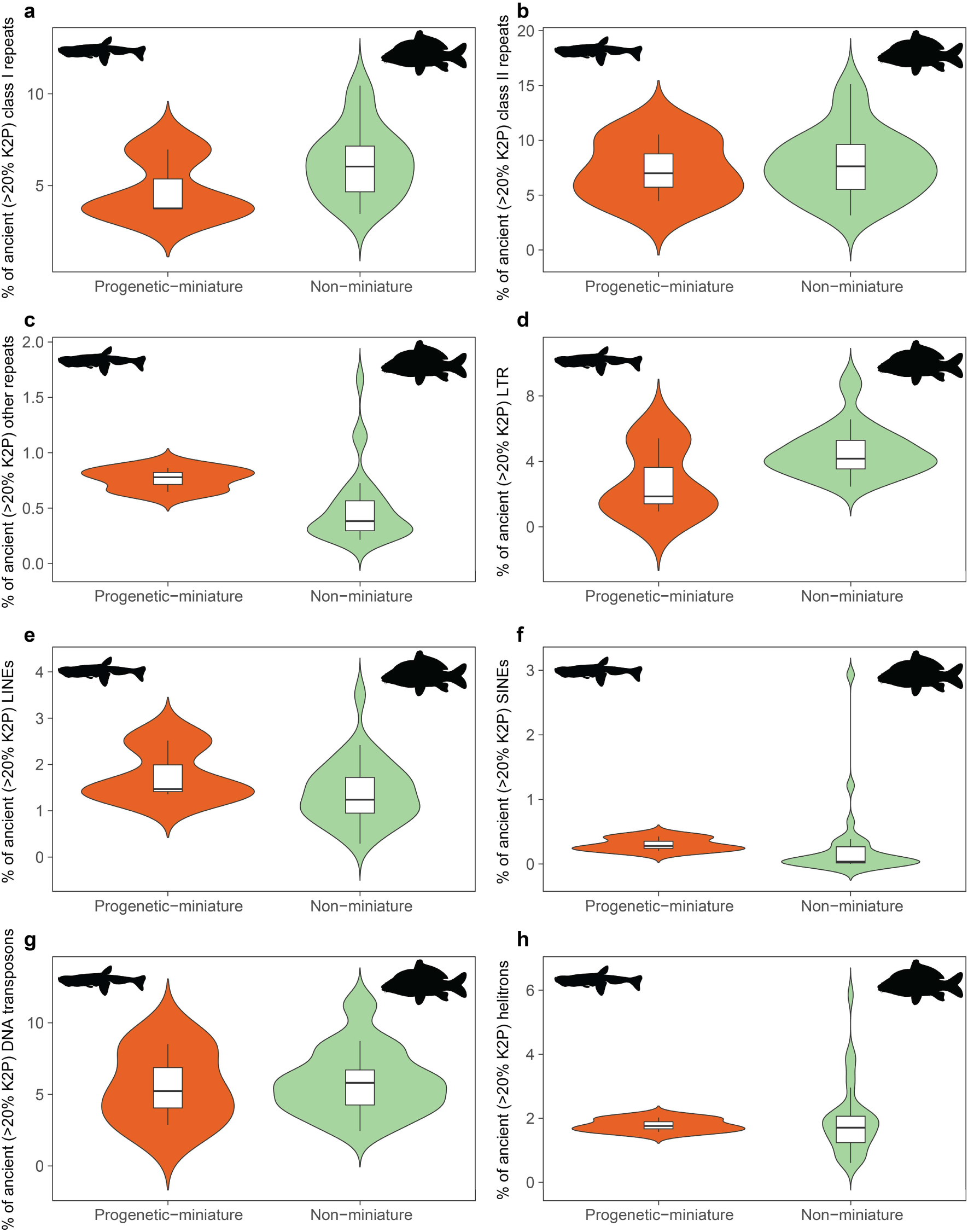
Comparison of ancient transposable element (TE) insertions (CpG-adjusted Kimura two-parameter divergence > 20% from TE consensus sequences) between progenetic-miniature (n = 3) and non-miniature (n = 29) cypriniform species included in the comparative genomic analyses of TE content. Panels show **a)** Class I repeats; **b)** Class II repeats; **c)** other repeats (small RNAs, satellites, simple repeats, low-complexity regions, and unclassified elements); **d)** LTR elements; **e)** LINEs; **f)** SINEs; **g)** DNA transposons; and **h)** helitrons.

**Supplementary Figure 17.**
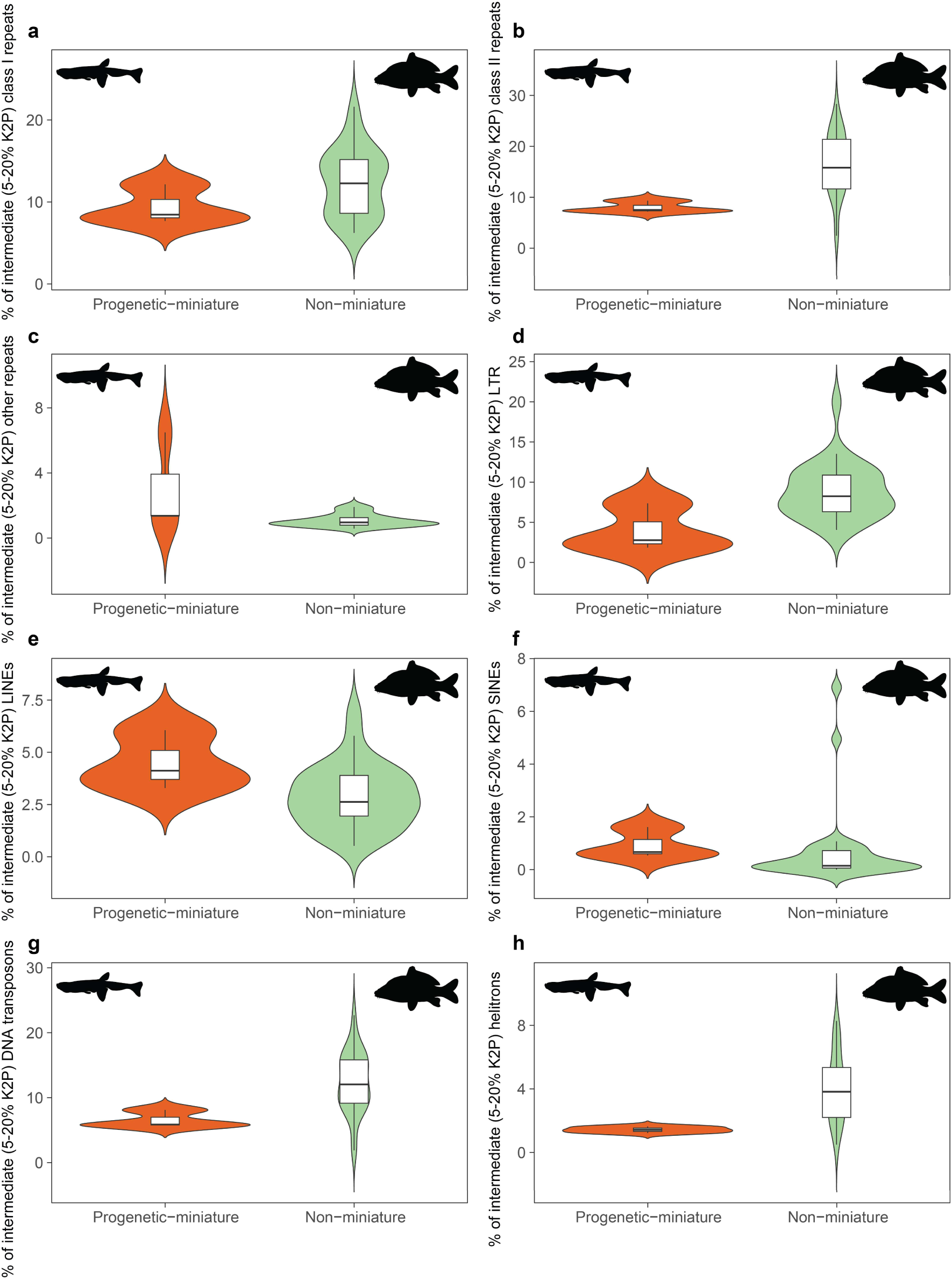
Comparison of intermediate transposable element (TE) insertions (CpG-adjusted Kimura two-parameter divergence 5-20% from TE consensus sequences) between progenetic-miniature (n = 3) and non-miniature (n = 29) cypriniform species included in the comparative genomic analyses of TE content. Panels show **a)** Class I repeats; **b)** Class II repeats; **c)** other repeats (small RNAs, satellites, simple repeats, low-complexity regions, and unclassified elements); **d)** LTR elements; **e)** LINEs; **f)** SINEs; **g)** DNA transposons; and **h)** helitrons.

**Supplementary Figure 18.**
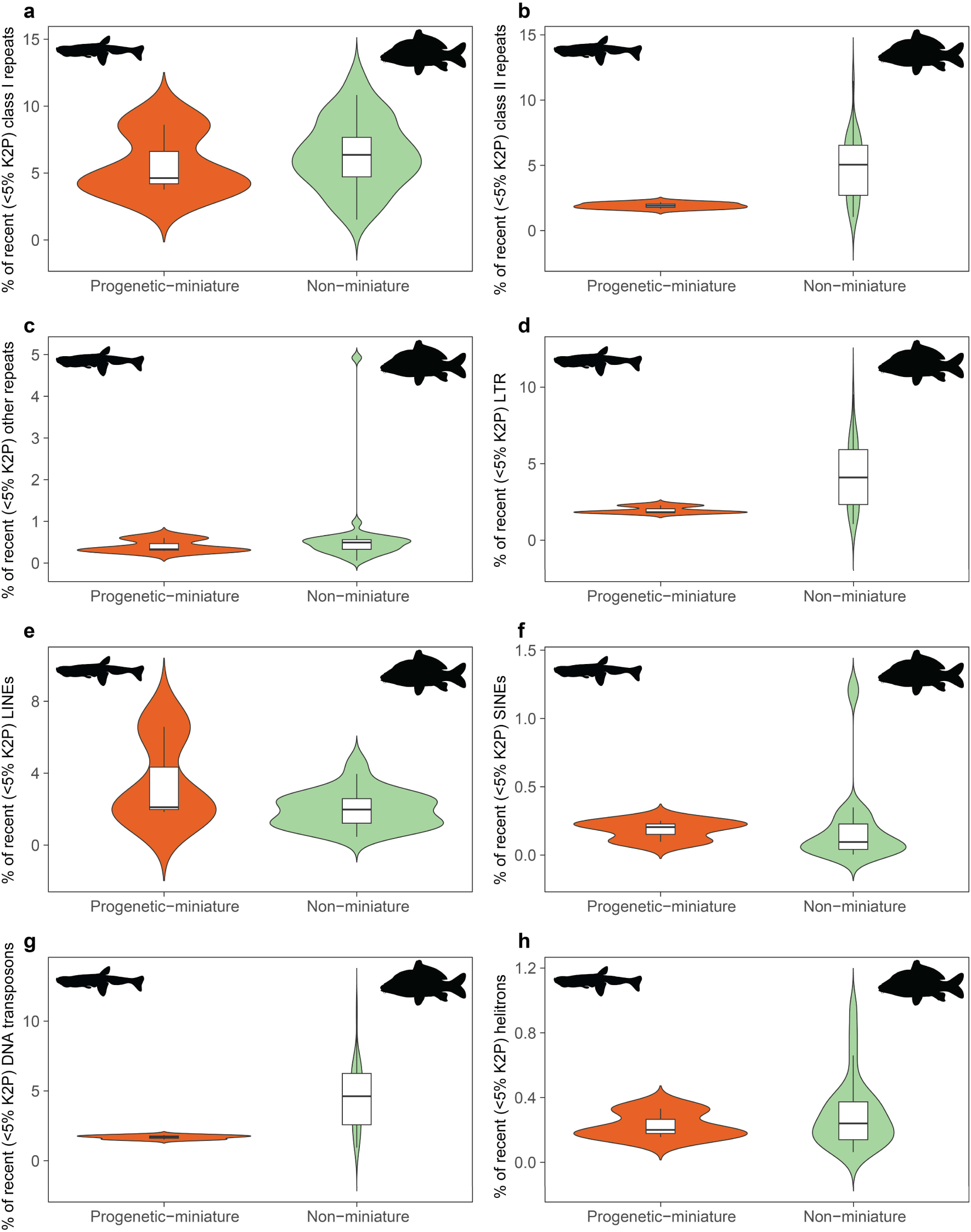
Comparison of recent transposable element (TE) insertions (CpG-adjusted Kimura two-parameter divergence < 5% from TE consensus sequences) between progenetic-miniature (n = 3) and non-miniature (n = 29) cypriniform species included in the comparative genomic analyses of TE content. Panels show **a)** Class I repeats; **b)** Class II repeats; **c)** other repeats (small RNAs, satellites, simple repeats, low-complexity regions, and unclassified elements); **d)** LTR elements; **e)** LINEs; **f)** SINEs; **g)** DNA transposons; and **h)** helitrons.

**Supplementary Figure 19.**
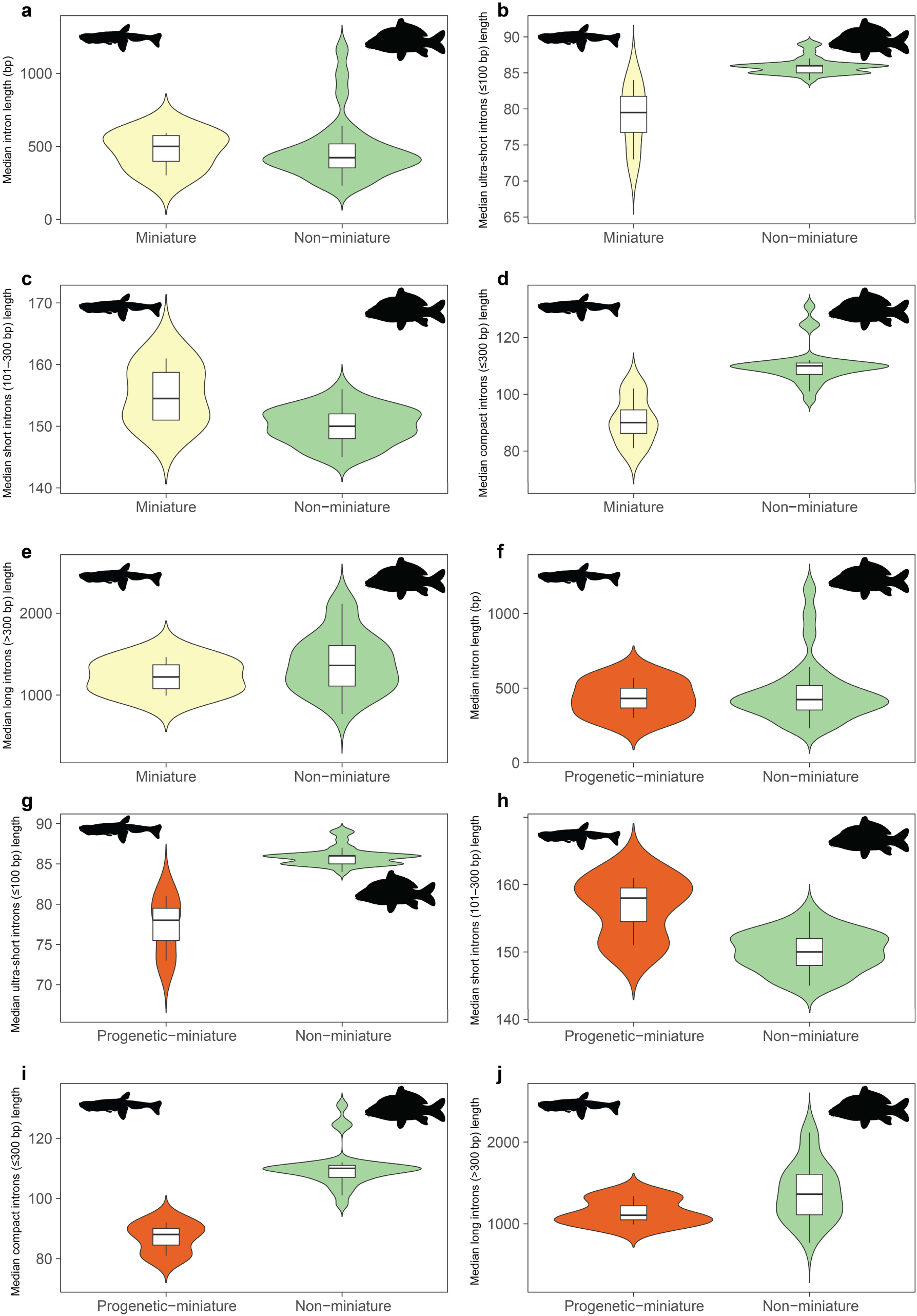
Comparison of median intron lengths by size category between **a–e)** miniature (n = 4) and non-miniature (n = 29) and **f–j)** progenetic miniature (n = 3) and non-miniature (n = 29) cypriniform species in the 33-species comparative genomic dataset. Panels show **a, f)** overall median intron length; **b, g)** median ultra-short intron length (≤100 bp); **c, h)** median short intron length (101–300 bp); **d, i)** median compact intron length (≤300 bp); and **e, j)** median long intron length (>300 bp).

**Supplementary Figure 20.**
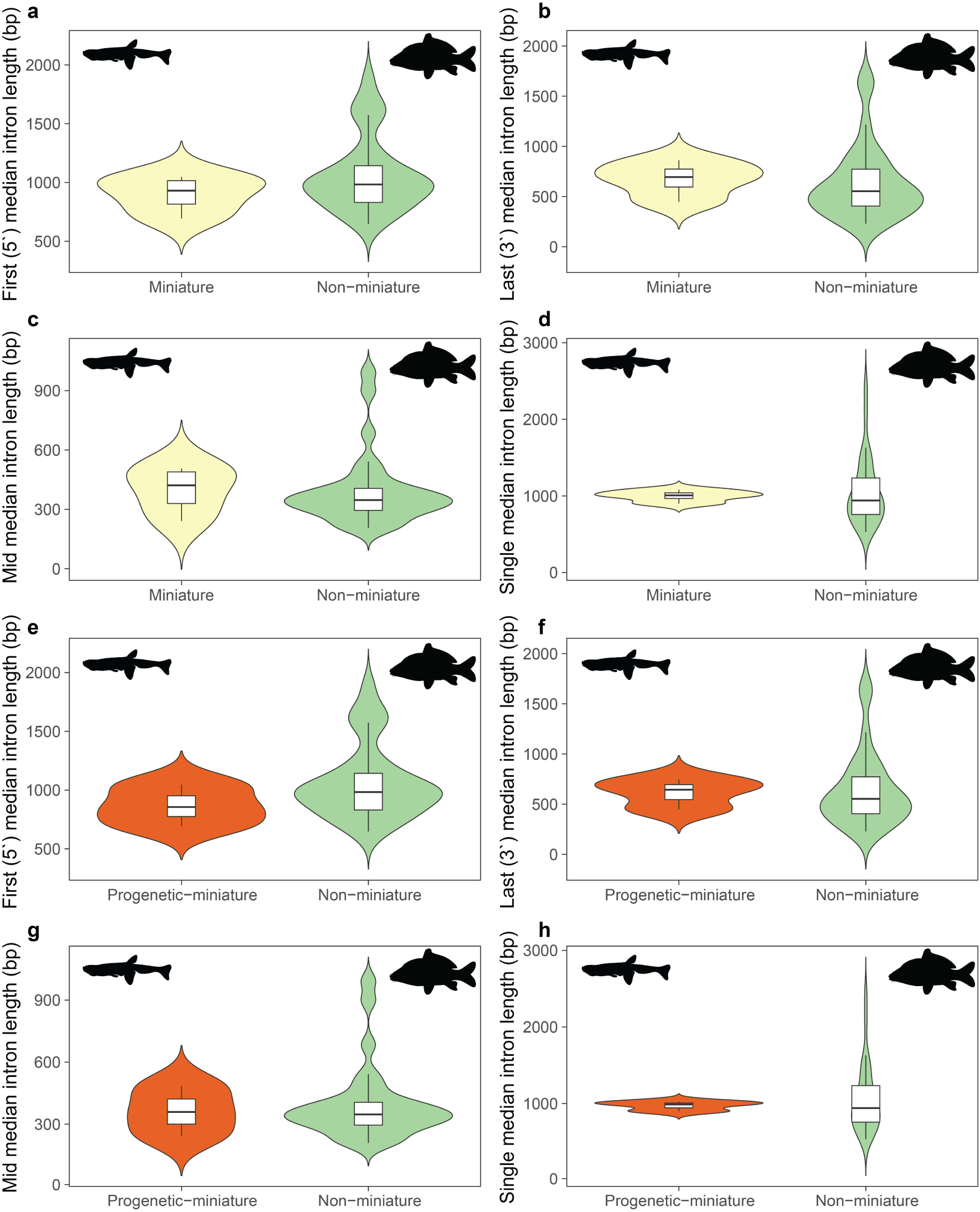
Comparison of median intron lengths by positional category between **a–d)** miniature (n = 4) and non-miniature (n = 29) and **e–h)** progenetic miniature (n = 3) and non-miniature (n = 29) cypriniform species in the 33-species comparative genomic dataset. Panels show **a, e)** first (5′) intron length; **b, f)** last (3′) intron length; **c, g)** middle intron length; and **d, h)** single intron length.

**Supplementary Figure 21.**
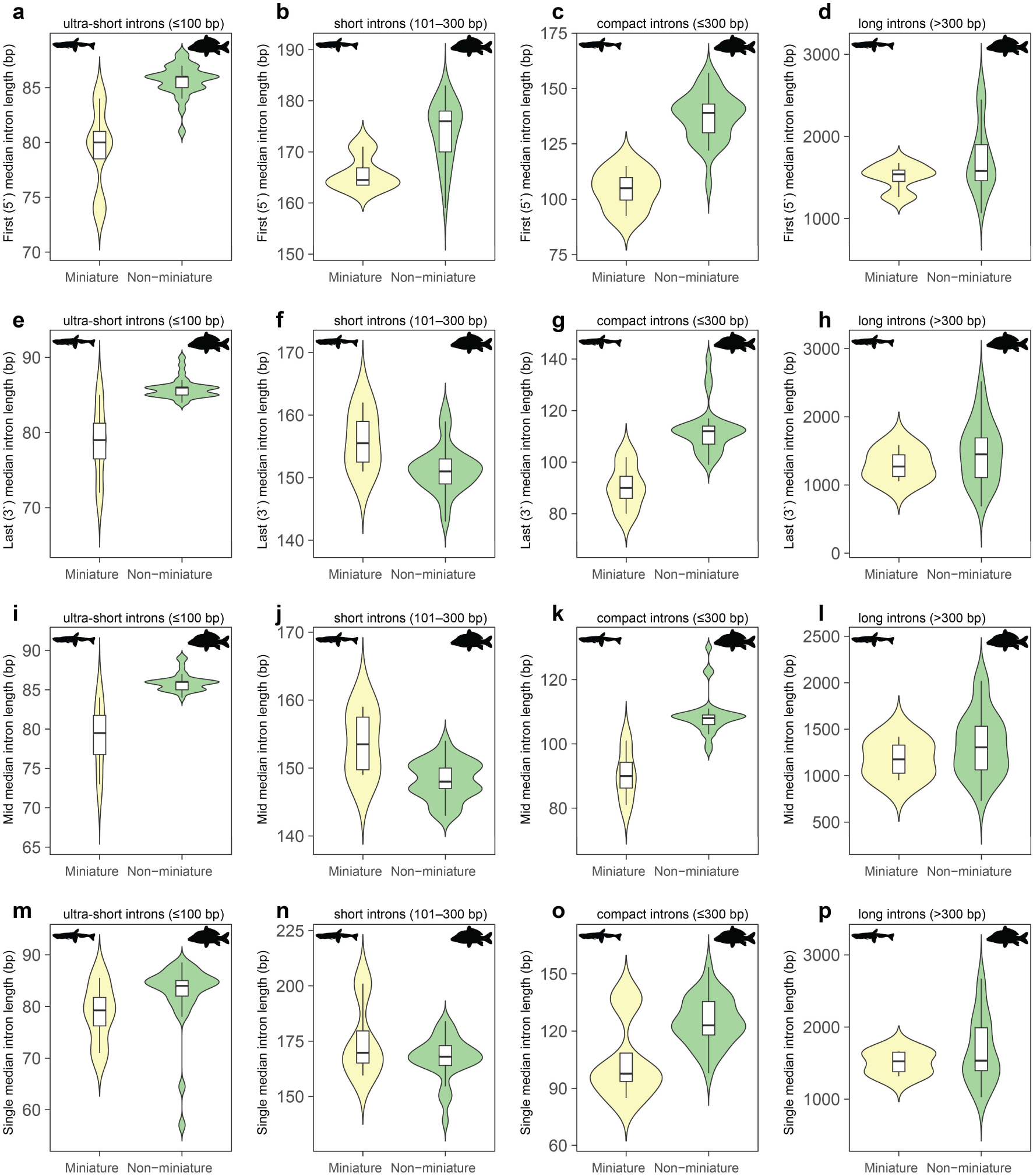
Comparison of median intron lengths by combined size class and positional category between miniature (n = 4) and non-miniature (n = 29) cypriniform species in the 33-species comparative genomic dataset. First (5′) intron position for **a)** ultra-short (≤100 bp), **b)** short (101–300 bp), **c)** compact (≤300 bp), and **d)** long (>300 bp) introns. Last (3′) intron position for **e)** ultra-short, **f)** short, **g)** compact, and **h)** long introns. Middle intron position for **i)** ultra-short, **j)** short, **k)** compact, and **l)** long introns. Single introns for **m)** ultra-short, **n)** short, **o)** compact, and **p)** long introns.

**Supplementary Figure 22.**
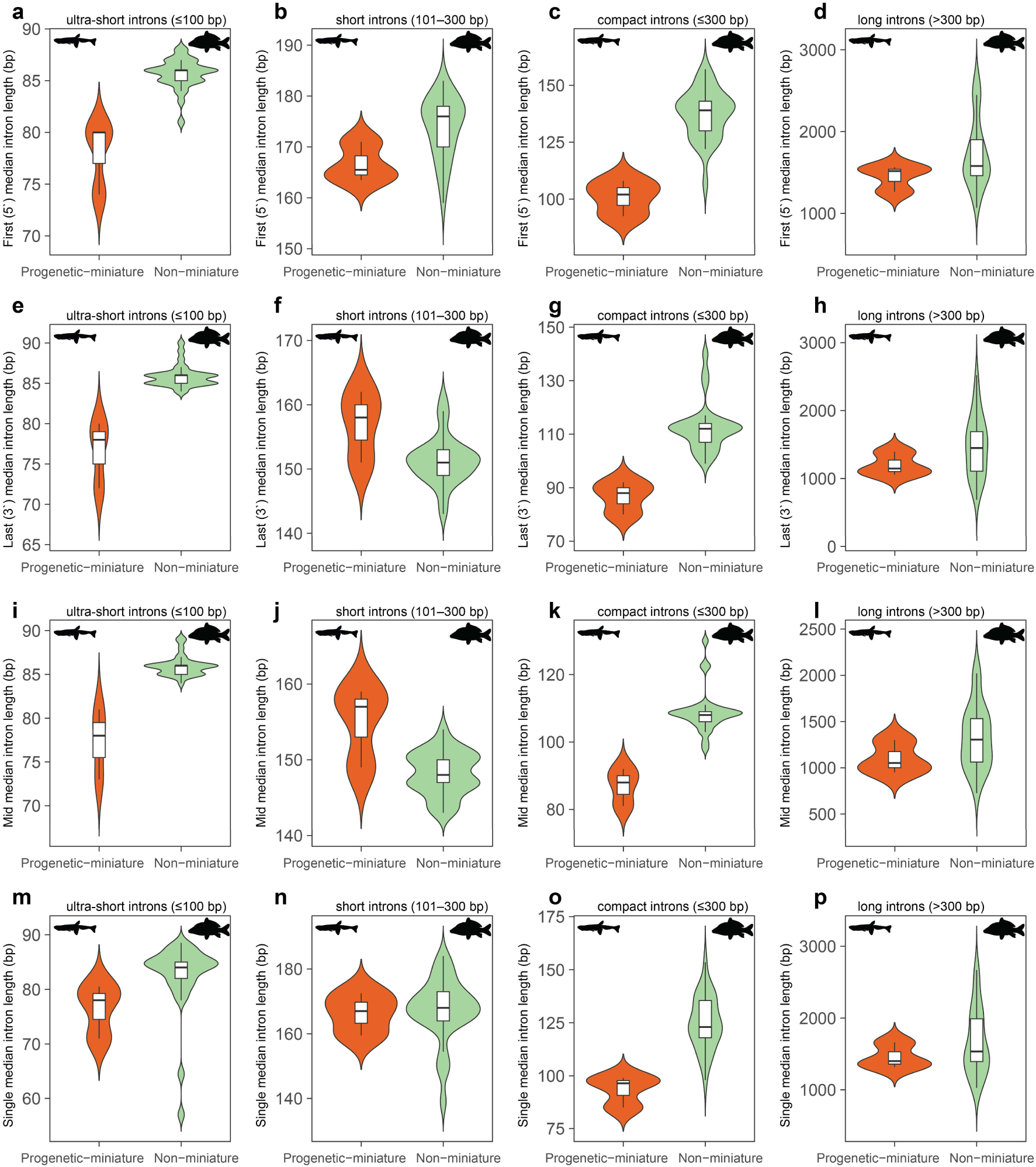
Comparison of median intron lengths by combined size class and positional category between progenetic miniature (n = 3) and non-miniature (n = 29) cypriniform species in the 33-species comparative genomic dataset. First (5′) intron position for **a)** ultra-short (≤100 bp), **b)** short (101–300 bp), **c)** compact (≤300 bp), and **d)** long (>300 bp) introns. Last (3′) intron position for **e)** ultra-short, **f)** short, **g)** compact, and **h)** long introns. Middle intron position for **i)** ultra-short, **j)** short, **k)** compact, and **l)** long introns. Single introns for **m)** ultra-short, **n)** short, **o)** compact, and **p)** long introns.

**Supplementary Figure 23.**
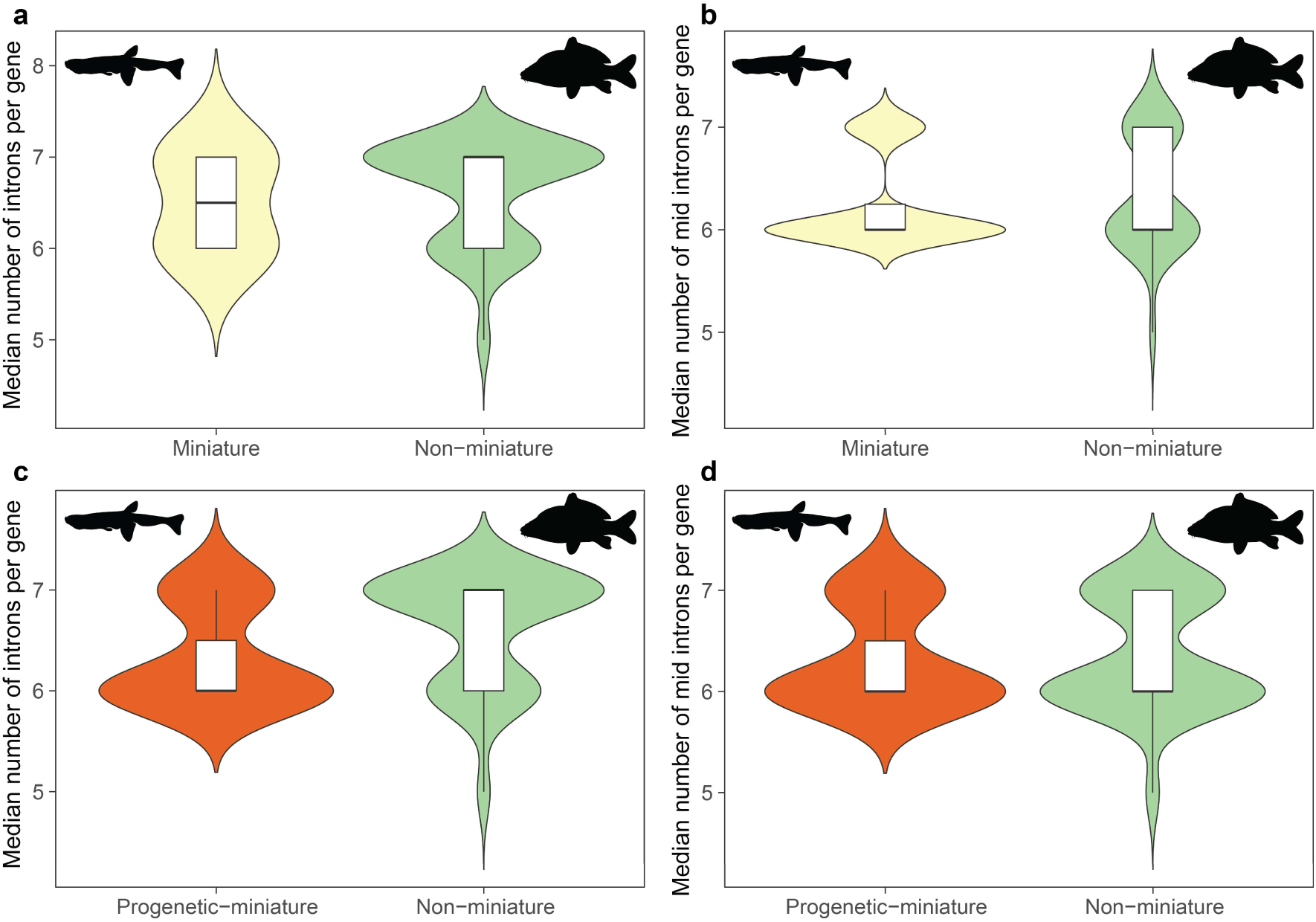
Median number of introns per gene in the cypriniform species in the 33-species comparative genomic dataset in **a)** miniature (n = 4) and non-miniature (n = 29) and **c)** progenetic miniature (n = 3) and non-miniature (n = 29) species. Median number of middle introns per gene in **b)** miniature (n = 4) and non-miniature (n = 29) and **d)** progenetic miniature (n = 3) and non-miniature (n = 29) species.

**Supplementary Figure 24.**
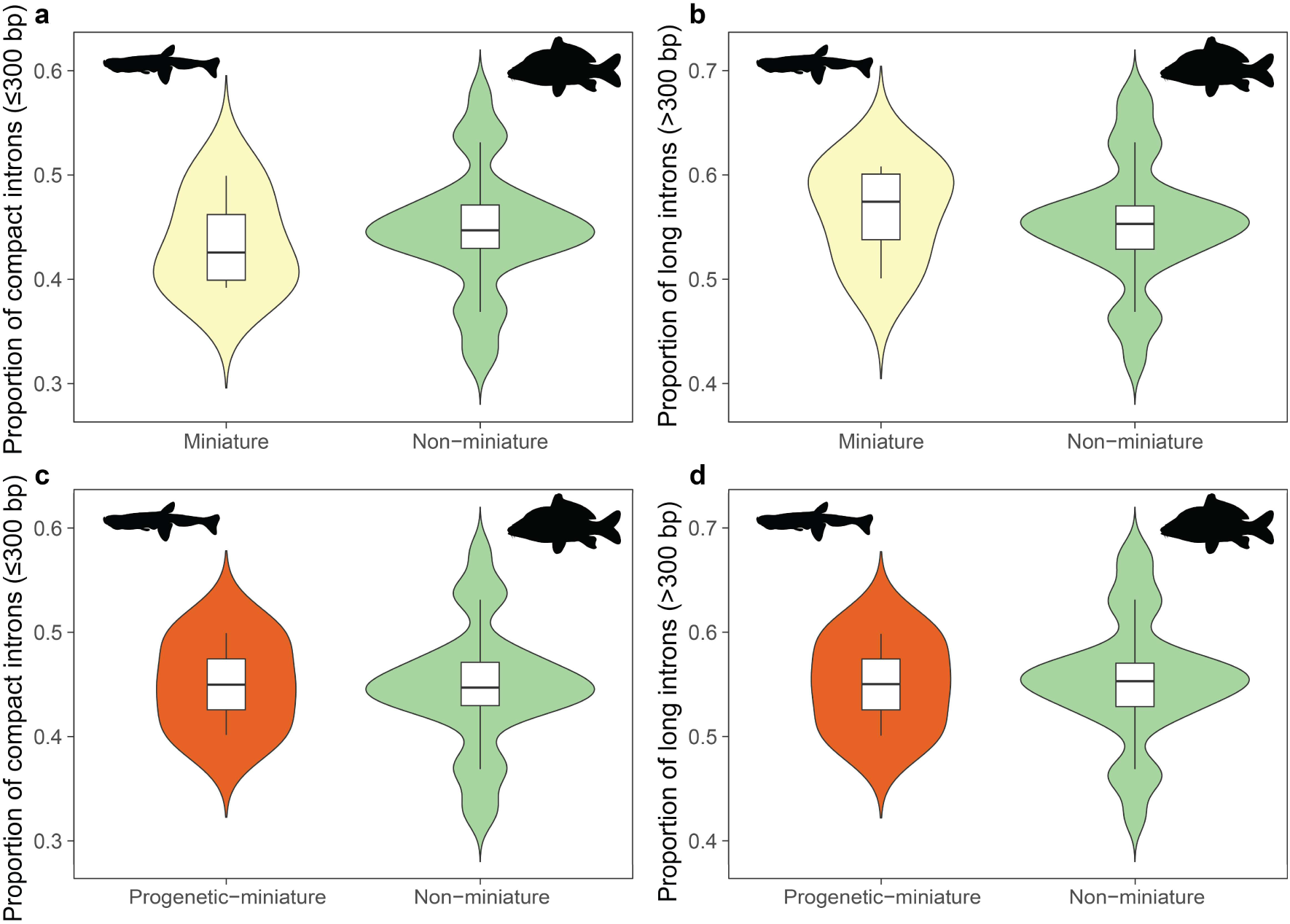
Proportion of compact (≤300 bp) and long (>300 bp) introns in the 33-species comparative genomic dataset. Proportion of compact introns in **a)** miniature (n = 4) and non-miniature (n = 29) and **c)** progenetic miniature (n = 3) and non-miniature (n = 29) species. Proportion of long introns in **b)** miniature (n = 4) and non-miniature (n = 29) and **d)** progenetic miniature (n = 3) and non-miniature (n = 29) species.

