## Supplementary Tables for "Convergent genome streamlining accompanies independent miniaturization in the world’s smallest fishes"

**Supplementary Information 1.** Fossil calibrations used in Bayesian divergence time estimation, including taxonomic assignment, minimum age, and justification for inclusion. Excluded calibrations and the reasons for their exclusion are also documented.

Our fossil calibration comprised the following ingroup fossils:

***†Paleogobio zhongyuanensis*** Zhou 1990 (Chang & Chen, 2008) as a stem representative of Cyprinoidei (Early Eocene, Fanxian, China, minimum age 47.0 Ma).

†*Palaeogobio zhongyuanensis* is an imperfectly known fossil described by Zhou (1990) based on an articulated almost complete skeleton from the Middle Miocene of the Shanwang Basin in China. We have no way of independently verifying the accuracy of its taxonomic assignment but note that Chang & Chen (2008) list this species as the earliest member of the Cyprinidae. We used a conservative approach treating this fossil as *incertae sedis* in Cyprinoidei and therefore used this fossil to calibrate the node of the last common ancestor of this taxon.

***†Cobitis nanningensis*** Chen, Liao & Lei 2015 (Chen et al., 2015) as a stem representative of Cobitidae (Early-Middle Oligocene, lover part of Yongning Formation, minimum age 28.1 Ma).

†*Cobitis nanningensis* was described by Chen et al. (2015) from Early-Middle Oligocene deposits of Nanning Basin, Guangxi Zhuang Autonomous Region, southern China. This oldest fossil cobitid species is based on a series of 1.8–3 mm long bifid spines, interpreted by the authors to represent cobitid suborbital spines. We agree that this type of bifid suborbital spine is unique among bony fishes and serves as a sufficiently convincing synapomorphy to assign this fossil fragment to the family Cobitidae and potentially to the genus *Cobitis* (Chen et al. 2015). We therefore calibrated the node of the last common ancestor of all Cobitidae with this fossil.

***†Amyzon aggregatum*** Wilson 1977 and ***†Wilsonium brevipinne*** (Cope, 1893) as stem representatives of Catostomoidei (early Eocene, Allenby formation and Horsefly beds in British Columbia, Canada, minimum age 47.8 Ma).

†*Wilsonium brevipinne* was originally described as a member of †*Amyzon* by Cope (1893) and later transferred to its own genus by Liu et al. (2016) based on the results of a phylogenetic analysis that placed this species as the sister group to all remaining members of Catostomidae, inclusive of the Asian taxa *Myxocyprinus* and *Plesiomyxocyprinus*. There is little doubt that †*Amyzon* and †*Wilsonium* belong to Catostomidae because they exhibit the pectinate pharyngeal teeth on ceratobranchial 5 characteristic of this family (Liu et al., 2016; Wilson, 1977). We therefore calibrated the node of the last common ancestor of all Catostomidae with one of these two fossils from the same time period.

Our fossil calibration comprised the following outgroup fossils:

***†Astephus antiquus*** (Leidy 1873) (Grande & Lundberg, 1988) constraining the split between *Ictalurus furcatus* and *Cranoglanis bouderius* (Early Eocene, Green River Formation, minimum age 46.0 Ma).

†*Astephus antiquus* was originally described as a member of *Pimelodus* by Leidy (1873) based on Eocene material from the Green River Bridger formations. In the same year, Cope (1873) transferred Leidy's species to the newly established †*Astephus*, a subgenus of †*Rhineastes* (Ictaluridae). Since description, †*A. antiquus* has been considered as either member of Ictaluridae (Cope, 1873; Grande & Lundberg, 1988; Lundberg, 1970, 1992), Ariidae (Jordan, 1923)

or the monogeneric †Astephidae (Arce-H. et al., 2017). The study of Arce-H et al. (2017) presented alternative hypotheses on the placement †*Astephus* (including †*A. antiquus*) within sisoroid catfishes resulting from the analyses of different datasets, including: (1) inside of Ictaluridae based on a morphological dataset of 209 characters (Arce-H, et al., 2017: figure 9a); and (2) as sister taxon to a larger group of catfishes including members of Ictaluridae, Schilbeidae, Bagridae, Siluridae and Plotosidae based on a combined dataset of morphological and molecular characters (Arce-H, et al., 2017: figure 7). In this second hypothesis, *Cranoglanis* is placed as the sister taxon to the monophyletic group that includes †*Astephus* plus the aforementioned families of catfishes, and not as the sister taxon of Ictaluridae, as is commonly reported (ref herein). This second hypothesis stems from the analysis of a combined dataset with large amounts of missing data for †*Astephus* and other extinct taxa and we consider the first hypothesis that places †*Astephus* as part of a monophyletic group that also includes all members of Ictaluridae and *Cranoglanis* as more plausible.

***†Rubiesichthys gregalis*** Wenz 1984 (Poyato-Ariza, 1996a) as a stem representative of Gonorynchiformes (Early Cretaceous, El Montsech and Las Hoyas (Spain), minimum age 132.6 Ma).

†*Rubiesichthys gregalis* was originally described by Wenz (1984) from the Lower Cretaceous of Montsech in Spain and subsequently assigned to the family Chanidae by (Poyato-Ariza, 1996a, 1996b) after the study of additional material from the type locality and Las Hoyas, Spain. This systematic placement has not been challenged and we have used †*Rubiesichthys* to calibrate the node uniting *Chanos* and Otophysi (i.e. the last common ancestor of Ostariophysi).

We refrained from using the below listed fossils that have sometimes been used as calibration points in molecular studies. In our opinion, morphological re-examination of these fossils is urgently needed before they can be used as reliable calibration points in molecular studies.

**†*Tischlingerichthys viohli*** Arratia 1997
This is a fossil fish from the Late Jurassic of Bavaria, Germany. It was described by Arratia (1997) based on a single specimen of about 126 mm standard length. It has since been used in numerous molecular analyses to calibrate time trees starting with Benton et al. (2015). In the original paper Arratia suggested that (p. 100) "at present I interpret †*Tischlingerichthys* as a probably ostariophysan because certain shared synapomorphies..."

The analysis of Arratia (1997), however, has a number of issues. The first issue concerns the approach. Arratia (1997) performed two different sets of analysis, one with a more inclusive taxon set with 32 terminal taxa, most of them represented by fossils and four fossil outgroups. The various analyses of this set differed in the inclusion of one additional fossil to test the influence of these on the resulting trees but were otherwise the same. The only ostariophysan taxa in all these analyses are †*Gordichthys*, a fossil chanid, and *Chanos*. No Otophysi were included in the analysis, and many of the traditionally used synapomorphies of Ostariophsi were not considered thus greatly limiting the significance of the result. These analyses recovered †*Tischlingerichthys* as the sister group to †*Gordichthys* plus *Chanos*.

The second set of analyses to "test the position of †*Tischlingerichthys* among ostariophysans" considered only five terminal taxa plus †*Tischlingerichthys* and a "combined" all-zero outgroup. Again, according to her own admission (p. 99), "Numerous ostariophysan characters which are unknown in fossils were not considered." In this restrictive analysis, *Tischlingerichthys* was recovered as the sister group to Gonorynchiformes or as the sister group to Otophysi.
The lack of sufficient overlap in terms of characters and taxa makes these two analyses unsuitable to evaluate the phylogenetic position of †*Tischlingerichthys*.

The second issue about the analyses of Arratia (1997) concerns the characters and their description. It is beyond the scope of this paper to discuss all characters and their problems in detail, but we note here that Patterson (1998) already commented on this issue.

We are thus not convinced by the evidence presented in Arratia (1997) that †*Tischlingerichthys* is a stem group ostariophysan (Arratia 1997: fig. 68b) or stem group chanid (Arratia 1997: fig. 68c) and conclude that a re-study of this taxon is needed before it can be used as a reliable calibration point.

**†*Santanichthys diasii*** (Silva Santos 1958)
Originally described in the genus †*Leptolepis*, as †*L. diasii*, it was subsequently transferred to the genus †*Santanichthys* and considered to be a clupeomorph. This tiny fish, rarely exceeding 30 mm standard length was restudied by Filleul & Maisey (2004) who claimed it was a member of the otophysan order Characiformes. Their conclusion was met with considerable skepticism and Malabarba & Malabarba (2010) summarized their criticism and concluded that †*Santanichthys* is a stem Ostariophysan or Otophysan. Recently, Alvarado-Ortega et al. (2025) confirmed this stating that the "alignment of *Santanichthys* with the Otophysi clade is controversial at least...and there is no reason to retain this taxon as a crown Otophysi." As in the case of †*Tischlingerichthys* a restudy and reinterpretation of the skeletal characters of this tiny fish will be necessary to place it with confidence into the teleost tree. We therefore decided not to use †*Santanichthys* as a calibration point in our study.

**†*Jianghanichthys hubeiensis*** (Lei, 1977)
An Eocene fossil from China that was originally assigned to the family Cyprinidae and subsequently considered a catostomid fossil. Liu et al. (2015) classified it in its own family Jianghanichthyidae. The analyses of Liu et al. (2015) remained inconclusive as to where this fossil belongs among Cypriniformes and we therefore did not use it as a calibration point.

**†*Sooinichthys varii*** Alvarado-Ortega, Otero & Mayrinck 2025.
A small fish of up to 97.3 mm standard length was recently described from Lower Cretaceous marine limestone deposits in Mexico (Alvarado-Ortega et al., 2025). The species was interpreted as a member of the Otophysi, a conclusion based on the reported presence of a Weberian apparatus in these heavily crushed and fractured specimens. Their condition of preservation leaves ample room for interpretation and speculation, and we disagree with the identification of many of the key bone elements. A critical reinterpretation of the crucial characters of †*Sooinichthys* is beyond the scope of this paper, but we suspect that this fish could be a clupeomorph. In favour of this hypothesis is for example the presence of structures visible at the ventral midline in Alvarado-Ortega et al. 2025: fig. 4a, that we think represent ventral scutes, one of the unique synapomorphies of clupeomorphs. Because of the drastically different interpretation of where this fish belongs phylogenetically, we refrain from using it as a calibration point.

Cope, E. D. (1873). *On the extinct Vertebrata of the Eocene of Wyoming observed by the expedition of 1872, with notes on the geology* (Sixth Report; pp. 545–649). U.S. Geological and Geographical Survey of the Territories.

Cope, E. D. (1893). Fossil fishes from British Columbia. *Proceedings of the Academy of Natural Sciences of Philadelphia*, *45*(3), 401–402.

Filleul, A., & Maisey, J. G. (2004). Redescription of *Santanichthys diasii* (Otophysi, Characiformes) from the Albian of the Santana formation and comments on its implication for otophysan relationships. *American Museum Novitates*, *3455*, 1–21.

Grande, L., & Lundberg, J. G. (1988). Revision and redescription of the genus *Astephus* (Siluriformes: Ictaluridae) with a discussion of its phylogenetic relationships. *Journal of Vertebrate Paleontology*, *8*(2), 139–171. https://doi.org/10.1080/02724634.1988.10011694

Jordan, D. S. (1923). A classification of fishes including families and generas as far as known. *Standford University Publications Biological Sciences*, *3*(2), 79–243.

Leidy, J. (1873). Notice of remains of fishes in the Bridger Tertiary Formation of Wyoming. *Proceedings of the Academy of Natural Sciences of Philadelphia*, *25*, 97–99.

Liu, J., Chang, M.-M., Wilson, M. V. H., & Murray, A. M. (2015). A new family of Cypriniformes (Teleostei, Ostariophysi) based on a redescription of † *Jianghanichthys hubeiensis* (Lei, 1977) from the Eocene Yangxi Formation of China. *Journal of Vertebrate Paleontology*, *35*(6), e1004073. https://doi.org/10.1080/02724634.2015.1004073

Liu, J., Wilson, M. V. H., & Murray, A. M. (2016). A new catostomid fish (Ostariophysi, Cypriniformes) from the Eocene Kishenehn Formation and remarks on the North American species of †*Amyzon* Cope, 1872. *Journal of Paleontology*, *90*(2), 288–304. https://doi.org/10.1017/jpa.2016.28

Lundberg, J. G. (1970). *The evolutionary history of North American catfishes, family Ictaluridae* [Ph.D Thesis]. University of Michigan.

Lundberg, J. G. (1992). The phylogeny of ictalurid catfishes: A synthesis of recent work. In R. L. Mayden (Ed.), *Systematics, Historical Ecology, and North American Freshwater Fishes* (pp. 392–420). Stanford University Press.

Malabarba, M. C., & Malabarba, L. R. (2010). Biogeography of Characiformes: An evaluation of the available information of fossil and extant taxa. In *Origin and phylogenetic interrelationships of teleosts* (pp. 317–336). Verlag Dr. Friedrich Pfeil.

Patterson, C. (1998). Comments on basal teleosts and teleostean phylogeny, by Gloria Arratia. *Copeia*, *1998*(4), 1107. https://doi.org/10.2307/1447368

Poyato-Ariza, F. J. (1996a). The phylogenetic relationships of *Rubiesichthys gregalis* and *Gordichthys conquensis* (Ostariophysi, Chanidae), from the Early Cretaceous of Spain. In *Systematics and Paleoecology* (pp. 329–348). Verlag Dr. Friedrich Pfeil.

Poyato-Ariza, F. J. (1996b). A revision of *Rubiesichthys gregalis* Wenz 1984. In *Systematics and Paleoecology* (pp. 361–368). Verlag Dr. Friedrich Pfeil.

Wenz, S. (1984). *Rubiesichthys gregalis* ng, n. Sp., Pisces Gonorhynchiformes, du Crétacé inférieur du Montsech (Province de Lérida, Espagne). *Bulletin Du Muséum National d'histoire Naturelle. Section C, Sciences de La Terre, Paléontologie, Géologie, Minéralogie*, 6(3), 275–285.

Wilson, M. V. H. (1977). *Middle Eocene freshwater fishes from British Columbia*. Life Science Contribution Royal Ontario Museum.

Zhou, J. (1990). The Cyprinidae fossils from the middle Miocene of Shanwang Basin. *Vertebrata Palasiatica*, *28*(2), 95–127.

**Supplementary Table 1.** Comparison of key divergence time estimates for major clades inferred in the present study with those reported in previously published molecular dating analyses. Node ages in the present study are reported as median estimates with 95% highest posterior density (HPD) intervals in million years (Ma).

| Node | This study | Near et al. (2012) | Rabosky et al. (2013) | Chen et al. (2013) | Betancur-R et al. (2017) | Hughes et al. (2018) | Tao et al. (2019) | Šlechtová et al. (2021) | Šlechtová et al. (2025) | Lenglin et al. (2025) |
|---|---|---|---|---|---|---|---|---|---|---|
| Ostariophysi crown age (Gonorynchiformes & Otophysi) | 193.4 (175.4-210.8) | 215.2 (198-233.0) | 146 | 163.8 (146.0-180.1) | 200.3 | 160.8(154.0-170.3) | n/a | 202 | n/a | 128 |
| Otophysi crown age (Cypriniformes, Siluriformes, Gymnotiformes, Characiformes) | 169.5 (149.7-188.3) | 175.6 (156-187.8) | 131.9 | 153.1 (136.4-169.8) | 173.3 | 147.2 (138.2-156.7) | n/a | 172.9 | n/a | 106 |
| Siluriformes, Gymnotiformes, Characiformes crown age | 136.2 (116.8-155.9) | 135.7 (121.5-150.7) | 111.9 | 136.1 (120.2-151.8) | 162.2 | 127.9 (117.9-138.0) | n/a | 115.1 | n/a | 94 |
| Cypriniformes crown age (Paedocypris as sister group to the remaining Cypriniformes) | n/a | n/a | n/a | 117.7 (94.5-138.6) | n/a | n/a | n/a | n/a | n/a | n/a |
| Cypriniformes crown age (Paedocypris, if included, not as sister group to the remaining Cypriniformes) | 112.0 (91.6-133.0) | 98.9 (82.4-118.6) | 97.8 | 96.1 (77.2-115.1 | 100 | 97.4 (85.1-116.2) | 154.5 (129.7-181.7) | 92.3 | 77.0 (59.8-100.3) | 49.3 |
| Cobitoidei crown age | 77.0 (62.9-92.1) | n/a | 73.5 | 57.4 (39.7-75.4) | 69 | 62.9 (50.0-78.6) | 85.7 (52.4-115.6) | 78.6 | 57.7 (45.7-75.2) | 23.2 |
| Cyprinoidei crown age (including Paedocypris) | 88.3 (71.6-106.6) | n/a | n/a | n/a | 83 | n/a | n/a | n/a | n/a | n/a |
| Cyprinoidei crown age (excluding Paedocypris) | 68.1 (55.9-82.1) | 72.4 (53.9-91.4) Danio&Opsariichtys | 85.2 | 78.3 (60.6-95.6) | 71.4 | 63.2 (52.3-77.7) | 59.2 (42.1-73.9) | 82.9 | n/a | 41.4 |
| Cobitoidei Botiidae crown age | 61.9 (46.2–77.9) | n/a | 39.3 | n/a | 23.4 | n/a | 49.0 (18.2-68.5) | 66.4 | 47.2 (33.9-63.1) | 16.6 |
| Cobitoidei Cobitidae crown age | 46.5 (36.8-57.4) | n/a | 56.4 | n/a | 39.8 | n/a | 42.7 (32.6-57.0) | 52.3 | 36.0 (28.7-44.4) | 15 |
| Cobitoidei Nemacheilidae crown age | 43.5 (34.8-52.7) | n/a | 51 | n/a | 36.5 | n/a | 41.6 (25.6-55.0) | 50.1 | 39.1 (30.2-50.7) | 15.8 |
| Cobitoidei Balitoridae crown age | 31.2 (24.3-39.1) | n/a | 29.1 | n/a | n/a | n/a | n/a | n/a | n/a | n/a |
| Cobitoidei Gastromyzontidae crown age | 26.9 (20.4-34.1) | n/a | 29.1 | n/a | n/a | n/a | n/a | n/a | n/a | n/a |
| Cobitoidei Vaillantellidae stem age | 72.6 (59.4-87.3) | n/a | 65.8 | 57.0 (39.3-75.4) | 65.3 | n/a | n/a | n/a | 54.6 (42.7-70.4) | 22.6 |
| Cobitoidei Ellopostomatidae stem age | 57.6 (46.5-69.5) | n/a | 63.6 | 51.5 (34.2-68.0) | n/a | n/a | n/a | n/a | 42.0 (31.6-54.9) | n/a |
| Cobitoidei Serpenticobitidae stem age | 32.9 (25.0-41.9) | n/a | 66.0 | n/a | n/a | n/a | n/a | n/a | n/a | n/a |
| Cobitoidei Barbuccidae stem age | 40.8 (31.6-50.4) | n/a | 44.7 | n/a | n/a | n/a | n/a | n/a | n/a | n/a |
| Cyprinoidei Cyprinidae crown age | 34.7 (27.5-42.2) | n/a | 44.7 | n/a | 57.7 | 25.1 (17.6-34.5) | n/a | n/a | n/a | 23 |
| Cyprinoidei Danionidae crown age | 62.7 (50.9-75.4) | n/a | 76.8 | 61.4 (44.1-79.0) | 66.3 | n/a | n/a | n/a | n/a | 34.9 |
| Cyprinoidei Sundadanionidae crown age | 47.2 (33.1-61.4) | n/a | n/a | n/a | n/a | n/a | n/a | n/a | n/a | n/a |
| Cyprinoidei Sundadanionidae stem age | 64.9 (53.0-78.9) | n/a | 66.3 | 50.4 (29.8-70.6) | 68.4 | n/a | n/a | n/a | n/a | n/a |
| Cyprinoidei Paedocyprididae crown age | 24.4 (16.1-35.2) | n/a | n/a | n/a | n/a | n/a | n/a | n/a | n/a | n/a |
| Cyprinoidei Paedocyprididae stem age | 88.4 (71.6-106.6) | n/a | n/a | 117.7 (94.5-138.6) | 83.3 | n/a | n/a | n/a | n/a | n/a |
| Cyprinoidei Xenocyprididae crown age | 19.3 (14.1-25.7) | n/a | 35.6 | n/a | 47.4 | n/a | n/a | n/a | n/a | 10.7 |
| Cyprinoidei Gobionidae crown age | 19.4 (14.5-24.9) | n/a | 37.9 | n/a | 41 | n/a | n/a | n/a | n/a | n/a |
| Cyprinoidei Leuciscidae crown age | 16.9 (13.3-21.0) | n/a | 37.4 | n/a | 32.2 | 17.0 (10.5-26.1) | n/a | n/a | n/a | n/a |
| Cyprinoidei Acheilognathidae crown age | 15.2 (10.4-20.4) | n/a | 27.1 | n/a | n/a | n/a | n/a | n/a | n/a | 22.7 |
| Cyprinoidei Leptobarbidae stem age | 51.2 (36.5-67.3) | n/a | 60.9 | 50.4 (29.8-70.6) | n/a | n/a | n/a | n/a | n/a | n/a |
| Cyprinoidei Psilorhynchidae stem age | 48.1 (34.9-64.0) | n/a | 67.4 | 62.5 (36.4-86.0) | n/a | n/a | n/a | n/a | n/a | n/a |
| Cyprinoidei Tincidae stem age | 27.9 (21.4-35.0) | n/a | 30.0 | 24.5 (15.4-39.0) | n/a | 33.9 (24.5-46.0) | n/a | n/a | n/a | 13.5 |
| Cyprinoidei Tanichthyidae stem age | 27.8 (21.3-35.0) | n/a | 30.0 | 28.6 (16.5-41.5) | 51.1 | n/a | n/a | n/a | n/a | n/a |
| Catostomoidei Catostomidae crown age | 26.3 (18.4-35.8) | 52.8 (50.7-55.0)Hypentelium&Ictiobus calibration | 42.1 | n/a | 33.1 | n/a | 62.9 (60.2-70.2) | 45.8 | 73.1 (58.3-95.0) | 13.8 |
| Catostomoidei Catostomidae stem age | 97.8 (81.0-117.5) | 98.9 (82.4-118.6) Hypentelium&Ictiobus | 92.5 | 82.6 (62.9-100.0) | 92.1 | n/a | 154.5 (129.7-181.7) | 82.9 | 17.7 (11.2-24.9) | 26.6 |
| Gyrinocheiloidei Gyrinocheilidae stem age | 112.1 (91.6-133.0) | n/a | 87.3 | 73.2 (54.0-91.5) | 100 | 97.0 (85.0-116.1) | 86.1 (52.2-116.1) | n/a | 71.7 (62.4-93.2) | 26.6 |

**Supplementary Table 2.** Species excluded from downstream analyses.

| Voucher_reads | Voucher_original | Suborder | Family | Genus | Species | Remarks |
|---|---|---|---|---|---|---|
| LR15244 | TCWC 15787.02 | Catostomoidei | Catostomidae | Catostomus | commersoni | poor model fit in kmer profiling; genome size estimates do not converge |
| LR15246 | TCWC 17181.02 | Catostomoidei | Catostomidae | Ictiobus | bubalus | high read error and poor model fit in kmer profiling; genome size estimates has a % diff > 10% between k=21 and k=31 models |
| LR15340 | LR15340 | Cobitoidei | Cobitidae | Acantopsis | sp | high read error and poor model fit in kmer profiling; genome size estimates has a % diff > 10% between k=21 and k=31 models |
| KUFOS-AN-2019.3.1 | KUFOS-AN-2019.3.1 | Cyprinoidei | Cyprinidae | Eechathalakenda | ophicephalus | high read error and poor model fit in kmer profiling; genome size estimates has a % diff > 10% between k=21 and k=31 models |
| LR15296 | LR15296 | Cyprinoidei | Cyprinidae | Enteromius | sp | high read error and poor model fit in kmer profiling; genome size estimates has a % diff > 10% between k=21 and k=31 models; unrealistic high level of heterozygozity |
| KUFOS-AN-2021.7.1 | KUFOS-AN-2021.7.1 | Cyprinoidei | Cyprinidae | Neolissochilus | wynaadensis | high read error and poor model fit in kmer profiling; genome size estimates has a % diff > 10% between k=21 and k=31 models |
| LR15108 | LR15108 | Cyprinoidei | Cyprinidae | Puntioplites | bulu | high read error and poor model fit in kmer profiling; genome size estimates has a % diff > 10% between k=21 and k=31 models |
| LR15100 | LR15100 | Cyprinoidei | Cyprinidae | Tor | sp2 | poor model fit in kmer profiling; genome size estimates do not converge |
| LR14717 | LR14717 | Cyprinoidei | Danionidae | Devario | sp | high read error and poor model fit in kmer profiling; genome size estimates has a % diff > 10% between k=21 and k=31 models; unrealistic high level of heterozygozity |
| LR12807 | LR12807 | Cyprinoidei | Danionidae | Pectenocypris | sp | generic ID doubtful; probable sample contamination with Rasbora and Pectenocypris |
| LR10789 | LR10789 | Cyprinoidei | Danionidae | Rasbora | einthovenii | high read error and poor model fit in kmer profiling; genome size estimates has a % diff > 10% between k=21 and k=31 models; unrealistic high level of heterozygozity |
| 129LR02128 | LR02128 | Cyprinoidei | Danionidae | Salmostoma | sp | generic ID doubtful: probable library contamination with Salmostoma and Mastacembelus |
| 1736_2012 | NMBE 1064463 | Cyprinoidei | Gobionidae | Gobio | gobio | high read error and poor model fit in kmer profiling; genome size estimates has a % diff > 10% between k=21 and k=31 models; unrealistic high level of heterozygozity |
| 117LR15718 | LR15718 | Cyprinoidei | Xenocyprididae | Aphyocypris | chinensis | probable hybrid sample of Nipponocypris X Aphyocypris |

**Supplementary Table 3.** Genome size estimates for species included in the k-mer profiling dataset (n = 261).

| Voucher_reads | Suborder | Family | Genus | Species | k21_Homozyg_min | k21_Homozyg_max | k21_Heterozyg_min | k21_Heterozyg_max | k21_Haplo_genome_L_min | k21_Haplo_genome_L_max |
|---|---|---|---|---|---|---|---|---|---|---|
| LR15247 | Catostomoidei | Catostomidae | Cycleptus | sp | 97.5319 | 100 | 0 | 2.46809 | 2428473807 | 2518005383 |
| LR15245 | Catostomoidei | Catostomidae | Erimyzon | sucetta | 98.0834 | 100 | 0 | 1.91661 | 1816004201 | 1844422962 |
| LR15248 | Catostomoidei | Catostomidae | Moxostoma | congestum | 97.4142 | 100 | 0 | 2.58582 | 2037414763 | 2096220823 |
| LR15059 | Catostomoidei | Catostomidae | Myxocyprinus | asiaticus | 76.0225 | 100 | 0 | 23.9775 | 2050186445 | 2657827306 |
| LR15354 | Cobitoidei | Balitoridae | Balitora | sp | 99.511 | 99.565 | 0.435012 | 0.488953 | 488971700 | 491621381 |
| LR15094 | Cobitoidei | Balitoridae | Balitoropsis | zollingeri | 99.3147 | 99.3271 | 0.672857 | 0.68534 | 415672538 | 416013807 |
| KUFOS.AN.2019.1.1 | Cobitoidei | Balitoridae | Bhavania | australis | 99.6951 | 99.7105 | 0.289492 | 0.304929 | 432042357 | 432560947 |
| LR15176 | Cobitoidei | Balitoridae | Hemimyzon | formosanum | 99.4694 | 99.4849 | 0.515056 | 0.530585 | 507154358 | 507802604 |
| LR15139 | Cobitoidei | Balitoridae | Homaloptera | sp | 99.5569 | 99.5624 | 0.43761 | 0.443147 | 448512610 | 448641647 |
| LR15140 | Cobitoidei | Balitoridae | Homalopteroides | stephensoni | 99.4466 | 99.456 | 0.543956 | 0.553415 | 487578776 | 487897402 |
| LR15093 | Cobitoidei | Balitoridae | Homalopteroides | tweediei | 98.862 | 98.8917 | 1.10833 | 1.13802 | 415991756 | 417389942 |
| LR15091 | Cobitoidei | Balitoridae | Homalopterula | gymnogaster | 97.6584 | 97.7814 | 2.21861 | 2.3416 | 346333033 | 352182664 |
| LR15172 | Cobitoidei | Balitoridae | Lepturichthys | fimbriatus | 99.2304 | 99.2394 | 0.760614 | 0.769603 | 530454951 | 530699041 |
| LR15487 | Cobitoidei | Balitoridae | Neohomaloptera | johorensis | 99.3679 | 99.4057 | 0.594324 | 0.632121 | 358710504 | 360063548 |
| LR15088 | Cobitoidei | Balitoridae | Sinogastromyzon | puliensis | 99.5779 | 99.5862 | 0.413756 | 0.42206 | 536734321 | 536974491 |
| LR15181 | Cobitoidei | Balitoridae | Travancoria | elongata | 99.7463 | 99.75 | 0.250028 | 0.253738 | 415380832 | 415461772 |
| LR15360 | Cobitoidei | Barbuccidae | Barbucca | elongatus | 99.7546 | 99.7602 | 0.239791 | 0.245442 | 379892167 | 379988941 |
| LR15330 | Cobitoidei | Botiidae | Ambastaia | sidthimunki | 98.9026 | 99.1548 | 0.845172 | 1.09738 | 728923836 | 729478747 |
| LR15149 | Cobitoidei | Botiidae | Botia | kubotai | 99.1192 | 99.366 | 0.633994 | 0.880778 | 820614521 | 821099355 |
| LR15148 | Cobitoidei | Botiidae | Chromobotia | macracanthus | 99.5741 | 99.785 | 0.214972 | 0.425872 | 821927650 | 822514382 |
| LR15074 | Cobitoidei | Botiidae | Leptobotia | elongata | 99.7893 | 99.805 | 0.194991 | 0.21073 | 628595806 | 629323510 |
| 112LR15712 | Cobitoidei | Botiidae | Parabotia | bimaculatus | 99.7501 | 99.7623 | 0.237738 | 0.249871 | 617468550 | 617867478 |
| LR15151 | Cobitoidei | Botiidae | Sinibotia | robusta | 98.6227 | 98.883 | 1.11699 | 1.3773 | 756091722 | 756601654 |
| LR15446 | Cobitoidei | Botiidae | Syncrossus | berdmorei | 98.8009 | 99.1174 | 0.882581 | 1.1991 | 651748207 | 652100027 |
| 115LR15715 | Cobitoidei | Cobitidae | Acanthopsoides | robertsi | 99.745 | 99.753 | 0.247041 | 0.25497 | 594583641 | 594818374 |
| LR15193 | Cobitoidei | Cobitidae | Bibarba | sp | 99.629 | 99.6411 | 0.358851 | 0.370985 | 527293793 | 527708351 |
| LR15575 | Cobitoidei | Cobitidae | Canthophrys | gongota | 99.6133 | 99.6193 | 0.380685 | 0.386714 | 540796198 | 540949578 |
| LR15465 | Cobitoidei | Cobitidae | Cobitis | sp | 98.4939 | 98.7959 | 1.20408 | 1.50611 | 1589711267 | 1628758447 |

**Supplementary Table 4.** Cytometric genome size estimates for cypriniform species compiled from the Animal Genome Size Database, accessed on 11 November 2024. Records were curated to retain only genera represented in the k-mer profiling dataset of the present study.

| Suborder | Family | Genus | C-value | Method | Cell type | Std sp | genomesize_MB | References |
|---|---|---|---|---|---|---|---|---|
| Catostomoidei | Catostomidae | Erimyzon | 1.86 | FCM | RBC | CCR | 1819.08 | Ferris, S.D. (1984). Tetraploidy and the evolution of the catostomid fishes. In: Evolutionary Genetics of Fishes, edited by B.J. Turner. Plenum Press, New York. 55-93. |
| Catostomoidei | Catostomidae | Moxostoma | 2.14 | FCM | RBC | CCR | 2092.92 | Ferris, S.D. (1984). Tetraploidy and the evolution of the catostomid fishes. In: Evolutionary Genetics of Fishes, edited by B.J. Turner. Plenum Press, New York. 55-93. |
| Catostomoidei | Catostomidae | Myxocyprinus | 2.02 | FD | RBC | CP | 1975.56 | Suzuki, A. (1992b). Karyotype and DNA contents of the Chinese catostomid fish, Myxocyprinius asiaticus. Chromosome Information Service 53: 25-27. |
| Catostomoidei | Catostomidae | Myxocyprinus | 2.75 | FCM | RBC | GD | 2689.5 | Zhu, D., W. Song, K. Yang, X. Cao, Y. Gul, and W. Wang (2012). Flow cytometric determination of genome size for eight commercially important fish species in China. In Vitro CellularÃ,Â &Ã,Â Developmental BiologyÃ,Â -Ã,Â Animal 48: 507-517. |
| Cobitoidei | Balitoridae | Homaloptera | 0.47 | FD | RBC | CP | 459.66 | Suzuki, A. and Y. Taki (1991). Karyological and cytochemical studies of Pseudogastromyzon myersi and Homaloptera hoffmanni in the order Cypriniformes. Chromosome Information Service 52: 32-34. |
| Cobitoidei | Botiidae | Botia | 0.78 | FCM | RBC | GD | 762.84 | Kushwaha, B., N. S. Nagpure, S. Srivastava, M. Pandey, R. Kumar, S. Raizada, S. Agarwal, M. Singh, V. S. Basheer, R. G. Kumar, P. Das, S. P. Das, S. Patnaik, A. Bit, S. K. Srivastava, A. L. Vishwakarma, C. G. Joshi, D. Kumar, and J. K. Jena (2023). Genome |
| Cobitoidei | Botiidae | Botia | 0.8 | FCM | RBC | MM | 782.4 | Ojima, Y. and K. Yamamoto (1990). Cellular DNA contents of fishes determined by flow cytometry. La Kromosomo II 57: 1871-1888. |
| Cobitoidei | Botiidae | Botia | 0.84 | BFA | RBC | SP | 821.52 | Hinegardner, R. and D.E. Rosen (1972). Cellular DNA content and the evolution of teleostean fishes. American Naturalist 106: 621-644. |
| Cobitoidei | Botiidae | Botia | 0.93 | FD | RBC | HS | 909.54 | Muramoto, J., S. Ohno, and N.B. Atkin (1968). On the diploid state of the fish order Ostariophysi. Chromosoma 24: 59-66. |
| Cobitoidei | Botiidae | Botia | 0.95 | FD | RBC | CP | 929.1 | Suzuki, A. (1992a). Chromosome and DNA studies of eight species in the family Cobitidae (Pisces, Cypriniformes). La Kromosomo II 67/68: 2275-2282. |
| Cobitoidei | Botiidae | Botia | 1.03 | FCM | RBC | MM | 1007.34 | Ojima, Y. and K. Yamamoto (1990). Cellular DNA contents of fishes determined by flow cytometry. La Kromosomo II 57: 1871-1888. |
| Cobitoidei | Botiidae | Botia | 1.21 | FCM | RBC | GD | 1183.38 | Kushwaha, B., N. S. Nagpure, S. Srivastava, M. Pandey, R. Kumar, S. Raizada, S. Agarwal, M. Singh, V. S. Basheer, R. G. Kumar, P. Das, S. P. Das, S. Patnaik, A. Bit, S. K. Srivastava, A. L. Vishwakarma, C. G. Joshi, D. Kumar, and J. K. Jena (2023). Genome |
| Cobitoidei | Botiidae | Leptobotia | 0.56 | FD | RBC | CP | 547.68 | Suzuki, A. (1992a). Chromosome and DNA studies of eight species in the family Cobitidae (Pisces, Cypriniformes). La Kromosomo II 67/68: 2275-2282. |
| Cobitoidei | Cobitidae | Cobitis | 1.4 | FD | LV | CA | 1369.2 | Park, E.-H. and C.Y. Chung (1985). Genome and nuclear sizes of Korean cobitid fishes (Teleostomi: Cypriniformes). Korean Journal of Genetics 7: 111-118. |
| Cobitoidei | Cobitidae | Cobitis | 1.42 | FCM | RBC | MM | 1388.76 | Vasil'ev, V.P., A.E. Vinogradov, Y.M. Rozanov, and E.D. Vasil'eva (1999). Cellular DNA content in different forms of the bisexual-unisexual complex of spined loaches of the genus Cobitis and in Luther's spined loach C. lutheri (Cobitidae). Journal of Icht |
| Cobitoidei | Cobitidae | Cobitis | 1.54 | FCM | RBC | HS, RT | 1506.12 | Vinogradov, A.E. (1998a). Genome size and GC-percent in vertebrates as determined by flow cytometry: the triangular relationship. Cytometry 31: 100-109. |
| Cobitoidei | Cobitidae | Cobitis | 1.58 | FCM | RBC | GD | 1545.24 | Juchno, D., B. Lackowska, A. Boron, and W. Kilarski (2010). DNA content of hepatocyte and erythrocyte nuclei of the spined loach (Cobitis taenia L.) and its polyploid forms. Fish Physiology and Biochemistry 36: 523-529. |
| Cobitoidei | Cobitidae | Cobitis | 1.69 | FD | LV | CA | 1652.82 | Park, E.-H. and C.Y. Chung (1985). Genome and nuclear sizes of Korean cobitid fishes (Teleostomi: Cypriniformes). Korean Journal of Genetics 7: 111-118. |
| Cobitoidei | Cobitidae | Cobitis | 1.76 | FCM | RBC | MM | 1721.28 | Vasil'ev, V.P., A.E. Vinogradov, Y.M. Rozanov, and E.D. Vasil'eva (1999). Cellular DNA content in different forms of the bisexual-unisexual complex of spined loaches of the genus Cobitis and in Luther's spined loach C. lutheri (Cobitidae). Journal of Icht |
| Cobitoidei | Cobitidae | Cobitis | 1.76 | FCM | RBC | MM | 1721.28 | Vasil'ev, V.P., A.E. Vinogradov, Y.M. Rozanov, and E.D. Vasil'eva (1999). Cellular DNA content in different forms of the bisexual-unisexual complex of spined loaches of the genus Cobitis and in Luther's spined loach C. lutheri (Cobitidae). Journal of Icht |
| Cobitoidei | Cobitidae | Cobitis | 1.95 | FCM | RBC | HS, RT | 1907.1 | Vinogradov, A.E. (1998a). Genome size and GC-percent in vertebrates as determined by flow cytometry: the triangular relationship. Cytometry 31: 100-109. |
| Cobitoidei | Cobitidae | Cobitis | 1.97 | FCM | RBC | MM | 1926.66 | Ojima, Y. and K. Yamamoto (1990). Cellular DNA contents of fishes determined by flow cytometry. La Kromosomo II 57: 1871-1888. |
| Cobitoidei | Cobitidae | Cobitis | 2.15 | FD | RBC | CP | 2102.7 | Suzuki, A. (1992a). Chromosome and DNA studies of eight species in the family Cobitidae (Pisces, Cypriniformes). La Kromosomo II 67/68: 2275-2282. |
| Cobitoidei | Cobitidae | Misgurnus | 1.24 | FD | LV | CA | 1212.72 | Park, E.-H. and C.Y. Chung (1985). Genome and nuclear sizes of Korean cobitid fishes (Teleostomi: Cypriniformes). Korean Journal of Genetics 7: 111-118. |
| Cobitoidei | Cobitidae | Misgurnus | 1.37 | FD | LV | CA | 1339.86 | Park, E.-H. and C.Y. Chung (1985). Genome and nuclear sizes of Korean cobitid fishes (Teleostomi: Cypriniformes). Korean Journal of Genetics 7: 111-118. |
| Cobitoidei | Cobitidae | Misgurnus | 1.4 | BFA | RBC | SP | 1369.2 | Hinegardner, R. and D.E. Rosen (1972). Cellular DNA content and the evolution of teleostean fishes. American Naturalist 106: 621-644. |
| Cobitoidei | Cobitidae | Misgurnus | 1.86 | FD | RBC | CP | 1819.08 | Suzuki, A. (1992a). Chromosome and DNA studies of eight species in the family Cobitidae (Pisces, Cypriniformes). La Kromosomo II 67/68: 2275-2282. |
| Cobitoidei | Cobitidae | Misgurnus | 2.6 | BCA | S | NS | 2542.8 | Timofeeva, M.Y. and K.A. Kaviani (1964). Nucleic acids of unfertilized eggs and developing groundling embryos. Biokimiia 29: 96-100. |
| Cobitoidei | Cobitidae | Pangio | 1 | FD | RBC | CP | 978 | Suzuki, A. (1992a). Chromosome and DNA studies of eight species in the family Cobitidae (Pisces, Cypriniformes). La Kromosomo II 67/68: 2275-2282. |
| Cobitoidei | Cobitidae | Pangio | 1.01 | FD | RBC | HS | 987.78 | Muramoto, J., S. Ohno, and N.B. Atkin (1968). On the diploid state of the fish order Ostariophysi. Chromosoma 24: 59-66. |
| Cobitoidei | Cobitidae | Pangio | 1.2 | BFA | RBC | SP | 1173.6 | Hinegardner, R. and D.E. Rosen (1972). Cellular DNA content and the evolution of teleostean fishes. American Naturalist 106: 621-644. |
| Cobitoidei | Gastromyzontidae | Pseudogastromyzon | 0.44 | FD | RBC | CP | 430.32 | Suzuki, A. and Y. Taki (1991). Karyological and cytochemical studies of Pseudogastromyzon myersi and Homaloptera hoffmanni in the order Cypriniformes. Chromosome Information Service 52: 32-34. |
| Cobitoidei | Nemacheilidae | Barbatula | 0.54 | FD | RBC | CP | 528.12 | Suzuki, A. (1992a). Chromosome and DNA studies of eight species in the family Cobitidae (Pisces, Cypriniformes). La Kromosomo II 67/68: 2275-2282. |
| Cobitoidei | Nemacheilidae | Barbatula | 0.74 | FD | LV | CA | 723.72 | Park, E.-H. and C.Y. Chung (1985). Genome and nuclear sizes of Korean cobitid fishes (Teleostomi: Cypriniformes). Korean Journal of Genetics 7: 111-118. |
| Cobitoidei | Nemacheilidae | Lefua | 0.48 | FD | RBC | CP | 469.44 | Suzuki, A. (1992a). Chromosome and DNA studies of eight species in the family Cobitidae (Pisces, Cypriniformes). La Kromosomo II 67/68: 2275-2282. |
| Cobitoidei | Nemacheilidae | Lefua | 1.2 | FD | LV | CA | 1173.6 | Park, E.-H. and C.Y. Chung (1985). Genome and nuclear sizes of Korean cobitid fishes (Teleostomi: Cypriniformes). Korean Journal of Genetics 7: 111-118. |
| Cyprinoidei | Acheilognathidae | Acheilognathus | 0.9 | FD | RBC | CP | 880.2 | Suzuki, A., Y. Taki, M. Mochizuki, and J. Hirata (1995). Chromosomal speciation in Eurasian and Japanese Cyprinidae (Pisces, Cypriniformes). Cytobios 83: 171-186. |
| Cyprinoidei | Acheilognathidae | Acheilognathus | 0.94 | FD | LV, BR | CA | 919.32 | Ojima, Y., M. Hayashi, and K. Ueno (1972). Cytogenetic studies in lower vertebrates. X. Karyotype and DNA studies in 15 species of Japanese Cyprinidae. Japanese Journal of Genetics 47: 431-440. |
| Cyprinoidei | Acheilognathidae | Acheilognathus | 0.95 | FD | RBC | CP | 929.1 | Suzuki, A., Y. Taki, M. Mochizuki, and J. Hirata (1995). Chromosomal speciation in Eurasian and Japanese Cyprinidae (Pisces, Cypriniformes). Cytobios 83: 171-186. |
| Cyprinoidei | Acheilognathidae | Acheilognathus | 0.95 | FD | RBC | CP | 929.1 | Suzuki, A., Y. Taki, M. Mochizuki, and J. Hirata (1995). Chromosomal speciation in Eurasian and Japanese Cyprinidae (Pisces, Cypriniformes). Cytobios 83: 171-186. |
| Cyprinoidei | Acheilognathidae | Acheilognathus | 1 | FD | RBC | GD | 978 | Cui, J., X. Ren, and Q. Yu (1991). Nuclear DNA content variation in fishes. Cytologia 56: 425-429. |
| Cyprinoidei | Acheilognathidae | Acheilognathus | 1.02 | FD | LV, BR | CA | 997.56 | Ojima, Y., M. Hayashi, and K. Ueno (1972). Cytogenetic studies in lower vertebrates. X. Karyotype and DNA studies in 15 species of Japanese Cyprinidae. Japanese Journal of Genetics 47: 431-440. |
| Cyprinoidei | Acheilognathidae | Acheilognathus | 1.06 | FCM | RBC | MM | 1036.68 | Ojima, Y. and K. Yamamoto (1990). Cellular DNA contents of fishes determined by flow cytometry. La Kromosomo II 57: 1871-1888. |
| Cyprinoidei | Acheilognathidae | Acheilognathus | 1.16 | FCM | RBC | MM | 1134.48 | Ojima, Y. and K. Yamamoto (1990). Cellular DNA contents of fishes determined by flow cytometry. La Kromosomo II 57: 1871-1888. |
| Cyprinoidei | Acheilognathidae | Rhodeus | 0.95 | FD | RBC | CP | 929.1 | Suzuki, A., Y. Taki, M. Mochizuki, and J. Hirata (1995). Chromosomal speciation in Eurasian and Japanese Cyprinidae (Pisces, Cypriniformes). Cytobios 83: 171-186. |
| Cyprinoidei | Acheilognathidae | Rhodeus | 1 | FD | LV, BR | CA | 978 | Ojima, Y., M. Hayashi, and K. Ueno (1972). Cytogenetic studies in lower vertebrates. X. Karyotype and DNA studies in 15 species of Japanese Cyprinidae. Japanese Journal of Genetics 47: 431-440. |
| Cyprinoidei | Acheilognathidae | Rhodeus | 1.06 | FD | RBC | HS | 1036.68 | Hafez, R., R. Labat, and R. Quillier (1978). Teneurs nuclÃ©aires en A.D.N. et relations Ã©volutives dans la famille des cyprinides (Teleostei). Bulletin de la SocietÃ© d'Histoire Naturelle de Toulouse 114: 71-84. |
| Cyprinoidei | Acheilognathidae | Rhodeus | 1.07 | FCM | RBC | GD | 1046.46 | Moreira da Costa, L., M.J. Collares-Pereira, S. Lusk, and P. Rab (1999). New data on genome size variation of Centro-European Cyprinidae (Pisces, Osteichthyes). Unpublished poster, Barcelona. |
| Cyprinoidei | Acheilognathidae | Rhodeus | 1.12 | FCM | RBC | MM | 1095.36 | Ojima, Y. and K. Yamamoto (1990). Cellular DNA contents of fishes determined by flow cytometry. La Kromosomo II 57: 1871-1888. |
| Cyprinoidei | Cyprinidae | Acrossocheilus | 1.25 | FD | RBC | NS | 1222.5 | Zan, R., Z. Song, and W. Liu (1986). Studies on karyotypes and nuclear DNA contents of some cyprinoid fishes, with notes on fish polyploids in China. In: Indo-Pacific Fish Biology, edited by T. Uyeno, R. Arai, T. Taniuchi and K. Matsuura. Ichthyological S |
| Cyprinoidei | Cyprinidae | Barbodes | 0.87 | FCM | RBC | HS, RT | 850.86 | Vinogradov, A.E. (1998a). Genome size and GC-percent in vertebrates as determined by flow cytometry: the triangular relationship. Cytometry 31: 100-109. |
| Cyprinoidei | Cyprinidae | Barbodes | 0.89 | FCM | RBC | HS, RT | 870.42 | Vinogradov, A.E. (1998a). Genome size and GC-percent in vertebrates as determined by flow cytometry: the triangular relationship. Cytometry 31: 100-109. |
| Cyprinoidei | Cyprinidae | Barbodes | 0.99 | BFA | RBC | SP | 968.22 | Hinegardner, R. and D.E. Rosen (1972). Cellular DNA content and the evolution of teleostean fishes. American Naturalist 106: 621-644. |
| Cyprinoidei | Cyprinidae | Barbodes | 1 | BFA | RBC | SP | 978 | Hinegardner, R. and D.E. Rosen (1972). Cellular DNA content and the evolution of teleostean fishes. American Naturalist 106: 621-644. |
| Cyprinoidei | Cyprinidae | Barbonymus | 1.1 | BFA | RBC | SP | 1075.8 | Hinegardner, R. and D.E. Rosen (1972). Cellular DNA content and the evolution of teleostean fishes. American Naturalist 106: 621-644. |
| Cyprinoidei | Cyprinidae | Barbonymus | 1.1 | FD | RBC | CP | 1075.8 | Suzuki, A., Y. Taki, M. Mochizuki, and J. Hirata (1995). Chromosomal speciation in Eurasian and Japanese Cyprinidae (Pisces, Cypriniformes). Cytobios 83: 171-186. |
| Cyprinoidei | Cyprinidae | Ceratogarra | 1.1 | BFA | RBC | SP | 1075.8 | Hinegardner, R. and D.E. Rosen (1972). Cellular DNA content and the evolution of teleostean fishes. American Naturalist 106: 621-644. |
| Cyprinoidei | Cyprinidae | Epalzeorhynchos | 1.22 | FIA | RBC | BS, GD, OM, RP | 1193.16 | Hardie, D.C. and P.D.N. Hebert (2003). The nucleotypic effects of cellular DNA content in cartilaginous and ray-finned fishes. Genome 46: 683-706.; Hardie, D.C. and P.D.N. Hebert (2004). Genome-size evolution in fishes. Canadian Journal of Fisheries and Aquatic Sciences 61: 1636-1646. Click here to download data appendix. |
| Cyprinoidei | Cyprinidae | Epalzeorhynchos | 1.23 | FIA | RBC | BS, GD, OM, RP | 1202.94 | Hardie, D.C. and P.D.N. Hebert (2003). The nucleotypic effects of cellular DNA content in cartilaginous and ray-finned fishes. Genome 46: 683-706.; Hardie, D.C. and P.D.N. Hebert (2004). Genome-size evolution in fishes. Canadian Journal of Fisheries and Aquatic Sciences 61: 1636-1646. Click here to download data appendix. |
| Cyprinoidei | Cyprinidae | Epalzeorhynchos | 1.3 | BFA | RBC | SP | 1271.4 | Hinegardner, R. and D.E. Rosen (1972). Cellular DNA content and the evolution of teleostean fishes. American Naturalist 106: 621-644. |
| Cyprinoidei | Cyprinidae | Garra | 1.08 | FCM | RBC | GD | 1056.24 | Kushwaha, B., N. S. Nagpure, S. Srivastava, M. Pandey, R. Kumar, S. Raizada, S. Agarwal, M. Singh, V. S. Basheer, R. G. Kumar, P. Das, S. P. Das, S. Patnaik, A. Bit, S. K. Srivastava, A. L. Vishwakarma, C. G. Joshi, D. Kumar, and J. K. Jena (2023). Genome |
| Cyprinoidei | Cyprinidae | Haludaria | 0.77 | FD | RBC | HS | 753.06 | Ohno, S., J. Muramoto, L. Christian, and N.B. Atkin (1967). Diploid-tetraploid relationship among old-world members of the fish family Cyprinidae. Chromosoma 23: 1-9. |
| Cyprinoidei | Cyprinidae | Labeo | 0.64 | FCM | RBC | GD | 625.92 | Kushwaha, B., N. S. Nagpure, S. Srivastava, M. Pandey, R. Kumar, S. Raizada, S. Agarwal, M. Singh, V. S. Basheer, R. G. Kumar, P. Das, S. P. Das, S. Patnaik, A. Bit, S. K. Srivastava, A. L. Vishwakarma, C. G. Joshi, D. Kumar, and J. K. Jena (2023). Genome |
| Cyprinoidei | Cyprinidae | Labeo | 0.95 | FCM | RBC | GD | 929.1 | Kushwaha, B., N. S. Nagpure, S. Srivastava, M. Pandey, R. Kumar, S. Raizada, S. Agarwal, M. Singh, V. S. Basheer, R. G. Kumar, P. Das, S. P. Das, S. Patnaik, A. Bit, S. K. Srivastava, A. L. Vishwakarma, C. G. Joshi, D. Kumar, and J. K. Jena (2023). Genome |
| Cyprinoidei | Cyprinidae | Labeo | 1.04 | FD | RBC | ON | 1017.12 | Patel, A., P. Das, A. Barat, and N. Sarangi (2009). Estimation of genome size in Indian major carps Labeo rohita (Hamilton), Catla catla (Hamilton), Cirrhinus mrigala (Hamilton) and Labeo calbasu (Hamilton) by Feulgen microdensitometry method. Indian Jour |
| Cyprinoidei | Cyprinidae | Labeo | 1.05 | FCM | RBC | GD | 1026.9 | Kushwaha, B., N. S. Nagpure, S. Srivastava, M. Pandey, R. Kumar, S. Raizada, S. Agarwal, M. Singh, V. S. Basheer, R. G. Kumar, P. Das, S. P. Das, S. Patnaik, A. Bit, S. K. Srivastava, A. L. Vishwakarma, C. G. Joshi, D. Kumar, and J. K. Jena (2023). Genome |
| Cyprinoidei | Cyprinidae | Labeo | 1.1 | FIA | RBC | BS, GD, OM, RP | 1075.8 | Hardie, D.C. and P.D.N. Hebert (2003). The nucleotypic effects of cellular DNA content in cartilaginous and ray-finned fishes. Genome 46: 683-706.; Hardie, D.C. and P.D.N. Hebert (2004). Genome-size evolution in fishes. Canadian Journal of Fisheries and Aquatic Sciences 61: 1636-1646. Click here to download data appendix. |
| Cyprinoidei | Cyprinidae | Labeo | 1.2 | BFA | RBC | SP | 1173.6 | Hinegardner, R. and D.E. Rosen (1972). Cellular DNA content and the evolution of teleostean fishes. American Naturalist 106: 621-644. |
| Cyprinoidei | Cyprinidae | Labeo | 1.22 | FCM | RBC | GD | 1193.16 | Kushwaha, B., N. S. Nagpure, S. Srivastava, M. Pandey, R. Kumar, S. Raizada, S. Agarwal, M. Singh, V. S. Basheer, R. G. Kumar, P. Das, S. P. Das, S. Patnaik, A. Bit, S. K. Srivastava, A. L. Vishwakarma, C. G. Joshi, D. Kumar, and J. K. Jena (2023). Genome |
| Cyprinoidei | Cyprinidae | Labeo | 1.36 | FCM | RBC | GD | 1330.08 | Kushwaha, B., N. S. Nagpure, S. Srivastava, M. Pandey, R. Kumar, S. Raizada, S. Agarwal, M. Singh, V. S. Basheer, R. G. Kumar, P. Das, S. P. Das, S. Patnaik, A. Bit, S. K. Srivastava, A. L. Vishwakarma, C. G. Joshi, D. Kumar, and J. K. Jena (2023). Genome |
| Cyprinoidei | Cyprinidae | Labeo | 1.4 | FD | RBC | HS | 1369.2 | Muramoto, J., S. Ohno, and N.B. Atkin (1968). On the diploid state of the fish order Ostariophysi. Chromosoma 24: 59-66. |
| Cyprinoidei | Cyprinidae | Labeo | 1.45 | FCM | RBC | GD | 1418.1 | Kushwaha, B., N. S. Nagpure, S. Srivastava, M. Pandey, R. Kumar, S. Raizada, S. Agarwal, M. Singh, V. S. Basheer, R. G. Kumar, P. Das, S. P. Das, S. Patnaik, A. Bit, S. K. Srivastava, A. L. Vishwakarma, C. G. Joshi, D. Kumar, and J. K. Jena (2023). Genome |
| Cyprinoidei | Cyprinidae | Labeo | 1.99 | FD | RBC | ON | 1946.22 | Patel, A., P. Das, A. Barat, and N. Sarangi (2009). Estimation of genome size in Indian major carps Labeo rohita (Hamilton), Catla catla (Hamilton), Cirrhinus mrigala (Hamilton) and Labeo calbasu (Hamilton) by Feulgen microdensitometry method. Indian Jour |
| Cyprinoidei | Cyprinidae | Labeo | 2.45 | FD | RBC | ON | 2396.1 | Patel, A., P. Das, A. Barat, and N. Sarangi (2009). Estimation of genome size in Indian major carps Labeo rohita (Hamilton), Catla catla (Hamilton), Cirrhinus mrigala (Hamilton) and Labeo calbasu (Hamilton) by Feulgen microdensitometry method. Indian Jour |
| Cyprinoidei | Cyprinidae | Paraspinibarbus | 0.98 | BFA | RBC | SP | 958.44 | Hinegardner, R. and D.E. Rosen (1972). Cellular DNA content and the evolution of teleostean fishes. American Naturalist 106: 621-644. |
| Cyprinoidei | Cyprinidae | Paraspinibarbus | 1.08 | FCM | RBC | MM | 1056.24 | Ojima, Y. and K. Yamamoto (1990). Cellular DNA contents of fishes determined by flow cytometry. La Kromosomo II 57: 1871-1888. |
| Cyprinoidei | Cyprinidae | Pethia | 0.7 | FCM | RBC | GD | 684.6 | Kushwaha, B., N. S. Nagpure, S. Srivastava, M. Pandey, R. Kumar, S. Raizada, S. Agarwal, M. Singh, V. S. Basheer, R. G. Kumar, P. Das, S. P. Das, S. Patnaik, A. Bit, S. K. Srivastava, A. L. Vishwakarma, C. G. Joshi, D. Kumar, and J. K. Jena (2023). Genome |
| Cyprinoidei | Cyprinidae | Pethia | 0.84 | FCM | RBC | HS, RT | 821.52 | Vinogradov, A.E. (1998a). Genome size and GC-percent in vertebrates as determined by flow cytometry: the triangular relationship. Cytometry 31: 100-109. |
| Cyprinoidei | Cyprinidae | Pethia | 0.91 | FCM | RBC | HS, RT | 889.98 | Vinogradov, A.E. (1998a). Genome size and GC-percent in vertebrates as determined by flow cytometry: the triangular relationship. Cytometry 31: 100-109. |
| Cyprinoidei | Cyprinidae | Pethia | 0.97 | BFA | RBC | SP | 948.66 | Hinegardner, R. and D.E. Rosen (1972). Cellular DNA content and the evolution of teleostean fishes. American Naturalist 106: 621-644. |
| Cyprinoidei | Cyprinidae | Pethia | 0.99 | FCM | RBC | MM | 968.22 | Ojima, Y. and K. Yamamoto (1990). Cellular DNA contents of fishes determined by flow cytometry. La Kromosomo II 57: 1871-1888. |
| Cyprinoidei | Cyprinidae | Poropuntius | 1.15 | FD | RBC | NS | 1124.7 | Zan, R., Z. Song, and W. Liu (1986). Studies on karyotypes and nuclear DNA contents of some cyprinoid fishes, with notes on fish polyploids in China. In: Indo-Pacific Fish Biology, edited by T. Uyeno, R. Arai, T. Taniuchi and K. Matsuura. Ichthyological S |
| Cyprinoidei | Cyprinidae | Puntigrus | 0.7 | FD | RBC | HS | 684.6 | Wolf, U., H. Ritter, N.B. Atkin, and S. Ohno (1969). Polyploidization in the fish family Cyprinidae, Order Cypriniformes. I. DNA-content and chromosome sets in various species of Cyprinidae. Humangenetik 7: 240-244. |
| Cyprinoidei | Cyprinidae | Puntigrus | 0.71 | FD | RBC | HS | 694.38 | Ohno, S., J. Muramoto, L. Christian, and N.B. Atkin (1967). Diploid-tetraploid relationship among old-world members of the fish family Cyprinidae. Chromosoma 23: 1-9. |
| Cyprinoidei | Cyprinidae | Puntigrus | 0.75 | FCM | RBC | HS, RT | 733.5 | Vinogradov, A.E. (1998a). Genome size and GC-percent in vertebrates as determined by flow cytometry: the triangular relationship. Cytometry 31: 100-109. |
| Cyprinoidei | Cyprinidae | Puntigrus | 0.96 | BFA | RBC | SP | 938.88 | Hinegardner, R. and D.E. Rosen (1972). Cellular DNA content and the evolution of teleostean fishes. American Naturalist 106: 621-644. |
| Cyprinoidei | Cyprinidae | Puntigrus | 1 | FD | RBC | CP | 978 | Suzuki, A., Y. Taki, M. Mochizuki, and J. Hirata (1995). Chromosomal speciation in Eurasian and Japanese Cyprinidae (Pisces, Cypriniformes). Cytobios 83: 171-186. |
| Cyprinoidei | Cyprinidae | Puntius | 0.71 | FCM | RBC | GD | 694.38 | Kushwaha, B., N. S. Nagpure, S. Srivastava, M. Pandey, R. Kumar, S. Raizada, S. Agarwal, M. Singh, V. S. Basheer, R. G. Kumar, P. Das, S. P. Das, S. Patnaik, A. Bit, S. K. Srivastava, A. L. Vishwakarma, C. G. Joshi, D. Kumar, and J. K. Jena (2023). Genome |
| Cyprinoidei | Cyprinidae | Rohanella | 1.2 | BFA | RBC | SP | 1173.6 | Hinegardner, R. and D.E. Rosen (1972). Cellular DNA content and the evolution of teleostean fishes. American Naturalist 106: 621-644. |
| Cyprinoidei | Cyprinidae | Sahyadria | 1.26 | FCM | RBC | GD | 1232.28 | Kushwaha, B., N. S. Nagpure, S. Srivastava, M. Pandey, R. Kumar, S. Raizada, S. Agarwal, M. Singh, V. S. Basheer, R. G. Kumar, P. Das, S. P. Das, S. Patnaik, A. Bit, S. K. Srivastava, A. L. Vishwakarma, C. G. Joshi, D. Kumar, and J. K. Jena (2023). Genome |
| Cyprinoidei | Cyprinidae | Systomus | 0.67 | FCM | RBC | GD | 655.26 | Kushwaha, B., N. S. Nagpure, S. Srivastava, M. Pandey, R. Kumar, S. Raizada, S. Agarwal, M. Singh, V. S. Basheer, R. G. Kumar, P. Das, S. P. Das, S. Patnaik, A. Bit, S. K. Srivastava, A. L. Vishwakarma, C. G. Joshi, D. Kumar, and J. K. Jena (2023). Genome |

| | | | | | | | | |
|---|---|---|---|---|---|---|---|---|
| Cyprinoidei | Cyprinidae | Systomus | 0.77 | FD | RBC | CP | 753.06 | Suzuki, A. and Y. Taki (1988). Karyotype and DNA content in the cyprinid Catlocarpio siamensis. Japanese Journal of Ichthyology 35: 389-391. |
| Cyprinoidei | Cyprinidae | Tor | 1.21 | FCM | RBC | GD | 1183.38 | Kushwaha, B., N. S. Nagpure, S. Srivastava, M. Pandey, R. Kumar, S. Raizada, S. Agarwal, M. Singh, V. S. Basheer, R. G. Kumar, P. Das, S. P. Das, S. Patnaik, A. Bit, S. K. Srivastava, A. L. Vishwakarma, C. G. Joshi, D. Kumar, and J. K. Jena (2023). Genome |
| Cyprinoidei | Danionidae | Amblypharyngodon | 1.22 | FCM | RBC | GD | 1193.16 | Kushwaha, B., N. S. Nagpure, S. Srivastava, M. Pandey, R. Kumar, S. Raizada, S. Agarwal, M. Singh, V. S. Basheer, R. G. Kumar, P. Das, S. P. Das, S. Patnaik, A. Bit, S. K. Srivastava, A. L. Vishwakarma, C. G. Joshi, D. Kumar, and J. K. Jena (2023). Genome |
| Cyprinoidei | Danionidae | Barilius | 1.22 | FCM | RBC | GD | 1193.16 | Kushwaha, B., N. S. Nagpure, S. Srivastava, M. Pandey, R. Kumar, S. Raizada, S. Agarwal, M. Singh, V. S. Basheer, R. G. Kumar, P. Das, S. P. Das, S. Patnaik, A. Bit, S. K. Srivastava, A. L. Vishwakarma, C. G. Joshi, D. Kumar, and J. K. Jena (2023). Genome |
| Cyprinoidei | Danionidae | Danio | 1.42 | FD | RBC | CP | 1388.76 | Suzuki, A., Y. Taki, M. Mochizuki, and J. Hirata (1995). Chromosomal speciation in Eurasian and Japanese Cyprinidae (Pisces, Cypriniformes). Cytobios 83: 171-186. |
| Cyprinoidei | Danionidae | Danio | 1.68 | FCM | RC | HS | 1643.04 | Ciudad, J., E. Cid, A. Velasco, J.M. Lara, J. AijÃ³n, and A. Orfao (2002). Flow cytometry measurement of the DNA contents of G0/G1 diploid cells from three different teleost fish species. Cytometry 48: 20-25. |
| Cyprinoidei | Danionidae | Danio | 1.75 | FCM | RBC | HS, RT | 1711.5 | Vinogradov, A.E. (1998a). Genome size and GC-percent in vertebrates as determined by flow cytometry: the triangular relationship. Cytometry 31: 100-109. |
| Cyprinoidei | Danionidae | Danio | 1.78 | FCM | RBC | MM | 1740.84 | Ojima, Y. and K. Yamamoto (1990). Cellular DNA contents of fishes determined by flow cytometry. La Kromosomo II 57: 1871-1888. |
| Cyprinoidei | Danionidae | Danio | 1.8 | BFA | RBC | SP | 1760.4 | Hinegardner, R. (1968). Evolution of cellular DNA content in teleost fishes. American Naturalist 102: 517-523.; Hinegardner, R. and D.E. Rosen (1972). Cellular DNA content and the evolution of teleostean fishes. American Naturalist 106: 621-644. |
| Cyprinoidei | Danionidae | Devario | 0.79 | FCM | RBC | GD | 772.62 | Kushwaha, B., N. S. Nagpure, S. Srivastava, M. Pandey, R. Kumar, S. Raizada, S. Agarwal, M. Singh, V. S. Basheer, R. G. Kumar, P. Das, S. P. Das, S. Patnaik, A. Bit, S. K. Srivastava, A. L. Vishwakarma, C. G. Joshi, D. Kumar, and J. K. Jena (2023). Genome |
| Cyprinoidei | Danionidae | Devario | 1.4 | FIA | RBC | BS, GD, OM, RP | 1369.2 | Hardie, D.C. and P.D.N. Hebert (2004). Genome-size evolution in fishes. Canadian Journal of Fisheries and Aquatic Sciences 61: 1636-1646. Click here to download data appendix. |
| Cyprinoidei | Danionidae | Laubuka | 1.6 | BFA | RBC | SP | 1564.8 | Hinegardner, R. (1968). Evolution of cellular DNA content in teleost fishes. American Naturalist 102: 517-523.; Hinegardner, R. and D.E. Rosen (1972). Cellular DNA content and the evolution of teleostean fishes. American Naturalist 106: 621-644. |
| Cyprinoidei | Danionidae | Opsarius | 1.22 | FCM | RBC | GD | 1193.16 | Kushwaha, B., N. S. Nagpure, S. Srivastava, M. Pandey, R. Kumar, S. Raizada, S. Agarwal, M. Singh, V. S. Basheer, R. G. Kumar, P. Das, S. P. Das, S. Patnaik, A. Bit, S. K. Srivastava, A. L. Vishwakarma, C. G. Joshi, D. Kumar, and J. K. Jena (2023). Genome |
| Cyprinoidei | Danionidae | Rasbora | 0.85 | FCM | RBC | GD | 831.3 | Kushwaha, B., N. S. Nagpure, S. Srivastava, M. Pandey, R. Kumar, S. Raizada, S. Agarwal, M. Singh, V. S. Basheer, R. G. Kumar, P. Das, S. P. Das, S. Patnaik, A. Bit, S. K. Srivastava, A. L. Vishwakarma, C. G. Joshi, D. Kumar, and J. K. Jena (2023). Genome |
| Cyprinoidei | Danionidae | Rasbora | 1.44 | FD | RBC | CP | 1408.32 | Suzuki, A., Y. Taki, M. Mochizuki, and J. Hirata (1995). Chromosomal speciation in Eurasian and Japanese Cyprinidae (Pisces, Cypriniformes). Cytobios 83: 171-186. |
| Cyprinoidei | Gobionidae | Hemibarbus | 0.95 | FD | RBC | CP | 929.1 | Suzuki, A., Y. Taki, M. Mochizuki, and J. Hirata (1995). Chromosomal speciation in Eurasian and Japanese Cyprinidae (Pisces, Cypriniformes). Cytobios 83: 171-186. |
| Cyprinoidei | Gobionidae | Hemibarbus | 1.05 | FD | LV | CA | 1026.9 | Kang, Y.S. and E.H. Park (1973). Studies on the karyotypes and comparative DNA values in several Korean cyprinid fishes. Korean Journal of Zoology 16: 97-108. |
| Cyprinoidei | Gobionidae | Hemibarbus | 1.08 | FD | LV | CA | 1056.24 | Kang, Y.S. and E.H. Park (1973). Studies on the karyotypes and comparative DNA values in several Korean cyprinid fishes. Korean Journal of Zoology 16: 97-108. |
| Cyprinoidei | Gobionidae | Hemibarbus | 1.15 | FD | RBC | GD | 1124.7 | Cui, J., X. Ren, and Q. Yu (1991). Nuclear DNA content variation in fishes. Cytologia 56: 425-429. |
| Cyprinoidei | Gobionidae | Hemibarbus | 1.17 | FD | RBC | GD | 1144.26 | Cui, J., X. Ren, and Q. Yu (1991). Nuclear DNA content variation in fishes. Cytologia 56: 425-429. |
| Cyprinoidei | Gobionidae | Hemibarbus | 1.28 | FCM | RBC | MM | 1251.84 | Ojima, Y. and K. Yamamoto (1990). Cellular DNA contents of fishes determined by flow cytometry. La Kromosomo II 57: 1871-1888. |
| Cyprinoidei | Gobionidae | Sarcocheilichthys | 1.2 | FD | RBC | CP | 1173.6 | Suzuki, A., Y. Taki, M. Mochizuki, and J. Hirata (1995). Chromosomal speciation in Eurasian and Japanese Cyprinidae (Pisces, Cypriniformes). Cytobios 83: 171-186. |
| Cyprinoidei | Gobionidae | Sarcocheilichthys | 1.21 | FD | LV, BR | CA | 1183.38 | Ojima, Y., M. Hayashi, and K. Ueno (1972). Cytogenetic studies in lower vertebrates. X. Karyotype and DNA studies in 15 species of Japanese Cyprinidae. Japanese Journal of Genetics 47: 431-440. |
| Cyprinoidei | Gobionidae | Sarcocheilichthys | 1.21 | FD | RBC | CP | 1183.38 | Zhou, T. (1984). Chromosome studies in Chinese freshwater fishes [in Chinese]. Zoological Research (Tung Wu Hsueh Yen Chiu) 5: 38-51. |
| Cyprinoidei | Gobionidae | Sarcocheilichthys | 1.25 | FD | RBC | HS | 1222.5 | Li, Y., K. Li, and T. Zhou (1983). Cellular DNA content of fourteen species of freshwater fishes [in Chinese]. Acta Genetica Sinica 10: 384-389. |
| Cyprinoidei | Gobionidae | Sarcocheilichthys | 1.39 | FCM | RBC | MM | 1359.42 | Ojima, Y. and K. Yamamoto (1990). Cellular DNA contents of fishes determined by flow cytometry. La Kromosomo II 57: 1871-1888. |
| Cyprinoidei | Gobionidae | Sarcocheilichthys | 1.54 | FD | RBC | GD | 1506.12 | Cui, J., X. Ren, and Q. Yu (1991). Nuclear DNA content variation in fishes. Cytologia 56: 425-429. |
| Cyprinoidei | Gobionidae | Squalidus | 1.25 | FD | RBC | CP | 1222.5 | Suzuki, A., Y. Taki, M. Mochizuki, and J. Hirata (1995). Chromosomal speciation in Eurasian and Japanese Cyprinidae (Pisces, Cypriniformes). Cytobios 83: 171-186. |
| Cyprinoidei | Leuciscidae | Chrosomus | 1.27 | FCM | RBC | GD | 1242.06 | Goddard, K.A., O. Megwinoff, L.L. Wessner, and F. Giaimo (1998). Confirmation of gynogenesis in Phoxinus eos-neogaeus (Pisces: Cyprinidae). Journal of Heredity 89: 151-157. |
| Cyprinoidei | Leuciscidae | Chrosomus | 1.32 | FD | RBC | GD | 1290.96 | Gold, J.R. and C.T. Amemiya (1987). Genome size variation in North American minnows (Cyprinidae). II. Variation among 20 species. Genome 29: 481-489. |
| Cyprinoidei | Leuciscidae | Chrosomus | 1.41 | FIA | RBC | BS, GD, OM, RP | 1378.98 | Hardie, D.C. and P.D.N. Hebert (2003). The nucleotypic effects of cellular DNA content in cartilaginous and ray-finned fishes. Genome 46: 683-706.; Hardie, D.C. and P.D.N. Hebert (2004). Genome-size evolution in fishes. Canadian Journal of Fisheries and Aquatic Sciences 61: 1636-1646. Click here to download data appendix. |
| Cyprinoidei | Leuciscidae | Chrosomus | 1.42 | FCM | RBC | GD | 1388.76 | Goddard, K.A., O. Megwinoff, L.L. Wessner, and F. Giaimo (1998). Confirmation of gynogenesis in Phoxinus eos-neogaeus (Pisces: Cyprinidae). Journal of Heredity 89: 151-157. |
| Cyprinoidei | Leuciscidae | Chrosomus | 1.57 | FCM | RBC | GD | 1535.46 | Goddard, K.A., O. Megwinoff, L.L. Wessner, and F. Giaimo (1998). Confirmation of gynogenesis in Phoxinus eos-neogaeus (Pisces: Cyprinidae). Journal of Heredity 89: 151-157. |
| Cyprinoidei | Leuciscidae | Chrosomus | 1.64 | FCM | RBC | GD | 1603.92 | Dawley, R.M. and K.A. Goddard (1988). Diploid-triploid mosaics among unisexual hybrids of the minnows Phoxinus eos and Phoxinus neogaeus. Evolution 42: 649-659. |
| Cyprinoidei | Leuciscidae | Chrosomus | 1.75 | FCM | RBC | GD | 1711.5 | Dawley, R.M. and K.A. Goddard (1988). Diploid-triploid mosaics among unisexual hybrids of the minnows Phoxinus eos and Phoxinus neogaeus. Evolution 42: 649-659. |
| Cyprinoidei | Leuciscidae | Chrosomus | 1.83 | FCM | RBC | GD | 1789.74 | Dawley, R.M. and K.A. Goddard (1988). Diploid-triploid mosaics among unisexual hybrids of the minnows Phoxinus eos and Phoxinus neogaeus. Evolution 42: 649-659. |
| Cyprinoidei | Leuciscidae | Chrosomus | 2.13 | FCM | RBC | GD | 2083.14 | Goddard, K.A., O. Megwinoff, L.L. Wessner, and F. Giaimo (1998). Confirmation of gynogenesis in Phoxinus eos-neogaeus (Pisces: Cyprinidae). Journal of Heredity 89: 151-157. |
| Cyprinoidei | Leuciscidae | Chrosomus | 2.14 | FIA | RBC | BS, GD, OM, RP | 2092.92 | Hardie, D.C. and P.D.N. Hebert (2003). The nucleotypic effects of cellular DNA content in cartilaginous and ray-finned fishes. Genome 46: 683-706.; Hardie, D.C. and P.D.N. Hebert (2004). Genome-size evolution in fishes. Canadian Journal of Fisheries and Aquatic Sciences 61: 1636-1646. Click here to download data appendix. |
| Cyprinoidei | Leuciscidae | Chrosomus | 2.49 | FCM | RBC | GD | 2435.22 | Dawley, R.M. and K.A. Goddard (1988). Diploid-triploid mosaics among unisexual hybrids of the minnows Phoxinus eos and Phoxinus neogaeus. Evolution 42: 649-659. |
| Cyprinoidei | Leuciscidae | Cyprinella | 1.03 | FD | RBC | GD | 1007.34 | Gold, J.R., C.J. Ragland, and L.J. Schliesing (1990). Genome size variation and evolution in North American cyprinid fishes. Genetics, Selection, Evolution 22: 11-29. |
| Cyprinoidei | Leuciscidae | Cyprinella | 1.14 | FD | RBC | GD | 1114.92 | Gold, J.R., C.J. Ragland, and L.J. Schliesing (1990). Genome size variation and evolution in North American cyprinid fishes. Genetics, Selection, Evolution 22: 11-29. |
| Cyprinoidei | Leuciscidae | Cyprinella | 1.19 | FD | RBC | GD | 1163.82 | Gold, J.R. and H.J. Price (1985). Genome size variation among North American minnows (Cyprinidae). I. Distribution of the variation in five species. Heredity 54: 297-305. |
| Cyprinoidei | Leuciscidae | Cyprinella | 1.2 | FD | RBC | GD | 1173.6 | Gold, J.R., C.J. Ragland, and L.J. Schliesing (1990). Genome size variation and evolution in North American cyprinid fishes. Genetics, Selection, Evolution 22: 11-29. |
| Cyprinoidei | Leuciscidae | Cyprinella | 1.21 | FD | RBC | GD | 1183.38 | Gold, J.R. and H.J. Price (1985). Genome size variation among North American minnows (Cyprinidae). I. Distribution of the variation in five species. Heredity 54: 297-305. |
| Cyprinoidei | Leuciscidae | Cyprinella | 1.25 | FD | RBC | GD | 1222.5 | Gold, J.R. and C.T. Amemiya (1987). Genome size variation in North American minnows (Cyprinidae). II. Variation among 20 species. Genome 29: 481-489. |
| Cyprinoidei | Leuciscidae | Cyprinella | 1.27 | FD | RBC | GD | 1242.06 | Gold, J.R., C.J. Ragland, and L.J. Schliesing (1990). Genome size variation and evolution in North American cyprinid fishes. Genetics, Selection, Evolution 22: 11-29. |
| Cyprinoidei | Leuciscidae | Cyprinella | 1.45 | FD | RBC | GD | 1418.1 | Gold, J.R., C.J. Ragland, and L.J. Schliesing (1990). Genome size variation and evolution in North American cyprinid fishes. Genetics, Selection, Evolution 22: 11-29. |
| Cyprinoidei | Leuciscidae | Dionda | 0.99 | FCM | RBC | GD, CP | 968.22 | Gold, J.R., Y. Li, M.C. Birkner, and J.D. Jenkin (1992). Chromosomal NOR karyotypes and genome sizes in Dionda (Osteichthyes: Cyprinidae) from Texas and New Mexico. Southwestern Naturalist 37: 217-222. |
| Cyprinoidei | Leuciscidae | Dionda | 1 | FCM | RBC | GD, CP | 978 | Gold, J.R., Y. Li, M.C. Birkner, and J.D. Jenkin (1992). Chromosomal NOR karyotypes and genome sizes in Dionda (Osteichthyes: Cyprinidae) from Texas and New Mexico. Southwestern Naturalist 37: 217-222. |
| Cyprinoidei | Leuciscidae | Dionda | 1.01 | FCM | RBC | GD, CP | 987.78 | Gold, J.R., Y. Li, M.C. Birkner, and J.D. Jenkin (1992). Chromosomal NOR karyotypes and genome sizes in Dionda (Osteichthyes: Cyprinidae) from Texas and New Mexico. Southwestern Naturalist 37: 217-222. |
| Cyprinoidei | Leuciscidae | Dionda | 1.04 | FCM | RBC | GD, CP | 1017.12 | Gold, J.R., Y. Li, M.C. Birkner, and J.D. Jenkin (1992). Chromosomal NOR karyotypes and genome sizes in Dionda (Osteichthyes: Cyprinidae) from Texas and New Mexico. Southwestern Naturalist 37: 217-222. |
| Cyprinoidei | Leuciscidae | Dionda | 1.04 | FCM | RBC | GD, CP | 1017.12 | Gold, J.R., Y. Li, M.C. Birkner, and J.D. Jenkin (1992). Chromosomal NOR karyotypes and genome sizes in Dionda (Osteichthyes: Cyprinidae) from Texas and New Mexico. Southwestern Naturalist 37: 217-222. |
| Cyprinoidei | Leuciscidae | Dionda | 1.07 | FD | RBC | GD | 1046.46 | Gold, J.R., C.J. Ragland, and L.J. Schliesing (1990). Genome size variation and evolution in North American cyprinid fishes. Genetics, Selection, Evolution 22: 11-29. |
| Cyprinoidei | Leuciscidae | Hybognathus | 1.41 | FD | RBC | GD | 1378.98 | Gold, J.R., C.J. Ragland, and L.J. Schliesing (1990). Genome size variation and evolution in North American cyprinid fishes. Genetics, Selection, Evolution 22: 11-29. |
| Cyprinoidei | Leuciscidae | Leuciscus | 1.19 | SCF | RBC | HS | 1163.82 | De Smet, W.H.O. (1981). The nuclear Feulgen-DNA content of the vertebrates (especially reptiles), as measured by fluorescence cytophotometry, with notes on the cell and chromosome size. Acta Zoologica et Pathologica Antverpiensia 76: 119-167. |
| Cyprinoidei | Leuciscidae | Leuciscus | 1.24 | FCM | RBC | GD | 1212.72 | Moreira da Costa, L., M.J. Collares-Pereira, S. Lusk, and P. Rab (1999). New data on genome size variation of Centro-European Cyprinidae (Pisces, Osteichthyes). Unpublished poster, Barcelona. |
| Cyprinoidei | Leuciscidae | Leuciscus | 1.28 | FD | RBC | HS | 1251.84 | Hafez, R., R. Labat, and R. Quillier (1978). Teneurs nuclÃ©aires en A.D.N. et relations Ã©volutives dans la famille des cyprinides (Teleostei). Bulletin de la SocietÃ© d'Histoire Naturelle de Toulouse 114: 71-84. |
| Cyprinoidei | Leuciscidae | Leuciscus | 1.3 | FCM | RBC | GD | 1271.4 | Moreira da Costa, L., M.J. Collares-Pereira, S. Lusk, and P. Rab (1999). New data on genome size variation of Centro-European Cyprinidae (Pisces, Osteichthyes). Unpublished poster, Barcelona. |
| Cyprinoidei | Leuciscidae | Leuciscus | 1.5 | BFA | RBC | SP | 1467 | Hinegardner, R. and D.E. Rosen (1972). Cellular DNA content and the evolution of teleostean fishes. American Naturalist 106: 621-644. |
| Cyprinoidei | Leuciscidae | Notropis | 1.04 | FD | RBC | GD | 1017.12 | Gold, J.R., C.J. Ragland, and L.J. Schliesing (1990). Genome size variation and evolution in North American cyprinid fishes. Genetics, Selection, Evolution 22: 11-29. |
| Cyprinoidei | Leuciscidae | Notropis | 1.1 | FD | RBC | GD | 1075.8 | Gold, J.R. and C.T. Amemiya (1987). Genome size variation in North American minnows (Cyprinidae). II. Variation among 20 species. Genome 29: 481-489. |
| Cyprinoidei | Leuciscidae | Notropis | 1.13 | FD | RBC | GD | 1105.14 | Gold, J.R., C.J. Ragland, and L.J. Schliesing (1990). Genome size variation and evolution in North American cyprinid fishes. Genetics, Selection, Evolution 22: 11-29. |
| Cyprinoidei | Leuciscidae | Notropis | 1.17 | FD | RBC | GD | 1144.26 | Gold, J.R. and C.T. Amemiya (1987). Genome size variation in North American minnows (Cyprinidae). II. Variation among 20 species. Genome 29: 481-489. |
| Cyprinoidei | Leuciscidae | Notropis | 1.18 | FD | RBC | GD | 1154.04 | Gold, J.R., C.J. Ragland, and L.J. Schliesing (1990). Genome size variation and evolution in North American cyprinid fishes. Genetics, Selection, Evolution 22: 11-29. |
| Cyprinoidei | Leuciscidae | Notropis | 1.19 | FD | RBC | GD | 1163.82 | Gold, J.R. and C.T. Amemiya (1987). Genome size variation in North American minnows (Cyprinidae). II. Variation among 20 species. Genome 29: 481-489. |
| Cyprinoidei | Leuciscidae | Notropis | 1.19 | FD | RBC | GD | 1163.82 | Gold, J.R. and C.T. Amemiya (1987). Genome size variation in North American minnows (Cyprinidae). II. Variation among 20 species. Genome 29: 481-489. |
| Cyprinoidei | Leuciscidae | Notropis | 1.19 | FD | RBC | GD | 1163.82 | Gold, J.R., C.J. Ragland, and L.J. Schliesing (1990). Genome size variation and evolution in North American cyprinid fishes. Genetics, Selection, Evolution 22: 11-29. |
| Cyprinoidei | Leuciscidae | Notropis | 1.23 | FD | RBC | GD | 1202.94 | Gold, J.R., C.J. Ragland, and L.J. Schliesing (1990). Genome size variation and evolution in North American cyprinid fishes. Genetics, Selection, Evolution 22: 11-29. |
| Cyprinoidei | Leuciscidae | Notropis | 1.24 | FD | RBC | GD | 1212.72 | Gold, J.R., C.J. Ragland, and L.J. Schliesing (1990). Genome size variation and evolution in North American cyprinid fishes. Genetics, Selection, Evolution 22: 11-29. |
| Cyprinoidei | Leuciscidae | Notropis | 1.25 | FD | RBC | GD | 1222.5 | Gold, J.R., C.J. Ragland, and L.J. Schliesing (1990). Genome size variation and evolution in North American cyprinid fishes. Genetics, Selection, Evolution 22: 11-29. |
| Cyprinoidei | Leuciscidae | Notropis | 1.26 | FD | RBC | GD | 1232.28 | Gold, J.R. and C.T. Amemiya (1987). Genome size variation in North American minnows (Cyprinidae). II. Variation among 20 species. Genome 29: 481-489. |
| Cyprinoidei | Leuciscidae | Notropis | 1.26 | FD | RBC | GD | 1232.28 | Gold, J.R., C.J. Ragland, and L.J. Schliesing (1990). Genome size variation and evolution in North American cyprinid fishes. Genetics, Selection, Evolution 22: 11-29. |
| Cyprinoidei | Leuciscidae | Notropis | 1.27 | FD | RBC | GD | 1242.06 | Gold, J.R., C.J. Ragland, and L.J. Schliesing (1990). Genome size variation and evolution in North American cyprinid fishes. Genetics, Selection, Evolution 22: 11-29. |
| Cyprinoidei | Leuciscidae | Notropis | 1.36 | FD | RBC | GD | 1330.08 | Gold, J.R. and C.T. Amemiya (1987). Genome size variation in North American minnows (Cyprinidae). II. Variation among 20 species. Genome 29: 481-489. |
| Cyprinoidei | Leuciscidae | Notropis | 1.37 | FD | RBC | GD | 1339.86 | Gold, J.R., C.J. Ragland, and L.J. Schliesing (1990). Genome size variation and evolution in North American cyprinid fishes. Genetics, Selection, Evolution 22: 11-29. |
| Cyprinoidei | Leuciscidae | Notropis | 1.4 | FD | RBC | GD | 1369.2 | Gold, J.R., C.J. Ragland, and L.J. Schliesing (1990). Genome size variation and evolution in North American cyprinid fishes. Genetics, Selection, Evolution 22: 11-29. |
| Cyprinoidei | Leuciscidae | Notropis | 1.45 | FD | RBC | GD | 1418.1 | Gold, J.R., C.J. Ragland, and L.J. Schliesing (1990). Genome size variation and evolution in North American cyprinid fishes. Genetics, Selection, Evolution 22: 11-29. |
| Cyprinoidei | Leuciscidae | Phoxinus | 1.15 | FD | RBC | HS | 1124.7 | Hafez, R., R. Labat, and R. Quillier (1978). Teneurs nuclÃ©aires en A.D.N. et relations Ã©volutives dans la famille des cyprinides (Teleostei). Bulletin de la SocietÃ© d'Histoire Naturelle de Toulouse 114: 71-84. |
| Cyprinoidei | Leuciscidae | Ptychocheilus | 1.26 | FCM | RBC | GD, CP | 1232.28 | Gold, J.R. (1994). Cytosystematic evidence that the genus Richardsonius belongs in the western clade of phoxinin cyprinids. Copeia 1994: 815-818. |
| Cyprinoidei | Leuciscidae | Ptychocheilus | 1.29 | FCM | RBC | GD, CP | 1261.62 | Gold, J.R. (1994). Cytosystematic evidence that the genus Richardsonius belongs in the western clade of phoxinin cyprinids. Copeia 1994: 815-818. |
| Cyprinoidei | Leuciscidae | Ptychocheilus | 1.38 | FCM | RBC | GD, CP | 1349.64 | Gold, J.R. (1994). Cytosystematic evidence that the genus Richardsonius belongs in the western clade of phoxinin cyprinids. Copeia 1994: 815-818. |
| Cyprinoidei | Leuciscidae | Rhinichthys | 1.29 | FD | RBC | GD | 1261.62 | Gold, J.R., C.J. Ragland, and L.J. Schliesing (1990). Genome size variation and evolution in North American cyprinid fishes. Genetics, Selection, Evolution 22: 11-29. |
| Cyprinoidei | Leuciscidae | Rhynchocypris | 1.02 | FD | LV | CA | 997.56 | Kang, Y.S. and E.H. Park (1973). Studies on the karyotypes and comparative DNA values in several Korean cyprinid fishes. Korean Journal of Zoology 16: 97-108. |
| Cyprinoidei | Leuciscidae | Rhynchocypris | 1.11 | FD | LV | CA | 1085.58 | Kang, Y.S. and E.H. Park (1973). Studies on the karyotypes and comparative DNA values in several Korean cyprinid fishes. Korean Journal of Zoology 16: 97-108. |
| Cyprinoidei | Leuciscidae | Rhynchocypris | 1.52 | FCM | RBC | MM | 1486.56 | Ojima, Y. and K. Yamamoto (1990). Cellular DNA contents of fishes determined by flow cytometry. La Kromosomo II 57: 1871-1888. |
| Cyprinoidei | Leuciscidae | Rutilus | 0.97 | FD | RBC | HS | 948.66 | Hafez, R., R. Labat, and R. Quillier (1978). Teneurs nuclÃ©aires en A.D.N. et relations Ã©volutives dans la famille des cyprinides (Teleostei). Bulletin de la SocietÃ© d'Histoire Naturelle de Toulouse 114: 71-84. |
| Cyprinoidei | Leuciscidae | Rutilus | 0.98 | FD | RBC | HS | 958.44 | Wolf, U., H. Ritter, N.B. Atkin, and S. Ohno (1969). Polyploidization in the fish family Cyprinidae, Order Cypriniformes. I. DNA-content and chromosome sets in various species of Cyprinidae. Humangenetik 7: 240-244. |
| Cyprinoidei | Leuciscidae | Rutilus | 1.08 | BCA | RBC | GD | 1056.24 | Vendrely, R. and C. Vendrely (1953). Argenine an deoxyribonucleic acid content of erythrocyte nuclei and sperms of some species of fishes. Nature 172: 30-31.; Vendrely, R. and C. Vendrely (1952a). Sur la teneur comparÃ©e en arginine et en acide dÃ©soxyribonuclÃ©ique des noyaux d'Ã©rythrocytes de quelques espÃ¨ces de pois |
| Cyprinoidei | Leuciscidae | Rutilus | 1.11 | BCA | RBC | HS | 1085.58 | Vendrely, R. and C. Vendrely (1950). Sur la teneur absolue en acide dÃ©soxyribonuclÃ©ique du noyau cellulaire chez quelques espÃ¨ces d'oiseaux et de poissons. Comptes Rendus de l'AcadÃ©mie des Sciences 230: 788-790. |
| Cyprinoidei | Leuciscidae | Rutilus | 1.21 | FCM | RBC | GD | 1183.38 | Moreira da Costa, L., M.J. Collares-Pereira, S. Lusk, and P. Rab (1999). New data on genome size variation of Centro-European Cyprinidae (Pisces, Osteichthyes). Unpublished poster, Barcelona. |
| Cyprinoidei | Leuciscidae | Rutilus | 1.32 | FCM | RBC | HS, RT | 1290.96 | Vinogradov, A.E. (1998a). Genome size and GC-percent in vertebrates as determined by flow cytometry: the triangular relationship. Cytometry 31: 100-109. |
| Cyprinoidei | Leuciscidae | Rutilus | 1.51 | SCF | RBC | HS | 1476.78 | De Smet, W.H.O. (1981). The nuclear Feulgen-DNA content of the vertebrates (especially reptiles), as measured by fluorescence cytophotometry, with notes on the cell and chromosome size. Acta Zoologica et Pathologica Antverpiensia 76: 119-167. |

| | | | | | | | | |
|---|---|---|---|---|---|---|---|---|
| Cyprinoidei | Leuciscidae | Scardinius | 0.94 | BCA | RBC | NS | 919.32 | Mauro, M.L. and G. Micheli (1979). DNA reassociation kinetics in diploid and phylogenetically tetraploid cyprinidae. Journal of Experimental Zoology 208: 407-416. |
| Cyprinoidei | Leuciscidae | Scardinius | 0.98 | FD | RBC | HS | 958.44 | Hafez, R., R. Labat, and R. Quillier (1978). Teneurs nuclÃ©aires en A.D.N. et relations Ã©volutives dans la famille des cyprinides (Teleostei). Bulletin de la SocietÃ© d'Histoire Naturelle de Toulouse 114: 71-84. |
| Cyprinoidei | Leuciscidae | Scardinius | 1.41 | FCM | RBC | GD | 1378.98 | Moreira da Costa, L., M.J. Collares-Pereira, S. Lusk, and P. Rab (1999). New data on genome size variation of Centro-European Cyprinidae (Pisces, Osteichthyes). Unpublished poster, Barcelona. |
| Cyprinoidei | Leuciscidae | Scardinius | 1.71 | FD | RBC | CP | 1672.38 | Suzuki, A., Y. Taki, M. Mochizuki, and J. Hirata (1995). Chromosomal speciation in Eurasian and Japanese Cyprinidae (Pisces, Cypriniformes). Cytobios 83: 171-186. |
| Cyprinoidei | Leuciscidae | Semotilus | 1.23 | FIA | RBC | BS, GD, OM, RP | 1202.94 | Hardie, D.C. and P.D.N. Hebert (2003). The nucleotypic effects of cellular DNA content in cartilaginous and ray-finned fishes. Genome 46: 683-706.; Hardie, D.C. and P.D.N. Hebert (2004). Genome-size evolution in fishes. Canadian Journal of Fisheries and Aquatic Sciences 61: 1636-1646. Click here to download data appendix. |
| Cyprinoidei | Leuciscidae | Semotilus | 1.26 | FD | RBC | GD | 1232.28 | Gold, J.R. and C.T. Amemiya (1987). Genome size variation in North American minnows (Cyprinidae). II. Variation among 20 species. Genome 29: 481-489. |
| Cyprinoidei | Leuciscidae | Squalius | 1.18 | FCM | RBC | GD | 1154.04 | Collares-Pereira, M.J. and L. Moreira da Costa (1999). Intraspecific and interspecific genome size variation in Iberian Cyprinidae and the problem of diploidy and polyploidy, with review of genome sizes within the family. Folia Zoologica 48: 61-76. |
| Cyprinoidei | Leuciscidae | Squalius | 1.21 | FCM | RBC | GD | 1183.38 | Collares-Pereira, M.J. and L. Moreira da Costa (1999). Intraspecific and interspecific genome size variation in Iberian Cyprinidae and the problem of diploidy and polyploidy, with review of genome sizes within the family. Folia Zoologica 48: 61-76. |
| Cyprinoidei | Leuciscidae | Squalius | 1.21 | FCM | RBC | GD | 1183.38 | Collares-Pereira, M.J. and L. Moreira da Costa (1999). Intraspecific and interspecific genome size variation in Iberian Cyprinidae and the problem of diploidy and polyploidy, with review of genome sizes within the family. Folia Zoologica 48: 61-76. |
| Cyprinoidei | Leuciscidae | Squalius | 1.22 | FCM | RBC | GD | 1193.16 | Collares-Pereira, M.J. and L. Moreira da Costa (1999). Intraspecific and interspecific genome size variation in Iberian Cyprinidae and the problem of diploidy and polyploidy, with review of genome sizes within the family. Folia Zoologica 48: 61-76. |
| Cyprinoidei | Leuciscidae | Squalius | 1.23 | FCM | RBC | GD | 1202.94 | Prospero, M.I. and M.J. Collares-Pereira (2000). Nuclear DNA content variation in the diploid-polyploid Leuciscus alburnoides complex (Teleostei, Cyprinidae) assessed by flow cytometry. Folia Zoologica 49: 53-58. |
| Cyprinoidei | Leuciscidae | Squalius | 1.27 | BCA | RBC | NS | 1242.06 | Mauro, M.L. and G. Micheli (1979). DNA reassociation kinetics in diploid and phylogenetically tetraploid cyprinidae. Journal of Experimental Zoology 208: 407-416. |
| Cyprinoidei | Leuciscidae | Squalius | 1.33 | FD | RBC | HS | 1300.74 | Hafez, R., R. Labat, and R. Quillier (1978). Teneurs nuclÃ©aires en A.D.N. et relations Ã©volutives dans la famille des cyprinides (Teleostei). Bulletin de la SocietÃ© d'Histoire Naturelle de Toulouse 114: 71-84. |
| Cyprinoidei | Leuciscidae | Squalius | 1.33 | FD | RBC | HS | 1300.74 | Wolf, U., H. Ritter, N.B. Atkin, and S. Ohno (1969). Polyploidization in the fish family Cyprinidae, Order Cypriniformes. I. DNA-content and chromosome sets in various species of Cyprinidae. Humangenetik 7: 240-244. |
| Cyprinoidei | Tincidae | Tinca | 0.81 | BCA | RBC | NS | 792.18 | Mauro, M.L. and G. Micheli (1979). DNA reassociation kinetics in diploid and phylogenetically tetraploid cyprinidae. Journal of Experimental Zoology 208: 407-416. |
| Cyprinoidei | Tincidae | Tinca | 0.85 | FD | RBC | TT | 831.3 | Fontana, F. (1976). Nuclear DNA content and cytometry of erythrocytes of Huso huso L., Acipenser sturio L. and Acipenser naccarii Bonaparte. Caryologia 29: 127-137. |
| Cyprinoidei | Tincidae | Tinca | 0.93 | NS | RBC | CP | 909.54 | Gerzeli, G., C. Casati, and A. Meneghetti Gennaro (1956). I volumi nucleari e cellulari in relazione al tenore di acido desossiribonucleico in eritrociti di alcune specie di Vertebrati. Rivista di Istochimica Normale e Patologica 2: 149-155. |
| Cyprinoidei | Tincidae | Tinca | 0.95 | NS | NS | GD | 929.1 | Vialli, M. (1957a). Volume et contenu en ADN par noyau. Experimental Cell Research Suppl. 4: 284-293. |
| Cyprinoidei | Tincidae | Tinca | 0.97 | BCA | RBC | GD | 948.66 | Vendrely, R. and C. Vendrely (1953). Argenine an deoxyribonucleic acid content of erythrocyte nuclei and sperms of some species of fishes. Nature 172: 30-31.; Vendrely, R. and C. Vendrely (1952a). Sur la teneur comparÃ©e en arginine et en acide dÃ©soxyribonuclÃ©ique des noyaux d'Ã©rythrocytes de quelques espÃ¨ces de pois |
| Cyprinoidei | Tincidae | Tinca | 0.98 | BCA | S | BT | 958.44 | Vendrely, R. and C. Vendrely (1952b). Sur la teneur individuelle en arginine des spermatozoÃ¯des comparÃ©e Ã la teneur individuelle en arginine des noyaux d'Ã©rythrocytes chez quelques espÃ¨ces de poissons. Comptes Rendus de l'AcadÃ©mie des Sciences 235: |
| Cyprinoidei | Tincidae | Tinca | 1.01 | FIA | RBC | GD | 987.78 | Bytyutskyy, D. and M. Flajshans (2014). Use of diploid and triploid tench (Tinca tinca) blood as standards for genome size measurements. Journal of Applied Ichthyology 30 (Suppl. 1): 12-14. |
| Cyprinoidei | Tincidae | Tinca | 1.02 | FIA | RBC | GD | 997.56 | Bytyutskyy, D., J. Srp, and M. Flajshans (2012). Use of Feulgen image analysis densitometry to study the effect of genome size on nuclear size in polyploid sturgeons. Journal of Applied Ichthyology 28: 704-708. |
| Cyprinoidei | Tincidae | Tinca | 1.04 | FD | RBC | HS | 1017.12 | Hafez, R., R. Labat, and R. Quillier (1978). Teneurs nuclÃ©aires en A.D.N. et relations Ã©volutives dans la famille des cyprinides (Teleostei). Bulletin de la SocietÃ© d'Histoire Naturelle de Toulouse 114: 71-84. |
| Cyprinoidei | Tincidae | Tinca | 1.05 | BCA | RBC | HS | 1026.9 | Vendrely, R. and C. Vendrely (1950). Sur la teneur absolue en acide dÃ©soxyribonuclÃ©ique du noyau cellulaire chez quelques espÃ¨ces d'oiseaux et de poissons. Comptes Rendus de l'AcadÃ©mie des Sciences 230: 788-790. |
| Cyprinoidei | Tincidae | Tinca | 1.05 | FD | RBC | HS | 1026.9 | Wolf, U., H. Ritter, N.B. Atkin, and S. Ohno (1969). Polyploidization in the fish family Cyprinidae, Order Cypriniformes. I. DNA-content and chromosome sets in various species of Cyprinidae. Humangenetik 7: 240-244. |
| Cyprinoidei | Tincidae | Tinca | 1.13 | FCM | RBC | GD | 1105.14 | Collares-Pereira, M.J. and L. Moreira da Costa (1999). Intraspecific and interspecific genome size variation in Iberian Cyprinidae and the problem of diploidy and polyploidy, with review of genome sizes within the family. Folia Zoologica 48: 61-76. |
| Cyprinoidei | Xenocyprididae | Hemiculter | 1.22 | FD | RBC | GD | 1193.16 | Cui, J., X. Ren, and Q. Yu (1991). Nuclear DNA content variation in fishes. Cytologia 56: 425-429. |
| Cyprinoidei | Xenocyprididae | Opsariichthys | 1.15 | FCM | RBC | MM | 1124.7 | Ojima, Y. and K. Yamamoto (1990). Cellular DNA contents of fishes determined by flow cytometry. La Kromosomo II 57: 1871-1888. |
| Cyprinoidei | Xenocyprididae | Opsariichthys | 1.67 | FD | LV, BR | CA | 1633.26 | Ojima, Y., M. Hayashi, and K. Ueno (1972). Cytogenetic studies in lower vertebrates. X. Karyotype and DNA studies in 15 species of Japanese Cyprinidae. Japanese Journal of Genetics 47: 431-440. |
| Gyrinocheiloidei | Gyrinocheilidae | Gyrinocheilus | 0.51 | FD | RBC | CP | 498.78 | Arai, R., A. Suzuki, and Y. Akai (1988). The karyotype and DNA value of a cypriniform algae eater, Gyrinocheilus aymonieri. Japanese Journal of Ichthyology 34: 515-517. |
| Gyrinocheiloidei | Gyrinocheilidae | Gyrinocheilus | 0.61 | FCM | RBC | MM | 596.58 | Ojima, Y. and K. Yamamoto (1990). Cellular DNA contents of fishes determined by flow cytometry. La Kromosomo II 57: 1871-1888. |
| Gyrinocheiloidei | Gyrinocheilidae | Gyrinocheilus | 0.62 | FIA | RBC | BS, GD, OM, RP | 606.36 | Hardie, D.C. and P.D.N. Hebert (2003). The nucleotypic effects of cellular DNA content in cartilaginous and ray-finned fishes. Genome 46: 683-706.; Hardie, D.C. and P.D.N. Hebert (2004). Genome-size evolution in fishes. Canadian Journal of Fisheries and Aquatic Sciences 61: 1636-1646. Click here to download data appendix. |
| Gyrinocheiloidei | Gyrinocheilidae | Gyrinocheilus | 0.65 | BFA | RBC | SP | 635.7 | Hinegardner, R. (1968). Evolution of cellular DNA content in teleost fishes. American Naturalist 102: 517-523.; Hinegardner, R. and D.E. Rosen (1972). Cellular DNA content and the evolution of teleostean fishes. American Naturalist 106: 621-644. |

**Supplementary Table 5.** Comparative dataset of 309 cypriniform species used in the phylogenetic comparative analyses of miniaturization.

| Voucher_reads | Suborder | Family | Genus | Species | Source | Genome_assembly_Mb_NCBI | Ploidy | Miniature | SL.max.cm | mean_k31_genomesize_MB |
|---|---|---|---|---|---|---|---|---|---|---|
| LR15247 | Catostomoidei | Catostomidae | Cycleptus | sp | This study | NA | polyploid | non-miniature | 66.5 | 2509.817213 |
| LR15245 | Catostomoidei | Catostomidae | Erimyzon | sucetta | This study | NA | polyploid | non-miniature | 31.4 | 1788.172173 |
| LR15248 | Catostomoidei | Catostomidae | Moxostoma | congestum | This study | NA | polyploid | non-miniature | 41.1 | 2017.530905 |
| GCF_019703515.2 | Catostomoidei | Catostomidae | Myxocyprinus | asiaticus | NCBI | NA | polyploid | non-miniature | 50 | 2309.856512 |
| LR15354 | Cobitoidei | Balitoridae | Balitora | sp | This study | NA | diploid | non-miniature | 7.7 | 492.1509565 |
| LR15094 | Cobitoidei | Balitoridae | Balitoropsis | zollingeri | This study | NA | diploid | non-miniature | 6.77 | 416.506414 |
| KUFOS.AN.2019.1.1 | Cobitoidei | Balitoridae | Bhavania | australis | This study | NA | diploid | non-miniature | 9 | 432.345153 |
| LR15176 | Cobitoidei | Balitoridae | Hemimyzon | formosanum | This study | NA | diploid | non-miniature | 10 | 509.5217015 |
| LR15139 | Cobitoidei | Balitoridae | Homaloptera | sp | This study | NA | diploid | non-miniature | 13 | 450.377603 |
| LR15140 | Cobitoidei | Balitoridae | Homalopteroides | stephensoni | This study | NA | diploid | non-miniature | 6.5 | 488.628322 |
| LR15093 | Cobitoidei | Balitoridae | Homalopteroides | tweediei | This study | NA | diploid | proportioned-dwarf | 4 | 413.243012 |
| LR15091 | Cobitoidei | Balitoridae | Homalopterula | gymnogaster | This study | NA | diploid | non-miniature | 6.6 | 347.959084 |
| LR15172 | Cobitoidei | Balitoridae | Lepturichthys | fimbriatus | This study | NA | diploid | non-miniature | 16.1 | 530.93438 |
| LR15487 | Cobitoidei | Balitoridae | Neohomaloptera | johorensis | This study | NA | diploid | proportioned-dwarf | 2.05 | 357.9961275 |
| LR15088 | Cobitoidei | Balitoridae | Sinogastromyzon | puliensis | This study | NA | diploid | non-miniature | 7.45 | 537.5911635 |
| LR15181 | Cobitoidei | Balitoridae | Travancoria | elongata | This study | NA | diploid | non-miniature | 11.4 | 419.2010845 |
| LR15360 | Cobitoidei | Barbuccidae | Barbucca | elongatus | This study | NA | diploid | proportioned-dwarf | 2.7 | 379.578087 |
| LR15330 | Cobitoidei | Botiidae | Ambastaia | sidthimunki | This study | NA | polyploid | non-miniature | 5.5 | 728.1004125 |
| GCA_036877575.1 | Cobitoidei | Botiidae | Botia | almorhae | NCBI | 894.1 | polyploid | non-miniature | 15.5 | 894.1 |
| LR15149 | Cobitoidei | Botiidae | Botia | kubotai | This study | NA | polyploid | non-miniature | 8.49 | 820.033105 |
| LR15148 | Cobitoidei | Botiidae | Chromobotia | macracanthus | This study | NA | polyploid | non-miniature | 25 | 821.307172 |
| LR15074 | Cobitoidei | Botiidae | Leptobotia | elongata | This study | NA | diploid | non-miniature | 50 | 626.037003 |
| 112LR15712 | Cobitoidei | Botiidae | Parabotia | bimaculatus | This study | NA | diploid | non-miniature | 15.8 | 615.909513 |
| LR15151 | Cobitoidei | Botiidae | Sinibotia | robusta | This study | NA | polyploid | non-miniature | 6.63 | 755.627618 |
| LR15446 | Cobitoidei | Botiidae | Syncrossus | berdmorei | This study | NA | polyploid | non-miniature | 11.1 | 650.9152395 |
| GCA_036877545.1 | Cobitoidei | Botiidae | Yasuhikotakia | modesta | NCBI | 636.8 | polyploid | non-miniature | 25 | 636.8 |
| 115LR15715 | Cobitoidei | Cobitidae | Acanthopsoides | robertsi | This study | NA | diploid | non-miniature | 5.5 | 595.835145 |
| LR15193 | Cobitoidei | Cobitidae | Bibarba | sp | This study | NA | diploid | non-miniature | 5.5 | 527.2919795 |
| LR15575 | Cobitoidei | Cobitidae | Canthophrys | gongota | This study | NA | diploid | non-miniature | 6.51 | 540.609599 |
| LR15465 | Cobitoidei | Cobitidae | Cobitis | sp | This study | NA | diploid | non-miniature | 10.9 | 1612.45102 |
| LR15170 | Cobitoidei | Cobitidae | Kottelatlimia | katik | This study | NA | diploid | proportioned-dwarf | 1.35 | 1043.902574 |
| 114LR15714 | Cobitoidei | Cobitidae | Kottelatlimia | pristes | This study | NA | diploid | non-miniature | 4 | 892.8929045 |
| LR15467 | Cobitoidei | Cobitidae | Lepidocephalichthys | sp | This study | NA | diploid | non-miniature | 6.28 | 656.0133925 |
| GCF_027580225.1 | Cobitoidei | Cobitidae | Misgurnus | anguillicaudatus | NCBI | 1104 | diploid | non-miniature | 28 | 1104 |
| GCA_025389145.1 | Cobitoidei | Cobitidae | Misgurnus | mizolepis | NCBI | 1112 | diploid | non-miniature | 16.7 | 1112 |
| BNHS.FWF.981 | Cobitoidei | Cobitidae | Pangio | bhujia | This study | NA | diploid | proportioned-dwarf | 2.6 | 676.7185705 |
| LR15141 | Cobitoidei | Cobitidae | Pangio | sp | This study | NA | diploid | non-miniature | 6.62 | 813.269303 |
| LR15609 | Cobitoidei | Cobitidae | Sabanejewia | balcanica | This study | NA | diploid | non-miniature | 9 | 1039.00824 |
| LR15146 | Cobitoidei | Cobitidae | Theriodes | sandakanensis | This study | NA | diploid | non-miniature | 3.5 | 487.8762655 |
| 141LR15748 | Cobitoidei | Ellopostomatidae | Ellopostoma | mystax | This study | NA | diploid | non-miniature | 5.53 | 452.711028 |
| LR15180 | Cobitoidei | Gastromyzontidae | Annamia | sp | This study | NA | diploid | non-miniature | 7.91 | 417.169605 |
| GCA_019155185.1 | Cobitoidei | Gastromyzontidae | Beaufortia | kweichowensis | NCBI | NA | diploid | non-miniature | 5.4 | 462.685559 |
| LR15491 | Cobitoidei | Gastromyzontidae | Erromyzon | sp | This study | NA | diploid | non-miniature | 5.2 | 446.541696 |
| LR15090 | Cobitoidei | Gastromyzontidae | Glaniopsis | muliradiatus | This study | NA | diploid | non-miniature | 6 | 472.8609795 |
| LR15133 | Cobitoidei | Gastromyzontidae | Katibasia | insidiosa | This study | NA | diploid | non-miniature | 3.4 | 427.0149425 |
| LR15175 | Cobitoidei | Gastromyzontidae | Liniparhomaloptera | disparis | This study | NA | diploid | non-miniature | 6.3 | 431.0346715 |
| LR15476 | Cobitoidei | Gastromyzontidae | Plesiomyzon | baotingensis | This study | NA | diploid | non-miniature | 3.94 | 428.9316375 |

| | | | | | | | | | | | |
|---|---|---|---|---|---|---|---|---|---|---|---|
| LR15425 | Cobitoidei | Gastromyzontidae | Protomyzon | borneensis | This study | NA | | diploid | non-miniature | 5.4 | 416.6464905 |
| LR15067 | Cobitoidei | Gastromyzontidae | Pseudogastromyzon | sp | This study | NA | | diploid | non-miniature | 8.2 | 438.344733 |
| LR15561 | Cobitoidei | Gastromyzontidae | Sewellia | elongata | This study | NA | | diploid | non-miniature | 7.3 | 411.1789435 |
| 113LR15713 | Cobitoidei | Gastromyzontidae | Yaoshania | pachychilus | This study | NA | | diploid | non-miniature | 5.8 | 435.496029 |
| LR15343 | Cobitoidei | Nemacheilidae | Aborichthys | elongatus | This study | NA | | diploid | non-miniature | 7.4 | 518.9789115 |
| GCA_947034865.1 | Cobitoidei | Nemacheilidae | Barbatula | barbatula | NCBI | | 617.6 | diploid | non-miniature | 12 | 617.6 |
| LR15190 | Cobitoidei | Nemacheilidae | Barbatula | sp | This study | NA | | diploid | non-miniature | 7 | 647.9074045 |
| LR15073 | Cobitoidei | Nemacheilidae | Homatula | variegata | This study | NA | | diploid | non-miniature | 12.4 | 697.3544255 |
| KUFOS.AN.2019.2.1 | Cobitoidei | Nemacheilidae | Indoreonectes | keralensis | This study | NA | | diploid | non-miniature | 4.65 | 393.362597 |
| LR15462 | Cobitoidei | Nemacheilidae | Lefua | sp | This study | NA | | diploid | non-miniature | 5.3 | 551.812393 |
| KUFOS.AN.2021.1.1 | Cobitoidei | Nemacheilidae | Mesonoemacheilus | triangularis | This study | NA | | diploid | non-miniature | 5.8 | 493.90045 |
| LR15337 | Cobitoidei | Nemacheilidae | Micronemacheilus | sp | This study | NA | | diploid | non-miniature | 5.8 | 551.353682 |
| LR15459 | Cobitoidei | Nemacheilidae | Nemacheilus | selangoricus | This study | NA | | diploid | non-miniature | 4.8 | 475.761951 |
| LR15611 | Cobitoidei | Nemacheilidae | Neonoemacheilus | sp | This study | NA | | diploid | non-miniature | 7.32 | 611.2010315 |
| LR15490 | Cobitoidei | Nemacheilidae | Oreonectes | platycephalus | This study | NA | | diploid | non-miniature | 4.38 | 606.3790165 |
| LR15504 | Cobitoidei | Nemacheilidae | Paracanthocobitis | botia | This study | NA | | diploid | non-miniature | 5.77 | 550.526874 |
| LR15189 | Cobitoidei | Nemacheilidae | Petruichthys | brevis | This study | NA | | diploid | non-miniature | 3.21 | 573.6448505 |
| LR15152 | Cobitoidei | Nemacheilidae | Schistura | fasciolata | This study | NA | | diploid | non-miniature | 6 | 697.99144 |
| LR15454 | Cobitoidei | Nemacheilidae | Schistura | sp1 | This study | NA | | diploid | non-miniature | 5.4 | 521.701272 |
| LR15496 | Cobitoidei | Nemacheilidae | Schistura | sp2 | This study | NA | | diploid | non-miniature | 4.2 | 689.9526065 |
| LR15455 | Cobitoidei | Nemacheilidae | Sundoreonectes | sabanus | This study | NA | | diploid | non-miniature | 5.41 | 655.1661915 |
| LR15072 | Cobitoidei | Nemacheilidae | Traccatichthys | pulcher | This study | NA | | diploid | non-miniature | 5.39 | 580.90548 |
| GCF_015846415.1 | Cobitoidei | Nemacheilidae | Triplophysa | dalaica | NCBI | | 607.9 | diploid | non-miniature | 8.2 | 607.9 |
| GCF_024868665.1 | Cobitoidei | Nemacheilidae | Triplophysa | rosa | NCBI | | 684.8 | diploid | non-miniature | 5.6 | 684.8 |
| GCA_006030095.1 | Cobitoidei | Nemacheilidae | Triplophysa | siluroides | NCBI | | 583.4 | diploid | non-miniature | 50 | 583.4 |
| GCA_008369825.1 | Cobitoidei | Nemacheilidae | Triplophysa | tibetana | NCBI | | 652.9 | diploid | non-miniature | 13.9 | 652.9 |
| GCA_033220385.1 | Cobitoidei | Nemacheilidae | Triplophysa | yarkandensis | NCBI | | 520.8 | diploid | non-miniature | 17.7 | 520.8 |
| LR15357 | Cobitoidei | Serpenticobitidae | Serpenticobitis | zonata | This study | NA | | diploid | non-miniature | 4.3 | 440.308335 |
| LR15334 | Cobitoidei | Vaillantellidae | Vaillantella | maassii | This study | NA | | diploid | non-miniature | 12.5 | 427.556523 |
| LR15562 | Cyprinoidei | Acheilognathidae | Acheilognathus | deignani | This study | NA | | diploid | non-miniature | 5.2 | 752.796849 |
| LR15063 | Cyprinoidei | Acheilognathidae | Acheilognathus | sp | This study | NA | | diploid | non-miniature | 5.2 | 830.2436615 |
| LR15636 | Cyprinoidei | Acheilognathidae | Paratanakia | himantegus | This study | NA | | diploid | non-miniature | 6.3 | 703.096636 |
| LR15661 | Cyprinoidei | Acheilognathidae | Rhodeus | amarus | This study | NA | | diploid | non-miniature | 9.5 | 840.9927755 |
| GCA_037043235.1 | Cyprinoidei | Acheilognathidae | Rhodeus | uyekii | NCBI | | 894.5 | diploid | non-miniature | 5 | 894.5 |
| LR15266 | Cyprinoidei | Cyprinidae | Acapoeta | tanganicae | This study | NA | | polyploid | non-miniature | 29 | 1296.278517 |
| GCA_039880705.1 | Cyprinoidei | Cyprinidae | Acrossocheilus | fasciatus | NCBI | NA | | diploid | non-miniature | 19.2 | 937.795839 |
| LR15615 | Cyprinoidei | Cyprinidae | Ageneiogarra | sp | This study | NA | | diploid | non-miniature | 34.5 | 1156.287975 |
| GCA_023376895.1 | Cyprinoidei | Cyprinidae | Aspiorhynchus | laticeps | NCBI | | 1583 | polyploid | non-miniature | 100 | 1583 |
| LR15639 | Cyprinoidei | Cyprinidae | Balantiocheilos | melanopterus | This study | NA | | diploid | non-miniature | 35 | 738.167276 |
| LR15079 | Cyprinoidei | Cyprinidae | Barbichthys | laevis | This study | NA | | diploid | non-miniature | 30 | 887.40424 |
| LR15102 | Cyprinoidei | Cyprinidae | Barbodes | lateristriga | This study | NA | | diploid | non-miniature | 7 | 740.2849255 |
| 120LR15721 | Cyprinoidei | Cyprinidae | Barbodes | sellifer | This study | NA | | diploid | non-miniature | 9.8 | 778.466779 |
| LR15256 | Cyprinoidei | Cyprinidae | Barboides | gracilis | This study | NA | | diploid | proportioned-dwarf | 1.8 | 636.29766 |
| EA0124 | Cyprinoidei | Cyprinidae | Barbonymus | altus | This study | NA | | diploid | non-miniature | 15 | 809.7455375 |
| GCA_936440315.1 | Cyprinoidei | Cyprinidae | Barbus | barbus | NCBI | | 1584 | polyploid | non-miniature | 90 | 1584 |
| GCF_003368295.1 | Cyprinoidei | Cyprinidae | Carassius | auratus | NCBI | | 1820 | polyploid | non-miniature | 26 | 1820 |
| GCF_963082965.1 | Cyprinoidei | Cyprinidae | Carassius | carassius | NCBI | | 1684 | polyploid | non-miniature | 55 | 1684 |
| GCA_023724075.1 | Cyprinoidei | Cyprinidae | Carassius | gibelio | NCBI | | 1949 | polyploid | non-miniature | 35 | 1949 |

| | | | | | | | | | | | |
|---|---|---|---|---|---|---|---|---|---|---|---|
| LR15111 | Cyprinoidei | Cyprinidae | Ceratogarra | cambodgiensis | This study | NA | | diploid | non-miniature | 10 | 847.3830295 |
| EA0401 | Cyprinoidei | Cyprinidae | Chagunius | sp | This study | NA | | diploid | non-miniature | 25 | 720.7493135 |
| LR15270 | Cyprinoidei | Cyprinidae | Clypeobarbus | sp | This study | NA | | diploid | non-miniature | 3.3 | 829.8340545 |
| LR10955 | Cyprinoidei | Cyprinidae | Crossocheilus | reticulatus | This study | NA | | diploid | non-miniature | 17 | 1296.106392 |
| LR10341 | Cyprinoidei | Cyprinidae | Cyclocheilichthys | apogon | This study | NA | | diploid | non-miniature | 16 | 833.4889075 |
| GCF_018340385.1 | Cyprinoidei | Cyprinidae | Cyprinus | carpio | NCBI | | 1680 | polyploid | non-miniature | 40 | 1680 |
| KUFOS.AN.2021.4.1 | Cyprinoidei | Cyprinidae | Dawkinsia | filamentosa | This study | NA | | diploid | non-miniature | 12 | 638.710795 |
| 140LR15737 | Cyprinoidei | Cyprinidae | Desmopuntius | sp | This study | NA | | diploid | non-miniature | 3 | 765.758192 |
| 122LR15725 | Cyprinoidei | Cyprinidae | Discherodontus | ashmeadi | This study | NA | | diploid | non-miniature | 13.6 | 723.054833 |
| 119LR15720 | Cyprinoidei | Cyprinidae | Eirmotus | sp1 | This study | NA | | diploid | non-miniature | 3.6 | 648.622376 |
| LR09582 | Cyprinoidei | Cyprinidae | Eirmotus | sp2 | This study | NA | | diploid | non-miniature | 3.6 | 658.8349075 |
| GCA_034779775.1 | Cyprinoidei | Cyprinidae | Enteromius | afrohamiltoni | NCBI | | 663.5 | diploid | non-miniature | 17.5 | 663.5 |
| GCA_032359395.1 | Cyprinoidei | Cyprinidae | Enteromius | anoplus | NCBI | | 737.7 | diploid | non-miniature | 10 | 737.7 |
| GCA_032357425.1 | Cyprinoidei | Cyprinidae | Enteromius | argenteus | NCBI | | 681.6 | diploid | non-miniature | 17.5 | 681.6 |
| GCA_032359235.1 | Cyprinoidei | Cyprinidae | Enteromius | mattozi | NCBI | | 727.4 | diploid | non-miniature | 40 | 727.4 |
| GCA_032357445.1 | Cyprinoidei | Cyprinidae | Enteromius | radiatus | NCBI | | 616.7 | diploid | non-miniature | 12 | 616.7 |
| LR15606 | Cyprinoidei | Cyprinidae | Enteromius | sp1 | This study | NA | | diploid | non-miniature | 15 | 859.7960585 |
| GCA_032359435.1 | Cyprinoidei | Cyprinidae | Enteromius | unitaeniatus | NCBI | NA | | diploid | non-miniature | 17.5 | 878.386854 |
| LR01655 | Cyprinoidei | Cyprinidae | Epalzeorhynchos | kalopterum | This study | NA | | diploid | non-miniature | 12 | 1362.220601 |
| LR15563 | Cyprinoidei | Cyprinidae | Folifer | brevifilis | This study | NA | | diploid | non-miniature | 19.2 | 900.587643 |
| KUFOS.RR.2022.P.4 | Cyprinoidei | Cyprinidae | Garra | periyarensis | This study | NA | | diploid | non-miniature | 16 | 967.001756 |
| GCA_027564155.1 | Cyprinoidei | Cyprinidae | Gymnocypris | eckloni-scoliostomus | NCBI | | 948.1 | polyploid | non-miniature | 39.5 | 948.1 |
| LR15635 | Cyprinoidei | Cyprinidae | Haludaria | fasciata | This study | NA | | diploid | non-miniature | 6 | 590.2574145 |
| LR15078 | Cyprinoidei | Cyprinidae | Hampala | ampalong | This study | NA | | diploid | non-miniature | 15 | 667.0398945 |
| KUFOS.RR.2022.P.3 | Cyprinoidei | Cyprinidae | Hypselobarbus | periyarensis | This study | NA | | polyploid | non-miniature | 40 | 752.678211 |
| GCA_012976165.1 | Cyprinoidei | Cyprinidae | Labeo | catla | NCBI | | 1019 | diploid | non-miniature | 120 | 1019 |
| GCF_022985175.1 | Cyprinoidei | Cyprinidae | Labeo | rohita | NCBI | | 1126 | diploid | non-miniature | 150 | 1126 |
| LR15278 | Cyprinoidei | Cyprinidae | Labeo | cylindricus | This study | NA | | diploid | non-miniature | 25 | 1028.751666 |
| GCA_034780575.1 | Cyprinoidei | Cyprinidae | Labeobarbus | marequensis | NCBI | NA | | polyploid | non-miniature | 35 | 1263.354553 |
| LR12884 | Cyprinoidei | Cyprinidae | Labiobarbus | ocellatus | This study | NA | | diploid | non-miniature | 15.5 | 1143.454791 |
| KUFOS.RR.2022.P.2 | Cyprinoidei | Cyprinidae | Lepidopygopsis | typus | This study | NA | | polyploid | non-miniature | 20 | 1406.545258 |
| LR15104 | Cyprinoidei | Cyprinidae | Lobocheilos | aureolineatus | This study | NA | | diploid | non-miniature | 5.99 | 1011.286788 |
| LR15565 | Cyprinoidei | Cyprinidae | Mystacoleucus | argenteus | This study | NA | | diploid | non-miniature | 15 | 683.1196445 |
| LR15612 | Cyprinoidei | Cyprinidae | Neolissochilus | sp | This study | NA | | polyploid | non-miniature | 60 | 857.157747 |
| 143LR15682 | Cyprinoidei | Cyprinidae | Oliotius | oligolepis | This study | NA | | diploid | non-miniature | 5 | 605.885024 |
| GCF_012432095.1 | Cyprinoidei | Cyprinidae | Onychostoma | macrolepis | NCBI | | 886.6 | diploid | non-miniature | 26.5 | 886.6 |
| EA1291 | Cyprinoidei | Cyprinidae | Oreichthys | sp | This study | NA | | diploid | non-miniature | 3 | 679.6391495 |
| LR09367 | Cyprinoidei | Cyprinidae | Osteobrama | cunma | This study | NA | | diploid | non-miniature | 12 | 718.0802345 |
| LR15096 | Cyprinoidei | Cyprinidae | Osteochilus | vittatoides | This study | NA | | diploid | non-miniature | 11.9 | 1105.531316 |
| GCA_003573665.1 | Cyprinoidei | Cyprinidae | Oxygymnocypris | stewartii | NCBI | | 1849 | polyploid | non-miniature | 54.6 | 1849 |
| LR01656 | Cyprinoidei | Cyprinidae | Paracrossochilus | sp | This study | NA | | diploid | non-miniature | 9.5 | 1281.936214 |
| LR15618 | Cyprinoidei | Cyprinidae | Paraspinibarbus | sp | This study | NA | | diploid | non-miniature | 40 | 823.683848 |
| KUFOS.AN.2021.5.1 | Cyprinoidei | Cyprinidae | Pethia | conchonius | This study | NA | | diploid | non-miniature | 5.6 | 601.786245 |
| LR06584 | Cyprinoidei | Cyprinidae | Pethia | sp1 | This study | NA | | diploid | non-miniature | 6 | 686.5376455 |
| LR15646 | Cyprinoidei | Cyprinidae | Pethia | sp2 | This study | NA | | diploid | non-miniature | 4.12 | 683.0203955 |
| GCA_004124795.1 | Cyprinoidei | Cyprinidae | Poropuntius | huangchuchieni | NCBI | | 760.2 | diploid | non-miniature | 25.6 | 760.2 |
| LR15083 | Cyprinoidei | Cyprinidae | Poropuntius | sp | This study | NA | | diploid | non-miniature | 20 | 844.2840135 |
| LR15065 | Cyprinoidei | Cyprinidae | Probarbus | jullieni | This study | NA | | polyploid | non-miniature | 150 | 1350.403991 |

| | | | | | | | | | | | |
|---|---|---|---|---|---|---|---|---|---|---|---|
| 125LR15728 | Cyprinoidei | Cyprinidae | Puntigrus | partipentazona | This study | NA | | diploid | non-miniature | 2.95 | 597.0353715 |
| GCF_018831695.1 | Cyprinoidei | Cyprinidae | Puntigrus | tetrazona | NCBI | NA | | diploid | non-miniature | 4.78 | 579.1733795 |
| EA1300 | Cyprinoidei | Cyprinidae | Puntius | sp | This study | NA | | diploid | non-miniature | 14.7 | 797.7085135 |
| LR15257 | Cyprinoidei | Cyprinidae | Rohanella | titteya | This study | NA | | diploid | non-miniature | 3.66 | 752.468901 |
| 124LR15727 | Cyprinoidei | Cyprinidae | Rohtee | ogilbii | This study | NA | | diploid | non-miniature | 40 | 707.9410645 |
| 145LR15691 | Cyprinoidei | Cyprinidae | Sahyadria | denisonii | This study | NA | | diploid | non-miniature | 19 | 697.7408185 |
| LR15061 | Cyprinoidei | Cyprinidae | Sawbwa | resplendens | This study | NA | | diploid | proportioned-dwarf | 2.17 | 792.2622485 |
| LR15564 | Cyprinoidei | Cyprinidae | Scaphiodonichthys | acanthopterus | This study | NA | | diploid | non-miniature | 31 | 780.9871525 |
| LR15101 | Cyprinoidei | Cyprinidae | Schismatorhynchos | heterorhynchos | This study | NA | | diploid | non-miniature | 28.3 | 892.758514 |
| LR09392 | Cyprinoidei | Cyprinidae | Semiplotus | sp | This study | NA | | diploid | non-miniature | 18.5 | 790.364518 |
| GCF_001515605.1 | Cyprinoidei | Cyprinidae | Sinocyclocheilus | anshuiensis | NCBI | | 1632 | polyploid | non-miniature | 10 | 1632 |
| GCF_001515645.1 | Cyprinoidei | Cyprinidae | Sinocyclocheilus | grahami | NCBI | | 1750 | polyploid | non-miniature | 15 | 1750 |
| GCA_018148995.1 | Cyprinoidei | Cyprinidae | Sinocyclocheilus | maitianheensis | NCBI | | 1680 | polyploid | non-miniature | 9.4 | 1680 |
| GCF_001515625.1 | Cyprinoidei | Cyprinidae | Sinocyclocheilus | rhinocerous | NCBI | | 1655 | polyploid | non-miniature | 9.7 | 1655 |
| 111LR15711 | Cyprinoidei | Cyprinidae | Spinibarbichthys | denticulatus | This study | NA | | polyploid | non-miniature | 41.5 | 1725.003298 |
| LR15617 | Cyprinoidei | Cyprinidae | Spinibarbichthys | sp | This study | NA | | polyploid | non-miniature | 41.5 | 1769.692711 |
| 126LR15731 | Cyprinoidei | Cyprinidae | Striuntius | lineatus | This study | NA | | diploid | non-miniature | 5.95 | 652.6217305 |
| LR06590 | Cyprinoidei | Cyprinidae | Systomus | binduchitra | This study | NA | | diploid | non-miniature | 6 | 649.663832 |
| KUFOS.RR.2022.P.1 | Cyprinoidei | Cyprinidae | Tariqilabeo | periyarensis | This study | NA | | diploid | non-miniature | 13.1 | 1038.771302 |
| LR12880 | Cyprinoidei | Cyprinidae | Thynnichthys | sp | This study | NA | | diploid | non-miniature | 20 | 946.176263 |
| LR09401 | Cyprinoidei | Cyprinidae | Tor | sp1 | This study | NA | | polyploid | non-miniature | 150 | 1807.676521 |
| LR01638 | Cyprinoidei | Danionidae | Amblypharyngodon | chulabhornae | This study | NA | | diploid | non-miniature | 4 | 561.2414775 |
| fAmbSpp1 | Cyprinoidei | Danionidae | Amblypharyngodon | sp | NCBI-SRA | NA | | diploid | non-miniature | 5.5 | 582.919701 |
| LR15576 | Cyprinoidei | Danionidae | Barilius | mesopotamicus | This study | NA | | diploid | non-miniature | 5 | 632.689895 |
| LR15871 | Cyprinoidei | Danionidae | Boraras | brigittae | This study | NA | | diploid | proportioned-dwarf | 1.8 | 989.810021 |
| fBorMac1 | Cyprinoidei | Danionidae | Boraras | maculatus | NCBI-SRA | NA | | diploid | proportioned-dwarf | 2 | 923.9557445 |
| LR12206 | Cyprinoidei | Danionidae | Boraras | micros | This study | NA | | diploid | proportioned-dwarf | 1.3 | 953.8686275 |
| LR12190 | Cyprinoidei | Danionidae | Boraras | urophthalmoides | This study | NA | | diploid | proportioned-dwarf | 1.8 | 1043.537416 |
| LR09154 | Cyprinoidei | Danionidae | Brevibora | cheeya1 | This study | NA | | diploid | proportioned-dwarf | 2.8 | 1057.686109 |
| LR10130 | Cyprinoidei | Danionidae | Brevibora | cheeya2 | This study | NA | | diploid | proportioned-dwarf | 2.8 | 1029.485158 |
| LR11616 | Cyprinoidei | Danionidae | Brevibora | dorsiocellata | This study | NA | | diploid | proportioned-dwarf | 2.3 | 866.844275 |
| LR15567 | Cyprinoidei | Danionidae | Cabdio | morar | This study | NA | | diploid | non-miniature | 10 | 549.1380975 |
| fCheCac1 | Cyprinoidei | Danionidae | Chela | cachius | NCBI-SRA | NA | | diploid | non-miniature | 5 | 1008.955852 |
| 131LR08589 | Cyprinoidei | Danionidae | Chelaethiops | elongatus | This study | NA | | diploid | non-miniature | 4.8 | 579.350842 |
| LR15596 | Cyprinoidei | Danionidae | Chelaethiops | sp1 | This study | NA | | diploid | non-miniature | 5 | 608.825888 |
| LR15597 | Cyprinoidei | Danionidae | Chelaethiops | sp2 | This study | NA | | diploid | non-miniature | 5 | 593.233103 |
| GCF_903798145.1 | Cyprinoidei | Danionidae | Danio | aesculapii | NCBI | | 1381 | diploid | non-miniature | 3 | 1381 |
| GCA_903798035.1 | Cyprinoidei | Danionidae | Danio | albolineatus | NCBI | | 1463 | diploid | non-miniature | 3.4 | 1463 |
| GCA_903798125.1 | Cyprinoidei | Danionidae | Danio | choprae | NCBI | | 1102 | diploid | non-miniature | 3.1 | 1102 |
| LR01642 | Cyprinoidei | Danionidae | Danio | erythromicron | This study | NA | | diploid | proportioned-dwarf | 2.37 | 1006.792028 |
| GCA_903798115.1 | Cyprinoidei | Danionidae | Danio | jaintianensis | NCBI | | 1122 | diploid | non-miniature | 8 | 1122 |
| GCA_903798195.1 | Cyprinoidei | Danionidae | Danio | kyathit | NCBI | | 1708 | diploid | non-miniature | 3.5 | 1708 |
| LR02056 | Cyprinoidei | Danionidae | Danio | margaritatus | This study | NA | | diploid | proportioned-dwarf | 2.1 | 928.546121 |
| GCA_944039275.1 | Cyprinoidei | Danionidae | Danio | rerio | NCBI | | 1413 | diploid | non-miniature | 3.8 | 1413 |
| GCA_903798205.1 | Cyprinoidei | Danionidae | Danio | tinwini | NCBI | | 1497 | diploid | proportioned-dwarf | 2.6 | 1497 |
| GCA_903798025.1 | Cyprinoidei | Danionidae | Danionella | cerebrum | NCBI | | 657.5 | diploid | progenetic-miniature | 1.4 | 657.5 |
| GCA_900490495.1 | Cyprinoidei | Danionidae | Danionella | dracula | NCBI | | 665.2 | diploid | progenetic-miniature | 1.7 | 665.2 |
| fDevDev2 | Cyprinoidei | Danionidae | Devario | devario | NCBI-SRA | NA | | diploid | non-miniature | 7 | 1617.316438 |

| | | | | | | | | | | |
|---|---|---|---|---|---|---|---|---|---|---|
| 144LR15698 | Cyprinoidei | Danionidae | Devario | sondhii | This study | NA | diploid | non-miniature | 6.5 | 1064.408718 |
| fDanAeq2 | Cyprinoidei | Danionidae | Devario | sp | NCBI-SRA | NA | diploid | non-miniature | 10 | 1613.603733 |
| LR15598 | Cyprinoidei | Danionidae | Engraulicypris | sp1 | This study | NA | diploid | non-miniature | 7.5 | 574.0788735 |
| LR15656 | Cyprinoidei | Danionidae | Engraulicypris | sp2 | This study | NA | diploid | non-miniature | 7.5 | 554.3224095 |
| 116LR15716 | Cyprinoidei | Danionidae | Esomus | danricus | This study | NA | diploid | non-miniature | 6 | 521.562969 |
| LR01653 | Cyprinoidei | Danionidae | Esomus | metallicus | This study | NA | diploid | non-miniature | 7.5 | 513.0350255 |
| LR12503 | Cyprinoidei | Danionidae | Esomus | sp | This study | NA | diploid | non-miniature | 7.5 | 519.440578 |
| fHorAtu1 | Cyprinoidei | Danionidae | Horadandia | atukorali | NCBI-SRA | NA | diploid | proportioned-dwarf | 1.93 | 585.840953 |
| LR01646 | Cyprinoidei | Danionidae | Inlecypris | auropurpurea | This study | NA | diploid | non-miniature | 5 | 1316.965625 |
| LR01674 | Cyprinoidei | Danionidae | Kottelatia | brittani | This study | NA | diploid | non-miniature | 4.13 | 689.6199065 |
| fLauMye1 | Cyprinoidei | Danionidae | Laubuka | siamensis | NCBI-SRA | NA | diploid | non-miniature | 6.1 | 1336.049704 |
| 132LR09413 | Cyprinoidei | Danionidae | Laubuka | sp1 | This study | NA | diploid | non-miniature | 8 | 1300.971656 |
| KUFOS.AN.2021.3.1 | Cyprinoidei | Danionidae | Laubuka | sp2 | This study | NA | diploid | non-miniature | 8 | 1271.61059 |
| LR15601 | Cyprinoidei | Danionidae | Leptocypris | sp | This study | NA | diploid | non-miniature | 11.8 | 620.067572 |
| fLucSpp1 | Cyprinoidei | Danionidae | Luciosoma | sp1 | NCBI-SRA | NA | diploid | non-miniature | 26 | 599.554698 |
| LR01659 | Cyprinoidei | Danionidae | Luciosoma | sp2 | This study | NA | diploid | non-miniature | 26 | 618.508899 |
| LR15098 | Cyprinoidei | Danionidae | Malayochela | maassi | This study | NA | diploid | non-miniature | 4 | 828.8339765 |
| LR15568 | Cyprinoidei | Danionidae | Microdevario | kubotai | This study | NA | diploid | proportioned-dwarf | 1.9 | 1010.972865 |
| LR15650 | Cyprinoidei | Danionidae | Microrasbora | rubescens | This study | NA | diploid | proportioned-dwarf | 2.63 | 1019.876315 |
| 121LR15723 | Cyprinoidei | Danionidae | Nematabramis | everetti | This study | NA | diploid | non-miniature | 10.5 | 861.4281705 |
| LR15099 | Cyprinoidei | Danionidae | Nematabramis | steindachneri | This study | NA | diploid | non-miniature | 11.2 | 831.6380315 |
| LR14062 | Cyprinoidei | Danionidae | Neochela | dadiburjori | This study | NA | diploid | non-miniature | 3 | 761.266707 |
| LR15289 | Cyprinoidei | Danionidae | Opsaridium | sp1 | This study | NA | diploid | non-miniature | 12 | 672.00933 |
| LR15290 | Cyprinoidei | Danionidae | Opsaridium | sp2 | This study | NA | diploid | non-miniature | 12 | 644.8919075 |
| LR15602 | Cyprinoidei | Danionidae | Opsaridium | sp3 | This study | NA | diploid | non-miniature | 12 | 686.125769 |
| 142LR15666 | Cyprinoidei | Danionidae | Opsarius | sp1 | This study | NA | diploid | non-miniature | 9.1 | 596.7035535 |
| LR02136 | Cyprinoidei | Danionidae | Opsarius | sp2 | This study | NA | diploid | non-miniature | 9.1 | 565.8439115 |
| LR12240 | Cyprinoidei | Danionidae | Opsarius | sp3 | This study | NA | diploid | non-miniature | 11 | 588.440035 |
| LR15081 | Cyprinoidei | Danionidae | Pectenocypris | sp | This study | NA | diploid | non-miniature | 3.5 | 783.181521 |
| fRaiGut1 | Cyprinoidei | Danionidae | Raiamas | guttatus | NCBI-SRA | NA | diploid | non-miniature | 30 | 623.62552 |
| LR15292 | Cyprinoidei | Danionidae | Raiamas | moorii | This study | NA | diploid | non-miniature | 20 | 619.3998015 |
| 105LR15605 | Cyprinoidei | Danionidae | Raiamas | sp | This study | NA | diploid | non-miniature | 20 | 686.7324995 |
| LR10786 | Cyprinoidei | Danionidae | Rasbora | cephalotaenia | This study | NA | diploid | non-miniature | 7.73 | 766.9291225 |
| KUFOS.AN.2021.2.1 | Cyprinoidei | Danionidae | Rasbora | dandia | This study | NA | diploid | non-miniature | 9.9 | 540.6536305 |
| fRasDan1 | Cyprinoidei | Danionidae | Rasbora | daniconius | NCBI-SRA | NA | diploid | non-miniature | 11 | 675.4717815 |
| LR15347 | Cyprinoidei | Danionidae | Rasbora | kalbarensis | This study | NA | diploid | proportioned-dwarf | 2.5 | 814.203926 |
| LR15877 | Cyprinoidei | Danionidae | Rasbora | kalochroma | This study | NA | diploid | non-miniature | 6 | 1290.849861 |
| LR10804 | Cyprinoidei | Danionidae | Rasbora | kottelati | This study | NA | diploid | non-miniature | 7 | 1246.322742 |
| LR09739 | Cyprinoidei | Danionidae | Rasbora | myersi | This study | NA | diploid | non-miniature | 7.3 | 766.9803475 |
| fRasRas1 | Cyprinoidei | Danionidae | Rasbora | rasbora | NCBI-SRA | NA | diploid | non-miniature | 10 | 914.083791 |
| LR09847 | Cyprinoidei | Danionidae | Rasbora | trilineata | This study | NA | diploid | non-miniature | 6 | 885.3081555 |
| LR05463 | Cyprinoidei | Danionidae | Rasboroides | pallidus | This study | NA | diploid | non-miniature | 3.6 | 628.5166625 |
| LR12212 | Cyprinoidei | Danionidae | Rasbosoma | spilocerca | This study | NA | diploid | proportioned-dwarf | 2.6 | 901.6240925 |
| KUFOS.AN.2021.6.1 | Cyprinoidei | Danionidae | Salmostoma | balookee | This study | NA | diploid | non-miniature | 14.5 | 583.0889565 |
| EA0634 | Cyprinoidei | Danionidae | Thryssocypris | tonlesapensis | This study | NA | diploid | non-miniature | 6.4 | 447.905259 |
| LR08172 | Cyprinoidei | Danionidae | Trigonopoma | gracile | This study | NA | diploid | non-miniature | 5.5 | 872.3102705 |
| LR10181 | Cyprinoidei | Danionidae | Trigonopoma | pauciperforatum | This study | NA | diploid | non-miniature | 5 | 984.3918645 |
| LR11656 | Cyprinoidei | Danionidae | Trigonopoma | sp | This study | NA | diploid | non-miniature | 5 | 1076.334203 |

| | | | | | | | | | | | |
|---|---|---|---|---|---|---|---|---|---|---|---|
| LR08812 | Cyprinoidei | Danionidae | Trigonostigma | espei | This study | NA | | diploid | proportioned-dwarf | 2.5 | 1111.510812 |
| LR11086 | Cyprinoidei | Danionidae | Trigonostigma | heteromorpha | This study | NA | | diploid | proportioned-dwarf | 2.84 | 1115.305677 |
| GCA_949357685.1 | Cyprinoidei | Gobionidae | Gobio | gobio | NCBI | | 1460 | diploid | non-miniature | 13 | 1460 |
| GCA_023029165.1 | Cyprinoidei | Gobionidae | Gobiocypris | rarus | NCBI | | 1134 | diploid | non-miniature | 5.9 | 1134 |
| LR15569 | Cyprinoidei | Gobionidae | Hemibarbus | maculatus | This study | NA | | diploid | non-miniature | 34 | 1025.660767 |
| LR15570 | Cyprinoidei | Gobionidae | Ladislavia | taczanowskii | This study | NA | | diploid | non-miniature | 12 | 1113.559064 |
| LR15062 | Cyprinoidei | Gobionidae | Microphysogobio | tafangensis | This study | NA | | diploid | non-miniature | 7.8 | 947.9214225 |
| GCA_018749465.1 | Cyprinoidei | Gobionidae | Paracanthobrama | guichenoti | NCBI | | 1087 | diploid | non-miniature | 41.5 | 1087 |
| GCA_024679245.1 | Cyprinoidei | Gobionidae | Pseudorasbora | parva | NCBI | | 1228 | diploid | non-miniature | 9.5 | 1228 |
| GCA_023566225.1 | Cyprinoidei | Gobionidae | Romanogobio | albipinnatus | NCBI | | 1428 | diploid | non-miniature | 11.5 | 1428 |
| LR15571 | Cyprinoidei | Gobionidae | Sarcocheilichthys | hainanensis | This study | NA | | diploid | non-miniature | 7.5 | 1208.260729 |
| LR15579 | Cyprinoidei | Gobionidae | Squalidus | sp | This study | NA | | diploid | non-miniature | 18.8 | 994.7189 |
| LR15106 | Cyprinoidei | Leptobarbidae | Leptobarbus | sp | This study | NA | | diploid | non-miniature | 100 | 1001.959424 |
| GCA_963993115.1 | Cyprinoidei | Leuciscidae | Abramis | brama | NCBI | NA | | diploid | non-miniature | 70 | 1081.577457 |
| 2180_2012 | Cyprinoidei | Leuciscidae | Alburnoides | bipunctatus | This study | NA | | diploid | non-miniature | 13 | 1096.164941 |
| 108LR15742 | Cyprinoidei | Leuciscidae | Chrosomus | erythrogaster | This study | NA | | diploid | non-miniature | 6.1 | 1134.19284 |
| GCA_027497735.1 | Cyprinoidei | Leuciscidae | Cyprinella | lepida | NCBI | | 768.2 | diploid | non-miniature | 7.1 | 768.2 |
| GCA_027497675.1 | Cyprinoidei | Leuciscidae | Cyprinella | lutrensis | NCBI | NA | | diploid | non-miniature | 7.1 | 1010.965464 |
| GCA_027361055.1 | Cyprinoidei | Leuciscidae | Cyprinella | venusta | NCBI | | 805.4 | diploid | non-miniature | 15.5 | 805.4 |
| LR15252 | Cyprinoidei | Leuciscidae | Dionda | flavipinnis | This study | NA | | diploid | non-miniature | 7.6 | 817.7215975 |
| 110LR15746 | Cyprinoidei | Leuciscidae | Gila | pandora | This study | NA | | diploid | non-miniature | 15 | 1133.136695 |
| GCA_030378365.1 | Cyprinoidei | Leuciscidae | Hybognathus | amarus | NCBI | NA | | diploid | non-miniature | 9.3 | 937.2453165 |
| GCA_023718395.1 | Cyprinoidei | Leuciscidae | Leucaspius | delineatus | NCBI | | 1153 | diploid | non-miniature | 9 | 1153 |
| GCA_021554675.1 | Cyprinoidei | Leuciscidae | Leuciscus | idus | NCBI | | 1104 | diploid | non-miniature | 85 | 1104 |
| 2223_2013 | Cyprinoidei | Leuciscidae | Leuciscus | leuciscus | This study | NA | | diploid | non-miniature | 25 | 1159.306152 |
| GCA_030578275.1 | Cyprinoidei | Leuciscidae | Meda | fulgida | NCBI | | 882.1 | diploid | non-miniature | 6.5 | 882.1 |
| LR15254 | Cyprinoidei | Leuciscidae | Notropis | chalybaeus | This study | NA | | diploid | non-miniature | 5 | 1230.957706 |
| GCA_949152265.1 | Cyprinoidei | Leuciscidae | Phoxinus | phoxinus | NCBI | NA | | diploid | non-miniature | 10 | 980.731219 |
| GCF_016745375.1 | Cyprinoidei | Leuciscidae | Pimephales | promelas | NCBI | | 1066 | diploid | non-miniature | 8.2 | 1066 |
| 109LR15744 | Cyprinoidei | Leuciscidae | Ptychocheilus | lucius | This study | NA | | diploid | non-miniature | 18 | 1094.060096 |
| LR15251 | Cyprinoidei | Leuciscidae | Rhinichthys | cataractae | This study | NA | | diploid | non-miniature | 10.4 | 1101.099443 |
| LR15572 | Cyprinoidei | Leuciscidae | Rhynchocypris | lagowskii | This study | NA | | diploid | non-miniature | 18.5 | 1230.470575 |
| GCA_951802725.1 | Cyprinoidei | Leuciscidae | Rutilus | rutilus | NCBI | NA | | diploid | non-miniature | 50 | 1063.445747 |
| GCA_024453875.1 | Cyprinoidei | Leuciscidae | Scardinius | erythrophthalmus | NCBI | NA | | diploid | non-miniature | 35 | 1103.761005 |
| GCA_031834385.1 | Cyprinoidei | Leuciscidae | Semotilus | atromaculatus | NCBI | NA | | diploid | non-miniature | 24.5 | 1174.221529 |
| GCA_949319135.1 | Cyprinoidei | Leuciscidae | Squalius | cephalus | NCBI | NA | | diploid | non-miniature | 60 | 1084.922084 |
| 1450_2013 | Cyprinoidei | Leuciscidae | Telestes | muticellus | This study | NA | | diploid | non-miniature | 17 | 1160.188897 |
| GCA_030578255.1 | Cyprinoidei | Leuciscidae | Tiaroga | cobitis | NCBI | | 1320 | diploid | non-miniature | 6 | 1320 |
| GCA_022828995.1 | Cyprinoidei | Leuciscidae | Vimba | vimba | NCBI | | 1054 | diploid | non-miniature | 35 | 1054 |
| LR12004 | Cyprinoidei | Paedocyprididae | Paedocypris | carbunculus | NCBI-SRA | NA | | diploid | progenetic-miniature | 1.15 | 422.0606155 |
| GCA_028031935.1 | Cyprinoidei | Paedocyprididae | Paedocypris | micromegethes1 | NCBI | | 421.3 | diploid | progenetic-miniature | 1.3 | 421.3 |
| LR7898 | Cyprinoidei | Paedocyprididae | Paedocypris | micromegethes2 | NCBI-SRA | NA | | diploid | progenetic-miniature | 1.3 | 407.596544 |
| LR15325 | Cyprinoidei | Paedocyprididae | Paedocypris | sp2 | This study | NA | | diploid | progenetic-miniature | 1.3 | 517.668667 |
| LR15863A | Cyprinoidei | Paedocyprididae | Paedocypris | sp1 | This study | NA | | diploid | progenetic-miniature | 1.3 | 401.451712 |
| LR09467 | Cyprinoidei | Psilorhynchidae | Psilorhynchus | sp1 | This study | NA | | diploid | non-miniature | 5.1 | 482.2281655 |
| LR15089 | Cyprinoidei | Psilorhynchidae | Psilorhynchus | sp2 | This study | NA | | diploid | non-miniature | 4.9 | 389.6027315 |
| 100LR08982 | Cyprinoidei | Sundadanionidae | Fangfangia | spinicleithralis | This study | NA | | diploid | progenetic-miniature | 2.3 | 761.3466415 |
| LR15870 | Cyprinoidei | Sundadanionidae | Sundadanio | atomus | This study | NA | | diploid | progenetic-miniature | 1.6 | 654.471117 |

| | | | | | | | | | | | |
|---|---|---|---|---|---|---|---|---|---|---|---|
| LR10372 | Cyprinoidei | Sundadanionidae | Sundadanio | goblinus | This study | NA | | diploid | progenetic-miniature | 1.9 | 679.917281 |
| LR15082 | Cyprinoidei | Sundadanionidae | Sundadanio | margarition | This study | NA | | diploid | progenetic-miniature | 2 | 716.7551635 |
| 101LR12751 | Cyprinoidei | Sundadanionidae | Sundadanio | retiarius | This study | NA | | diploid | progenetic-miniature | 1.9 | 752.2425955 |
| LR15110 | Cyprinoidei | Tanichthyidae | Tanichthys | albonudes | This study | NA | | diploid | proportioned-dwarf | 2.77 | 1071.863989 |
| LR02462 | Cyprinoidei | Tanichthyidae | Tanichthys | micagemmae | This study | NA | | diploid | proportioned-dwarf | 2.3 | 1093.443826 |
| 1545_2010 | Cyprinoidei | Tincidae | Tinca | tinca | This study | NA | | diploid | non-miniature | 70 | 1048.69225 |
| GCA_003731715.1 | Cyprinoidei | Xenocyprididae | Anabarilius | grahami | NCBI | | 991.9 | diploid | non-miniature | 15.6 | 991.9 |
| 118LR15719 | Cyprinoidei | Xenocyprididae | Candidia | pingtungensis | This study | NA | | diploid | non-miniature | 11.4 | 780.192702 |
| GCA_024489055.1 | Cyprinoidei | Xenocyprididae | Chanodichthys | erythropterus | NCBI | | 1085 | diploid | non-miniature | 80 | 1085 |
| GCF_019924925.1 | Cyprinoidei | Xenocyprididae | Ctenopharyngodon | idella | NCBI | | 893.2 | diploid | non-miniature | 80 | 893.2 |
| GCA_028476615.1 | Cyprinoidei | Xenocyprididae | Culter | alburnus | NCBI | | 1055 | diploid | non-miniature | 60 | 1055 |
| GCA_037101425.1 | Cyprinoidei | Xenocyprididae | Elopichthys | bambusa | NCBI | | 827.7 | diploid | non-miniature | 180 | 827.7 |
| 103LR15622 | Cyprinoidei | Xenocyprididae | Hemiculter | sp | This study | NA | | diploid | non-miniature | 20.5 | 977.935087 |
| GCA_004764525.1 | Cyprinoidei | Xenocyprididae | Hypophthalmichthys | molitrix | NCBI | | 1104 | diploid | non-miniature | 100 | 1104 |
| GCA_019925145.1 | Cyprinoidei | Xenocyprididae | Hypophthalmichthys | nobilis | NCBI | | 857 | diploid | non-miniature | 146 | 857 |
| GCF_018812025.1 | Cyprinoidei | Xenocyprididae | Megalobrama | amblycephala | NCBI | | 1109 | diploid | non-miniature | 180 | 1109 |
| 102LR15621 | Cyprinoidei | Xenocyprididae | Opsariichthys | bidens | This study | NA | | diploid | non-miniature | 16 | 905.3265225 |
| 128LR15724 | Cyprinoidei | Xenocyprididae | Opsariichthys | pachycephalus | This study | NA | | diploid | non-miniature | 12.2 | 734.149207 |
| LR15574 | Cyprinoidei | Xenocyprididae | Oxygaster | pointoni | This study | NA | | diploid | non-miniature | 15.5 | 862.965098 |
| GCA_034642465.1 | Cyprinoidei | Xenocyprididae | Zacco | platypus | NCBI | | 815 | diploid | non-miniature | 13 | 815 |
| LR15097 | Gyrinocheiloidei | Gyrinocheilidae | Gyrinocheilus | pustulosus | This study | NA | | diploid | non-miniature | 28 | 478.3126755 |
| 104EA0138 | Gyrinocheiloidei | Gyrinocheilidae | Gyrinocheilus | sp | This study | NA | | diploid | non-miniature | 28 | 475.3712995 |

**Supplementary Table 6.** Miniature fishes within Cypriniformes, comprising progenetic miniatures and proportioned dwarfs, with indication of which species were included in the present study.

| Suborder | Family | Genus | Miniature species | Miniaturization type | Included_309Cyp_dataset | Distribution | Reference | Remarks |
|---|---|---|---|---|---|---|---|---|
| Cobitoidei | Balitoridae | Homalopteroides | *Homalopteroides tweediei* | Proportional dwarf | yes | Asian freshwaters | Kottelat & Vidthayanon (1993) | Genus comprises both miniature (n=1) and non-miniature species (n=12) |
| Cobitoidei | Balitoridae | Neohomaloptera | *Neohomaloptera johorensis* | Proportional dwarf | yes | Asian freshwaters | Kottelat & Vidthayanon (1993) | Monotypic genus comprising a miniature species (n=1) |
| Cobitoidei | Barbuccidae | Barbucca | *Barbucca elongatus* | Proportional dwarf | yes | Asian freshwaters | Vasil'eva & Vasil'ev (2013) | Genus comprises only miniature species (n=3) |
| Cobitoidei | Barbuccidae | Barbucca | *Barbucca diabolica* | Proportional dwarf | no | Asian freshwaters | Kottelat & Vidthayanon (1993) | |
| Cobitoidei | Barbuccidae | Barbucca | *Barbucca heokhuii* | Proportional dwarf | no | Asian freshwaters | Kottelat (2025) | |
| Cobitoidei | Cobitidae | Kottelatlimia | *Kottelatlimia katik* | Proportional dwarf | yes | Asian freshwaters | Kottelat & Vidthayanon (1993) | Genus comprises both miniature (n=1) and non-miniature species (n=2) |
| Cobitoidei | Cobitidae | Pangio | *Pangio bhujia* | Proportional dwarf | yes | Asian freshwaters | Anoop et al. (2019) | Genus comprises both miniature (n=4) and non-miniature species (n=31) |
| Cobitoidei | Cobitidae | Pangio | *Pangio ammophila* | Proportional dwarf | no | Asian freshwaters | Britz et al. (2012) | |
| Cobitoidei | Cobitidae | Pangio | *Pangio longimanus* | Proportional dwarf | no | Asian freshwaters | Britz & Kottelat (2010) | |
| Cobitoidei | Cobitidae | Pangio | *Pangio pathala* | Proportional dwarf | no | Asian freshwaters | Sundar et al. (2022) | |
| Cobitoidei | Cobitidae | Gitchak | *Gitchak nakana* | Proportional dwarf | no | Asian freshwaters | Britz et al. (2026) | Monotypic genus comprising a miniature species (n=1) |
| Cobitoidei | Nemacheilidae | Tuberoschistura | *Tuberoschistura cambodgiensis* | Proportional dwarf | no | Asian freshwaters | Kottelat & Vidthayanon (1993) | Genus comprises both miniature (n=1) and non-miniature species (n=1) |
| Cobitoidei | Nemacheilidae | Physoschistura | *Physoschistura mango* | Proportional dwarf | no | Asian freshwaters | Conway & Kottelat (2023) | Genus comprises both miniature (n=1) and non-miniature species (n=8) |
| Cobitoidei | Nemacheilidae | Schistura | *Schistura diminuta* | Proportional dwarf | no | Asian freshwaters | Ou et al. (2011) | Genus comprises majority of non-miniature species (n=244) |
| Cobitoidei | Nemacheilidae | Schistura | *Schistura rikiki* | Proportional dwarf | no | Asian freshwaters | Kottelat (2000) | |
| Cyprinoidei | Cyprinidae | Sawbwa | *Sawbwa resplendens* | Proportional dwarf | yes | Asian freshwaters | Kottelat & Vidthayanon (1993) | Monotypic genus comprising a single species (n=1); considered here a miniature based on Kottelat & Vidthayanon (1993) but may grow bigger |
| Cyprinoidei | Cyprinidae | Garra | *Garra lancrenonensis* | Proportional dwarf | no | African freshwaters | Conway & Moritz (2006) and ref herein | Genus comprises majority of non-miniature species (n=192) |
| Cyprinoidei | Cyprinidae | Enteromius | *Enteromius candens* | Proportional dwarf | no | African freshwaters | Conway & Moritz (2006) and ref herein | Genus comprises majority of non-miniature species (n=223); the number of miniature species reported maybe underrepresented ? |
| Cyprinoidei | Cyprinidae | Enteromius | *Enteromius erythrozonus* | Proportional dwarf | no | African freshwaters | Conway & Moritz (2006) and ref herein | |
| Cyprinoidei | Cyprinidae | Enteromius | *Enteromius kissiensis* | Proportional dwarf | no | African freshwaters | Conway & Moritz (2006) and ref herein | |
| Cyprinoidei | Cyprinidae | Enteromius | *Enteromius nigrifilis* | Proportional dwarf | no | African freshwaters | Conway & Moritz (2006) and ref herein | |
| Cyprinoidei | Cyprinidae | Enteromius | *Enteromius nigroluteus* | Proportional dwarf | no | African freshwaters | Conway & Moritz (2006) and ref herein | |
| Cyprinoidei | Cyprinidae | Enteromius | *Enteromius papilio* | Proportional dwarf | no | African freshwaters | Conway & Moritz (2006) and ref herein | |
| Cyprinoidei | Cyprinidae | Enteromius | *Enteromius pumilus* | Proportional dwarf | no | African freshwaters | Conway & Moritz (2006) and ref herein | |
| Cyprinoidei | Cyprinidae | Enteromius | *Enteromius sylvaticus* | Proportional dwarf | no | African freshwaters | Conway & Moritz (2006) and ref herein | |
| Cyprinoidei | Cyprinidae | Enteromius | *Enteromius tongaensis* | Proportional dwarf | no | African freshwaters | Conway & Moritz (2006) and ref herein | |
| Cyprinoidei | Cyprinidae | Barboides | *Barboides gracilis* | Proportional dwarf | yes | African freshwaters | Conway & Moritz (2006) | Genus comprises only miniature species (n=2); here considered as a proportioned dwarf but the two species of Barboides display some progenetic features (Conway et al. 2017) |
| Cyprinoidei | Cyprinidae | Barboides | *Barboides britzi* | Proportional dwarf | no | African freshwaters | Conway & Moritz (2006) | |
| Cyprinoidei | Danionidae | Danio | *Danio erythromicron* | Proportional dwarf | yes | Asian freshwaters | Kottelat & Vidthayanon (1993) | Genus comprises both miniature (n=3) and non-miniature species (n=24) |
| Cyprinoidei | Danionidae | Danio | *Danio margaritatus* | Proportional dwarf | yes | Asian freshwaters | Roberts (2007) | |
| Cyprinoidei | Danionidae | Danio | *Danio tinwini* | Proportional dwarf | yes | Asian freshwaters | Kullander & Fang (2009) | |
| Cyprinoidei | Danionidae | Danionella | *Danionella cerebrum* | Progenetic miniature | yes | Asian freshwaters | Britz et al. (2021) | Genus comprises only miniature species (n=5) |
| Cyprinoidei | Danionidae | Danionella | *Danionella dracula* | Progenetic miniature | yes | Asian freshwaters | Britz et al. (2021) | |
| Cyprinoidei | Danionidae | Danionella | *Danionella mirifica* | Progenetic miniature | no | Asian freshwaters | Britz et al. (2021) | |
| Cyprinoidei | Danionidae | Danionella | *Danionella priapus* | Progenetic miniature | no | Asian freshwaters | Britz et al. (2021) | |
| Cyprinoidei | Danionidae | Danionella | *Danionella translucida* | Progenetic miniature | no | Asian freshwaters | Britz et al. (2021) | |
| Cyprinoidei | Danionidae | Microdevario | *Microdevario kubotai* | Proportional dwarf | yes | Asian freshwaters | Kottelat & Witte (1999) | Genus comprises only miniature species (n=4) |
| Cyprinoidei | Danionidae | Microdevario | *Microdevario gatesi* | Proportional dwarf | no | Asian freshwaters | Kottelat & Vidthayanon (1993) | |
| Cyprinoidei | Danionidae | Microdevario | *Microdevario microphthalma* | Proportional dwarf | no | Asian freshwaters | Jiang et al. (2008) | |
| Cyprinoidei | Danionidae | Microdevario | *Microdevario nanus* | Proportional dwarf | no | Asian freshwaters | Kottelat & Witte (1999) | |
| Cyprinoidei | Danionidae | Microrasbora | *Microrasbora rubescens* | Proportional dwarf | yes | Asian freshwaters | Kottelat & Vidthayanon (1993) | Monotypic genus comprising a miniature species (n=1); considered here a miniature based on Kottelat & Vidthayanon (1993) but may grow bigger |
| Cyprinoidei | Danionidae | Neobola | *Neobola stellae* | Proportional dwarf | no | African freshwaters | Daget et al. (1984) | Genus comprises both miniature (n=1) and non-miniature species (n=4) |
| Cyprinoidei | Danionidae | Boraras | *Boraras brigittae* | Proportional dwarf | yes | Asian freshwaters | Kottelat & Vidthayanon (1993) | Genus comprises only miniature species (n=6) |
| Cyprinoidei | Danionidae | Boraras | *Boraras maculatus* | Proportional dwarf | yes | Asian freshwaters | Kottelat & Vidthayanon (1993) | |
| Cyprinoidei | Danionidae | Boraras | *Boraras merah* | Proportional dwarf | no | Asian freshwaters | Kottelat & Vidthayanon (1993) | |
| Cyprinoidei | Danionidae | Boraras | *Boraras micros* | Proportional dwarf | yes | Asian freshwaters | Kottelat & Vidthayanon (1993) | |
| Cyprinoidei | Danionidae | Boraras | *Boraras naevus* | Proportional dwarf | no | Asian freshwaters | Conway & Kottelat (2011) | |
| Cyprinoidei | Danionidae | Boraras | *Boraras urophthalmoides* | Proportional dwarf | yes | Asian freshwaters | Kottelat & Vidthayanon (1993) | |
| Cyprinoidei | Danionidae | Brevibora | *Brevibora cheeya* | Proportional dwarf | yes | Asian freshwaters | Liao & Tan (2011) | Genus comprises only miniature species (n=3) |
| Cyprinoidei | Danionidae | Brevibora | *Brevibora dorsiocellata* | Proportional dwarf | yes | Asian freshwaters | Liao & Tan (2011) | |
| Cyprinoidei | Danionidae | Brevibora | *Brevibora exilis* | Proportional dwarf | no | Asian freshwaters | Liao & Tan (2014) | |
| Cyprinoidei | Danionidae | Horadandia | *Horadandia atukorali* | Proportional dwarf | yes | Asian freshwaters | Kottelat & Vidthayanon (1993) | Genus comprises only miniature species (n=2) |
| Cyprinoidei | Danionidae | Horadandia | *Horadandia brittani* | Proportional dwarf | no | Asian freshwaters | Batuwita et al. (2013) | |
| Cyprinoidei | Danionidae | Rasbora | *Rasbora kalbarensis* | Proportional dwarf | yes | Asian freshwaters | Kottelat & Vidthayanon (1993) | Genus comprises majority of non-miniature species (n=89) |
| Cyprinoidei | Danionidae | Rasbosoma | *Rasbosoma spilocerca* | Proportional dwarf | yes | Asian freshwaters | Kottelat & Vidthayanon (1993) | Monotypic genus comprising a miniature species (n=1) |
| Cyprinoidei | Danionidae | Trigonostigma | *Trigonostigma espei* | Proportional dwarf | yes | Asian freshwaters | Kottelat & Witte (1999) | Genus comprises only miniature species (n=5) |
| Cyprinoidei | Danionidae | Trigonostigma | *Trigonostigma hengeli* | Proportional dwarf | no | Asian freshwaters | Kottelat & Witte (1999) | |
| Cyprinoidei | Danionidae | Trigonostigma | *Trigonostigma heteromorpha* | Proportional dwarf | yes | Asian freshwaters | Kottelat & Witte (1999) | |
| Cyprinoidei | Danionidae | Trigonostigma | *Trigonostigma somphongsi* | Proportional dwarf | no | Asian freshwaters | Kottelat & Witte (1999) | |
| Cyprinoidei | Danionidae | Trigonostigma | *Trigonostigma truncata* | Proportional dwarf | no | Asian freshwaters | Tan (2020) | |
| Cyprinoidei | Paedocyprididae | Paedocypris | *Paedocypris carbunculus* | Progenetic miniature | yes | Asian freshwaters | Britz & Kottelat (2008) | Genus comprises only miniature species (n=3) |
| Cyprinoidei | Paedocyprididae | Paedocypris | *Paedocypris micromegethes* | Progenetic miniature | yes | Asian freshwaters | Kottelat et al. (2006) | |
| Cyprinoidei | Paedocyprididae | Paedocypris | *Paedocypris progenetica* | Progenetic miniature | no | Asian freshwaters | Kottelat et al. (2006) | |
| Cyprinoidei | Sundadanionidae | Fangfangia | *Fangfangia spinicleithralis* | Progenetic miniature | yes | Asian freshwaters | Britz et al. (2011) | Monotypic genus comprising a miniature species (n=1) |
| Cyprinoidei | Sundadanionidae | Sundadanio | *Sundadanio atomus* | Progenetic miniature | yes | Asian freshwaters | Conway et al. (2011) | Genus comprises only miniature species (n=8) |
| Cyprinoidei | Sundadanionidae | Sundadanio | *Sundadanio axelrodi* | Progenetic miniature | no | Asian freshwaters | Conway et al. (2011) | |
| Cyprinoidei | Sundadanionidae | Sundadanio | *Sundadanio echinus* | Progenetic miniature | no | Asian freshwaters | Conway et al. (2011) | |
| Cyprinoidei | Sundadanionidae | Sundadanio | *Sundadanio gargula* | Progenetic miniature | no | Asian freshwaters | Conway et al. (2011) | |
| Cyprinoidei | Sundadanionidae | Sundadanio | *Sundadanio goblinus* | Progenetic miniature | yes | Asian freshwaters | Conway et al. (2011) | |
| Cyprinoidei | Sundadanionidae | Sundadanio | *Sundadanio margarition* | Progenetic miniature | yes | Asian freshwaters | Conway et al. (2011) | |
| Cyprinoidei | Sundadanionidae | Sundadanio | *Sundadanio retiarius* | Progenetic miniature | yes | Asian freshwaters | Conway et al. (2011) | |
| Cyprinoidei | Sundadanionidae | Sundadanio | *Sundadanio rubellus* | Progenetic miniature | no | Asian freshwaters | Conway et al. (2011) | |
| Cyprinoidei | Tanichthyidae | Tanichthys | *Tanichthys albonubes* | Proportional dwarf | yes | Asian freshwaters | Freyhof & Herder (2001) | Genus comprises only miniature species (n=9) |
| Cyprinoidei | Tanichthyidae | Tanichthys | *Tanichthys micagemmae* | Proportional dwarf | yes | Asian freshwaters | Freyhof & Herder (2001) | |
| Cyprinoidei | Tanichthyidae | Tanichthys | *Tanichthys albiventris* | Proportional dwarf | no | Asian freshwaters | Li et al. (2022) | |
| Cyprinoidei | Tanichthyidae | Tanichthys | *Tanichthys flavianalis* | Proportional dwarf | no | Asian freshwaters | Li et al. (2022) | |

| | | | | | | | |
|---|---|---|---|---|---|---|---|
| Cyprinoidei | Tanichthyidae | Tanichthys | *Tanichthys guipingensis* | Proportional dwarf | no | Asian freshwaters | Jin et al. (2022) |
| Cyprinoidei | Tanichthyidae | Tanichthys | *Tanichthys huidongensis* | Proportional dwarf | no | Asian freshwaters | Jin et al. (2022) |
| Cyprinoidei | Tanichthyidae | Tanichthys | *Tanichthys kuehnei* | Proportional dwarf | no | Asian freshwaters | Bohlen et al. (2019) |
| Cyprinoidei | Tanichthyidae | Tanichthys | *Tanichthys luheensis* | Proportional dwarf | no | Asian freshwaters | Jin et al. (2022) |
| Cyprinoidei | Tanichthyidae | Tanichthys | *Tanichthys shenzhenensis* | Proportional dwarf | no | Asian freshwaters | Jin et al. (2022) |

**Supplementary Table 7.** Comparison of evolutionary model fit for log-transformed genome size and body size across Cypriniformes. Models were fitted using the fitContinuous function in the R package geiger.

| Trait | Model | sigsq | alpha | a | log-likelihood | AIC | AICc | deltaAICc | weight |
|---|---|---|---|---|---|---|---|---|---|
| log10GenomeSize | **BM** | **0.000324** | | | **344.596126** | **-685.192253** | **-685.153** | **0.619** | **0.3672** |
| | OU | 0.000324 | 0 | | 344.596126 | -683.192253 | -683.114 | 2.659 | 0.1324 |
| | **EB** | **0.00091** | | **-0.010328** | **345.925513** | **-685.851025** | **-685.772** | **0** | **0.5004** |
| | WN | 0.02905 | | | 108.282147 | -212.564293 | -212.525 | 473.247 | 0 |
| log10BodySize | BM | 0.007054 | | | -131.503969 | 267.007938 | 267.047 | 35.717 | 0 |
| | **OU** | **0.010609** | **0.026546** | | **-112.625856** | **231.251713** | **231.33** | **0** | **1** |
| | EB | 0.007055 | | -0.000001 | -131.50473 | 269.009461 | 269.088 | 37.758 | 0 |
| | WN | 0.206863 | | | -195.006364 | 394.012727 | 394.052 | 162.722 | 0 |

**Supplementary Table 8.** Results of phylogenetic generalized least squares (PGLS) regression analyses for log-transformed genome size and body size across Cypriniformes under a Brownian motion (BM) model of trait evolution.

| Model | Predictor | β (estimate) | SE (Standard error) | t value | p-value | $R^2$ | Adjusted $R^2$ | N |
|---|---|---|---|---|---|---|---|---|
| log10GenomeSize ~ miniature vs non-miniature | Intercept | 2.8456642 | 0.0822538 | 34.5961 | <2e-16 | 0.0005666 | -0.002689 | 309 |
| | miniature | -0.0078134 | 0.0187284 | -0.4172 | 0.6768 | | | |
| log10GenomeSize ~ non-miniature vs proportioned-dwarf vs progenetic-miniature | Intercept | 2.863487 | 0.082979 | 34.5085 | <2e-16 | 0.007635 | 0.001149 | 309 |
| | progenetic-miniature | -0.125775 | 0.08206 | -1.5327 | 0.1264 | | | |
| | proportioned-dwarf | -0.002135 | 0.019084 | -0.1119 | 0.911 | | | |
| log10GenomeSize ~ Ploidy | Intercept | 2.790532 | 0.073569 | 37.9309 | 2.20E-16 | 0.2043 | 0.2017 | 309 |
| | polyploid | 0.259152 | 0.029188 | 8.8787 | 2.20E-16 | | | |
| log10GenomeSize ~ miniature vs non-miniature + Ploidy | Intercept | 2.7915258 | 0.0737551 | 37.8486 | 2.00E-16 | 0.2045 | 0.1993 | 309 |
| | miniature | -0.0049448 | 0.0167387 | -0.2954 | 0.7679 | | | |
| | polyploid | 0.2589849 | 0.0292369 | 8.8582 | 2.00E-16 | | | |
| log10GenomeSize ~ non-miniature vs proportioned-dwarf vs progenetic-miniature + Ploidy | Intercept | 2.80609131 | 0.07448703 | 37.6722 | 2.00E-16 | 0.2091 | 0.2013 | 309 |
| | progenetic-miniature | -0.09937014 | 0.07344123 | -1.3531 | 0.177 | | | |
| | proportioned-dwarf | -0.00041616 | 0.01706662 | -0.0244 | 0.9806 | | | |
| | polyploid | 0.25754284 | 0.02922184 | 8.8134 | 2.00E-16 | | | |
| log10GenomeSize ~ log10BodySize | Intercept | 2.823171 | 0.082791 | 34.1001 | 2.00E-16 | 0.008961 | 0.005733 | 309 |
| | log10BodySize | 0.020279 | 0.012171 | 1.6662 | 0.0967 | | | |
| log10GenomeSize ~ log10BodySize + miniature vs non-miniature | Intercept | 2.821967 | 0.083338 | 33.8617 | 2.00E-16 | 0.00903 | 0.002553 | 309 |
| | log10BodySize | 0.0209039 | 0.0129311 | 1.6166 | 0.107 | | | |
| | miniature | 0.0028718 | 0.0198144 | 0.1449 | 0.8849 | | | |
| log10GenomeSize ~ log10BodySize + non-miniature vs proportioned-dwarf vs progenetic-miniature | Intercept | 2.8398846 | 0.0841886 | 33.7324 | 2.00E-16 | 0.01534 | 0.005659 | 309 |
| | log10BodySize | 0.0199781 | 0.0129279 | 1.5453 | 0.1233 | | | |
| | progenetic-miniature | -0.1092464 | 0.0825704 | -1.3231 | 0.1868 | | | |
| | proportioned-dwarf | 0.0077728 | 0.0200915 | 0.3869 | 0.6991 | | | |
| log10GenomeSize ~ log10BodySize + miniature vs non-miniature + Ploidy | Intercept | 2.78684 | 0.0748799 | 37.2174 | 2.20E-16 | 0.2049 | 0.1971 | 309 |
| | log10BodySize | 0.0044689 | 0.0117556 | 0.3802 | 0.7041 | | | |
| | miniature | -0.0026806 | 0.0177889 | -0.1507 | 0.8803 | | | |
| | polyploid | 0.2571661 | 0.0296662 | 8.6687 | 2.63E-16 | | | |
| log10GenomeSize ~ log10BodySize + non-miniature vs proportioned-dwarf vs progenetic-miniature + Ploidy | Intercept | 2.8019784 | 0.0756924 | 37.0179 | 2.20E-16 | 0.2093 | 0.1989 | 309 |
| | log10BodySize | 0.0037676 | 0.0117545 | 0.3205 | 0.7488 | | | |
| | progenetic-miniature | -0.0964086 | 0.0741276 | -1.3006 | 0.1944 | | | |
| | proportioned-dwarf | 0.0014422 | 0.0180484 | 0.0799 | 0.9364 | | | |
| | polyploid | 0.2560255 | 0.0296453 | 8.6363 | 3.34E-16 | | | |
| log10GenomeSize ~ log10BodySize * miniature vs non-miniature | Intercept | 2.8232972 | 0.0836524 | 33.7503 | 2.00E-16 | 0.009215 | -0.0005307 | 309 |
| | log10BodySize | 0.0206078 | 0.0130103 | 1.584 | 0.1142 | | | |
| | miniature | -0.0096016 | 0.0558882 | -0.1718 | 0.8637 | | | |
| | log10BodySize:miniature | 0.0348413 | 0.1459368 | 0.2387 | 0.8115 | | | |
| log10GenomeSize ~ log10BodySize * non-miniature vs proportioned-dwarf vs progenetic-miniature | Intercept | 2.842059 | 0.08445 | 33.6536 | 2.00E-16 | 0.01775 | 0.00154 | 309 |
| | log10BodySize | 0.020101 | 0.013003 | 1.5459 | 0.1232 | | | |
| | progenetic-miniature | -0.173465 | 0.119207 | -1.4552 | 0.1467 | | | |
| | proportioned-dwarf | 0.031958 | 0.0623 | 0.513 | 0.6083 | | | |
| | log10BodySize:progenetic-miniature | 0.293969 | 0.389548 | 0.7546 | 0.451 | | | |
| | log10BodySize:proportioned-dwarf | -0.067048 | 0.162355 | -0.413 | 0.6799 | | | |
| log10BodySize ~ miniature vs non-miniature | Intercept | 1.133629 | 0.362087 | 3.1308 | 0.001911 | 0.1113 | 0.1084 | 309 |
| | miniature | -0.511156 | 0.082444 | -6.2 | 1.82E-09 | | | |
| log10BodySize ~ non-miniature vs proportioned-dwarf vs progenetic-miniature | Intercept | 1.181398 | 0.366097 | 3.227 | 0.001387 | 0.1136 | 0.1078 | 309 |
| | progenetic-miniature | -0.827324 | 0.362043 | -2.2852 | 0.022988 | | | |
| | proportioned-dwarf | -0.495936 | 0.084198 | -5.8901 | 1.02E-08 | | | |

**Supplementary Table 9.** Results of phylogenetic generalized least squares (PGLS) regression analyses for log-transformed genome size and body size across Cypriniformes under an Ornstein–Uhlenbeck (OU) model of trait evolution.

| Model | Predictor | β (estimate) | SE (Standard error) | t value | p-value | $R^2$ | Adjusted $R^2$ | N |
|---|---|---|---|---|---|---|---|---|
| log10GenomeSize ~ miniature vs non-miniature | Intercept | 2.8464091 | 0.4107606 | 6.9296 | 2.49E-11 | 0.0006119 | -0.002643 | 309 |
| | miniature | -0.0081419 | 0.0187795 | -0.4336 | 0.6649 | | | |
| log10GenomeSize ~ non-miniature vs proportioned-dwarf vs progenetic-miniature | Intercept | 2.8641829 | 0.3877374 | 7.3869 | 1.44E-12 | 0.008068 | 0.001585 | 309 |
| | progenetic-miniature | -0.1259239 | 0.079893 | -1.5762 | 0.116 | | | |
| | proportioned-dwarf | -0.0021376 | 0.0191639 | -0.1115 | 0.9113 | | | |
| log10GenomeSize ~ Ploidy | Intercept | 2.792146 | 0.251733 | 11.0917 | 2.20E-16 | 0.2073 | 0.2047 | 309 |
| | polyploid | 0.259632 | 0.028978 | 8.9595 | 2.20E-16 | | | |
| log10GenomeSize ~ miniature vs non-miniature + Ploidy | Intercept | 2.7932193 | 0.2510412 | 11.1265 | 2.00E-16 | 0.2075 | 0.2024 | 309 |
| | miniature | -0.0052409 | 0.0167967 | -0.312 | 0.7552 | | | |
| | polyploid | 0.2594415 | 0.0290262 | 8.9382 | 2.00E-16 | | | |
| log10GenomeSize ~ non-miniature vs proportioned-dwarf vs progenetic-miniature + Ploidy | Intercept | 2.80767608 | 0.23605797 | 11.894 | 2.00E-16 | 0.2128 | 0.205 | 309 |
| | progenetic-miniature | -0.09860622 | 0.06972523 | -1.4142 | 0.1583 | | | |
| | proportioned-dwarf | -0.00023076 | 0.01717182 | -0.0134 | 0.9893 | | | |
| | polyploid | 0.25788167 | 0.02898577 | 8.8968 | 2.00E-16 | | | |
| log10GenomeSize ~ log10BodySize | Intercept | 2.823184 | 0.374153 | 7.5455 | 5.15E-13 | 0.009636 | 0.00641 | 309 |
| | log10BodySize | 0.021167 | 0.012247 | 1.7283 | 0.08495 | | | |
| log10GenomeSize ~ log10BodySize + miniature vs non-miniature | Intercept | 2.8219254 | 0.3745409 | 7.5344 | 5.58E-13 | 0.00971 | 0.003237 | 309 |
| | log10BodySize | 0.0218258 | 0.0130231 | 1.6759 | 0.09477 | | | |
| | miniature | 0.0029916 | 0.0198879 | 0.1504 | 0.88053 | | | |
| log10GenomeSize ~ log10BodySize + non-miniature vs proportioned-dwarf vs progenetic-miniature | Intercept | 2.839728 | 0.354962 | 8.0001 | 2.61E-14 | 0.01645 | 0.006777 | 309 |
| | log10BodySize | 0.020916 | 0.013028 | 1.6055 | 0.1094 | | | |
| | progenetic-miniature | -0.10861 | 0.080079 | -1.3563 | 0.176 | | | |
| | proportioned-dwarf | 0.008247 | 0.020192 | 0.4084 | 0.6832 | | | |
| log10GenomeSize ~ log10BodySize + miniature vs non-miniature + Ploidy | Intercept | 2.7877549 | 0.2455278 | 11.3541 | 2.00E-16 | 0.2082 | 0.2004 | 309 |
| | log10BodySize | 0.0053284 | 0.0118881 | 0.4482 | 0.6543 | | | |
| | miniature | -0.002542 | 0.0178756 | -0.1422 | 0.887 | | | |
| | polyploid | 0.2572796 | 0.0294603 | 8.7331 | 2.00E-16 | | | |
| log10GenomeSize ~ log10BodySize + non-miniature vs proportioned-dwarf vs progenetic-miniature + Ploidy | Intercept | 2.8027354 | 0.2319475 | 12.0835 | 2.00E-16 | 0.2133 | 0.2029 | 309 |
| | log10BodySize | 0.0046224 | 0.0118986 | 0.3885 | 0.6979 | | | |
| | progenetic-miniature | -0.0949436 | 0.0703324 | -1.3499 | 0.178 | | | |
| | proportioned-dwarf | 0.0020589 | 0.0181759 | 0.1133 | 0.9099 | | | |
| | polyploid | 0.2560225 | 0.0294153 | 8.7037 | 2.00E-16 | | | |
| log10GenomeSize ~ log10BodySize * miniature vs non-miniature | Intercept | 2.823346 | 0.373466 | 7.5598 | 4.77E-13 | 0.009929 | 0.0001903 | 309 |
| | log10BodySize | 0.02151 | 0.013105 | 1.6414 | 0.1017 | | | |
| | miniature | -0.010417 | 0.055955 | -0.1862 | 0.8524 | | | |
| | log10BodySize:miniature | 0.037505 | 0.146245 | 0.2565 | 0.7978 | | | |
| log10GenomeSize ~ log10BodySize * non-miniature vs proportioned-dwarf vs progenetic-miniature | Intercept | 2.841875 | 0.346977 | 8.1904 | 7.36E-15 | 0.01907 | 0.00288 | 309 |
| | log10BodySize | 0.021084 | 0.013107 | 1.6087 | 0.1087 | | | |
| | progenetic-miniature | -0.174827 | 0.117309 | -1.4903 | 0.1372 | | | |
| | proportioned-dwarf | 0.033439 | 0.062601 | 0.5342 | 0.5936 | | | |
| | log10BodySize:progenetic-miniature | 0.304054 | 0.389928 | 0.7798 | 0.4361 | | | |
| | log10BodySize:proportioned-dwarf | -0.069796 | 0.163169 | -0.4278 | 0.6691 | | | |
| log10BodySize ~ miniature vs non-miniature | Intercept | 1.002546 | 0.049468 | 20.267 | 2.20E-16 | 0.1827 | 0.1801 | 309 |
| | miniature | -0.59795 | 0.072173 | -8.285 | 3.71E-15 | | | |
| log10BodySize ~ non-miniature vs proportioned-dwarf vs progenetic-miniature | Intercept | 1.016456 | 0.050152 | 20.2677 | 2.20E-16 | 0.1892 | 0.1839 | 309 |
| | progenetic-miniature | -0.810526 | 0.154112 | -5.2593 | 2.72E-07 | | | |
| | proportioned-dwarf | -0.545696 | 0.079413 | -6.8716 | 3.56E-11 | | | |

**Supplementary Table 10.** Comparison of Brownian motion (BM) and Ornstein–Uhlenbeck (OU) model fit within the phylogenetic generalized least squares (PGLS) regression framework for log-transformed genome size and body size across Cypriniformes.

| | | Brownian motion model | | | Ornstein Uhlenbeck model | | | |
|---|---|---|---|---|---|---|---|---|
| **Trait** | **Predictor** | **BM_log likelihood** | **BM_AIC** | **BM_AICw** | **OU_log likelihood** | **OU_AIC** | **OU_AICw** | **delta AIC** |
| log10GenomeSize | miniature vs non-miniature | **344.7** | **-683.4** | **0.95843673** | 342.5 | -677.1 | 0.04156327 | 6.276173 |
| log10GenomeSize | non-miniature vs proportioned-dwarf vs progenetic-miniature | **345.8** | **-683.6** | **0.95611298** | 343.7 | -677.4 | 0.04388702 | 6.162515 |
| log10GenomeSize | diploid vs polyploid | **379.9** | **-753.8** | **0.93879306** | 378.2 | -748.4 | 0.06120694 | 5.460669 |
| log10GenomeSize | miniature vs non-miniature + Ploidy | **380** | **-751.9** | **0.93850085** | 378.2 | -746.5 | 0.06149915 | 5.450521 |
| log10GenomeSize | non-miniature vs proportioned-dwarf vs progenetic-miniature + Ploidy | **380.8** | **-751.7** | **0.93396206** | 379.2 | -746.4 | 0.06603794 | 5.298413 |
| log10GenomeSize | log10BodySize | **346** | **-686** | **0.95472774** | 343.9 | -679.9 | 0.04527226 | 6.097464 |
| log10GenomeSize | log10BodySize + miniature vs non-miniature | **346** | **-684** | **0.95469239** | 343.9 | -677.9 | 0.04530761 | 6.095828 |
| log10GenomeSize | log10BodySize + non-miniature vs proportioned-dwarf vs progenetic-miniature | **347** | **-684** | **0.95226423** | 345 | -678 | 0.04773577 | 5.986323 |
| log10GenomeSize | log10BodySize + miniature vs non-miniature + Ploidy | **380** | **-750.1** | **0.93686869** | 378.3 | -744.7 | 0.06313131 | 5.394653 |
| log10GenomeSize | log10BodySize + non-miniature vs proportioned-dwarf vs progenetic-miniature + Ploidy | **380.9** | **-749.8** | **0.93246352** | 379.3 | -744.5 | 0.06753648 | 5.250324 |
| log10GenomeSize | log10BodySize * miniature vs non-miniature | **346** | **-682.1** | **0.95450069** | 344 | -676 | 0.04549931 | 6.086983 |
| log10GenomeSize | log10BodySize * non-miniature vs proportioned-dwarf vs progenetic-miniature | **347.4** | **-680.7** | **0.95109488** | 345.4 | -674.8 | 0.04890512 | 5.935463 |
| log10BodySize | miniature vs non-miniature | -113.3 | 232.6 | 3.10E-13 | **-83.47** | **174.95** | **1.00E+00** | 57.60758 |
| log10BodySize | non-miniature vs proportioned-dwarf vs progenetic-miniature | -112.9 | 233.7 | 1.37E-13 | **-82.25** | **174.5** | **1.00E+00** | 59.24452 |

**Supplementary Table 11.** Reference genome assemblies used in the analyses of transposable element content and gene architecture across miniature and non-miniature cypriniform species.

| Group | Order | Suborder | Family | Genus | Species | Miniature | Source | Assembly.Accession | Assembly.Name | Annotation.Release.Date | Organism.Sex | Annotation.Name | Assembly.Stats.Total.Sequence.Length | Assembly.Stats.Total.Number.of.Chromosomes | Assembly.Level | Assembly.Release.Date | WGS.project.accession | Assembly.Stats.Contig.N50 | Assembly.Stats.Scaffold.N50 | Assembly.Stats.Number.of.Scaffolds | Assembly.Stats.GC.Percent | Assembly.Sequencing.Tech | Assembly.Submitter | Assembly.BioProject.Accession | Assembly.BioSample.Accession | Publication |
|---|---|---|---|---|---|---|---|---|---|---|---|---|---|---|---|---|---|---|---|---|---|---|---|---|---|---|
| Ingroup | Cypriniformes | Catostomoidei | Catostomidae | Myxocyprinus | asiaticus | non-miniature | NCBI | GCF_019703515.2 | UBuf_Myxa_2 | 25/02/2023 | | GCF_019703515.2-RS_2023_02 | 2340848162 | 50 | Chromosome | 26/10/2022 | JAIGOD02 | 35119607 | 47849308 | 352 | 39 | Oxford Nanopore GridION; Illumina | University at Buffalo | PRJNA739167 | SAMN19774990 | https://doi.org/10.1093/gbe/evab190 |
| Ingroup | Cypriniformes | Catostomoidei | Catostomidae | Xyrauchen | texanus | non-miniature | NCBI | GCF_025860055.1 | RBS_HiC_50CHRs | 25/02/2023 | | GCF_025860055.1-RS_2023_02 | 2186806939 | 50 | Chromosome | 26/10/2022 | JAOTQA01 | 5095822 | 43767000 | 1745 | 39.5 | Oxford Nanopore GridION | University at Buffalo | PRJNA885108 | SAMN31066171 | |
| Ingroup | Cypriniformes | Cobitoidei | Cobitidae | Misgurnus | anguillicaudatus | non-miniature | NCBI | GCF_027580225.2 | ASM2758022v2 | 24/04/2025 | | GCF_027580225.2-RS_2025_04 | 1120296596 | 25 | Chromosome | 13/11/2024 | JALDXJ02 | 41472457 | 41472457 | 60 | 38 | PacBio RSII | Huazhong Agriculture University | PRJNA812369 | SAMN26377316 | https://doi.org/10.1186/s12915-023-01517-1 |
| Ingroup | Cypriniformes | Cobitoidei | Cobitidae | Paramisgurnus | dabryanus | non-miniature | NCBI | GCF_030506205.2 | PD_genome_1.1 | 24/04/2025 | | GCF_030506205.2-RS_2025_04 | 1043469091 | 24 | Chromosome | 03/09/2024 | JAUJEC02 | 1000000 | 41695668 | 219 | 38.5 | Pacbio Hifi; HIC | ihb | PRJNA980909 | SAMN35654119 | https://doi.org/10.1038/s41597-025-04498-8 |
| Ingroup | Cypriniformes | Cobitoidei | Nemacheilidae | Triplophysa | dalaica | non-miniature | NCBI | GCF_015846415.1 | ASM1584641v1 | 01/06/2023 | female | GCF_015846415.1-RS_2023_06 | 607923607 | 25 | Chromosome | 11/12/2020 | JABZFK01 | 9273388 | 23627495 | 50 | 39 | PacBio Sequel II | HeNan Normal University and Wuhan diggers | PRJNA624716 | SAMN14582390 | https://doi.org/10.1093/gbe/evab153 |
| Ingroup | Cypriniformes | Cobitoidei | Nemacheilidae | Triplophysa | rosa | non-miniature | NCBI | GCF_024868665.1 | Trosa_1v2 | 09/06/2023 | female | GCF_024868665.1-RS_2023_06 | 684805028 | 25 | Chromosome | 14/09/2022 | JAFHDT01 | 377582 | 24840772 | 1038 | 39.5 | Illumina; PacBio RSII | the National Natural Science Foundation of China | PRJNA673076 | SAMN16587819 | https://doi.org/10.1111/mec.16700 |
| Ingroup | Cypriniformes | Cobitoidei | Nemacheilidae | Triplophysa | tibetana | non-miniature | NCBI | GCA_008369825.1 | ASM836982v1 | 12/09/2019 | female | Annotation submitted by Gooalgene | 652930317 | 25 | Chromosome | 12/09/2019 | SOYY01 | 2833080 | 24885746 | 292 | 39 | PacBio Sequel; Illumina HiSeq | Gooalgene | PRJNA526292 | SAMN11094431 | https://doi.org/10.1111/1755-0998.13021 |
| Ingroup | Cypriniformes | Cyprinoidei | Cyprinidae | Carassius | auratus | non-miniature | NCBI | GCF_003368295.1 | ASM336829v1 | 18/06/2025 | | GCF_003368295.1-RS_2025_06 | 1820618472 | 59 | Chromosome | 09/08/2018 | QPKE01 | 821153 | 22763433 | 6215 | 37.5 | PacBio RSII | National Institutes of Health | PRJNA481500 | SAMN09670328 | https://doi.org/10.1126/sciadv.aav0547 |
| Ingroup | Cypriniformes | Cyprinoidei | Cyprinidae | Carassius | carassius | non-miniature | NCBI | GCF_963082965.1 | fCarCar2.1 | 07/09/2023 | | GCF_963082965.1-RS_2023_09 | 1684296438 | 50 | Chromosome | 09/08/2023 | CAUJCD01 | 3822391 | 32663704 | 277 | 38.5 | PacBio,Arima2 | WELLCOME SANGER INSTITUTE | PRJEB64837 | SAMEA11296534 | |
| Ingroup | Cypriniformes | Cyprinoidei | Cyprinidae | Carassius | gibelio | non-miniature | NCBI | GCF_023724105.1 | carGib1.2-hapl.c | 25/02/2023 | | GCF_023724105.1-RS_2023_02 | 1583334954 | 50 | Chromosome | 16/06/2022 | JAJQLC01 | 5134100 | 30677984 | 50 | 37.5 | PacBio HiFi; OmniC Illumina reads | IGB Leibniz-Institute of Freshwater Ecology and Inland Fisheries | PRJNA780760 | SAMN23046545 | https://doi.org/10.1038/s41467-022-31515-w |
| Ingroup | Cypriniformes | Cyprinoidei | Cyprinidae | Cirrhinus | molitorella | non-miniature | NCBI | GCA_033026305.1 | C_molitorella v1.0 | 25/10/2023 | | Annotation submitted by Pearl River Fisheries Research Institute Chinese Academy of Fishery Sciences | 1033411071 | 25 | Chromosome | 25/10/2023 | JAUYZG01 | 33095397 | 39863030 | 42 | 36 | PacBio HiFi | Pearl River Fisheries Research Institute Chinese Academy of Fishery Sciences | PRJNA1001164 | SAMN36797183 | https://doi.org/10.1038/s41597-024-04075-5 |
| Ingroup | Cypriniformes | Cyprinoidei | Cyprinidae | Cyprinus | carpio | non-miniature | NCBI | GCF_018340385.1 | ASM1834038v1 | 20/07/2021 | | NCBI Cyprinus carpio Annotation Release 101 | 1680118328 | 50 | Chromosome | 12/05/2021 | JAEOAB01 | 1558716 | 29545497 | 6700 | 37 | PacBio; Oxford Nanopore; Illumina HiSeq | Chinese Academy of Fishery Sciences | PRJNA682709 | SAMN17005855 | https://doi.org/10.1038/s41588-021-00933-9 |
| Ingroup | Cypriniformes | Cyprinoidei | Cyprinidae | Garra | rufa | non-miniature | NCBI | GCF_049309525.1 | GarRuf1.0 | 24/04/2025 | | GCF_049309525.1-RS_2025_04 | 1391285389 | 25 | Chromosome | 05/04/2025 | JBLIWC01 | 132089 | 49275370 | 1000 | 37 | PacBio Revio; DNBSEQ-T7; Illumina NovaSeq X Plus (Hi-C) | University of Vienna | PRJNA1204062 | SAMN46009517 | https://doi.org/10.1038/s41597-025-05101-w |
| Ingroup | Cypriniformes | Cyprinoidei | Cyprinidae | Labeo | rohita | non-miniature | NCBI | GCF_022985175.1 | IGBB_LRoh.1.0 | 26/02/2023 | male | GCF_022985175.1-RS_2023_02 | 1126541939 | 25 | Chromosome | 19/04/2022 | JACTAM01 | 1265157 | 37901974 | 2869 | 36 | Oxford Nanopore GridION; Illumina HiSeq | Mississippi State University | PRJNA650519 | SAMN15716776 | https://doi.org/10.1093/g3journal/jkad009 |
| Ingroup | Cypriniformes | Cyprinoidei | Cyprinidae | Onychostoma | macrolepis | non-miniature | NCBI | GCF_012432095.1 | ASM1243209v1 | 02/08/2023 | female | GCF_012432095.1-RS_2023_08 | 886569841 | 25 | Chromosome | 17/04/2020 | JAAMOB01 | 10809623 | 34067727 | 352 | 37 | Oxford Nanopore PromethION; Illumina HiSeq; Hi-C; Bionano optical mapping | SOUTHWEST UNIVERSITY | PRJNA608076 | SAMN14163390 | https://doi.org/10.1111/1755-0998.13190 |
| Ingroup | Cypriniformes | Cyprinoidei | Cyprinidae | Puntigrus | tetrazona | non-miniature | NCBI | GCF_018831695.1 | ASM1883169v1 | 12/08/2021 | | NCBI Puntigrus tetrazona Annotation Release 100 | 730801986 | 25 | Chromosome | 10/06/2021 | JAEQBD01 | 1423374 | 25085836 | 1121 | 38 | Illumina HiSeq; Nanopore | China Academy of Fishery Sciencey | PRJNA683758 | SAMN17036344 | https://doi.org/10.1038/s41588-021-00933-9 |
| Ingroup | Cypriniformes | Cyprinoidei | Danionidae | Boraras | brigittae | proportioned-dwarf | Sudasinghe et al. (2026) | | LR15871 | | | | | | | | | | | | | | | | | |
| Ingroup | Cypriniformes | Cyprinoidei | Danionidae | Danio | aesculapii | non-miniature | NCBI | GCF_903798145.1 | fDanAes4.1 | 25/05/2023 | | GCF_903798145.1-RS_2023_05 | 1381485597 | 25 | Chromosome | 27/06/2020 | CAIGQR01 | 1517104 | 55996547 | 342 | 36.5 | PacBio,Illumina,BioNano,Dovetail | SC | PRJEB38583 | SAMEA104240212 | |
| Ingroup | Cypriniformes | Cyprinoidei | Danionidae | Danio | rerio | non-miniature | NCBI | GCF_049306965.1 | GRCz12tu | 11/04/2025 | | GCF_049306965.1-RS_2025_04 | 1448791966 | 25 | Complete Genome | 04/04/2025 | JBMGRA01 | 59305174 | 59305174 | 25 | 37 | Oxford Nanopore PromethION; PacBio | National Human Genome Research Institute | PRJNA1193077 | SAMN43570788 | https://doi.org/10.1101/2025.11.17.688901 |
| Ingroup | Cypriniformes | Cyprinoidei | Danionidae | Danionella | cerebrum | progenetic-miniature | NCBI | GCA_007224835.1 | ASM722483v1 | 22/07/2019 | pooled male and female | Annotation submitted by Charite - Universitatsmedizin Berlin | 735303417 | | Scaffold | 22/07/2019 | SRMA01 | 133131 | 340819 | 27639 | 38.5 | Illumina HiSeq; Oxford Nanopore MiniION | Charite - Universitatsmedizin Berlin | PRJNA445947 | SAMN11288624 | https://doi.org/10.1038/s41597-019-0161-z |
| Ingroup | Cypriniformes | Cyprinoidei | Danionidae | Rasbora | kalochroma | non-miniature | Sudasinghe et al. (2026) | | LR15877 | | | | | | | | | | | | | | | | | |
| Ingroup | Cypriniformes | Cyprinoidei | Gobionidae | Pseudorasbora | parva | non-miniature | NCBI | GCF_024679245.1 | ASM2467924v1 | 16/08/2024 | | GCF_024679245.1-RS_2024_08 | 1228852188 | 25 | Chromosome | 16/08/2022 | JANLNJ01 | 4648668 | 47421265 | 80 | 39 | PacBio Sequel; Illumina NovaSeq | The Max Planck Institute of Molecular Cell Biology and Genetics | PRJNA843929 | SAMN28772710 | |
| Ingroup | Cypriniformes | Cyprinoidei | Leuciscidae | Leuciscus | chuanchicus | non-miniature | NCBI | GCA_965140175.1 | ASM96514017v1 | 13/02/2025 | | Annotation submitted by Chinese Academy of Fishery Sciences | 1160198222 | | Scaffold | 14/02/2025 | CBCSLH01 | 31116631 | 43855677 | 130 | 39 | Revio | Chinese Academy of Fishery Sciences | PRJEB85705 | SAMEA117683954 | https://doi.org/10.1038/s41597-025-04787-2 |
| Ingroup | Cypriniformes | Cyprinoidei | Leuciscidae | Leuciscus | waleckii | non-miniature | NCBI | GCA_041200155.1 | ASM4120015v1 | 11/08/2024 | | Annotation submitted by Heilongjiang River Fisheries Research Institute | 1098969243 | 25 | Chromosome | 11/08/2024 | | 1827993 | 42397162 | 25 | 39 | PacBio Sequel | Heilongjiang River Fisheries Research Institute | PRJNA929316 | SAMN32957118 | |
| Ingroup | Cypriniformes | Cyprinoidei | Leuciscidae | Phoxinus | phoxinus | non-miniature | NCBI | GCA_037504875.1 | fPhoPho.hap1 | 25/03/2024 | male | Annotation submitted by Leibniz Institute for the analysis of biodiversity change | 939831526 | 25 | Chromosome | 25/03/2024 | JAYKXG01 | 3852692 | 36381605 | 101 | 39 | PacBio Sequel | Leibniz Institute for the analysis of biodiversity change | PRJNA1030284 | SAMN37908444 | https://doi.org/10.1093/gigascience/giae116 |
| Ingroup | Cypriniformes | Cyprinoidei | Leuciscidae | Pimephales | promelas | non-miniature | NCBI | GCF_016745375.1 | EPA_FHM_2.0 | 19/02/2021 | | NCBI Pimephales promelas Annotation Release 100 | 1066412313 | | Scaffold | 24/01/2021 | WIOS01 | 295770 | 11952773 | 910 | 38.5 | PacBio RSII; Illumina HiSeq | US EPA | PRJNA565199 | SAMN12875914 | https://doi.org/10.1002/etc.5266 |
| Ingroup | Cypriniformes | Cyprinoidei | Paedocyprididae | Paedocypris | sp | progenetic-miniature | Sudasinghe et al. (2026) | | LR15863A | | | | | | | | | | | | | | | | | |
| Ingroup | Cypriniformes | Cyprinoidei | Sundadanionidae | Sundadanio | atomus | progenetic-miniature | Sudasinghe et al. (2026) | | LR15870 | | | | | | | | | | | | | | | | | |
| Ingroup | Cypriniformes | Cyprinoidei | Xenocyprididae | Chanodichthys | erythropterus | non-miniature | NCBI | GCF_024489055.1 | ASM2448905v1 | 16/08/2024 | | GCF_024489055.1-RS_2024_08 | 1085510300 | 24 | Chromosome | 29/07/2022 | JALPSW01 | 23286394 | 42399299 | 50 | 38 | Oxford Nanopore PromethION | Qufu Normal University | PRJNA827856 | SAMN27624467 | https://doi.org/10.1038/s41597-022-01648-0 |
| Ingroup | Cypriniformes | Cyprinoidei | Xenocyprididae | Ctenopharyngodon | idella | non-miniature | NCBI | GCF_019924925.1 | HZGC01 | 25/02/2023 | female | GCF_019924925.1-RS_2023_02 | 893198495 | 24 | Chromosome | 14/09/2021 | JAIKLG01 | 19319444 | 35662356 | 30 | 37.5 | PacBio Sequel | HuaZhong Agricultural University | PRJNA745929 | SAMN20170273 | https://doi.org/10.1186/s12864-022-08503-x |
| Ingroup | Cypriniformes | Cyprinoidei | Xenocyprididae | Culter | alburnus | non-miniature | NCBI | GCA_040182925.1 | ASM4018292v1 | 17/06/2024 | | Annotation submitted by Freshwater Fisheries Research Center, Chinese Academy of Fishery Science | 1052771736 | 24 | Chromosome | 17/06/2024 | JAWDJR01 | 32920630 | 43094527 | 37 | 37.5 | Illumina NovaSeq 6000; PacBio Sequel | Freshwater Fisheries Research Center, Chinese Academy of Fishery Science | PRJNA1019486 | SAMN37486085 | |
| Ingroup | Cypriniformes | Cyprinoidei | Xenocyprididae | Megalobrama | amblycephala | non-miniature | NCBI | GCF_018812025.1 | ASM1881202v1 | 06/05/2022 | female | NCBI Megalobrama amblycephala Annotation Release 100 | 1109871993 | 24 | Chromosome | 09/06/2021 | JAGTWI01 | 2576066 | 42265598 | 242 | 37.5 | PacBio RSII | Huazhong Agricultural University | PRJNA343584 | SAMN18522313 | https://doi.org/10.1093/molbev/msab152 |
| Ingroup | Cypriniformes | Gyrinocheiloidei | Gyrinocheilidae | Gyrinocheilus | aymonieri | non-miniature | NCBI | GCA_051167395.1 | ASM5116739v1 | | | | 507040962 | 24 | Chromosome | 01/07/2025 | JBPGVV01 | 11731257 | 18993287 | 244 | 40.5 | PacBio | Institute of Hydrobiology, Chinese Academy of Sciences | PRJNA1269254 | SAMN48783300 | https://doi.org/10.1038/s41597-025-06070-w |

**Supplementary Table 12.** Phylogenetic signal estimates for transposable element content and gene architecture traits across Cypriniformes, reported as Pagel's λ and Blomberg's K with associated p values.

| Trait | Pagel's λ | Pagel's λ-P-value | Blomberg's K | Blomberg's K-P-value |
|---|---|---|---|---|
| median gene length | 0.802532 | 0.00245881 | 0.373982 | 0.013 |
| overall median intron length | 1.00969 | 3.02E-09 | 1.13564 | 0.001 |
| first (5`) median intron length | 0.992601 | 2.47E-06 | 0.762518 | 0.001 |
| last (3`) median intron length | 0.952229 | 2.47E-05 | 0.647167 | 0.001 |
| mid median intron length | 1.01547 | 1.49E-10 | 1.28209 | 0.001 |
| single median intron length | 0.509037 | 0.152696 | 0.277763 | 0.128 |
| ultra-short introns (≤100 bp) median length | 1.00366 | 1.88E-08 | 0.98057 | 0.003 |
| short introns (101–300 bp) median length | 1.00458 | 1.68E-06 | 1.13143 | 0.001 |
| compact introns (≤300 bp) median length | 1.01666 | 3.97E-12 | 1.38285 | 0.001 |
| long introns (>300 bp) median length | 1.01495 | 1.95E-07 | 0.916031 | 0.001 |
| first (5`) ultra-short introns (≤100 bp) median intron length | 0.908369 | 0.0019015 | 0.376057 | 0.046 |
| last (3`) ultra-short introns (≤100 bp) median intron length | 1.01666 | 5.24E-09 | 0.91926 | 0.003 |
| mid ultra-short introns (≤100 bp) median intron length | 1.00367 | 1.72E-08 | 0.987621 | 0.001 |
| single ultra-short introns (≤100 bp) median intron length | 7.45E-05 | 1 | 0.0779879 | 0.885 |
| first (5`) short introns (101–300 bp) median intron length | 0.482065 | 0.165687 | 0.298931 | 0.028 |
| last (3`) short introns (101–300 bp) median intron length | 0.908692 | 0.00215551 | 0.799045 | 0.001 |
| mid short introns (101–300 bp) median intron length | 1.0078 | 7.40E-07 | 1.17038 | 0.001 |
| single short introns (101–300 bp) median intron length | 7.45E-05 | 1 | 0.190763 | 0.353 |
| first (5`) compact introns (≤300 bp) median intron length | 0.922384 | 0.000444486 | 0.516034 | 0.002 |
| last (3`) compact introns (≤300 bp) median intron length | 0.995408 | 5.28E-09 | 1.25741 | 0.001 |
| mid compact introns (≤300 bp) median intron length | 1.0133 | 1.28E-10 | 1.35911 | 0.001 |
| single compact introns (≤300 bp) median intron length | 0.376595 | 0.574689 | 0.217382 | 0.166 |
| first (5`) long introns (>300 bp) median intron length | 0.995701 | 3.75E-06 | 0.764099 | 0.001 |
| last (3`) long introns (>300 bp) median intron length | 1.01666 | 5.82E-08 | 0.907078 | 0.001 |
| mid long introns (>300 bp) median intron length | 1.01618 | 9.21E-08 | 0.945564 | 0.001 |
| single long introns (>300 bp) median intron length | 0.726836 | 0.0284924 | 0.351664 | 0.012 |
| median number of introns per gene | 7.45E-05 | 1 | 0.107023 | 0.855 |
| median number of mid introns per gene | 7.45E-05 | 1 | 0.252534 | 0.071 |
| proportion of compact introns (≤300 bp) per gene | 1.00073 | 1.30E-10 | 1.61284 | 0.001 |
| proportion of long introns (>300 bp) per gene | 1.00073 | 1.30E-10 | 1.61284 | 0.001 |
| % of non-repititive DNA | 1.01036 | 6.81E-05 | 0.674322 | 0.001 |
| % of total repeat DNA | 1.01036 | 6.81E-05 | 0.674322 | 0.001 |
| % of class I repeats | 0.435864 | 0.624297 | 0.318678 | 0.013 |
| % of class II repeats | 0.957014 | 0.0017664 | 0.563171 | 0.001 |
| % of other repeats | 1.00736 | 1.18E-08 | 1.28411 | 0.001 |
| % of retroelements | 0.435864 | 0.624297 | 0.318678 | 0.022 |
| % of total interspersed repeats | 1.01163 | 2.61E-05 | 0.745715 | 0.001 |
| % of LINEs | 7.45E-05 | 1 | 0.178872 | 0.343 |
| % of SINEs | 1.00095 | 1.03E-06 | 0.774523 | 0.022 |
| % of LTR | 0.996143 | 0.207257 | 0.416149 | 0.004 |
| % of DNA transposons | 0.957351 | 0.00176658 | 0.563588 | 0.001 |
| % of helitrons | 1.01013 | 2.35E-05 | 0.650286 | 0.05 |
| % of small RNA | 0.796993 | 0.010617 | 0.490275 | 0.019 |
| % of satellites | 1.01666 | 0.000207951 | 0.445246 | 0.201 |
| % of simple repeats | 1.01666 | 2.32E-11 | 1.39963 | 0.001 |
| % of low complexity repeats | 1.01666 | 7.03E-13 | 2.28397 | 0.001 |
| % of unclassified repeats | 0.980167 | 4.60E-05 | 0.606783 | 0.001 |
| % of recent (<5% K2P) class I repeats | 7.45E-05 | 1 | 0.309686 | 0.017 |
| % of recent (<5% K2P) class II repeats | 0.963424 | 0.0231938 | 0.474302 | 0.001 |
| % of recent (<5% K2P) other repeats | 1.00932 | 0.189588 | 0.387764 | 0.222 |
| % of recent (<5% K2P) LINEs | 0.910883 | 0.30254 | 0.343248 | 0.033 |
| % of recent (<5% K2P) SINEs | 0.986788 | 6.95E-07 | 0.77696 | 0.012 |

| | | | | |
|---|---|---|---|---|
| % of recent (<5% K2P) LTR | 7.45E-05 | 1 | 0.369941 | 0.011 |
| % of recent (<5% K2P) DNA transposons | 0.944907 | 0.0575844 | 0.457263 | 0.004 |
| % of recent (<5% K2P) helitrons | 1.00512 | 3.12E-05 | 0.632993 | 0.002 |
| % of intermediate (5-20% K2P) class I repeats | 7.45E-05 | 1 | 0.220081 | 0.147 |
| % of intermediate (5-20% K2P) class II repeats | 0.979662 | 0.000848262 | 0.590309 | 0.001 |
| % of intermediate (5-20% K2P) other repeats | 1.01062 | 8.44E-09 | 1.14364 | 0.028 |
| % of intermediate (5-20% K2P) LINEs | 0.498798 | 0.490254 | 0.183523 | 0.32 |
| % of intermediate (5-20% K2P) SINEs | 1.0158 | 3.73E-08 | 0.910588 | 0.013 |
| % of intermediate (5-20% K2P) LTR | 7.45E-05 | 1 | 0.227275 | 0.154 |
| % of intermediate (5-20% K2P) DNA transposons | 1.00911 | 0.000589941 | 0.606215 | 0.001 |
| % of intermediate (5-20% K2P) helitrons | 0.947989 | 0.00377356 | 0.475019 | 0.001 |
| % of ancient (>20% K2P) class I repeats | 7.45E-05 | 1 | 0.302804 | 0.032 |
| % of ancient (>20% K2P) class II repeats | 0.985574 | 2.46E-06 | 0.75023 | 0.001 |
| % of ancient (>20% K2P) other repeats | 1.01646 | 0.00130776 | 0.449547 | 0.009 |
| % of ancient (>20% K2P) LINEs | 7.45E-05 | 1 | 0.216739 | 0.207 |
| % of ancient (>20% K2P) SINEs | 0.867002 | 0.00524729 | 0.406832 | 0.107 |
| % of ancient (>20% K2P) LTR | 1.01661 | 0.0846263 | 0.389663 | 0.015 |
| % of ancient (>20% K2P) DNA transposons | 1.01661 | 4.46E-07 | 0.770712 | 0.001 |
| % of ancient (>20% K2P) helitrons | 0.929811 | 0.00513947 | 0.404155 | 0.026 |

**Supplementary Table 13.** Results of phylogenetic generalized least squares (PGLS) regression analyses for transposable element content and gene architecture traits across Cypriniformes under a Brownian motion (BM) model of trait evolution, with miniaturization status as the categorical predictor.

| Trait | Predictor | Intercept | β (estimate) | SE (Standard error) | t value | p-value | $R^2$ | Adjusted $R^2$ | N |
|---|---|---|---|---|---|---|---|---|---|
| median gene length | miniature vs non-miniature | 5760.5 | 2955 | 1631.3 | 1.8114 | 0.07977 | 0.09572 | 0.06655 | 33 |
| overall median intron length | miniature vs non-miniature | 273.51 | 186.21 | 84.63 | 2.2003 | 0.03537 | 0.1351 | 0.1072 | 33 |
| first (5`) median intron length | miniature vs non-miniature | 625.44 | 449.37 | 144.58 | 3.1082 | 0.004012 | 0.2376 | 0.213 | 33 |
| last (3`) median intron length | miniature vs non-miniature | 248.14 | 378.84 | 193.35 | 1.9594 | 0.05911 | 0.1102 | 0.08149 | 33 |
| mid median intron length | miniature vs non-miniature | 242.305 | 143.579 | 68.917 | 2.0834 | 0.04556 | 0.1228 | 0.09452 | 33 |
| single median intron length | miniature vs non-miniature | 458.31 | 624.58 | 357.88 | 1.7452 | 0.09085 | 0.08946 | 0.06009 | 33 |
| ultra-short introns (≤100 bp) median length | miniature vs non-miniature | 81.0902 | 4.6592 | 1.0816 | 4.3075 | 0.0001543 | 0.3744 | 0.3543 | 33 |
| short introns (101–300 bp) median length | miniature vs non-miniature | 158.1352 | -6.1909 | 1.5537 | -3.9847 | 0.0003812 | 0.3387 | 0.3174 | 33 |
| compact introns (≤300 bp) median length | miniature vs non-miniature | 102.2557 | 10.95 | 3.3347 | 3.2836 | 0.002544 | 0.2581 | 0.2341 | 33 |
| long introns (>300 bp) median length | miniature vs non-miniature | 889.98 | 453.96 | 153.17 | 2.9637 | 0.0057966 | 0.2208 | 0.1957 | 33 |
| first (5`) ultra-short introns (≤100 bp) median intron length | miniature vs non-miniature | 80.8915 | 4.6347 | 1.9151 | 2.4201 | 0.02157 | 0.1589 | 0.1318 | 33 |
| last (3`) ultra-short introns (≤100 bp) median intron length | miniature vs non-miniature | 81.2226 | 4.5666 | 1.2663 | 3.6062 | 0.001077 | 0.2955 | 0.2728 | 33 |
| mid ultra-short introns (≤100 bp) median intron length | miniature vs non-miniature | 81.0869 | 4.6701 | 1.0781 | 4.332 | 0.0001441 | 0.3771 | 0.357 | 33 |
| single ultra-short introns (≤100 bp) median intron length | miniature vs non-miniature | 79.8125 | 2.8433 | 10.8576 | 0.2619 | 0.7952 | 0.002207 | -0.02998 | 33 |
| first (5`) short introns (101–300 bp) median intron length | miniature vs non-miniature | 172.3188 | 3.3047 | 5.5495 | 0.5955 | 0.5558 | 0.01131 | -0.02058 | 33 |
| last (3`) short introns (101–300 bp) median intron length | miniature vs non-miniature | 159.8659 | -5.6716 | 2.6002 | -2.1812 | 0.03688 | 0.1331 | 0.1051 | 33 |
| mid short introns (101–300 bp) median intron length | miniature vs non-miniature | 156.7372 | -6.7037 | 1.4555 | -4.6057 | 6.63E-05 | 0.4063 | 0.3871 | 33 |
| single short introns (101–300 bp) median intron length | miniature vs non-miniature | 182.904 | -14.041 | 11.911 | -1.1788 | 0.2474 | 0.0429 | 0.01203 | 33 |
| first (5`) compact introns (≤300 bp) median intron length | miniature vs non-miniature | 120.8675 | 17.9587 | 9.2845 | 1.9343 | 0.06225 | 0.1077 | 0.07891 | 33 |
| last (3`) compact introns (≤300 bp) median intron length | miniature vs non-miniature | 104.7157 | 12.5398 | 4.6039 | 2.7237 | 0.01051 | 0.1931 | 0.1671 | 33 |
| mid compact introns (≤300 bp) median intron length | miniature vs non-miniature | 101.3276 | 10.7677 | 3.2184 | 3.3457 | 0.002161 | 0.2653 | 0.2416 | 33 |
| single compact introns (≤300 bp) median intron length | miniature vs non-miniature | 117.0336 | 5.8117 | 14.9545 | 0.3886 | 0.7002 | 0.004848 | -0.02725 | 33 |
| first (5`) long introns (>300 bp) median intron length | miniature vs non-miniature | 1126.62 | 597.24 | 194.2 | 3.0753 | 0.004365 | 0.2338 | 0.2091 | 33 |
| last (3`) long introns (>300 bp) median intron length | miniature vs non-miniature | 814.52 | 590.36 | 199.02 | 2.9664 | 0.005758 | 0.2211 | 0.196 | 33 |
| mid long introns (>300 bp) median intron length | miniature vs non-miniature | 873.5 | 411.77 | 146.78 | 2.8054 | 0.0086024 | 0.2025 | 0.1768 | 33 |
| single long introns (>300 bp) median intron length | miniature vs non-miniature | 1078.04 | 586.19 | 331.1 | 1.7704 | 0.08649 | 0.09182 | 0.06253 | 33 |
| median number of introns per gene | miniature vs non-miniature | 6.86998 | -0.23253 | 0.79041 | -0.2942 | 0.7706 | 0.002784 | -0.02938 | 33 |
| median number of mid introns per gene | miniature vs non-miniature | 6.539729 | 0.098227 | 0.561621 | 0.1749 | 0.8623 | 0.0009858 | -0.03124 | 33 |
| proportion of compact introns (≤300 bp) per gene | miniature vs non-miniature | 0.489083 | -0.021446 | 0.022359 | -0.9591 | 0.3449 | 0.02882 | -0.002508 | 33 |
| proportion of long introns (>300 bp) per gene | miniature vs non-miniature | 0.510917 | 0.021446 | 0.022359 | 0.9591 | 0.3449 | 0.02882 | -0.002508 | 33 |
| % of non-repititive DNA | miniature vs non-miniature | 62.6829 | -9.4398 | 6.133 | -1.5392 | 0.1339 | 0.071 | 0.04103 | 33 |
| % of total repeat DNA | miniature vs non-miniature | 37.3171 | 9.4398 | 6.133 | 1.5392 | 0.1339079 | 0.071 | 0.04103 | 33 |
| % of class I repeats | miniature vs non-miniature | 18.1668 | 1.7246 | 5.1411 | 0.3354 | 0.73955 | 0.003617 | -0.02852 | 33 |
| % of class II repeats | miniature vs non-miniature | 13.0937 | 9.4497 | 4.7328 | 1.9966 | 0.05471 | 0.1139 | 0.08536 | 33 |
| % of other repeats | miniature vs non-miniature | 6.05659 | -1.73452 | 0.84801 | -2.0454 | 0.04938 | 0.1189 | 0.09049 | 33 |
| % of retroelements | miniature vs non-miniature | 18.1668 | 1.7246 | 5.1411 | 0.3354 | 0.73955 | 0.003617 | -0.02852 | 33 |
| % of total interspersed repeats | miniature vs non-miniature | 33.8746 | 11.5361 | 6.1733 | 1.8687 | 0.0711398 | 0.1012 | 0.07225 | 33 |
| % of LINEs | miniature vs non-miniature | 6.21825 | -0.97423 | 2.69598 | -0.3614 | 0.7203 | 0.004195 | -0.02793 | 33 |
| % of SINEs | miniature vs non-miniature | 0.61457 | 0.34539 | 0.96903 | 0.3564 | 0.7239 | 0.004081 | -0.02804 | 33 |
| % of LTR | miniature vs non-miniature | 11.334 | 2.3534 | 3.7717 | 0.624 | 0.53722 | 0.0124 | -0.01945 | 33 |
| % of DNA transposons | miniature vs non-miniature | 13.0476 | 9.4899 | 4.7324 | 2.0053 | 0.05373 | 0.1148 | 0.08627 | 33 |
| % of helitrons | miniature vs non-miniature | 0.046154 | -0.0402 | 0.012203 | -3.2944 | 0.002473 | 0.2593 | 0.2354 | 33 |
| % of small RNA | miniature vs non-miniature | 0.255961 | -0.036109 | 0.110487 | -0.3268 | 0.746 | 0.003434 | -0.02871 | 33 |
| % of satellites | miniature vs non-miniature | 1.46E-04 | -1.71E-04 | 3.82E-05 | -4.4808 | 9.45E-05 | 0.3931 | 0.3735 | 33 |
| % of simple repeats | miniature vs non-miniature | 2.87053 | -1.89276 | 0.56842 | -3.3299 | 0.0022527 | 0.2634 | 0.2397 | 33 |
| % of low complexity repeats | miniature vs non-miniature | 0.269772 | -0.127133 | 0.049815 | -2.5521 | 0.0158553 | 0.1736 | 0.147 | 33 |
| % of unclassified repeats | miniature vs non-miniature | 2.66018 | 0.32166 | 0.47096 | 0.683 | 0.4996902 | 0.01482 | -0.01696 | 33 |
| % of recent (<5% K2P) class I repeats | miniature vs non-miniature | 6.571 | -1.5601 | 2.3635 | -0.6601 | 0.51407 | 0.01386 | -0.01795 | 33 |
| % of recent (<5% K2P) class II repeats | miniature vs non-miniature | 1.9705 | 2.4639 | 1.6839 | 1.4632 | 0.1535 | 0.0646 | 0.03443 | 33 |
| % of recent (<5% K2P) other repeats | miniature vs non-miniature | -0.81488 | 1.81493 | 0.49916 | 3.636 | 0.0009932 | 0.299 | 0.2763 | 33 |
| % of recent (<5% K2P) LINEs | miniature vs non-miniature | 2.92467 | -1.28544 | 0.96754 | -1.3286 | 0.19369 | 0.05387 | 0.02335 | 33 |
| % of recent (<5% K2P) SINEs | miniature vs non-miniature | 0.121824 | 0.077911 | 0.146235 | 0.5328 | 0.598 | 0.009074 | -0.02289 | 33 |
| % of recent (<5% K2P) LTR | miniature vs non-miniature | 3.5245 | -0.3526 | 1.851 | -0.1905 | 0.8502 | 0.001169 | -0.03105 | 33 |
| % of recent (<5% K2P) DNA transposons | miniature vs non-miniature | 1.6266 | 2.4957 | 1.6155 | 1.5449 | 0.1325 | 0.07149 | 0.04153 | 33 |
| % of recent (<5% K2P) helitrons | miniature vs non-miniature | 0.343859 | -0.031814 | 0.128197 | -0.2482 | 0.80564 | 0.001983 | -0.03021 | 33 |
| % of intermediate (5-20% K2P) class I repeats | miniature vs non-miniature | 10.6977 | 2.021 | 3.9074 | 0.5172 | 0.60867 | 0.008556 | -0.02343 | 33 |
| % of intermediate (5-20% K2P) class II repeats | miniature vs non-miniature | 7.0056 | 7.9482 | 3.7754 | 2.1053 | 0.04347 | 0.1251 | 0.09687 | 33 |
| % of intermediate (5-20% K2P) other repeats | miniature vs non-miniature | 2.2075 | -0.70616 | 0.45382 | -1.556 | 0.129855 | 0.07244 | 0.04252 | 33 |
| % of intermediate (5-20% K2P) LINEs | miniature vs non-miniature | 3.64957 | -0.62507 | 1.65299 | -0.3781 | 0.7079 | 0.004592 | -0.02752 | 33 |
| % of intermediate (5-20% K2P) SINEs | miniature vs non-miniature | 0.43762 | 0.20708 | 0.68801 | 0.301 | 0.7654 | 0.002914 | -0.02925 | 33 |
| % of intermediate (5-20% K2P) LTR | miniature vs non-miniature | 6.6105 | 2.439 | 3.4158 | 0.714 | 0.4805 | 0.01618 | -0.01556 | 33 |
| % of intermediate (5-20% K2P) DNA transposons | miniature vs non-miniature | 5.2581 | 6.0612 | 2.7516 | 2.2028 | 0.03517 | 0.1353 | 0.1075 | 33 |
| % of intermediate (5-20% K2P) helitrons | miniature vs non-miniature | 1.7475 | 1.887 | 1.4889 | 1.2674 | 0.2144 | 0.04927 | 0.0186 | 33 |
| % of ancient (>20% K2P) class I repeats | miniature vs non-miniature | 4.5681 | 1.4224 | 1.5913 | 0.8939 | 0.37827 | 0.02513 | -0.00632 | 33 |
| % of ancient (>20% K2P) class II repeats | miniature vs non-miniature | 6.36973 | 0.83362 | 1.63719 | 0.5092 | 0.614236 | 0.008294 | -0.0237 | 33 |
| % of ancient (>20% K2P) other repeats | miniature vs non-miniature | 0.63224 | -0.12663 | 0.21796 | -0.581 | 0.56546 | 0.01077 | -0.02114 | 33 |
| % of ancient (>20% K2P) LINEs | miniature vs non-miniature | 1.307504 | -0.019199 | 0.674233 | -0.0285 | 0.9775 | 2.62E-05 | -0.03223 | 33 |
| % of ancient (>20% K2P) SINEs | miniature vs non-miniature | 0.047395 | 0.193171 | 0.391003 | 0.494 | 0.6248 | 0.007812 | -0.02419 | 33 |
| % of ancient (>20% K2P) LTR | miniature vs non-miniature | 3.2132 | 1.2485 | 1.2155 | 1.0271 | 0.3123 | 0.03291 | 0.001717 | 33 |
| % of ancient (>20% K2P) DNA transposons | miniature vs non-miniature | 4.61582 | 0.80958 | 1.25794 | 0.6436 | 0.52459 | 0.01318 | -0.01865 | 33 |
| % of ancient (>20% K2P) helitrons | miniature vs non-miniature | 1.753908 | 0.024039 | 0.790573 | 0.0304 | 0.9759 | 2.98E-05 | -0.03223 | 33 |
| median gene length | progenetic-miniature vs non-miniature | 6021.8 | 2645.8 | 2148.5 | 1.2315 | 0.22772 | 0.04812 | 0.01639 | 32 |
| overall median intron length | progenetic-miniature vs non-miniature | 278.11 | 180.76 | 111.55 | 1.6205 | 0.11559 | 0.08049 | 0.04984 | 32 |
| first (5`) median intron length | progenetic-miniature vs non-miniature | 682.81 | 381.49 | 189.58 | 2.0123 | 0.053243 | 0.1189 | 0.08956 | 32 |
| last (3`) median intron length | progenetic-miniature vs non-miniature | 341.76 | 268.08 | 252.88 | 1.0601 | 0.2976 | 0.03611 | 0.003979 | 32 |
| mid median intron length | progenetic-miniature vs non-miniature | 232.416 | 155.28 | 90.783 | 1.7105 | 0.09751 | 0.08886 | 0.05848 | 32 |
| single median intron length | progenetic-miniature vs non-miniature | 702.31 | 335.89 | 464.42 | 0.7232 | 0.4751 | 0.01714 | -0.01563 | 32 |
| ultra-short introns (≤100 bp) median length | progenetic-miniature vs non-miniature | 78.2938 | 7.9678 | 1.0642 | 7.487 | 2.41E-08 | 0.6514 | 0.6398 | 32 |
| short introns (101–300 bp) median length | progenetic-miniature vs non-miniature | 159.5703 | -7.8889 | 1.9893 | -3.9657 | 0.0004196 | 0.3439 | 0.3221 | 32 |

| | | | | | | | | | |
|---|---|---|---|---|---|---|---|---|---|
| compact introns (≤300 bp) median length | progenetic-miniature vs non-miniature | 94.173 | 20.5133 | 3.4353 | 5.9714 | 1.51E-06 | 0.5431 | 0.5279 | 32 |
| long introns (>300 bp) median length | progenetic-miniature vs non-miniature | 911.05 | 429.04 | 201.78 | 2.1263 | 0.041822 | 0.131 | 0.102 | 32 |
| first (5`) ultra-short introns (≤100 bp) median intron length | progenetic-miniature vs non-miniature | 78.6006 | 7.3452 | 2.4017 | 3.0583 | 0.004652 | 0.2377 | 0.2123 | 32 |
| last (3`) ultra-short introns (≤100 bp) median intron length | progenetic-miniature vs non-miniature | 77.8117 | 8.6023 | 1.2029 | 7.1514 | 5.89E-08 | 0.6303 | 0.618 | 32 |
| mid ultra-short introns (≤100 bp) median intron length | progenetic-miniature vs non-miniature | 78.2871 | 7.9827 | 1.0569 | 7.5532 | 2.02E-08 | 0.6554 | 0.6439 | 32 |
| single ultra-short introns (≤100 bp) median intron length | progenetic-miniature vs non-miniature | 77.1179 | 6.0315 | 14.283 | 0.4223 | 0.6758 | 0.005909 | -0.02723 | 32 |
| first (5`) short introns (101–300 bp) median intron length | progenetic-miniature vs non-miniature | 169.911 | 6.1535 | 7.2694 | 0.8465 | 0.404 | 0.02333 | -0.009228 | 32 |
| last (3`) short introns (101–300 bp) median intron length | progenetic-miniature vs non-miniature | 160.4558 | -6.3695 | 3.4217 | -1.8615 | 0.0725 | 0.1035 | 0.07366 | 32 |
| mid short introns (101–300 bp) median intron length | progenetic-miniature vs non-miniature | 157.8575 | -8.0292 | 1.8806 | -4.2695 | 0.0001812 | 0.378 | 0.3572 | 32 |
| single short introns (101–300 bp) median intron length | progenetic-miniature vs non-miniature | 169.094 | 2.2985 | 14.9852 | 0.1534 | 0.8791 | 0.0007836 | -0.03252 | 32 |
| first (5`) compact introns (≤300 bp) median intron length | progenetic-miniature vs non-miniature | 109.093 | 31.89 | 11.568 | 2.7567 | 0.009838 | 0.2021 | 0.1755 | 32 |
| last (3`) compact introns (≤300 bp) median intron length | progenetic-miniature vs non-miniature | 96.3233 | 22.4694 | 5.3592 | 4.1927 | 0.0002243 | 0.3695 | 0.3484 | 32 |
| mid compact introns (≤300 bp) median intron length | progenetic-miniature vs non-miniature | 93.8584 | 19.6049 | 3.4022 | 5.7624 | 2.71E-06 | 0.5254 | 0.5095 | 32 |
| single compact introns (≤300 bp) median intron length | progenetic-miniature vs non-miniature | 97.517 | 28.903 | 18.567 | 1.5567 | 0.1300332 | 0.07474 | 0.0439 | 32 |
| first (5`) long introns (>300 bp) median intron length | progenetic-miniature vs non-miniature | 1197.41 | 513.49 | 254.87 | 2.0148 | 0.0529699 | 0.1192 | 0.08982 | 32 |
| last (3`) long introns (>300 bp) median intron length | progenetic-miniature vs non-miniature | 878.49 | 514.68 | 261.44 | 1.9686 | 0.0583 | 0.1144 | 0.08488 | 32 |
| mid long introns (>300 bp) median intron length | progenetic-miniature vs non-miniature | 877.45 | 407.1 | 193.47 | 2.1042 | 0.0438477 | 0.1286 | 0.09956 | 32 |
| single long introns (>300 bp) median intron length | progenetic-miniature vs non-miniature | 1209.56 | 430.58 | 434.17 | 0.9917 | 0.32926 | 0.03174 | -0.0005315 | 32 |
| median number of introns per gene | progenetic-miniature vs non-miniature | 6.47319 | 0.23695 | 1.03316 | 0.2293 | 0.8202 | 0.00175 | -0.03152 | 32 |
| median number of mid introns per gene | progenetic-miniature vs non-miniature | 6.541825 | 0.095746 | 0.740315 | 0.1293 | 0.898 | 0.0005572 | -0.03276 | 32 |
| proportion of compact introns (≤300 bp) per gene | progenetic-miniature vs non-miniature | 0.495843 | -0.029443 | 0.029384 | -1.002 | 0.3244 | 0.03238 | 0.0001289 | 32 |
| proportion of long introns (>300 bp) per gene | progenetic-miniature vs non-miniature | 0.504157 | 0.029443 | 0.029384 | 1.002 | 0.3244 | 0.03238 | 0.0001289 | 32 |
| % of non-repititive DNA | progenetic-miniature vs non-miniature | 66.4331 | -13.8769 | 7.9836 | -1.7382 | 0.09244 | 0.09149 | 0.06121 | 32 |
| % of total repeat DNA | progenetic-miniature vs non-miniature | 33.5669 | 13.8769 | 7.9836 | 1.7382 | 0.092435 | 0.09149 | 0.06121 | 32 |
| % of class I repeats | progenetic-miniature vs non-miniature | 14.348 | 6.2429 | 6.6518 | 0.9385 | 0.35547 | 0.02852 | -0.003859 | 32 |
| % of class II repeats | progenetic-miniature vs non-miniature | 11.9181 | 10.8407 | 6.2259 | 1.7412 | 0.09189 | 0.09179 | 0.06151 | 32 |
| % of other repeats | progenetic-miniature vs non-miniature | 7.3009 | -3.2067 | 1.035 | -3.0982 | 0.004204 | 0.2424 | 0.2171 | 32 |
| % of retroelements | progenetic-miniature vs non-miniature | 14.348 | 6.2429 | 6.6518 | 0.9385 | 0.35547 | 0.02852 | -0.003859 | 32 |
| % of total interspersed repeats | progenetic-miniature vs non-miniature | 29.2872 | 16.9637 | 7.9873 | 2.1238 | 0.042042 | 0.1307 | 0.1017 | 32 |
| % of LINEs | progenetic-miniature vs non-miniature | 6.8904 | -1.7695 | 3.5464 | -0.499 | 0.6214 | 0.00823 | -0.02483 | 32 |
| % of SINEs | progenetic-miniature vs non-miniature | 0.44367 | 0.5476 | 1.27604 | 0.4291 | 0.6709 | 0.006101 | -0.02703 | 32 |
| % of LTR | progenetic-miniature vs non-miniature | 7.0139 | 7.4648 | 4.7508 | 1.5713 | 0.1266 | 0.07604 | 0.04524 | 32 |
| % of DNA transposons | progenetic-miniature vs non-miniature | 11.8587 | 10.8966 | 6.225 | 1.7504 | 0.09026 | 0.09267 | 0.06243 | 32 |
| % of helitrons | progenetic-miniature vs non-miniature | 0.059397 | -0.055868 | 0.015445 | -3.6173 | 0.00108 | 0.3037 | 0.2805 | 32 |
| % of small RNA | progenetic-miniature vs non-miniature | 0.243218 | -0.021032 | 0.145577 | -0.1445 | 0.8861 | 0.0006952 | -0.03261 | 32 |
| % of satellites | progenetic-miniature vs non-miniature | -2.07E-06 | 4.01E-06 | 3.40E-06 | 1.1816 | 0.2466 | 0.04447 | 0.01262 | 32 |
| % of simple repeats | progenetic-miniature vs non-miniature | 3.65344 | -2.81907 | 0.7006 | -4.0238 | 0.0003577 | 0.3505 | 0.3289 | 32 |
| % of low complexity repeats | progenetic-miniature vs non-miniature | 0.323591 | -0.19081 | 0.063075 | -3.0252 | 0.005059 | 0.2337 | 0.2082 | 32 |
| % of unclassified repeats | progenetic-miniature vs non-miniature | 3.08063 | -0.1758 | 0.60419 | -0.291 | 0.773075 | 0.002814 | -0.03043 | 32 |
| % of recent (<5% K2P) class I repeats | progenetic-miniature vs non-miniature | 5.0783 | 0.206 | 3.0741 | 0.067 | 0.947 | 0.0001497 | -0.03318 | 32 |
| % of recent (<5% K2P) class II repeats | progenetic-miniature vs non-miniature | 1.4116 | 3.1251 | 2.2115 | 1.4131 | 0.1679 | 0.06241 | 0.03116 | 32 |
| % of recent (<5% K2P) other repeats | progenetic-miniature vs non-miniature | 0.17584 | 0.64275 | 0.56563 | 1.1363 | 0.2648 | 0.04127 | 0.009308 | 32 |
| % of recent (<5% K2P) LINEs | progenetic-miniature vs non-miniature | 3.5599 | -2.037 | 1.257 | -1.6205 | 0.1156 | 0.08048 | 0.04983 | 32 |
| % of recent (<5% K2P) SINEs | progenetic-miniature vs non-miniature | 0.039282 | 0.175572 | 0.190718 | 0.9206 | 0.3646 | 0.02747 | -0.004944 | 32 |
| % of recent (<5% K2P) LTR | progenetic-miniature vs non-miniature | 1.4791 | 2.0674 | 2.3392 | 0.8838 | 0.3838 | 0.02538 | -0.00711 | 32 |
| % of recent (<5% K2P) DNA transposons | progenetic-miniature vs non-miniature | 1.1627 | 3.0445 | 2.1236 | 1.4336 | 0.162 | 0.06412 | 0.03292 | 32 |
| % of recent (<5% K2P) helitrons | progenetic-miniature vs non-miniature | 0.248851 | 0.080596 | 0.165883 | 0.4859 | 0.6306 | 0.007807 | -0.02527 | 32 |
| % of intermediate (5-20% K2P) class I repeats | progenetic-miniature vs non-miniature | 8.6221 | 4.4767 | 5.1023 | 0.8774 | 0.3872 | 0.02502 | -0.007481 | 32 |
| % of intermediate (5-20% K2P) class II repeats | progenetic-miniature vs non-miniature | 5.5782 | 9.6371 | 4.953 | 1.9457 | 0.06111 | 0.1121 | 0.08245 | 32 |
| % of intermediate (5-20% K2P) other repeats | progenetic-miniature vs non-miniature | 2.84707 | -1.46287 | 0.55747 | -2.6241 | 0.01353 | 0.1867 | 0.1596 | 32 |
| % of intermediate (5-20% K2P) LINEs | progenetic-miniature vs non-miniature | 4.0823 | -1.1371 | 2.174 | -0.5231 | 0.6048 | 0.009037 | -0.02399 | 32 |
| % of intermediate (5-20% K2P) SINEs | progenetic-miniature vs non-miniature | 0.19531 | 0.49377 | 0.90318 | 0.5467 | 0.5886 | 0.009865 | -0.02314 | 32 |
| % of intermediate (5-20% K2P) LTR | progenetic-miniature vs non-miniature | 4.3445 | 5.1201 | 4.4365 | 1.1541 | 0.2576 | 0.04251 | 0.01059 | 32 |
| % of intermediate (5-20% K2P) DNA transposons | progenetic-miniature vs non-miniature | 4.8014 | 6.6016 | 3.6237 | 1.8218 | 0.07848 | 0.09961 | 0.06959 | 32 |
| % of intermediate (5-20% K2P) helitrons | progenetic-miniature vs non-miniature | 0.7768 | 3.0355 | 1.9347 | 1.569 | 0.1271 | 0.07583 | 0.04503 | 32 |
| % of ancient (>20% K2P) class I repeats | progenetic-miniature vs non-miniature | 4.2552 | 1.7926 | 2.0949 | 0.8557 | 0.39895 | 0.02383 | -0.008714 | 32 |
| % of ancient (>20% K2P) class II repeats | progenetic-miniature vs non-miniature | 7.04772 | 0.03144 | 2.14581 | 0.0147 | 0.988407 | 7.16E-06 | -0.03333 | 32 |
| % of ancient (>20% K2P) other repeats | progenetic-miniature vs non-miniature | 0.64769 | -0.14491 | 0.28727 | -0.5044 | 0.61763 | 0.008411 | -0.02464 | 32 |
| % of ancient (>20% K2P) LINEs | progenetic-miniature vs non-miniature | 1.55262 | -0.30921 | 0.88486 | -0.3495 | 0.7292 | 0.004054 | -0.02914 | 32 |
| % of ancient (>20% K2P) SINEs | progenetic-miniature vs non-miniature | 0.031162 | 0.212378 | 0.515381 | 0.4121 | 0.6832 | 0.005628 | -0.02752 | 32 |
| % of ancient (>20% K2P) LTR | progenetic-miniature vs non-miniature | 2.6715 | 1.8895 | 1.5916 | 1.1871 | 0.2445 | 0.04487 | 0.01303 | 32 |
| % of ancient (>20% K2P) DNA transposons | progenetic-miniature vs non-miniature | 5.256079 | 0.052044 | 1.643893 | 0.0317 | 0.97495 | 3.34E-05 | -0.0333 | 32 |
| % of ancient (>20% K2P) helitrons | progenetic-miniature vs non-miniature | 1.79164 | -0.020605 | 1.042035 | -0.0198 | 0.9844 | 1.30E-05 | -0.03332 | 32 |

**Supplementary Table 14.** Results of phylogenetic generalized least squares (PGLS) regression analyses for transposable element content and gene architecture traits across Cypriniformes under an Ornstein–Uhlenbeck (OU) model of trait evolution, with miniaturization status as the categorical predictor.

| Trait | Predictor | Intercept | β (estimate) | SE (Standard error) | t value | p-value | $R^2$ | Adjusted $R^2$ | N |
|---|---|---|---|---|---|---|---|---|---|
| median gene length | miniature vs non-miniature | 6553.6 | 2433.5 | 1291.6 | 1.8841 | 0.06897 | 0.1027 | 0.0738 | 33 |
| overall median intron length | miniature vs non-miniature | 284.74 | 178.165 | 85.274 | 2.0893 | 0.04498 | 0.1234 | 0.09516 | 33 |
| first (5`) median intron length | miniature vs non-miniature | 657.53 | 424.69 | 144.34 | 2.9423 | 0.006118 | 0.2183 | 0.1931 | 33 |
| last (3`) median intron length | miniature vs non-miniature | 365.03 | 281.77 | 185.51 | 1.5189 | 0.1389 | 0.06926 | 0.03924 | 33 |
| mid median intron length | miniature vs non-miniature | 248.927 | 139.069 | 69.576 | 1.9988 | 0.05446 | 0.1142 | 0.08559 | 33 |
| single median intron length | miniature vs non-miniature | 956.21 | 146.86 | 244.79 | 0.6 | 0.552898 | 0.01148 | -0.02041 | 33 |
| ultra-short introns (≤100 bp) median length | miniature vs non-miniature | 80.7725 | 4.9859 | 1.0848 | 4.5962 | 6.81E-05 | 0.4053 | 0.3861 | 33 |
| short introns (101–300 bp) median length | miniature vs non-miniature | 157.7984 | -6.0813 | 1.5463 | -3.9329 | 0.0004401 | 0.3329 | 0.3113 | 33 |
| compact introns (≤300 bp) median length | miniature vs non-miniature | 101.621 | 11.45 | 3.369 | 3.3985 | 0.001879 | 0.2714 | 0.2479 | 33 |
| long introns (>300 bp) median length | miniature vs non-miniature | 918.56 | 435.65 | 153.81 | 2.8324 | 0.008048 | 0.2056 | 0.18 | 33 |
| first (5`) ultra-short introns (≤100 bp) median intron length | miniature vs non-miniature | 79.5 | 6.175 | 1.00612 | 6.1374 | 8.30E-07 | 0.5486 | 0.534 | 33 |
| last (3`) ultra-short introns (≤100 bp) median intron length | miniature vs non-miniature | 80.788 | 5.0011 | 1.2675 | 3.9455 | 0.0004249 | 0.3343 | 0.3128 | 33 |
| mid ultra-short introns (≤100 bp) median intron length | miniature vs non-miniature | 80.7738 | 4.9929 | 1.0814 | 4.6169 | 6.42E-05 | 0.4074 | 0.3883 | 33 |
| single ultra-short introns (≤100 bp) median intron length | miniature vs non-miniature | 78.75 | 3.7901 | 3.5103 | 1.0797 | 0.2886 | 0.03624 | 0.005154 | 33 |
| first (5`) short introns (101–300 bp) median intron length | miniature vs non-miniature | 165.9076 | 8.1078 | 3.1646 | 2.562 | 0.01549 | 0.1747 | 0.1481 | 33 |
| last (3`) short introns (101–300 bp) median intron length | miniature vs non-miniature | 157.7819 | -4.8387 | 2.303 | -2.101 | 0.04387 | 0.1246 | 0.09641 | 33 |
| mid short introns (101–300 bp) median intron length | miniature vs non-miniature | 156.4888 | -6.6116 | 1.4589 | -4.5321 | 8.17E-05 | 0.3985 | 0.3791 | 33 |
| single short introns (101–300 bp) median intron length | miniature vs non-miniature | 175 | -7.545 | 5.8181 | -1.2968 | 0.2043 | 0.05146 | 0.02086 | 33 |
| first (5`) compact introns (≤300 bp) median intron length | miniature vs non-miniature | 104.633 | 32.9961 | 5.9607 | 5.5356 | 4.62E-06 | 0.4971 | 0.4809 | 33 |
| last (3`) compact introns (≤300 bp) median intron length | miniature vs non-miniature | 103.3324 | 13.5511 | 4.6259 | 2.9294 | 0.006319 | 0.2168 | 0.1915 | 33 |
| mid compact introns (≤300 bp) median intron length | miniature vs non-miniature | 100.7136 | 11.2389 | 3.2511 | 3.457 | 0.001608 | 0.2782 | 0.255 | 33 |
| single compact introns (≤300 bp) median intron length | miniature vs non-miniature | 104.3751 | 21.0359 | 7.3408 | 2.8656 | 0.007412 | 0.2094 | 0.1839 | 33 |
| first (5`) long introns (>300 bp) median intron length | miniature vs non-miniature | 1172.23 | 564.17 | 193.88 | 2.9099 | 0.006636 | 0.2145 | 0.1892 | 33 |
| last (3`) long introns (>300 bp) median intron length | miniature vs non-miniature | 860.55 | 558.48 | 199.5 | 2.7994 | 0.008732 | 0.2018 | 0.176 | 33 |
| mid long introns (>300 bp) median intron length | miniature vs non-miniature | 899.44 | 396.25 | 147.4 | 2.6882 | 0.01145 | 0.189 | 0.1629 | 33 |
| single long introns (>300 bp) median intron length | miniature vs non-miniature | 1407.36 | 300.62 | 255.56 | 1.1763 | 0.2484 | 0.04273 | 0.01185 | 33 |
| median number of introns per gene | miniature vs non-miniature | 6.5 | 0.13323 | 0.30012 | 0.4439 | 0.6602 | 0.006317 | -0.02574 | 33 |
| median number of mid introns per gene | miniature vs non-miniature | 6.25024 | 0.14972 | 0.29185 | 0.513 | 0.6116 | 0.008418 | -0.02357 | 33 |
| proportion of compact introns (≤300 bp) per gene | miniature vs non-miniature | 0.486839 | -0.020464 | 0.02258 | -0.9063 | 0.3718 | 0.02581 | -0.005614 | 33 |
| proportion of long introns (>300 bp) per gene | miniature vs non-miniature | 0.513161 | 0.020464 | 0.02258 | 0.9063 | 0.3718 | 0.02581 | -0.005614 | 33 |
| % of non-repititive DNA | miniature vs non-miniature | 60.7506 | -9.0898 | 5.8063 | -1.5655 | 0.1276 | 0.07327 | 0.04337 | 33 |
| % of total repeat DNA | miniature vs non-miniature | 39.2494 | 9.0898 | 5.8063 | 1.5655 | 0.1276 | 0.07327 | 0.04337 | 33 |
| % of class I repeats | miniature vs non-miniature | 18.7564 | 2.7343 | 3.3736 | 0.8105 | 0.4238 | 0.02075 | -0.01084 | 33 |
| % of class II repeats | miniature vs non-miniature | 15.2929 | 8.9797 | 4.1526 | 2.1624 | 0.038416 | 0.1311 | 0.103 | 33 |
| % of other repeats | miniature vs non-miniature | 6.22176 | -2.02216 | 0.84678 | -2.3881 | 0.023216 | 0.1554 | 0.1281 | 33 |
| % of retroelements | miniature vs non-miniature | 18.7564 | 2.7343 | 3.3736 | 0.8105 | 0.4238 | 0.02075 | -0.01084 | 33 |
| % of total interspersed repeats | miniature vs non-miniature | 35.4314 | 11.6487 | 5.9227 | 1.9668 | 0.0582131 | 0.1109 | 0.08226 | 33 |
| % of LINEs | miniature vs non-miniature | 6.9164 | -1.4028 | 1.291 | -1.0867 | 0.2856 | 0.03669 | 0.005619 | 33 |
| % of SINEs | miniature vs non-miniature | 0.78738 | 0.25555 | 0.9352 | 0.2733 | 0.7865 | 0.002403 | -0.02978 | 33 |
| % of LTR | miniature vs non-miniature | 10.5912 | 4.2241 | 2.8443 | 1.4851 | 0.1476177 | 0.06642 | 0.0363 | 33 |
| % of DNA transposons | miniature vs non-miniature | 15.2467 | 9.0191 | 4.1525 | 2.172 | 0.037628 | 0.1321 | 0.1041 | 33 |
| % of helitrons | miniature vs non-miniature | 0.046005 | -0.039916 | 0.012094 | -3.3006 | 0.002433 | 0.26 | 0.2362 | 33 |
| % of small RNA | miniature vs non-miniature | 0.168196 | -0.054382 | 0.069043 | -0.7877 | 0.43687 | 0.01962 | -0.012 | 33 |
| % of satellites | miniature vs non-miniature | 1.31E-04 | -1.52E-04 | 3.78E-05 | -4.0075 | 0.0003577 | 0.3413 | 0.32 | 33 |
| % of simple repeats | miniature vs non-miniature | 2.90529 | -1.97159 | 0.57331 | -3.4389 | 0.001687 | 0.2761 | 0.2528 | 33 |
| % of low complexity repeats | miniature vs non-miniature | 0.269832 | -0.130838 | 0.050404 | -2.5958 | 0.0143 | 0.1786 | 0.1521 | 33 |
| % of unclassified repeats | miniature vs non-miniature | 2.971195 | 0.017205 | 0.437854 | 0.0393 | 0.9689 | 4.98E-05 | -0.03221 | 33 |
| % of recent (<5% K2P) class I repeats | miniature vs non-miniature | 6.77744 | -0.52856 | 1.45396 | -0.3635 | 0.7187 | 0.004245 | -0.02788 | 33 |
| % of recent (<5% K2P) class II repeats | miniature vs non-miniature | 2.7953 | 2.0108 | 1.4059 | 1.4302 | 0.16266 | 0.0619 | 0.03164 | 33 |
| % of recent (<5% K2P) other repeats | miniature vs non-miniature | -0.51353 | 1.4413 | 0.49031 | 2.9396 | 0.00616 | 0.218 | 0.1928 | 33 |
| % of recent (<5% K2P) LINEs | miniature vs non-miniature | 3.11065 | -1.1898 | 0.65575 | -1.8144 | 0.0793 | 0.096 | 0.06684 | 33 |
| % of recent (<5% K2P) SINEs | miniature vs non-miniature | 0.142848 | 0.070457 | 0.140251 | 0.5024 | 0.619 | 0.008075 | -0.02392 | 33 |
| % of recent (<5% K2P) LTR | miniature vs non-miniature | 3.45711 | 0.56177 | 1.29416 | 0.4341 | 0.667236 | 0.006042 | -0.02602 | 33 |
| % of recent (<5% K2P) DNA transposons | miniature vs non-miniature | 2.4981 | 2.0061 | 1.3164 | 1.524 | 0.13764 | 0.0697 | 0.03969 | 33 |
| % of recent (<5% K2P) helitrons | miniature vs non-miniature | 0.347891 | -0.028669 | 0.119315 | -0.2403 | 0.8117 | 0.001859 | -0.03034 | 33 |
| % of intermediate (5-20% K2P) class I repeats | miniature vs non-miniature | 10.8397 | 1.5017 | 2.1577 | 0.696 | 0.4916 | 0.01538 | -0.01638 | 33 |
| % of intermediate (5-20% K2P) class II repeats | miniature vs non-miniature | 8.7812 | 7.4085 | 3.3593 | 2.2054 | 0.03497 | 0.1356 | 0.1077 | 33 |
| % of intermediate (5-20% K2P) other repeats | miniature vs non-miniature | 2.23827 | -0.75211 | 0.45843 | -1.6406 | 0.111 | 0.07989 | 0.05021 | 33 |
| % of intermediate (5-20% K2P) LINEs | miniature vs non-miniature | 4.08796 | -1.26257 | 0.80249 | -1.5733 | 0.1258 | 0.07395 | 0.04407 | 33 |
| % of intermediate (5-20% K2P) SINEs | miniature vs non-miniature | 0.50799 | 0.18195 | 0.68005 | 0.2676 | 0.7908 | 0.002304 | -0.02988 | 33 |
| % of intermediate (5-20% K2P) LTR | miniature vs non-miniature | 5.864 | 2.918 | 1.8657 | 1.564 | 0.127972 | 0.07313 | 0.04324 | 33 |
| % of intermediate (5-20% K2P) DNA transposons | miniature vs non-miniature | 6.4963 | 5.6251 | 2.5062 | 2.2445 | 0.03208 | 0.1398 | 0.112 | 33 |
| % of intermediate (5-20% K2P) helitrons | miniature vs non-miniature | 2.2852 | 1.7919 | 1.188 | 1.5083 | 0.1416 | 0.06837 | 0.03832 | 33 |
| % of ancient (>20% K2P) class I repeats | miniature vs non-miniature | 4.9934 | 1.27101 | 1.04385 | 1.2176 | 0.2326 | 0.04564 | 0.01486 | 33 |
| % of ancient (>20% K2P) class II repeats | miniature vs non-miniature | 6.44769 | 0.95619 | 1.59638 | 0.599 | 0.55354 | 0.01144 | -0.02045 | 33 |
| % of ancient (>20% K2P) other repeats | miniature vs non-miniature | 0.68907 | -0.16709 | 0.1753 | -0.9531 | 0.3478883 | 0.02847 | -0.002868 | 33 |
| % of ancient (>20% K2P) LINEs | miniature vs non-miniature | 1.60402 | -0.24238 | 0.35738 | -0.6782 | 0.5027 | 0.01462 | -0.01716 | 33 |
| % of ancient (>20% K2P) SINEs | miniature vs non-miniature | 0.201797 | 0.080626 | 0.309689 | 0.2603 | 0.7963 | 0.002182 | -0.03001 | 33 |
| % of ancient (>20% K2P) LTR | miniature vs non-miniature | 3.14378 | 1.48696 | 0.94241 | 1.5778 | 0.124755 | 0.07434 | 0.04448 | 33 |
| % of ancient (>20% K2P) DNA transposons | miniature vs non-miniature | 4.68757 | 0.85802 | 1.23946 | 0.6923 | 0.49393 | 0.01522 | -0.01654 | 33 |
| % of ancient (>20% K2P) helitrons | miniature vs non-miniature | 1.67531 | 0.25061 | 0.62248 | 0.4026 | 0.690011 | 0.005201 | -0.02689 | 33 |
| median gene length | progenetic-miniature vs non-miniature | 6673.5 | 2302.6 | 1520.5 | 1.5144 | 0.1404 | 0.07101 | 0.04005 | 32 |
| overall median intron length | progenetic-miniature vs non-miniature | 289.66 | 172.62 | 110.74 | 1.5588 | 0.1295 | 0.07493 | 0.04409 | 32 |
| first (5`) median intron length | progenetic-miniature vs non-miniature | 709.07 | 364.36 | 184.33 | 1.9767 | 0.05733 | 0.1152 | 0.08574 | 32 |
| last (3`) median intron length | progenetic-miniature vs non-miniature | 427.36 | 208.12 | 230.01 | 0.9048 | 0.3728 | 0.02657 | -0.005882 | 32 |
| mid median intron length | progenetic-miniature vs non-miniature | 239.85 | 149.927 | 90.601 | 1.6548 | 0.1084 | 0.08364 | 0.0531 | 32 |
| single median intron length | progenetic-miniature vs non-miniature | 966.69 | 138.37 | 285.55 | 0.4846 | 0.6314911 | 0.007767 | -0.02531 | 32 |
| ultra-short introns (≤100 bp) median length | progenetic-miniature vs non-miniature | 78.1422 | 8.0863 | 1.046 | 7.7309 | 1.26E-08 | 0.6658 | 0.6547 | 32 |
| short introns (101–300 bp) median length | progenetic-miniature vs non-miniature | 159.102 | -7.6148 | 1.9216 | -3.9628 | 0.0004229 | 0.3436 | 0.3217 | 32 |

| | | | | | | | | | |
|---|---|---|---|---|---|---|---|---|---|
| compact introns (≤300 bp) median length | progenetic-miniature vs non-miniature | 93.8708 | 20.6779 | 3.4514 | 5.9912 | 1.43E-06 | 0.5447 | 0.5296 | 32 |
| long introns (>300 bp) median length | progenetic-miniature vs non-miniature | 936.75 | 414.87 | 198.55 | 2.0895 | 0.04524 | 0.127 | 0.09795 | 32 |
| first (5`) ultra-short introns (≤100 bp) median intron length | progenetic-miniature vs non-miniature | 78 | 7.67245 | 1.00682 | 7.6205 | 1.69E-08 | 0.6594 | 0.648 | 32 |
| last (3`) ultra-short introns (≤100 bp) median intron length | progenetic-miniature vs non-miniature | 77.5983 | 8.7553 | 1.1743 | 7.4558 | 2.61E-08 | 0.6495 | 0.6378 | 32 |
| mid ultra-short introns (≤100 bp) median intron length | progenetic-miniature vs non-miniature | 78.1396 | 8.0986 | 1.0396 | 7.7897 | 1.08E-08 | 0.6692 | 0.6581 | 32 |
| single ultra-short introns (≤100 bp) median intron length | progenetic-miniature vs non-miniature | 76.5 | 6.0401 | 3.9629 | 1.5242 | 0.1379 | 0.07187 | 0.04093 | 32 |
| first (5`) short introns (101–300 bp) median intron length | progenetic-miniature vs non-miniature | 166.6667 | 7.3468 | 3.6352 | 2.021 | 0.05228 | 0.1198 | 0.0905 | 32 |
| last (3`) short introns (101–300 bp) median intron length | progenetic-miniature vs non-miniature | 158.2623 | -5.4271 | 2.749 | -1.9742 | 0.05762 | 0.115 | 0.08548 | 32 |
| mid short introns (101–300 bp) median intron length | progenetic-miniature vs non-miniature | 157.5322 | -7.8438 | 1.8439 | -4.2539 | 0.0001893 | 0.3762 | 0.3554 | 32 |
| single short introns (101–300 bp) median intron length | progenetic-miniature vs non-miniature | 166.3333 | 1.1217 | 5.8407 | 0.1921 | 0.849 | 0.001228 | -0.03206 | 32 |
| first (5`) compact introns (≤300 bp) median intron length | progenetic-miniature vs non-miniature | 100.883 | 36.9138 | 6.9612 | 5.3028 | 9.91E-06 | 0.4838 | 0.4666 | 32 |
| last (3`) compact introns (≤300 bp) median intron length | progenetic-miniature vs non-miniature | 95.3678 | 22.974 | 5.2942 | 4.3394 | 0.0001491 | 0.3856 | 0.3652 | 32 |
| mid compact introns (≤300 bp) median intron length | progenetic-miniature vs non-miniature | 93.544 | 19.7718 | 3.4153 | 5.7891 | 2.52E-06 | 0.5277 | 0.5119 | 32 |
| single compact introns (≤300 bp) median intron length | progenetic-miniature vs non-miniature | 93.5 | 31.853 | 7.3569 | 4.3296 | 0.0001533 | 0.3846 | 0.364 | 32 |
| first (5`) long introns (>300 bp) median intron length | progenetic-miniature vs non-miniature | 1234.73 | 491.08 | 247.75 | 1.9822 | 0.05669 | 0.1158 | 0.08633 | 32 |
| last (3`) long introns (>300 bp) median intron length | progenetic-miniature vs non-miniature | 916.47 | 493.05 | 256.38 | 1.9231 | 0.064 | 0.1098 | 0.08008 | 32 |
| mid long introns (>300 bp) median intron length | progenetic-miniature vs non-miniature | 902.14 | 393.84 | 190.36 | 2.0689 | 0.04726 | 0.1249 | 0.0957 | 32 |
| single long introns (>300 bp) median intron length | progenetic-miniature vs non-miniature | 1425.02 | 280.91 | 300.48 | 0.9349 | 0.3573 | 0.02831 | -0.004083 | 32 |
| median number of introns per gene | progenetic-miniature vs non-miniature | 6.33333 | 0.29989 | 0.34073 | 0.8801 | 0.3858 | 0.02517 | -0.007323 | 32 |
| median number of mid introns per gene | progenetic-miniature vs non-miniature | 6.333333 | 0.067018 | 0.335273 | 0.1999 | 0.8429 | 0.00133 | -0.03196 | 32 |
| proportion of compact introns (≤300 bp) per gene | progenetic-miniature vs non-miniature | 0.493354 | -0.028225 | 0.02936 | -0.9613 | 0.3441 | 0.02988 | -0.002453 | 32 |
| proportion of long introns (>300 bp) per gene | progenetic-miniature vs non-miniature | 0.506646 | 0.028225 | 0.02936 | 0.9613 | 0.3441 | 0.02988 | -0.002453 | 32 |
| % of non-repititive DNA | progenetic-miniature vs non-miniature | 63.9085 | -12.8731 | 6.9622 | -1.849 | 0.07433 | 0.1023 | 0.07238 | 32 |
| % of total repeat DNA | progenetic-miniature vs non-miniature | 36.0915 | 12.8731 | 6.9622 | 1.849 | 0.0743323 | 0.1023 | 0.07238 | 32 |
| % of class I repeats | progenetic-miniature vs non-miniature | 16.2367 | 5.3002 | 3.751 | 1.413 | 0.1679 | 0.0624 | 0.03115 | 32 |
| % of class II repeats | progenetic-miniature vs non-miniature | 13.9912 | 10.5068 | 4.8762 | 2.1547 | 0.039338 | 0.134 | 0.1052 | 32 |
| % of other repeats | progenetic-miniature vs non-miniature | 7.3656 | -3.3591 | 1.0058 | -3.3396 | 0.002254 | 0.271 | 0.2467 | 32 |
| % of retroelements | progenetic-miniature vs non-miniature | 16.2367 | 5.3002 | 3.751 | 1.413 | 0.1679 | 0.0624 | 0.03115 | 32 |
| % of total interspersed repeats | progenetic-miniature vs non-miniature | 31.5627 | 16.2805 | 7.1307 | 2.2832 | 0.029675 | 0.148 | 0.1196 | 32 |
| % of LINEs | progenetic-miniature vs non-miniature | 7.4508 | -1.9375 | 1.475 | -1.3136 | 0.199 | 0.05439 | 0.02287 | 32 |
| % of SINEs | progenetic-miniature vs non-miniature | 0.75112 | 0.3002 | 1.16281 | 0.2582 | 0.798 | 0.002217 | -0.03104 | 32 |
| % of LTR | progenetic-miniature vs non-miniature | 7.3962 | 7.5791 | 2.9843 | 2.5397 | 0.01651 | 0.177 | 0.1495 | 32 |
| % of DNA transposons | progenetic-miniature vs non-miniature | 13.9374 | 10.5549 | 4.8756 | 2.1648 | 0.038485 | 0.1351 | 0.1063 | 32 |
| % of helitrons | progenetic-miniature vs non-miniature | 0.057517 | -0.053438 | 0.014913 | -3.5834 | 0.001183 | 0.2997 | 0.2764 | 32 |
| % of small RNA | progenetic-miniature vs non-miniature | 0.162026 | -0.048213 | 0.07974 | -0.6046 | 0.54998 | 0.01204 | -0.02089 | 32 |
| % of satellites | progenetic-miniature vs non-miniature | -1.94E-06 | 3.90E-06 | 3.39E-06 | 1.1506 | 0.259 | 0.04226 | 0.01034 | 32 |
| % of simple repeats | progenetic-miniature vs non-miniature | 3.65306 | -2.84885 | 0.69891 | -4.0761 | 0.0003096 | 0.3564 | 0.335 | 32 |
| % of low complexity repeats | progenetic-miniature vs non-miniature | 0.322526 | -0.192931 | 0.063395 | -3.0433 | 0.004832 | 0.2359 | 0.2104 | 32 |
| % of unclassified repeats | progenetic-miniature vs non-miniature | 3.34673 | -0.40821 | 0.51089 | -0.799 | 0.4306 | 0.02084 | -0.0118 | 32 |
| % of recent (<5% K2P) class I repeats | progenetic-miniature vs non-miniature | 5.66959 | 0.59329 | 1.61455 | 0.3675 | 0.715852 | 0.004481 | -0.0287 | 32 |
| % of recent (<5% K2P) class II repeats | progenetic-miniature vs non-miniature | 1.9081 | 3.0015 | 1.5353 | 1.955 | 0.05996 | 0.113 | 0.08344 | 32 |
| % of recent (<5% K2P) other repeats | progenetic-miniature vs non-miniature | 0.19617 | 0.61828 | 0.55992 | 1.1042 | 0.2783 | 0.03906 | 0.007025 | 32 |
| % of recent (<5% K2P) LINEs | progenetic-miniature vs non-miniature | 3.50698 | -1.59005 | 0.73604 | -2.1603 | 0.03887 | 0.1346 | 0.1058 | 32 |
| % of recent (<5% K2P) SINEs | progenetic-miniature vs non-miniature | 0.086663 | 0.134859 | 0.173462 | 0.7775 | 0.443 | 0.01975 | -0.01292 | 32 |
| % of recent (<5% K2P) LTR | progenetic-miniature vs non-miniature | 1.9756 | 2.11 | 1.3497 | 1.5633 | 0.1285 | 0.07533 | 0.04451 | 32 |
| % of recent (<5% K2P) DNA transposons | progenetic-miniature vs non-miniature | 1.6809 | 2.9075 | 1.4318 | 2.0307 | 0.05123 | 0.1209 | 0.09155 | 32 |
| % of recent (<5% K2P) helitrons | progenetic-miniature vs non-miniature | 0.242729 | 0.089771 | 0.134502 | 0.6674 | 0.5096 | 0.01463 | -0.01821 | 32 |
| % of intermediate (5-20% K2P) class I repeats | progenetic-miniature vs non-miniature | 9.4273 | 2.9141 | 2.4334 | 1.1976 | 0.2404622 | 0.04562 | 0.01381 | 32 |
| % of intermediate (5-20% K2P) class II repeats | progenetic-miniature vs non-miniature | 7.4549 | 8.9667 | 3.9378 | 2.2771 | 0.03008 | 0.1474 | 0.1189 | 32 |
| % of intermediate (5-20% K2P) other repeats | progenetic-miniature vs non-miniature | 2.88117 | -1.51244 | 0.55555 | -2.7224 | 0.01069 | 0.1981 | 0.1714 | 32 |
| % of intermediate (5-20% K2P) LINEs | progenetic-miniature vs non-miniature | 4.49021 | -1.66486 | 0.91353 | -1.8224 | 0.07837 | 0.09967 | 0.06966 | 32 |
| % of intermediate (5-20% K2P) SINEs | progenetic-miniature vs non-miniature | 0.34258 | 0.37487 | 0.86132 | 0.4352 | 0.6665 | 0.006274 | -0.02685 | 32 |
| % of intermediate (5-20% K2P) LTR | progenetic-miniature vs non-miniature | 3.9955 | 4.7864 | 2.0328 | 2.3546 | 0.02528 | 0.156 | 0.1278 | 32 |
| % of intermediate (5-20% K2P) DNA transposons | progenetic-miniature vs non-miniature | 6.007 | 6.2083 | 3.0051 | 2.0659 | 0.04757 | 0.1245 | 0.09537 | 32 |
| % of intermediate (5-20% K2P) helitrons | progenetic-miniature vs non-miniature | 1.4023 | 2.7454 | 1.3089 | 2.0976 | 0.04447 | 0.1279 | 0.09883 | 32 |
| % of ancient (>20% K2P) class I repeats | progenetic-miniature vs non-miniature | 4.813 | 1.4553 | 1.2054 | 1.2073 | 0.2367607 | 0.04633 | 0.01454 | 32 |
| % of ancient (>20% K2P) class II repeats | progenetic-miniature vs non-miniature | 7.12106 | 0.19242 | 1.97792 | 0.0973 | 0.92315 | 0.0003154 | -0.03301 | 32 |
| % of ancient (>20% K2P) other repeats | progenetic-miniature vs non-miniature | 0.74272 | -0.22477 | 0.20375 | -1.1032 | 0.2787151 | 0.03899 | 0.006952 | 32 |
| % of ancient (>20% K2P) LINEs | progenetic-miniature vs non-miniature | 1.78139 | -0.42015 | 0.40631 | -1.0341 | 0.3094 | 0.03442 | 0.002231 | 32 |
| % of ancient (>20% K2P) SINEs | progenetic-miniature vs non-miniature | 0.251787 | 0.026861 | 0.362754 | 0.074 | 0.9415 | 0.0001827 | -0.03314 | 32 |
| % of ancient (>20% K2P) LTR | progenetic-miniature vs non-miniature | 2.7197 | 1.937 | 1.0832 | 1.7882 | 0.08385 | 0.09632 | 0.0662 | 32 |
| % of ancient (>20% K2P) DNA transposons | progenetic-miniature vs non-miniature | 5.30488 | 0.14979 | 1.54611 | 0.0969 | 0.92346 | 0.0003128 | -0.03301 | 32 |
| % of ancient (>20% K2P) helitrons | progenetic-miniature vs non-miniature | 1.7965 | 0.12188 | 0.72592 | 0.1679 | 0.86779 | 0.0009388 | -0.03236 | 32 |

**Supplementary Table 15.** Comparison of Brownian motion (BM) and Ornstein–Uhlenbeck (OU) model fit within the phylogenetic generalized least squares (PGLS) regression framework for transposable element content and gene architecture traits across Cypriniformes.

| | | Brownian motion model | | | Ornstein Uhlenbeck model | | | |
|---|---|---|---|---|---|---|---|---|
| **Trait** | **Predictor** | **BM_log likelihood** | **BM_AIC** | **BM_AICw** | **OU_log likelihood** | **OU_AIC** | **OU_AICw** | **delta AIC** |
| median gene length | miniature vs non-miniature | -299.7 | 605.4 | 0.09766831 | -296.5 | 601 | 0.90233169 | 4.44681 |
| overall median intron length | miniature vs non-miniature | -202.1 | 410.1 | 0.94325102 | -203.9 | 415.8 | 0.05674898 | 5.621389 |
| first (5`) median intron length | miniature vs non-miniature | -219.7 | 445.5 | 0.91786383 | -221.2 | 450.3 | 0.08213617 | 4.827341 |
| last (3`) median intron length | miniature vs non-miniature | -229.3 | 464.7 | 0.8139695 | -229.8 | 467.6 | 0.1860305 | 2.952024 |
| mid median intron length | miniature vs non-miniature | -195.3 | 396.6 | 0.95003862 | -197.2 | 402.5 | 0.04996138 | 5.890505 |
| single median intron length | miniature vs non-miniature | -249.7 | 505.3 | 0.0155038 | -244.5 | 497 | 0.9844962 | 8.302089 |
| ultra-short introns (≤100 bp) median length | miniature vs non-miniature | -58.2 | 122.4 | 0.92875084 | -59.76 | 127.53 | 0.07124916 | 5.135315 |
| short introns (101–300 bp) median length | miniature vs non-miniature | -70.15 | 146.29 | 0.91029618 | -71.46 | 150.93 | 0.08970382 | 4.634513 |
| compact introns (≤300 bp) median length | miniature vs non-miniature | -95.35 | 196.7 | 0.04782785 | -97.34 | 202.68 | 0.95217215 | 5.982275 |
| long introns (>300 bp) median length | miniature vs non-miniature | -221.6 | 449.3 | 0.93274375 | -223.3 | 454.6 | 0.06725625 | 5.259241 |
| first (5`) ultra-short introns (≤100 bp) median intron length | miniature vs non-miniature | -77.05 | 160.1 | 7.13E-05 | -66.5 | 141 | 1.00E+00 | 19.09733 |
| last (3`) ultra-short introns (≤100 bp) median intron length | miniature vs non-miniature | -63.4 | 132.8 | 0.92410055 | -64.9 | 137.8 | 0.07589945 | 4.998823 |
| mid ultra-short introns (≤100 bp) median intron length | miniature vs non-miniature | -58.09 | 122.17 | 0.92929131 | -59.66 | 127.33 | 0.07070869 | 5.151708 |
| single ultra-short introns (≤100 bp) median intron length | miniature vs non-miniature | -134.3 | 274.6 | 9.18E-12 | -107.9 | 223.8 | 1.00E+00 | 50.82785 |
| first (5`) short introns (101–300 bp) median intron length | miniature vs non-miniature | -112.2 | 230.3 | 0.00037479 | -103.3 | 214.5 | 0.99962521 | 15.77754 |
| last (3`) short introns (101–300 bp) median intron length | miniature vs non-miniature | -87.14 | 180.28 | 0.4867711 | -86.09 | 180.18 | 0.5132289 | 0.1058561 |
| mid short introns (101–300 bp) median intron length | miniature vs non-miniature | -67.99 | 141.99 | 0.9277965 | -69.55 | 147.09 | 0.0722035 | 5.106648 |
| single short introns (101–300 bp) median intron length | miniature vs non-miniature | -137.4 | 280.7 | 7.54E-06 | -124.6 | 257.1 | 1.00E+00 | 23.59124 |
| first (5`) compact introns (≤300 bp) median intron length | miniature vs non-miniature | -129.1 | 264.3 | 0.008993164 | -123.4 | 254.9 | 0.991006836 | 9.404513 |
| last (3`) compact introns (≤300 bp) median intron length | miniature vs non-miniature | -106 | 218 | 0.93441453 | -107.7 | 223.3 | 0.06558547 | 5.313132 |
| mid compact introns (≤300 bp) median intron length | miniature vs non-miniature | -94.18 | 194.36 | 0.95174567 | -96.16 | 200.32 | 0.04825433 | 5.963625 |
| single compact introns (≤300 bp) median intron length | miniature vs non-miniature | -144.9 | 295.7 | 7.09E-06 | -132 | 272 | 1.00E+00 | 23.71445 |
| first (5`) long introns (>300 bp) median intron length | miniature vs non-miniature | -229.5 | 465 | 0.91781899 | -230.9 | 469.8 | 0.08218101 | 4.826152 |
| last (3`) long introns (>300 bp) median intron length | miniature vs non-miniature | -230.3 | 466.6 | 0.92805717 | -231.8 | 471.7 | 0.07194283 | 5.114443 |
| mid long introns (>300 bp) median intron length | miniature vs non-miniature | -220.2 | 446.5 | 0.93300027 | -221.9 | 451.7 | 0.06699973 | 5.267434 |
| single long introns (>300 bp) median intron length | miniature vs non-miniature | -247.1 | 500.2 | 0.07861309 | -243.6 | 495.3 | 0.92138691 | 4.922684 |
| median number of introns per gene | miniature vs non-miniature | -47.84 | 101.69 | 1.73E-09 | -26.67 | 61.34 | 1.00E+00 | 40.35381 |
| median number of mid introns per gene | miniature vs non-miniature | -36.57 | 79.14 | 2.56698E-05 | -25 | 58 | 0.99997433 | 21.14034 |
| proportion of compact introns (≤300 bp) per gene | miniature vs non-miniature | 69.81 | -133.62 | 0.95169441 | 67.83 | -127.66 | 0.04830559 | 5.961394 |
| proportion of long introns (>300 bp) per gene | miniature vs non-miniature | 69.81 | -133.62 | 0.95169441 | 67.83 | -127.66 | 0.04830559 | 5.961394 |
| % of non-repititive DNA | miniature vs non-miniature | -115.5 | 236.9 | 0.7846175 | -115.8 | 239.5 | 0.2153825 | 2.585561 |
| % of total repeat DNA | miniature vs non-miniature | -115.5 | 236.9 | 0.7846175 | -115.8 | 239.5 | 0.2153825 | 2.585561 |
| % of class I repeats | miniature vs non-miniature | -109.6 | 225.3 | 0.003988913 | -103.1 | 214.2 | 0.996011087 | 11.04048 |
| % of class II repeats | miniature vs non-miniature | -106.9 | 219.8 | 0.50314 | -105.9 | 219.8 | 0.49686 | 0.0251203 |
| % of other repeats | miniature vs non-miniature | -50.17 | 106.33 | 0.91887745 | -51.59 | 111.19 | 0.08112255 | 4.854383 |
| % of retroelements | miniature vs non-miniature | -109.6 | 225.3 | 0.003988913 | -103.1 | 214.2 | 0.996011087 | 11.04048 |
| % of total interspersed repeats | miniature vs non-miniature | -115.7 | 237.3 | 0.8223273 | -116.2 | 240.4 | 0.1776727 | 3.06439 |
| % of LINEs | miniature vs non-miniature | -88.33 | 182.67 | 1.68E-06 | -74.04 | 156.07 | 1.00E+00 | 26.59604 |
| % of SINEs | miniature vs non-miniature | -54.57 | 115.14 | 0.8358171 | -55.2 | 118.4 | 0.1641829 | 3.254858 |
| % of LTR | miniature vs non-miniature | -99.41 | 204.83 | 0.07521554 | -95.91 | 199.81 | 0.92478446 | 5.018406 |
| % of DNA transposons | miniature vs non-miniature | -106.9 | 219.8 | 0.5034622 | -105.9 | 219.8 | 0.4965378 | 0.02769818 |
| % of helitrons | miniature vs non-miniature | 89.79 | -173.59 | 0.90293485 | 88.56 | -169.13 | 0.09706515 | 4.460536 |
| % of small RNA | miniature vs non-miniature | 17.09 | -28.18 | 0.0249112 | 21.75 | -35.51 | 0.9750888 | 7.334422 |
| % of satellites | miniature vs non-miniature | 280.1 | -554.2 | 0.90748221 | 278.8 | -549.7 | 0.09251779 | 4.566546 |
| % of simple repeats | miniature vs non-miniature | -36.96 | 79.93 | 0.94514209 | -38.81 | 85.62 | 0.05485791 | 5.693178 |
| % of low complexity repeats | miniature vs non-miniature | 43.37 | -80.75 | 0.9592773 | 41.22 | -74.43 | 0.0407227 | 6.318789 |
| % of unclassified repeats | miniature vs non-miniature | -30.76 | 67.52 | 0.7439525 | -30.82 | 69.65 | 0.2560475 | 2.133228 |
| % of recent (<5% K2P) class I repeats | miniature vs non-miniature | -83.99 | 173.98 | 0.001168581 | -76.24 | 160.48 | 0.998831419 | 13.50159 |
| % of recent (<5% K2P) class II repeats | miniature vs non-miniature | -72.8 | 151.6 | 0.2876433 | -70.9 | 149.8 | 0.7123567 | 1.813716 |
| % of recent (<5% K2P) other repeats | miniature vs non-miniature | -32.68 | 71.35 | 0.8926718 | -33.8 | 75.59 | 0.1073282 | 4.236655 |
| % of recent (<5% K2P) LINEs | miniature vs non-miniature | -54.52 | 115.04 | 0.01271682 | -49.17 | 106.33 | 0.98728318 | 8.704063 |
| % of recent (<5% K2P) SINEs | miniature vs non-miniature | 7.837 | -9.675 | 0.818014 | 7.334 | -6.669 | 0.181986 | 3.0059 |
| % of recent (<5% K2P) LTR | miniature vs non-miniature | -75.93 | 157.85 | 0.0157162 | -70.79 | 149.58 | 0.9842838 | 8.274444 |
| % of recent (<5% K2P) DNA transposons | miniature vs non-miniature | -71.43 | 148.87 | 0.2091536 | -69.1 | 146.2 | 0.7908464 | 2.66007 |
| % of recent (<5% K2P) helitrons | miniature vs non-miniature | 12.18 | -18.36 | 0.7247393 | 12.21 | -16.43 | 0.2752607 | 1.936187 |
| % of intermediate (5-20% K2P) class I repeats | miniature vs non-miniature | -100.6 | 207.2 | 0.000431174 | -91.83 | 191.67 | 0.999568827 | 15.49714 |
| % of intermediate (5-20% K2P) class II repeats | miniature vs non-miniature | -99.45 | 204.89 | 0.5621819 | -98.7 | 205.4 | 0.4378181 | 0.5000438 |
| % of intermediate (5-20% K2P) other repeats | miniature vs non-miniature | -29.53 | 65.07 | 0.95146393 | -31.51 | 71.02 | 0.04853607 | 5.951389 |
| % of intermediate (5-20% K2P) LINEs | miniature vs non-miniature | -72.19 | 150.38 | 4.56E-06 | -58.89 | 125.79 | 1.00E+00 | 24.59541 |
| % of intermediate (5-20% K2P) SINEs | miniature vs non-miniature | -43.27 | 92.53 | 0.8948223 | -44.41 | 96.81 | 0.1051777 | 4.281947 |
| % of intermediate (5-20% K2P) LTR | miniature vs non-miniature | -96.14 | 198.29 | 0.000300827 | -87.04 | 182.07 | 0.999699173 | 16.21735 |
| % of intermediate (5-20% K2P) DNA transposons | miniature vs non-miniature | -89.01 | 184.02 | 0.6565485 | -88.66 | 185.31 | 0.3434515 | 1.295902 |
| % of intermediate (5-20% K2P) helitrons | miniature vs non-miniature | -68.74 | 143.48 | 0.1526958 | -66.03 | 140.05 | 0.8473042 | 3.427225 |
| % of ancient (>20% K2P) class I repeats | miniature vs non-miniature | -70.94 | 147.87 | 0.004949424 | -64.63 | 137.27 | 0.995050576 | 10.60704 |
| % of ancient (>20% K2P) class II repeats | miniature vs non-miniature | -71.87 | 149.75 | 0.861369 | -72.7 | 153.4 | 0.138631 | 3.653414 |
| % of ancient (>20% K2P) other repeats | miniature vs non-miniature | -5.334 | 16.667 | 0.1598745 | -2.674 | 13.349 | 0.8401255 | 3.318324 |
| % of ancient (>20% K2P) LINEs | miniature vs non-miniature | -42.6 | 91.2 | 2.79081E-05 | -31.11 | 70.22 | 0.999972092 | 20.97313 |
| % of ancient (>20% K2P) SINEs | miniature vs non-miniature | -24.62 | 55.24 | 0.107492 | -21.5 | 51 | 0.892508 | 4.233239 |
| % of ancient (>20% K2P) LTR | miniature vs non-miniature | -62.05 | 130.09 | 0.1137902 | -58.99 | 125.99 | 0.8862098 | 4.105195 |
| % of ancient (>20% K2P) DNA transposons | miniature vs non-miniature | -63.18 | 132.36 | 0.8871803 | -64.24 | 136.48 | 0.1128197 | 4.124514 |
| % of ancient (>20% K2P) helitrons | miniature vs non-miniature | -47.85 | 101.7 | 0.1067453 | -44.73 | 97.45 | 0.8932547 | 4.248853 |
| median gene length | progenetic-miniature vs non-miniature | -291 | 588.1 | 0.1024631 | -287.9 | 583.7 | 0.8975369 | 4.340303 |
| overall median intron length | progenetic-miniature vs non-miniature | -196.4 | 398.8 | 0.94184815 | -198.2 | 404.3 | 0.05815185 | 5.569573 |
| first (5`) median intron length | progenetic-miniature vs non-miniature | -213.4 | 432.7 | 0.91673511 | -214.8 | 437.5 | 0.08326489 | 4.797583 |
| last (3`) median intron length | progenetic-miniature vs non-miniature | -222.6 | 451.2 | 0.8193098 | -223.1 | 454.2 | 0.1806902 | 3.023357 |
| mid median intron length | progenetic-miniature vs non-miniature | -189.8 | 385.6 | 0.94895375 | -191.7 | 391.4 | 0.05104625 | 5.845256 |
| single median intron length | progenetic-miniature vs non-miniature | -242 | 490.1 | 0.02861799 | -237.5 | 483 | 0.97138201 | 7.049368 |
| ultra-short introns (≤100 bp) median length | progenetic-miniature vs non-miniature | -47.51 | 101.03 | 0.93017909 | -49.1 | 106.2 | 0.06982091 | 5.178887 |
| short introns (101–300 bp) median length | progenetic-miniature vs non-miniature | -67.53 | 141.06 | 0.91001468 | -68.84 | 145.69 | 0.08998532 | 4.627628 |
| compact introns (≤300 bp) median length | progenetic-miniature vs non-miniature | -85.01 | 176.03 | 0.9594366 | -87.18 | 182.35 | 0.0405634 | 6.32696 |

| | | | | | | | | |
|---|---|---|---|---|---|---|---|---|
| long introns (>300 bp) median length | progenetic-miniature vs non-miniature | -215.4 | 436.7 | 0.93124272 | -217 | 441.9 | 0.06875728 | 5.211875 |
| first (5`) ultra-short introns (≤100 bp) median intron length | progenetic-miniature vs non-miniature | -73.56 | 153.12 | 5.44E-06 | -60.44 | 128.88 | 1.00E+00 | 24.24284 |
| last (3`) ultra-short introns (≤100 bp) median intron length | progenetic-miniature vs non-miniature | -51.43 | 108.87 | 0.92280647 | -52.91 | 113.83 | 0.07719353 | 4.962208 |
| mid ultra-short introns (≤100 bp) median intron length | progenetic-miniature vs non-miniature | -47.29 | 100.58 | 0.93115921 | -48.9 | 105.8 | 0.06884079 | 5.209268 |
| single ultra-short introns (≤100 bp) median intron length | progenetic-miniature vs non-miniature | -130.6 | 267.2 | 1.09E-11 | -104.4 | 216.7 | 1.00E+00 | 50.48064 |
| first (5`) short introns (101–300 bp) median intron length | progenetic-miniature vs non-miniature | -109 | 224 | 0.000553997 | -100.5 | 209 | 0.999446004 | 14.9956 |
| last (3`) short introns (101–300 bp) median intron length | progenetic-miniature vs non-miniature | -84.89 | 175.77 | 0.472501 | -83.78 | 175.55 | 0.527499 | 0.2202141 |
| mid short introns (101–300 bp) median intron length | progenetic-miniature vs non-miniature | -65.73 | 137.47 | 0.92710669 | -67.28 | 142.55 | 0.07289331 | 5.086144 |
| single short introns (101–300 bp) median intron length | progenetic-miniature vs non-miniature | -132.1 | 270.3 | 5.78E-07 | -116.8 | 241.6 | 1.00E+00 | 28.72754 |
| first (5`) compact introns (≤300 bp) median intron length | progenetic-miniature vs non-miniature | -123.9 | 253.7 | 0.02798835 | -119.3 | 246.6 | 0.97201165 | 7.095159 |
| last (3`) compact introns (≤300 bp) median intron length | progenetic-miniature vs non-miniature | -99.24 | 204.49 | 0.93538639 | -100.9 | 209.8 | 0.06461361 | 5.345069 |
| mid compact introns (≤300 bp) median intron length | progenetic-miniature vs non-miniature | -84.7 | 175.4 | 0.95799469 | -86.83 | 181.66 | 0.04200531 | 6.254092 |
| single compact introns (≤300 bp) median intron length | progenetic-miniature vs non-miniature | -139 | 284 | 8.47E-07 | -124 | 256.1 | 1.00E+00 | 27.96224 |
| first (5`) long introns (>300 bp) median intron length | progenetic-miniature vs non-miniature | -222.8 | 451.7 | 0.91679712 | -224.2 | 456.5 | 0.08320288 | 4.799209 |
| last (3`) long introns (>300 bp) median intron length | progenetic-miniature vs non-miniature | -223.6 | 453.3 | 0.92732428 | -225.2 | 458.4 | 0.07267572 | 5.092592 |
| mid long introns (>300 bp) median intron length | progenetic-miniature vs non-miniature | -214 | 434 | 0.93117813 | -215.6 | 439.2 | 0.06882187 | 5.209858 |
| single long introns (>300 bp) median intron length | progenetic-miniature vs non-miniature | -239.9 | 485.7 | 0.09495646 | -236.6 | 481.2 | 0.90504354 | 4.509129 |
| median number of introns per gene | progenetic-miniature vs non-miniature | -46.57 | 99.13 | 2.59E-09 | -25.79 | 59.59 | 1.00E+00 | 39.54329 |
| median number of mid introns per gene | progenetic-miniature vs non-miniature | -35.9 | 77.8 | 3.26E-05 | -24.57 | 57.14 | 1.00E+00 | 20.6641 |
| proportion of compact introns (≤300 bp) per gene | progenetic-miniature vs non-miniature | 67.35 | -128.7 | 0.95066866 | 65.39 | -122.79 | 0.04933134 | 5.917212 |
| proportion of long introns (>300 bp) per gene | progenetic-miniature vs non-miniature | 67.35 | -128.7 | 0.95066866 | 65.39 | -122.79 | 0.04933134 | 5.917212 |
| % of non-repititive DNA | progenetic-miniature vs non-miniature | -112 | 230 | 0.7581424 | -112.1 | 232.3 | 0.2418576 | 2.285045 |
| % of total repeat DNA | progenetic-miniature vs non-miniature | -112 | 230 | 0.7581424 | -112.1 | 232.3 | 0.2418576 | 2.285045 |
| % of class I repeats | progenetic-miniature vs non-miniature | -106.2 | 218.3 | 0.002958534 | -99.34 | 206.68 | 0.997041466 | 11.6402 |
| % of class II repeats | progenetic-miniature vs non-miniature | -104 | 214.1 | 0.4646398 | -102.9 | 213.8 | 0.5353602 | 0.283355 |
| % of other repeats | progenetic-miniature vs non-miniature | -46.62 | 99.25 | 0.91943902 | -48.06 | 104.12 | 0.08056098 | 4.869499 |
| % of retroelements | progenetic-miniature vs non-miniature | -106.2 | 218.3 | 0.002958534 | -99.34 | 206.68 | 0.997041466 | 11.6402 |
| % of total interspersed repeats | progenetic-miniature vs non-miniature | -112 | 230 | 0.8024675 | -112.4 | 232.8 | 0.1975325 | 2.803576 |
| % of LINEs | progenetic-miniature vs non-miniature | -86.03 | 178.06 | 2.09E-06 | -71.96 | 151.91 | 1.00E+00 | 26.15255 |
| % of SINEs | progenetic-miniature vs non-miniature | -53.32 | 112.64 | 0.8341534 | -53.94 | 115.88 | 0.1658466 | 3.230708 |
| % of LTR | progenetic-miniature vs non-miniature | -95.39 | 196.78 | 0.02765023 | -90.83 | 189.66 | 0.97234977 | 7.120163 |
| % of DNA transposons | progenetic-miniature vs non-miniature | -104 | 214.1 | 0.4647575 | -102.9 | 213.8 | 0.5352425 | 0.2824083 |
| % of helitrons | progenetic-miniature vs non-miniature | 87.93 | -169.87 | 0.91362301 | 86.57 | -165.15 | 0.08637699 | 4.717393 |
| % of small RNA | progenetic-miniature vs non-miniature | 16.14 | -26.29 | 0.03003646 | 20.62 | -33.24 | 0.96996354 | 6.949693 |
| % of satellites | progenetic-miniature vs non-miniature | 357.4 | -708.9 | 0.94785615 | 355.5 | -703.1 | 0.05214385 | 5.800393 |
| % of simple repeats | progenetic-miniature vs non-miniature | -34.14 | 74.27 | 0.94847113 | -36.05 | 80.1 | 0.05152887 | 5.825418 |
| % of low complexity repeats | progenetic-miniature vs non-miniature | 42.91 | -79.82 | 0.96006961 | 40.73 | -73.46 | 0.03993039 | 6.359736 |
| % of unclassified repeats | progenetic-miniature vs non-miniature | -29.4 | 64.8 | 0.6843999 | -29.17 | 66.34 | 0.3156001 | 1.548133 |
| % of recent (<5% K2P) class I repeats | progenetic-miniature vs non-miniature | -81.46 | 168.92 | 0.000747826 | -73.26 | 154.52 | 0.999252174 | 14.39519 |
| % of recent (<5% K2P) class II repeats | progenetic-miniature vs non-miniature | -70.92 | 147.84 | 0.1523485 | -68.2 | 144.4 | 0.8476515 | 3.432598 |
| % of recent (<5% K2P) other repeats | progenetic-miniature vs non-miniature | -27.29 | 60.58 | 0.93885055 | -29.02 | 66.04 | 0.06114945 | 5.462671 |
| % of recent (<5% K2P) LINEs | progenetic-miniature vs non-miniature | -52.84 | 111.69 | 0.0116678 | -47.4 | 102.8 | 0.9883322 | 8.878373 |
| % of recent (<5% K2P) SINEs | progenetic-miniature vs non-miniature | 7.501 | -9.001 | 0.8305054 | 6.911 | -5.823 | 0.1694946 | 3.178426 |
| % of recent (<5% K2P) LTR | progenetic-miniature vs non-miniature | -72.72 | 151.43 | 0.004685838 | -66.36 | 140.71 | 0.995314162 | 10.71703 |
| % of recent (<5% K2P) DNA transposons | progenetic-miniature vs non-miniature | -69.62 | 145.24 | 0.1055837 | -66.49 | 140.97 | 0.8944163 | 4.273336 |
| % of recent (<5% K2P) helitrons | progenetic-miniature vs non-miniature | 11.97 | -17.93 | 0.5583498 | 12.73 | -17.46 | 0.4416502 | 0.468935 |
| % of intermediate (5-20% K2P) class I repeats | progenetic-miniature vs non-miniature | -97.67 | 201.34 | 0.000368204 | -88.77 | 185.53 | 0.999631796 | 15.81301 |
| % of intermediate (5-20% K2P) class II repeats | progenetic-miniature vs non-miniature | -96.72 | 199.44 | 0.5151211 | -95.78 | 199.56 | 0.4848789 | 0.1210056 |
| % of intermediate (5-20% K2P) other repeats | progenetic-miniature vs non-miniature | -26.82 | 59.65 | 0.94707365 | -28.71 | 65.41 | 0.05292635 | 5.768951 |
| % of intermediate (5-20% K2P) LINEs | progenetic-miniature vs non-miniature | -70.37 | 146.74 | 4.88E-06 | -57.14 | 122.28 | 1.00E+00 | 24.46034 |
| % of intermediate (5-20% K2P) SINEs | progenetic-miniature vs non-miniature | -42.26 | 90.53 | 0.8967545 | -43.43 | 94.85 | 0.1032455 | 4.323344 |
| % of intermediate (5-20% K2P) LTR | progenetic-miniature vs non-miniature | -93.2 | 192.4 | 0.000102334 | -83.01 | 174.02 | 0.999897667 | 18.37434 |
| % of intermediate (5-20% K2P) DNA transposons | progenetic-miniature vs non-miniature | -86.72 | 179.44 | 0.6379921 | -86.29 | 180.58 | 0.3620079 | 1.133319 |
| % of intermediate (5-20% K2P) helitrons | progenetic-miniature vs non-miniature | -66.64 | 139.28 | 0.08574702 | -63.27 | 134.55 | 0.91425298 | 4.733412 |
| % of ancient (>20% K2P) class I repeats | progenetic-miniature vs non-miniature | -69.19 | 144.37 | 0.00580287 | -63.04 | 134.09 | 0.99419713 | 10.28717 |
| % of ancient (>20% K2P) class II repeats | progenetic-miniature vs non-miniature | -69.95 | 145.91 | 0.8495341 | -70.69 | 149.37 | 0.1504659 | 3.461903 |
| % of ancient (>20% K2P) other repeats | progenetic-miniature vs non-miniature | -5.607 | 17.214 | 0.1432287 | -2.818 | 13.637 | 0.8567713 | 3.577457 |
| % of ancient (>20% K2P) LINEs | progenetic-miniature vs non-miniature | -41.61 | 89.22 | 2.95608E-05 | -30.18 | 68.36 | 0.999970439 | 20.85806 |
| % of ancient (>20% K2P) SINEs | progenetic-miniature vs non-miniature | -24.31 | 54.62 | 0.1071604 | -21.19 | 50.38 | 0.8928396 | 4.240161 |
| % of ancient (>20% K2P) LTR | progenetic-miniature vs non-miniature | -60.39 | 126.79 | 0.1021285 | -57.22 | 122.44 | 0.8978715 | 4.34759 |
| % of ancient (>20% K2P) DNA transposons | progenetic-miniature vs non-miniature | -61.43 | 128.86 | 0.878801 | -62.41 | 132.82 | 0.121199 | 3.962249 |
| % of ancient (>20% K2P) helitrons | progenetic-miniature vs non-miniature | -46.84 | 99.68 | 0.1042534 | -43.69 | 95.38 | 0.8957466 | 4.301667 |
